# Comparative analysis of amplicon and metagenomic sequencing methods reveals key features in the evolution of animal metaorganisms

**DOI:** 10.1101/604314

**Authors:** Philipp Rausch, Malte Rühlemann, Britt Hermes, Shauni Doms, Tal Dagan, Katja Dierking, Hanna Domin, Sebastian Fraune, Jakob von Frieling, Ute Henschel Humeida, Femke-Anouska Heinsen, Marc Höppner, Martin Jahn, Cornelia Jaspers, Kohar Annie B. Kissoyan, Daniela Langfeldt, Ateeqr Rehman, Thorsten B. H. Reusch, Thomas Röder, Ruth A. Schmitz, Hinrich Schulenburg, Ryszard Soluch, Felix Sommer, Eva Stukenbrock, Nancy Weiland-Bräuer, Philip Rosenstiel, Andre Franke, Thomas Bosch, John F. Baines

**Author notes:** Authors contributed equally (Philipp Rausch, Malte Rühlemann). Corresponding authors: Philipp Rausch, John F. Baines. Email addresses: John F. Baines-, Thomas Bosch-, Tal Dagan-, Katja Dierking-, Hanna Domin-, Shauni Doms-, Andre Franke-, Sebastian Fraune-, Jakob von Frieling-, Femke-Anouska Heinsen-, Ute Henschel Humeida-, Britt Marie Hermes-, Marc Höppner-, Martin Jahn-, Cornelia Jaspers-, Kohar Annie B. Kissoyan-, Daniela Langfeldt-, Philipp Rausch-, Ateeqr Rehman-, Thorsten B. H. Reusch-, Thomas Röder-, Philip Rosenstiel-, Malte Rühlemann-, Ruth A. Schmitz-, Hinrich Schulenburg-, Ryszard Soluch-, Felix Sommer-, Eva Stukenbrock-, Nancy Weiland-Bräuer.

## Abstract

**Background:** The interplay between hosts and their associated microbiome is now recognized as a fundamental basis of the ecology, evolution and development of both players. These interdependencies inspired a new view of multicellular organisms as “metaorganisms”. The goal of the Collaborative Research Center “Origin and Function of Metaorganisms” is to understand why and how microbial communities form long-term associations with hosts from diverse taxonomic groups, ranging from sponges to humans in addition to plants.

**Methods:** In order to optimize the choice of analysis procedures, which may differ according to the host organism and question at hand, we systematically compared the two main technical approaches for profiling microbial communities, 16S rRNA gene amplicon- and metagenomic shotgun sequencing across our panel of ten host taxa. This includes two commonly used 16S rRNA gene regions and two amplification procedures, thus totaling five different microbial profiles per host sample.

**Conclusion:** While 16S rRNA gene-based analyses are subject to much skepticism, we demonstrate that many aspects of bacterial community characterization are consistent across methods and that metagenomic shotgun results are largely dependent on the employed pipeline. The resulting insight facilitates the selection of appropriate methods across a wide range of host taxa. Finally, by contrasting taxonomic and functional profiles and performing phylogenetic analysis, we provide important and novel insight into broad evolutionary patterns among metaorganisms, whereby the transition of animals from an aquatic to a terrestrial habitat marks a major event in the evolution of host-associated microbial composition.

## Background

Dynamic host-microbe interactions have shaped the evolution of life. Virtually all plants and animals are colonized by an interdependent complex of microorganisms, and there is growing recognition that the biological processes of hosts and their associated microbial communities function in tandem, often as biological partners comprising a collective entity known as the metaorganism [1]. For instance, symbiotic bacteria contribute to host health and development in critical ways, ranging from nutrient metabolism to regulating whole life cycles [2] and in turn benefit from habitats and resources the host provides. Moreover, it is well established that perturbations of the microbiome likely play an important role in many host disease states [3]. However, researchers have yet to elucidate the mechanisms driving these interactions, as the exact molecular and cellular processes are only poorly understood.

An integrated view on the metaorganism encompasses a cross-disciplinary approach that addresses how and why microbial communities form long-term associations with their hosts. Despite widespread agreement that the interdependencies of microbes and their hosts warrant elucidation, there remains considerable incongruity between researchers regarding the best methodologies to study host-microbe interactions. The development of standardized protocols for characterizing and analyzing host-associated microbiomes across the breadth of the tree of life are thus crucial to understand the evolution and function of metaorganisms without the issues of technical inconsistencies or data quality.

Rapidly growing interest in microbiome research has been bolstered by the ability to profile diverse microbial communities using next-generation sequencing (NGS). This culture-free, high-throughput technology enables identification and comparison of entire microbial communities [4]. Metagenomics typically encompasses two particular sequencing strategies: amplicon sequencing, most often of the 16S rRNA gene as a phylogenetic marker, or shotgun sequencing, which captures the complete breadth of DNA within a sample [4].

The use of the 16S ribosomal RNA gene as a phylogenetic marker has proven to be an efficient and cost-effective strategy for microbiome analysis, and even allows for the imputation of functional content based on taxon abundances [5]. However, PCR-based phylogenetic marker protocols are vulnerable to biases through sample preparation and sequencing errors, in particular the choice of which hypervariable regions of the 16S rRNA gene targeted seem to be among the biggest factors underlying technical differences in microbiome composition [6–8]. Furthermore, 16S rRNA gene amplicon sequencing is typically limited to taxonomic classification at the genus-level depending on the database and classifiers used [9], and provides only limited functional information [5]. These well-recognized limitations of amplicon-based microbial community analyses have raised concerns about the accuracy and reproducibility of 16S rRNA phylogenetic marker studies and have led to an increased interest in developing more reliable methods for amplicon library preparation and sequencing [8, 10].

Shotgun metagenomics, on the other hand, offers the advantage of species- and strain-level classification of bacteria. Additionally, it allows researchers to examine the functional relationships between hosts and bacteria by determining the functional content of samples directly [9, 11], and enables the exploration of yet unknown microbial life that would otherwise remain unclassifiable [12]. However, the relatively high costs of shotgun metagenomics and more demanding bioinformatic requirements have precluded its use for microbiome analysis on a wide scale [4, 9].

In this study, we set out to systematically compare experimental and analytical aspects of the two main technical approaches for profiling microbial communities, 16S rRNA gene amplicon- and shotgun sequencing, across a diverse array of host species studied in the Collaborative Research Center 1182, “Origin and Function of Metaorganisms”. The ten host species range from basal aquatic metazoans [*Aplysina aerophoba* (sponge) and *Mnemiopsis leidyi* (comb jelly)], to marine and limnic cnidarians (*Aurelia aurita, Nematostella vectensis*, *Hydra vulgaris)*, standard vertebrate (*Mus musculus*) and invertebrate model organisms (*Drosophila melanogaster*, *Caenorhabditis elegans*), to *Homo sapiens*, in addition to wheat (*Triticum aestivum*) and a standardized mock community. This setup provides a breadth of samples in terms of taxonomic composition and diversity. Conducting standardized data generation procedures on these diverse samples on the one hand provides a unique and powerful opportunity to systematically compare alternative methods, which display considerable heterogeneity in performance. On the other hand, this information enables researchers working on these or similar host species to choose the experimental (*e.g*. hypervariable region) or analytical pipelines that best suit their needs, which will be a valuable resource to the greater community of host-microbe researchers. Finally, we identified a number of interesting, broad scale patterns contrasting the aquatic and terrestrial environment of metaorganisms, which also reflect their evolutionary trajectories.

## Results

Our panel of hosts includes ten species, for which five biological replicates each were included (see Figure S1). The majority of hosts are metazoans, including the “gold sponge” (*Aplysina aerophoba)*, moon jellyfish (*Aurelia aurita*), comb jellyfish (*Mnemiopsis leidyi*), starlet sea anemone (*Nematostella vectensis)*, fresh-water polyp *Hydra vulgaris*, roundworm (*Ceanorhabditis elegans*), fruit fly (*Drosophila melanogaster*), mouse (*Mus musculus*), human (*Homo sapiens*), as well as the inclusion of wheat (*Triticum aestivum*), which can serve as an outgroup to the metazoan taxa. *Drosophila melanogaster* was additionally sampled using two different methods targeting feces and intestinal tissue. Nucleic acid extraction procedures were conducted according to the needs of the individual host species (see Methods and Supplementary Material), after which all DNA templates were subjected to a standard panel of sequencing procedures. For 16S rRNA gene amplicon sequencing we used primers flanking two commonly used variable regions, the V1V2 and V3V4 regions. Further, for each region we compared a single-step fusion-primer PCR to a two-step procedure designed to improve the accuracy of amplicon-based studies [8]. Finally, all samples were also subjected to shotgun sequencing, such that five different sequence profiles were generated for each sample. While a single classification pipeline was employed for all four 16S rRNA gene amplicon sequence profiles, community composition based on shotgun data was initially evaluated using five different classification methods (Kraken [13], MEGAN [14], MetaPhlan [15], MetaPhlan2 [16], and SortmeRNA [17]; see Supplementary Material for comparative descriptions). However, due to the advantage of simultaneously performing taxonomical and functional classification of shotgun reads, as well as overall good performance (see analyses of mock community below), MEGAN was used as a representative pipeline for most subsequent analyses.

### Performance of data processing and quality control

All data generated from amplicons were subject to the same stringent quality control pipeline including read-trimming, merging of forward and reverse reads, quality filtering based on sequence quality and estimated errors, and chimera removal (see Methods). The one step V1V2 amplicon data showed the highest rate of read-survival (62.13 ± 23.90%, mean ± s.d.) followed by the corresponding two step method (mean= 49.85 ± 23.90%, mean ± s.d.), in large part due to the greater coverage of this comparatively shorter amplicon (~312 bp). In contrast, 42.02 ± 16.41% and 36.88 ± 23.89% of the total reads were included in downstream analysis for the one step and two step V3V4 data, respectively. The longer V3V4 amplicon (~470 bp) was more affected by drops in quality at the end of the reads, which decreases the overlap of forward and reverse reads and thus increases the chances of sequencing errors (Figure S2, for final sample sizes see Table S1). Overall, aside from chimera removal, each quality control step resulted in a comparatively greater loss of V3V4-compared to V1V2 data. On the other hand, the V3V4 one step method yields the lowest number of chimeras, suggesting a lower rate of chimera formation- and/or detection in this approach (variable region- *F*_1,214_=3.8881, *P*=0.0499, PCR- *F*_1,214_=8.1751, *P*=0.0047, variable region×PCR- *F*_1,214_=6.4733, *P*=0.0117; Linear Mixed Model with organism as random factor). Among all host taxa we observe the highest proportion of retained reads in the V1V2 one step method and the lowest in the V3V4 two step method (Figure S2B; variable region- *F*_1,215_=74.9989, *P*<0.0001, PCR- *F*_1,215_=21.0743, *P*<0.0001; Linear Mixed Model with organism as random factor). After quality filtering and the identification of bacterial reads, an average of 0.46 Gb of shotgun reads per sample was achieved (range 0.03 to 2.1 Gb) (Figure S3A, for final sample sizes see Table S1). To provide an initial assessment and comparison between the amplicon and shotgun-based techniques, we plotted the discovered classifiable taxa and functions for the entire pooled dataset. Although the methods differ distinctly, each method shows a plateau in the number of discovered entities (see Figure S3C, S3D).

### Mock community

The analysis of standardized mock communities is an important measure to ensure general quality standards in microbial community analysis. In this study we employed a commercially available mixture of eight bacterial- and two yeast species. Comparison among the amplification procedures (one- and two step PCR), 16S rRNA gene regions (V1V2, V3V4) and shotgun data reveals varying degrees of similarity to the expected microbial community composition (Figure 1). One discrepancy is apparent due to the misclassification of *Escherichia/Shigella*, whose close relationship make delineation at the genus level difficult based on the V1V2 region are subsequently classified to *Enterobacteriaceae* (Figure 1A, Figure S4). Classification of this bacterial group also differs according to shotgun pipeline employed, due to different naming and taxonomic standards of the respective databases (*Escherichia*, *Shigella*, *Enterobacteriaceae* refer to the *Escherichia/ Shigella* cluster) [18]. However, overall the amplicon-based profiles show the closest matches to the expected community. The V1V2 one step method and Kraken show the lowest degree of deviation between observed and expected abundances of the focus taxa (Table 1, Figure S4). However, Kraken falsely detects a large number of taxa not present in the mock communities. In addition, the relative abundances of fungi in the mock community were relatively well predicted by MEGAN and Kraken, while MetaPhlan2 failed to identify *Cryptococcus* and replaced it with several other taxa (see Figure 1).

**Table 1:** Differences between expected and observed genus abundances in the mock communities (N_shotgun_=4, N_amplicon_=3) via a one-sample *t*-test (two-sided) of relative abundances (*P*-values are adjusted via Hommel procedure).

| Members mock community | shotgun |  |  |  |  | amplicon |  |  |  |
| --- | --- | --- | --- | --- | --- | --- | --- | --- | --- |
|  | MEGAN | Kraken | MetaPhlan | MetaPhlan2 | SortmeRNA | V1V2 one step | V3V4 one step | V1V2 two step | V3V4 two step |
| *Staphylococcus* | 0.00002 | 0.14039 | 0.07916 | 0.07010 | $1.1097 \times 10^{-6}$ | 0.52446 | 0.09200 | 0.03994 | 0.21564 |
| *Listeria* | 0.00395 | 0.06065 | 0.02306 | 0.06043 | $1.4751 \times 10^{-6}$ | 0.34964 | 0.53267 | 0.03003 | 0.00545 |
| *Bacillus* | 0.00006 | 0.09558 | 0.02219 | 0.03638 | $3.9824 \times 10^{-7}$ | 0.21420 | 0.02818 | 0.29671 | 0.30589 |
| *Pseudomonas* | 0.13668 | 0.40989 | 0.62649 | 0.46933 | $2.7877 \times 10^{-7}$ | 0.36721 | 0.05776 | 0.38147 | 0.59037 |
| *Escherichia/Shigella* * | NA | NA | NA | NA | $9.9378 \times 10^{-10}$ | 0.00462 | 0.45612 | 0.00237 | 0.59037 |
| *Shigella* * | $4.6372 \times 10^{-10}$ | NA | $8.0806 \times 10^{-8}$ | NA | NA | NA | NA | NA | NA |
| *Escherichia* * | 0.00001 | 0.00882 | 0.00710 | 0.28178 | NA | NA | NA | NA | NA |
| *Enterobacteriaceae* * | NA | NA | NA | NA | NA | 0.87898 | 0.00004 | 0.19274 | 0.00055 |
| *Salmonella* | $3.8092 \times 10^{-6}$ | 0.08772 | 0.02203 | 0.03361 | $4.9348 \times 10^{-7}$ | 0.34964 | 0.05838 | 0.09712 | 0.08851 |
| *Lactobacillus* | 0.00297 | 0.09704 | 0.05384 | 0.04043 | $1.4751 \times 10^{-6}$ | 0.87898 | 0.53267 | 0.38147 | 0.59037 |
| *Enterococcus* | 0.00012 | 0.18719 | 0.00353 | 0.07277 | $6.3719 \times 10^{-7}$ | 0.04816 | 0.03746 | 0.01159 | 0.00954 |

\* *Escherichia/Shigella* relatives counted as equivalent

**Figure 1:**
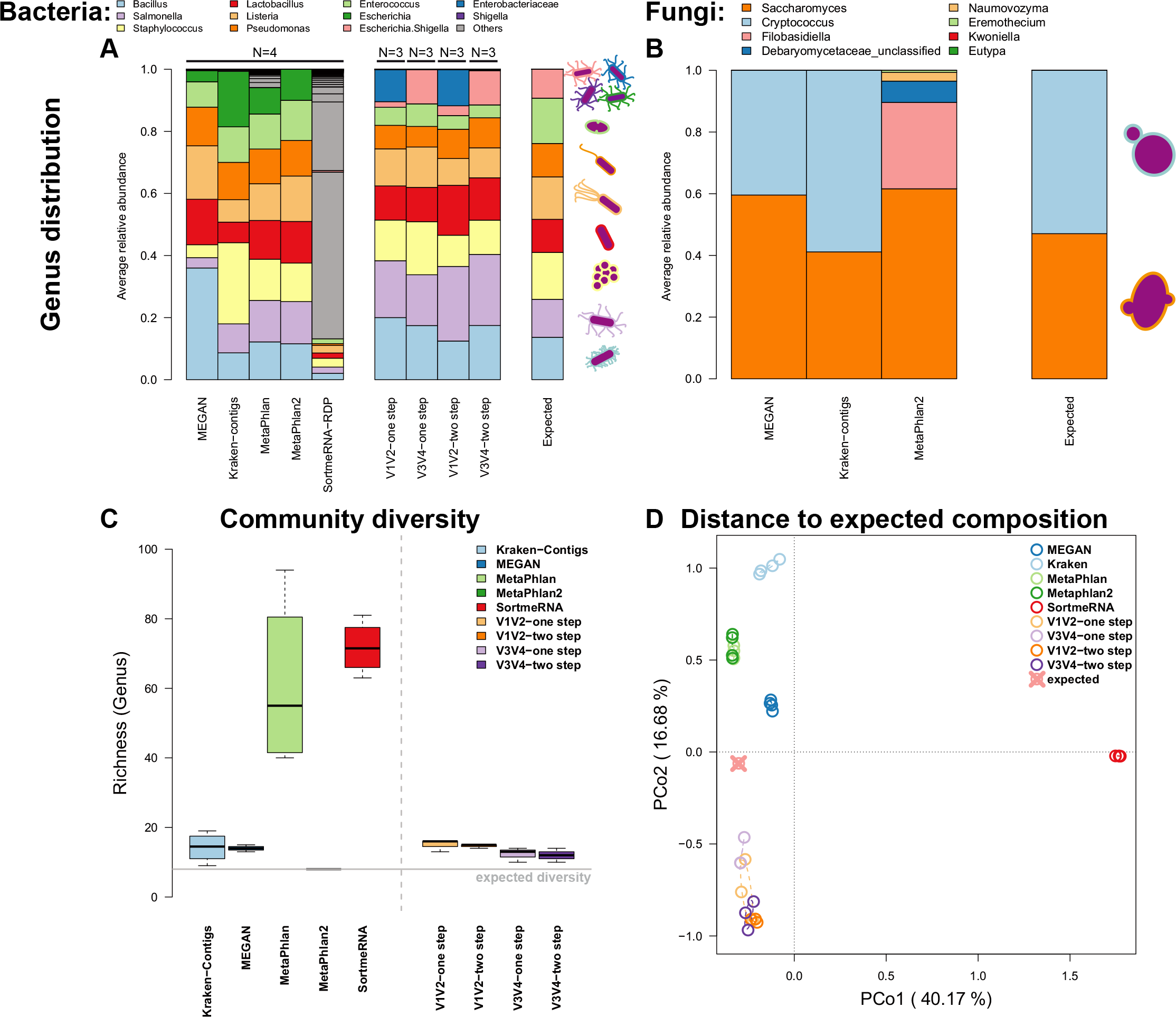
Average community composition of bacteria (A) and fungi (B) in the mock community samples sequenced via metagenomic shotgun- and 16S rRNA gene amplicon techniques (amplicon: V1V2, V3V4, one step, two step; shotgun: MEGAN based classification (short reads), MetaPhlan (short reads), MetaPhlan2 (short reads), Kraken based classification (contigs), SortmeRNA (short reads)). (C) Bacterial genus-level alpha diversity estimates in comparison to the expected community value. (D) Principle coordinate analysis of the Bray-Curtis distance between methods and the expected community. Ellipses represent standard deviations of points within the respective groups. Sample sizes for the different approaches are N_shotgun_=4, N_V1V2-one step_=3, N_V1V2-two step_=3, N_V3V4-one step_=3, and N_V1V2-two step_=3.

Next, we evaluated alpha and beta diversity across the different technical and analytical methods. Interestingly, most methods overestimate taxon richness but underestimate complexity (as measured by the Shannon index) of the mock community, which could reflect biases arising from grouping taxon abundances together (Figure 1, Figure S4, Figure S5, Table S2). Overall the amplicon methods appear to more accurately reflect alpha diversity, although significant differences are present with regard to the amplified region (species richness: variable region- *F*_1,10_=6.3657, *P*=0.0302; Shannon H: method- *F*_1,9_=3.330, *P*=0.1014, variable region- *F*_1,9_=6.110, *P*=0.0354). With regard to beta diversity, the largest distance to the expected composition is observed in SortmeRNA applied to shotgun sequencing of the mock community, while the amplicon-based techniques, MEGAN, and MetaPhlan2 show the lowest distance (Figure 1D, Figure S5, Table S3). Pairwise tests show almost no differences between the amplicon-based techniques, while all shotgun based methods significantly differ from each other (Table S4). Thus, in conclusion shotgun-based analysis pipelines yield a higher degree of variability/error compared to the amplicon-based approaches based on a simple mock community. For subsequent analyses we thus mainly focus on the amplicon-based data and MEGAN as a representative shotgun-based pipeline, for which eukaryotic (*e.g*. fungal) sequences were not included in the following analyses.

### Taxonomic diversity within and between hosts

To evaluate the performance of our panel of metagenomic methods over the range of complex host-associated communities in our consortium, we next employed a panel of alpha- and beta diversity analyses to these samples, which also provides an opportunity to infer broad patterns across animal taxa based on a standardized methodology. Measures of alpha diversity display overall consistent values with respect to host species, although many significant differences between technical methods are present, mostly in a host-specific manner (Figure 2A-B). However, several host taxa display high levels of consistency across methods including *A. aurita*, *C. elegans*, *D. melanogaster* and *H. sapiens*, which show almost no significant differences between methods. Discrepancies and individual recommendations for each host species are discussed in the Supplementary Material (see Figures S6-S16). An intriguing observation is the tendency of aquatic hosts to display higher alpha diversity values than those of terrestrial hosts, which is supported by average differences between aquatic and terrestrial hosts and by relative consistent comparisons among single host species as well (Figure 2C-D, Table S5). Finally, we also compared alpha diversity estimates based on the other shotgun-based classifiers, which in most cases display greater heterogeneity than among the 16S rRNA gene amplicon and MEGAN based estimates alone, but still recover similar trends (Figure S17).

**Figure 2:**
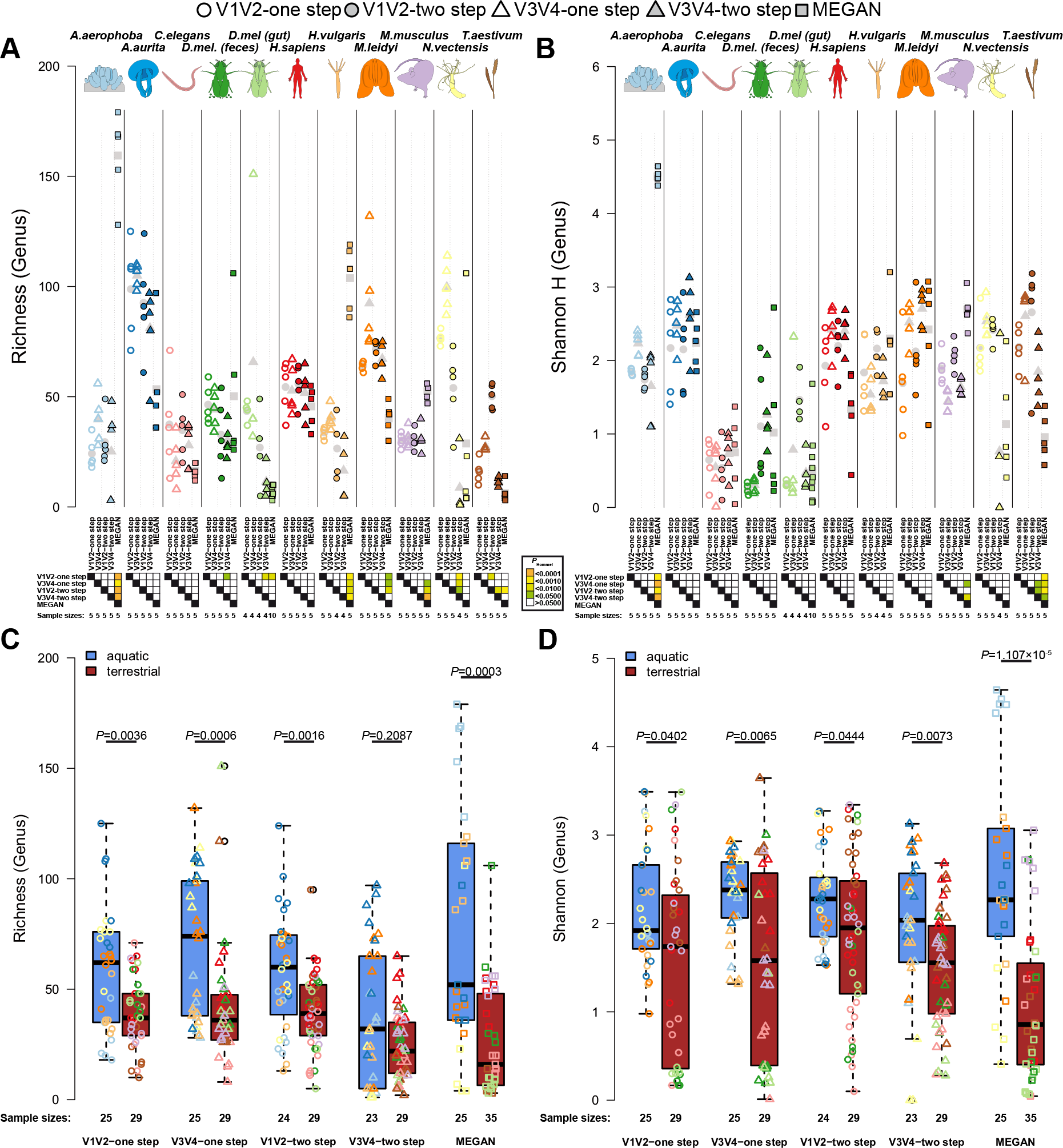
Comparison of bacterial genus richness (A) and Shannon H (B) based on 16S rRNA gene amplicon and shotgun derived genus profiles based on MEGAN highlighting the differences between variable regions, amplification methods, and metagenomic classifier, as well as between the different host organisms. Colors show significance of amplification methods (A, C) or pairwise comparisons of methods (B, D) based on pairwise *t*-tests with Hommel *P*-value adjustment (A, B), and approximate Wilcoxon test for the comparison between environmental categories (C, D). Mean values are shown in grey symbols in plots A and B. Sample sizes are indicated below the samples.

In order to investigate broad patterns of bacterial community similarity according to metagenomic procedure and host species, we performed beta diversity analyses including all host samples and each of their five different methodological profiles. This analysis reveals an overall strong signal of host species, irrespective of the method used to generate community profiles (Table 2, Figure 3). Pairwise comparisons between hosts are significant in all cases except for samples derived from the V3V4 two step protocol, which did not consistently reach significance after correction for multiple testing (Table S6). Further, complementary to the observations made for alpha diversity, we also find strong signals of community differentiation between the aquatic and terrestrial hosts (Table 2, Figure 3B and D). The separation between these environments appears to be stronger based on amplicon data, whereas the separation between hosts is stronger based on shotgun derived data (Table 2). Clustering of communities based on host environment is consistent irrespective of the underlying shotgun analysis method, although the topologies vary strongly (*e.g*. MetaPhlan2, see Figure S18). To further evaluate the variability among biological replicates, we evaluated intra-group distances according to host species, which reveals organisms with generally higher community variability (*i.e. C. elegans*, *A. aurita, H. sapiens*, *H. vulgaris, T. aestivum*, and *M. leidyi*) than other host organisms in our study (*N. vectensis*, *M. musculus, D. melanogaster*, *and A. aerophoba*; Figure S19A, C). Interestingly, intra-group distances also significantly differ between the aquatic and terrestrial environments, whereby aquatic organisms tend to display less variable communities than terrestrial ones (Figure S19B, D). The low performance of *T. aestivum* in subsequent analyses possibly originates from its commercial origin and low bacterial biomass relative to host material.

**Table 2:**
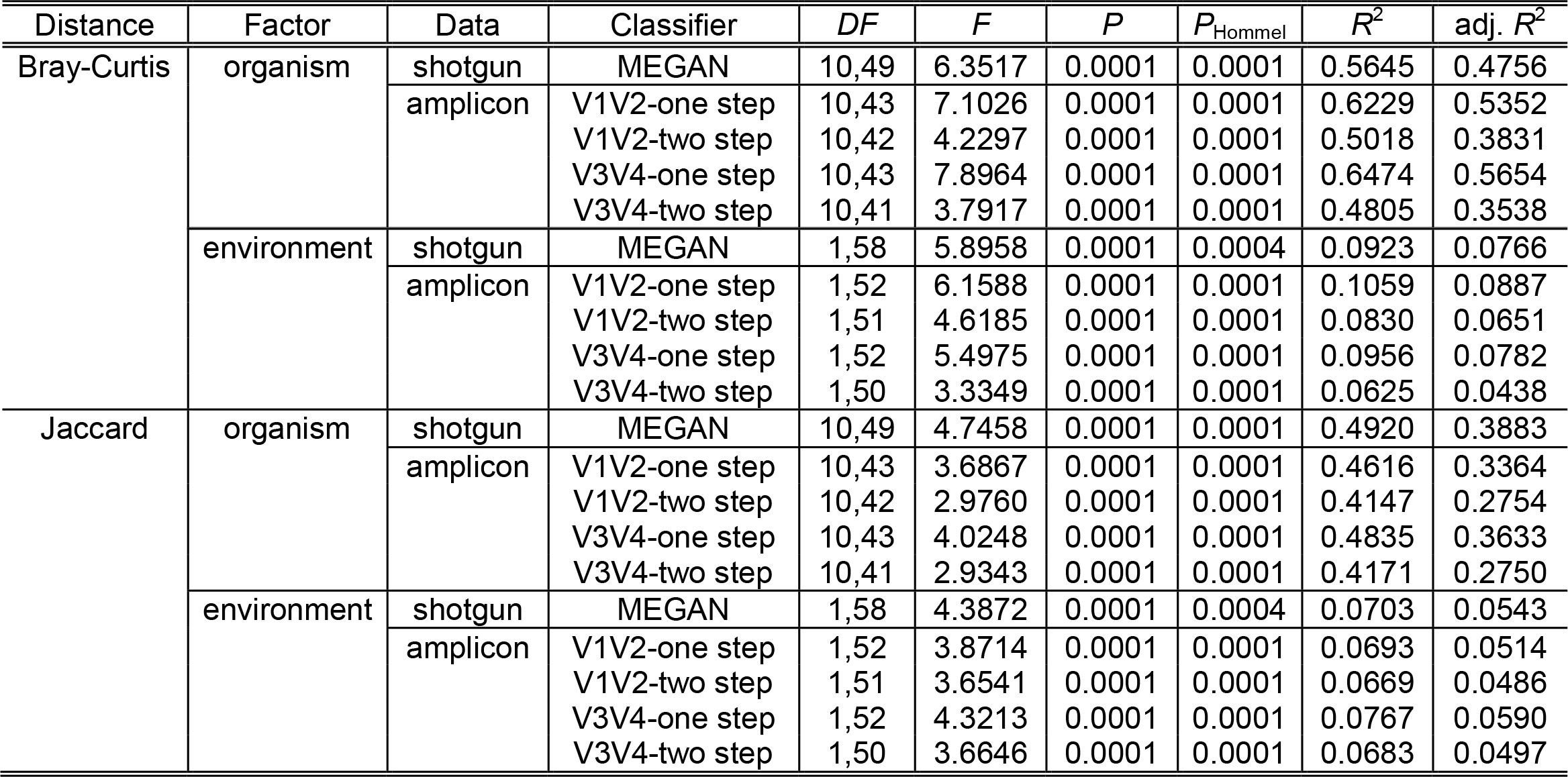
Taxonomic distance based PERMANOVA results for differences in community composition (genus level) between host species and host environments based on shared abundance (Bray-Curtis) and shared presence (Jaccard), based on whole genome shotgun and different amplicon strategies (*P*-values are adjusted via Hommel procedure).

**Figure 3:**
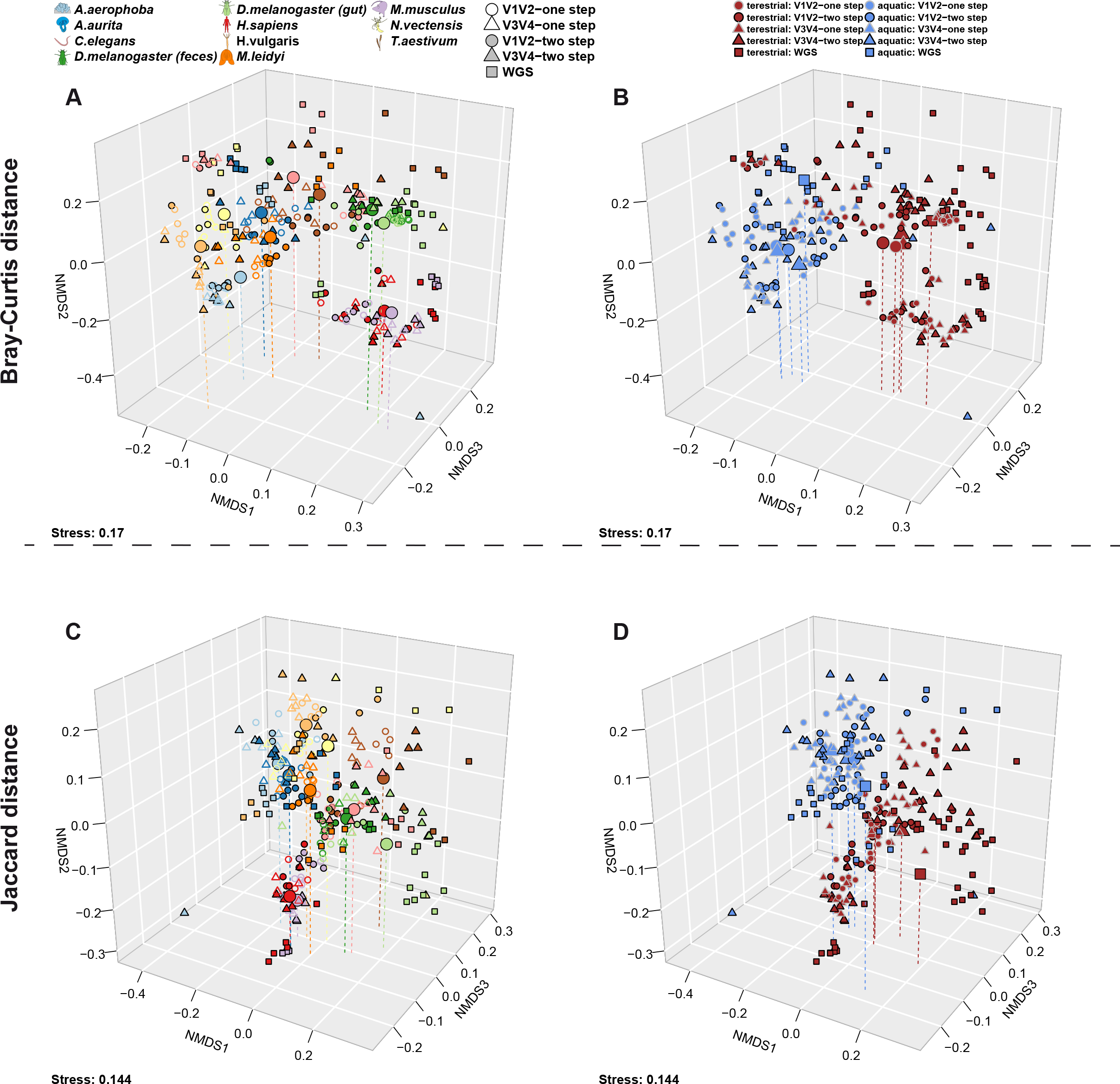
Non-metric Multidimensional Scaling of Bray-Curtis distances based on genus profiles derived from the different 16S rRNA gene amplicon methods (V1V2/ V3V4, one step/ two step) and shotgun derived genus profiles highlighting (A) host differences and (B) differences between host environments (terrestrial/aquatic; see Table 2). Non-metric Multidimensional Scaling of Jaccard distances based on genus profiles derived from the different 16S rRNA gene amplicon methods and shotgun derived genus profiles highlighting (C) host taxon differences and (D) differences between host environments (terrestrial/aquatic; see Table 2). Both panels show a separation based on host organisms and environments and not by method. Large symbols indicate the centroid of the respective host groups and vertical lines help to determine their position in space. Samples sizes are equal to Figure 2 (see also Table S1).

To identify individual drivers behind patterns of beta diversity, we performed indicator species analysis [19] at the genus level with respect to method, host species, and environment. Based on the amplicon data we identified 56 of 313 indicators to display consistent associations across all four amplicon techniques, such as *Bacteroides*, *Barnesiella*, *Clostridium IV*, and *Faecalibacterium* in *H. sapiens*, and *Helicobacter* and *Mucispirillum* in *M. musculus*, whereas other associations were limited to *e.g*. only one variable region (Table S7, S8). However, the overall pattern of host associations is largely consistent across methods (Figure S20). We also identified numerous indicator genera for aquatic and terrestrial hosts (Table S9, S10). Indicator analyses based on shotgun data reveals a smaller and less diverse set of host-specific indicators, which however show many congruencies with the amplicon-based data.

### Functional diversity within and between hosts

To examine the diversity (gene richness) of metagenomic functions across host species we evaluated EggNOG [20] annotations (assembly-based and MEGAN) to obtain a general functional spectrum (evolutionary genealogy of genes: Non-supervised Orthologous Groups), in addition to annotations derived from a database dedicated to functions interacting with carbohydrates (CAZY- Carbohydrate-Active enZYmes) [21]. Overall the individual host communities differ drastically in gene richness (EggNOG genes (MEGAN): *χ*^2^=52.202, *P*<2.10×10^−16^; EggNOG genes (assembly): *χ*^2^=49.986, *P*<2.10×10^−16^; CAZY: *χ*^2^=48.815, *P*<2.10×10^−16^; approximate Kruskal-Wallis test). Although the values also differ considerably between methods, overall the functional repertoires are most diverse in the vertebrate hosts, while only *H. vulgaris* and *A. aerophoba* as aquatic hosts carry a comparably diverse functional repertoire (Figure 4A, Figure S21). Interestingly, in contrast to taxonomic diversity we observe no difference in functional diversity between aquatic and terrestrial hosts.

**Figure 4:**
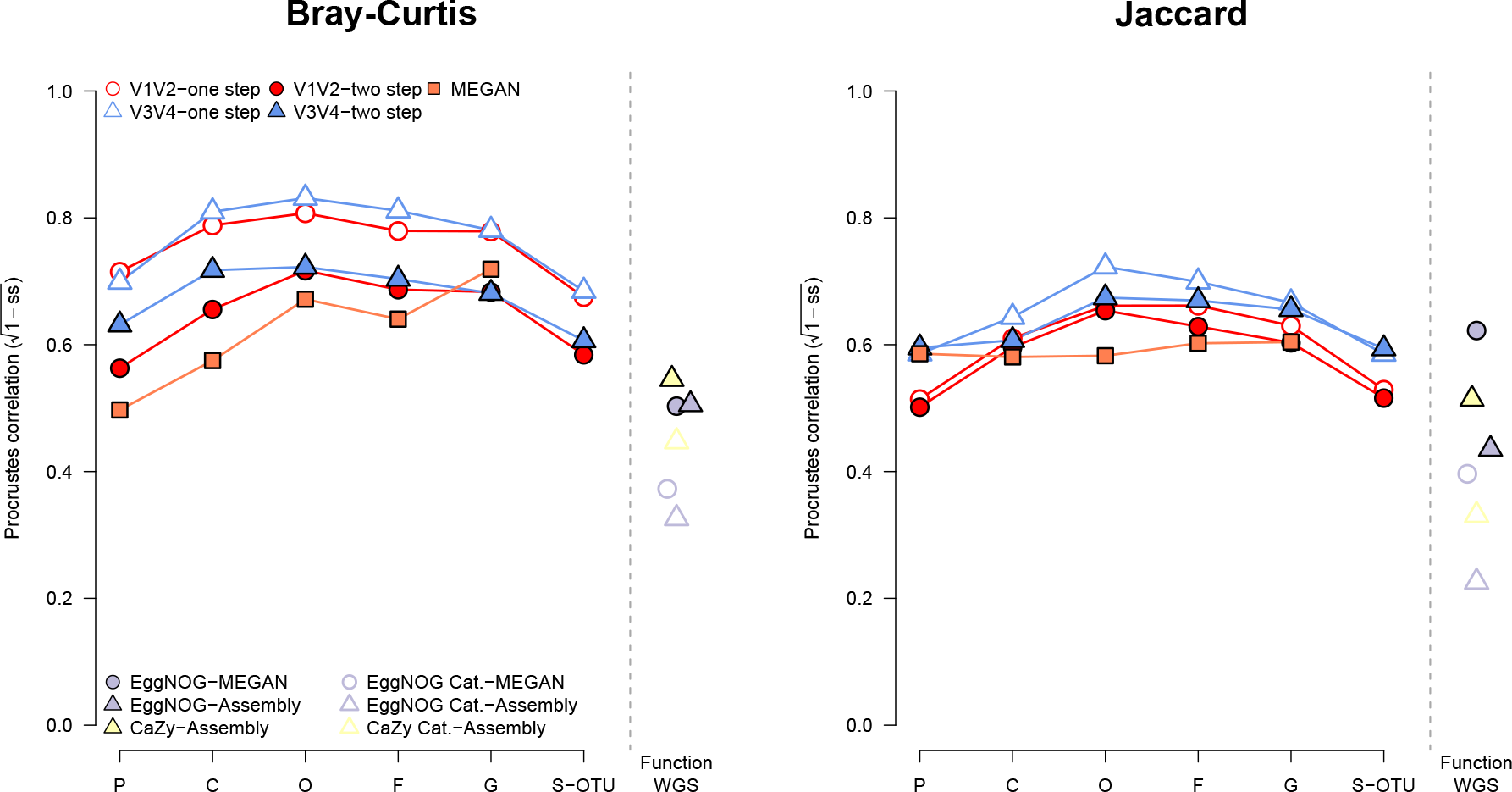
Multivariate correlation (Procrustes analyses) of phylogenetic distance among host organisms and community distances based on 16S rRNA gene amplicon- or shotgun derived community profiles at different taxonomic cutoffs, from Phylum to Genus and species level OTUs in the amplicon based profiles. Similar results are shown for the correspondence between functional composition based distances derived from imputed COGs and COG categories imputed from PICRUSt, and EggNOG derived genes and COG categories, as well as CAZY. All correlations are significant at *P* ≤ 0.05 (10’000 permutations). Large symbols indicate the centroid of the respective host groups and vertical lines help to determine their position in space.

Next we examined community differences (beta diversity) at the functional level, which are overall more pronounced (average adj. *R*^2^: 0.5084, Figure 4) than those based on taxonomic (genus level) classification (shotgun adj. *R*^2^: 0.4756; amplicon average adj. *R*^2^: 0.4594, see Table 2 and Table 3, Figure 3 and Figure 4, Figure S22). On the functional level aquatic and terrestrial hosts are considerably less distinct than observed at the taxonomic level (taxonomic shotgun adj. *R*^2^=0.0766; taxonomic amplicon average adj. *R*^2^=0.0690, functional shotgun average adj. *R*^2^=0.0441, see Table 2 and Table 3, Figure 4, S22). Variability of the functional repertoires was lowest in *A. aerophoba*, *D. melanogaster* feces and *M. musculus* gut contents, while *H. vulgaris*, *C. elegans*, and *D. melanogaster* gut samples displayed the highest intra-group distances, which translates to a higher amount of functional heterogeneity between replicates (Figure S23). This reflects in large part the patterns we observed in taxonomic variability of those host-associated communities (Figure S19).

**Table 3:**
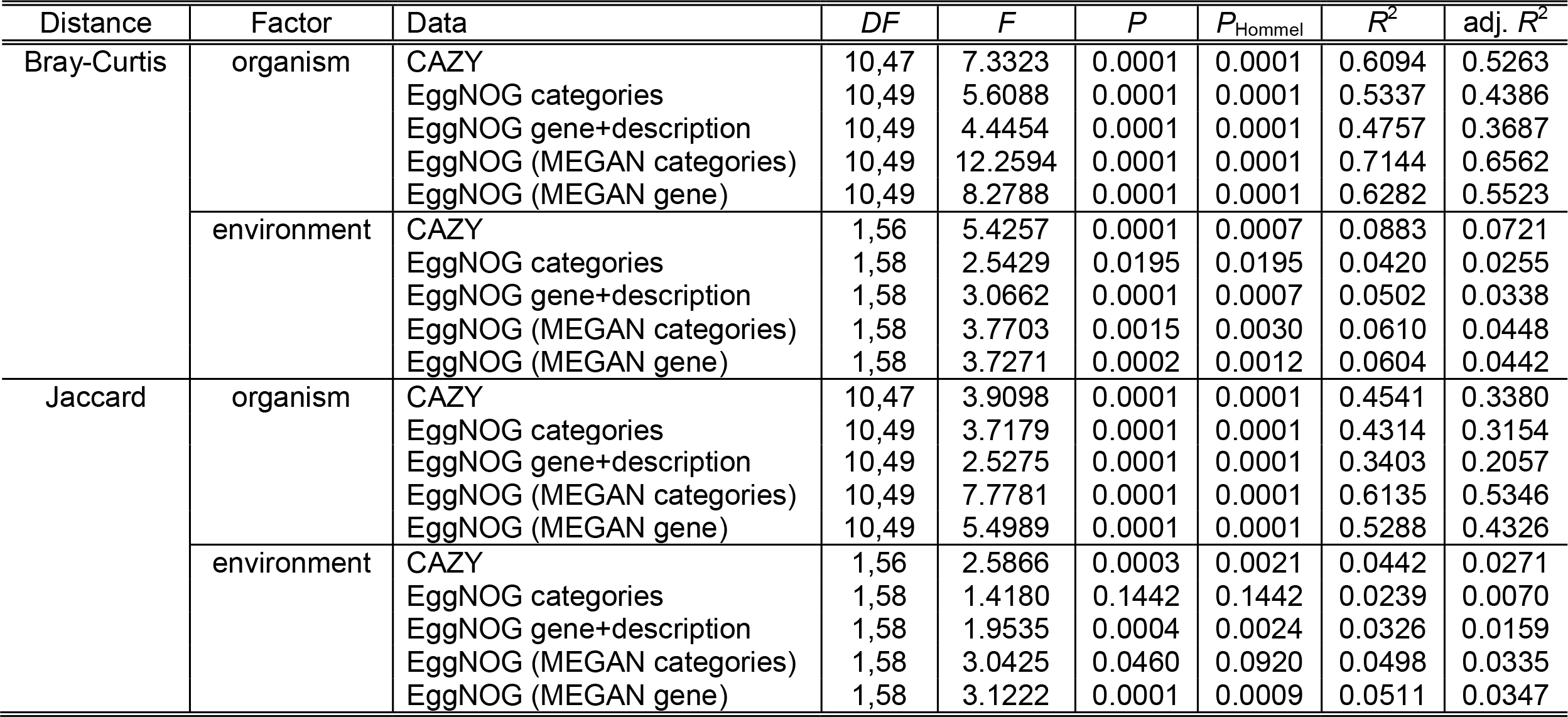
Functional distance based PERMANOVA results for differences in general functional community composition (EggNOG) and carbohydrate active enzymes (CAZY) between host species and host environments based on shared abundance (Bray-Curtis) and shared presence (Jaccard) of functions (*P*-values are adjusted via Hommel procedure).

### Indicator functions

To identify specific functions that are characteristic of individual hosts, we applied indicator analysis to functional categories. General functions in EggNOG reveal several interesting patterns, including CRISPR related genes in *A. aerophoba*, *H. sapiens*, and *H. vulgaris*, suggesting a particular importance of viruses in these communities. *A. aerophoba* possess a large set of characteristic genes involved in energy production and conversion, amino acid transport and metabolism, replication, recombination and repair. *M. musculus* and others appear to possess a large number of characteristic genes involved in carbohydrate transport and metabolism, energy production and conversion, transcription and cell wall/membrane/envelope biogenesis. *H. vulgaris* is characterized by a high number of genes involved in transcription, inorganic ion transport, metabolism, signal transduction mechanisms and cell wall/membrane/envelope biogenesis (Table S11-S13).

Analysis of carbohydrate-metabolizing functions based on CAZY [21] (Carbohydrate-Active enZYmes) reveals the highest number of characteristic glycoside hydrolases (GH) in *H. sapiens* and *M. musculus*, whereas polysaccharide lyases (PLs) for non-hydrolytic cleavage of glycosidic bonds are present in *A. aerophoba* and *H. sapiens* (Table S14). Parts of the cellulosome are only present in *A. aerophoba* and not in *M. musculus* or *H. sapiens*. Interestingly, only the freshwater *H. vulgaris* carries characteristic auxiliary CAZYs involved in lignin and chitin digestion, which may reflect dietary adaptations of the host.

### Performance of metagenome imputation from 16S rRNA gene amplicon data using PICRUSt across metaorganisms

Researchers often desire to obtain the insight gained from functional metagenomic information despite being limited to 16S rRNA gene data, for which imputation methods such as PICRUSt can be employed [5]. However, due to their dependence on variable region and database coverage [5], these imputations must be viewed with caution. Given our data set of both 16S amplicon- and shotgun metagenomic sequences, we systematically evaluated the performance of PICRUSt predictions across hosts and amplicon data type (V1V2, V3V4, one step/ two step protocol). Beginning with the mock community, the V1V2 region displays lower performance for imputing functions compared to V3V4, as indicated by a higher weighted Nearest Sequenced Taxon Index (NSTI) (*t*=17.812, *P*=1.119×10^−7^, Figure S24). High NSTI values imply low availability of genome representatives for the respective sample, due to either large phylogenetic distance for each OTU to its closest sequenced reference genome or a high frequency of poorly represented OTUs [5]. Comparing the distribution of functional categories based on Clusters of Orthologous Groups (COG) [22] between the different imputations (no cutoff applied) and the actual shotgun based repertoires reveals considerable overlap (Figure S24). Exceptions include the functional category R (general function prediction only), which is almost absent in the shotgun data, while the category S (function unknown) is more abundant among the shotgun based functional data (Figure S24).

Next we evaluated functional imputations for the different host species and amplification methods. We found no significant difference in average NSTI values or prediction success (NSTI < 0.15) between amplification protocols or variable region. However, approximately a third (31.8%) of the samples are lost due to incomplete imputation (NSTI > 0.15; Figure 5A). Notable problematic host taxa are *A. aerophoba* and *H. vulgaris*, for which no sample remained below the NSTI cutoff value. Other host taxa displayed clear differential performance with regard to the variable region used, whereby *H. sapiens*, *N. vectensis* and *T. aestivum* were successfully predicted based on V3V4, but not V1V2. However, when we employ Procrustes tests to compare community functional profiles based on shotgun sequencing (single assembly, MEGAN) and functional imputations at the COG-category level, we find a lower correspondence of the V3V4-based imputations compared to those based on V1V2 (Figure 5B), while the amplification methods displayed no significant difference. A similar pattern is observed when we correlate community differences based on shotgun results and lower level (single functions) COG annotations based on PICRUSt, although the difference is not significant (*F*_1,18_=0.6172, *P*=0.4423).

**Figure 5:**
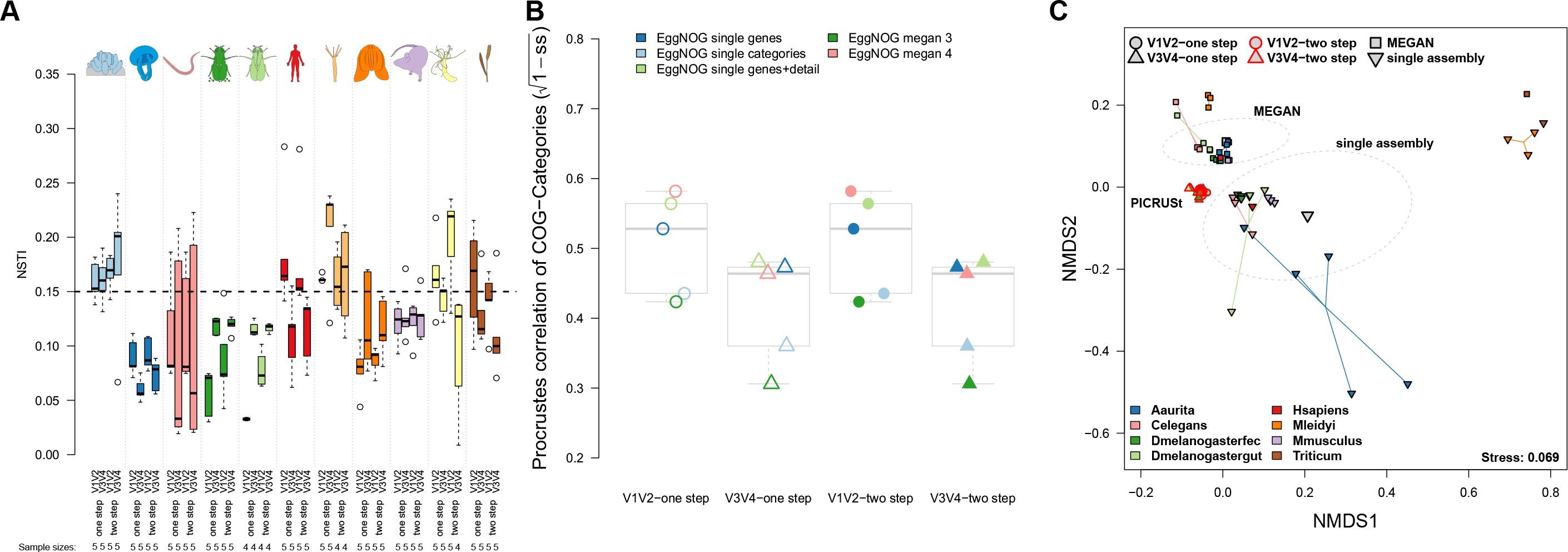
(A) Differences in Nearest Sequenced Taxon Index (imputation success) between variable regions (average: *Z*=0.3869, *P*=0.7017, approximate Wilcoxon test; probability: odds ratio=1.5941, *P*=0.1402, Fisher test) and amplification method (*Z*=0.0667, *P*=0.9472, approximate Wilcoxon test; probability: odds ratio=1.5511, *P*=0.1436, Fisher test). (B) Procrustes correlation of imputed and shotgun based COG categories among different techniques, with significantly higher correspondence between imputed and measured functional profiles in the V1V2 compared to the V3V4 region (*F*_1,18_=7.8537, *P*=0.0118, ANOVA). (C) Non-metric Multidimensional Scaling displays Bray-Curtis distances based on functional category abundances (COG categories) derived from PICRUSt (V1V2/ V3V4, one step/ two step) and shotgun based approaches (MEGAN, single assembly). Ellipses represent standard deviations of points within the respective groups.

To investigate the similarities among methods in more detail, we merged shotgun and PICRUSt based annotations at the level of COG categories. Principle coordinate analysis reveals only small differences between imputations with regard to amplification method or variable region (Figure 5C). However, large differences exist between the PICRUSt and shotgun based functional repertoires, as well as between the shotgun techniques (MEGAN, single assembly). Differences between the shotgun techniques were significant, but smaller than their distance to the imputed functional spectra (Figure 5C, Table S15). Finally, we examined the abundance of functional categories within single host taxa and the mock community, which reveals a higher relative abundance of functions related to energy production and conversion (C), replication, recombination and repair (L), and unknown functions (S) in the assembly-based annotations compared to the other techniques, which might be an important driver of the observed differences (Figure S24, S25).

Thus, in summary, the PICRUSt imputed functional repertoires significantly differ from actual shotgun profiles. While variation in imputation success is largely dependent on the identity of the particular host community, V3V4 appears to more often yield successful imputations. However, when successful, V1V2-derived imputations display closer similarity to actual functional profiles. Finally, the amplification method (one step, two step) appears to have no significant effect on the quality of functional imputation. These data therefore support the notion that metagenome imputations should be evaluated with care, as they depend on the underlying variable region and sample source.

### Phylogenetic patterns in microbial community composition

The term “phylosymbiosis” refers to the phenomenon where the pattern of similarity among host-associated microbial communities parallels the phylogeny of their hosts [23]. Highly divergent hosts with drastic differences in physiology and life history might be expected to overwhelm the likelihood of observing phylosymbiosis, which is typically observed within a given host clade [23]. However, the factors driving differences in composition among our panel of hosts may also be expected to vary in terms of the bacterial phylogenetic scale at which they are most readily observed [24]. Thus, we evaluated the degree to which bacterial community relationships (beta diversity) reflect the underlying phylogeny of our hosts at a range of bacterial taxonomic ranks, spanning from the genus to the phylum level.

In order to assess the general overlap between beta diversity and phylogenetic distance of the host species, we performed Procrustes analysis [25]. These analyses reveal that the strongest phylogenetic signal is observed when bacterial taxa are grouped at the order and/or family level, whereby the one step protocols and the V3V4 region display greater correlations to phylogenetic distance (Figure 6A). A similar pattern is observed for shotgun based community profiles (*i.e*. MEGAN), although its fit increases again at the genus level. Measuring beta diversity based on co-occurrence of bacterial taxa between hosts (Jaccard) displays a weaker correspondence to host phylogeny than the abundance-based measure (Bray-Curtis) (Figure 6).

**Figure 6:**
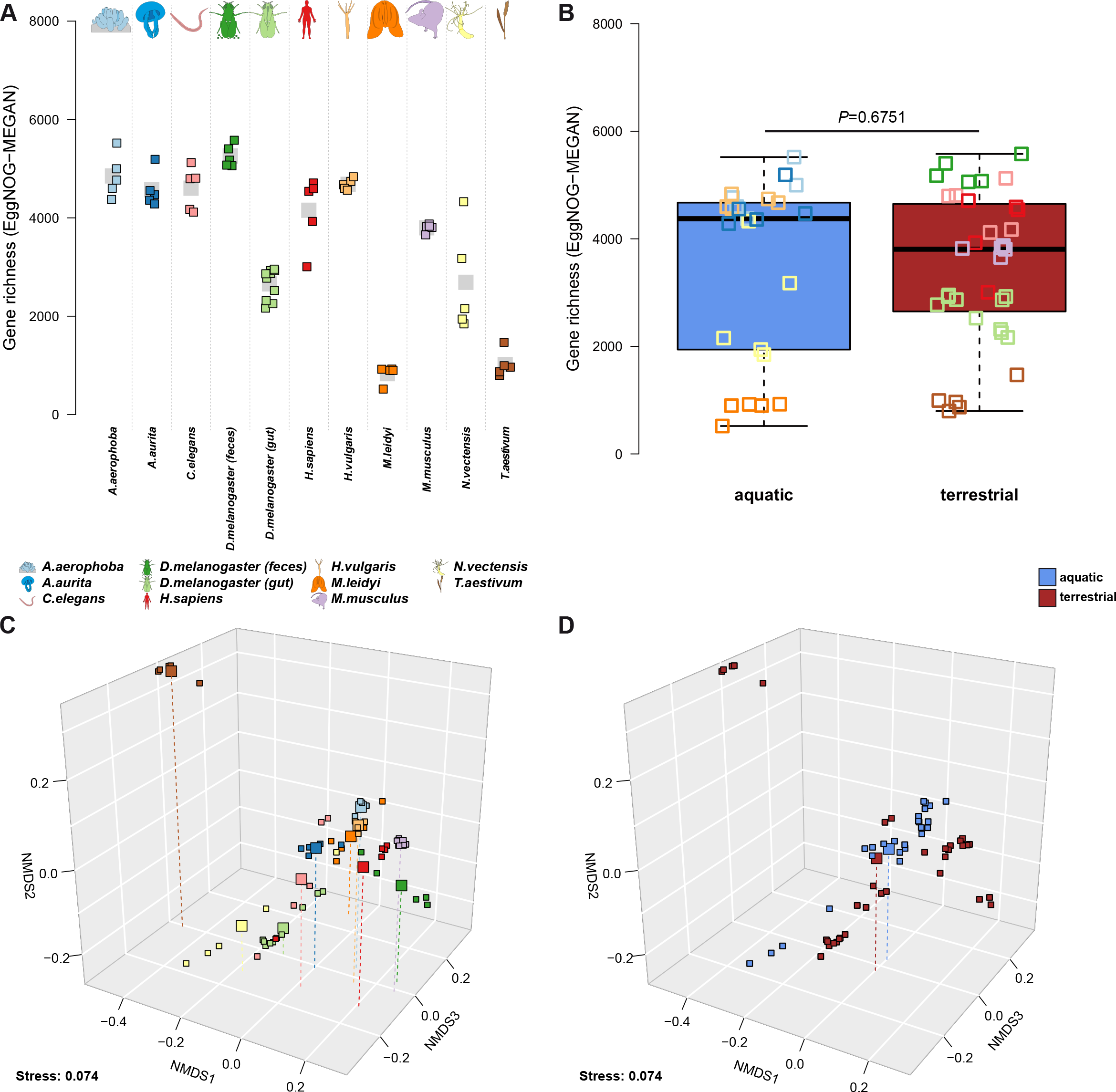
Functional diversities were derived from the number and abundances of MEGAN based EggNOG annotations. Functional richness between (**A**) host organisms and (**B**) host environmental groups based is displayed, as well as functional differences between hosts (**C**) and environmental groups (**D**). Non-metric Multidimensional Scaling is based on Bray-Curtis distances on the differences in functional composition between the host organisms is displayed (**C**, **D**; see Table 3). Large symbols indicate the centroid of the respective groups. Functional variation of communities based on pairwise Bray-Curtis distances within host organism groups and environmental groups. Samples sizes for the host taxa is N=5, except for D. melanogaster gut tissue (N=10; see Table S1).

To assess the fit of individual host taxa, we examined the residuals of the correlation between community composition and phylogenetic distance. This reveals a large variation in correspondence among host taxa, with *M. musculus*, *M. leidyi*, *H. sapiens* and *D. melanogaster* (feces) displaying the highest, while *H. vulgaris*, *C. elegans*, and *A. aerophoba* display the lowest correspondence between their microbiome composition and phylogenetic position (largest residuals Figure S26). Furthermore, terrestrial hosts display an overall better correspondence between co-occurrences of bacterial genera and host relatedness (V1V2 one step: *Z*=2.9578, *P*=0.0025), as do measurements based on V3V4 (one step: *Z*=2.7496, *P*=0.0054; two step: *Z*=2.8097, *P*=0.0046; approximate Wilcoxon test).

Next, given the peak of correspondence between bacterial community composition and host phylogeny observed at the order and/or family level, we set out to identify individual community members whose abundances best correlate to host phylogenetic distance using Moran’s eigenvector method [26]. This reveals 41 bacterial families and 36 orders with significant phylogenetic signal based on one or more amplicon data set, whereby 16 families and 18 orders display repeated associations across methods (*e.g. Clostridia, Ruminococcaceae*, *Helicobacteraceae*, *Lachnospiraceae*, *Coriobacteriaceae*, *Erysipelotrichaceae*, *Selenomonadales*, *Bacteroidales*, *Desulfovibrionales*; Table S16; Figure S27, S28). Analyzing communities based on shotgun data on the other hand identifies 215 bacterial families and 97 orders associated with phylogenetic distances, whereby 69 and 27 display repeated associations, respectively (Table S17; Figure S29, S30). The combined results of these analyses identify several families and orders with strong and consistent phylogenetic associations, in particular for the vertebrate hosts (*e.g. Bacteroidaceae*/ *Bacteroidales*, *Bifidobacteriaceae*/ *Bifidobacteriales*, *Coriobacteriaceae*/ *Coriobacteriales*, *Desulfovibrionaceae*/ *Desulfovibrionales*, *Erysipelotrichaceae*/ *Erysipelotrichales*, *Porphyromonadaceae*/ *Bacteroidales*, *Ruminococcaceae*/ *Clostridiales*, *Selenomonadales*; see Table S16). Other individual examples include bacteria related to *Helicobacteraceae/ Campylobacterales* in *A. aurita*, which are observed in other marine cnidarians and may be involved in sulfur oxidation [27]. *Alcanivoracaceae*, an alkane degrading bacterial group, is strongly associated to the coastal cnidarian *N. vectensis*. This association might originate from adaptation to a polluted coastal environment [28]. *Acidobacteria Gp6 and Gp9* specifically occur in *A. aerophoba* and are commonly associated to the core microbial community of sponges [29].

### Phylogenetic patterns in functional community composition

In order to contrast the patterns observed at the taxonomic level to those based on function we used Procrustes correlation to measure the overlap between phylogenetic distance and community distance based on the panel of functional categories in our analyses. Interestingly, the two functional categories displaying the greatest correspondence to host phylogeny are the CAZY and single EggNOG based functions (Figure 6). The remainder of patterns between phylogeny and bacterial functional spectra differed among the host species and functional categories (Figure S26), *T. aestivum* and *D. melanogaster* (feces) display the lowest correspondence, while *C. elegans, M. musculus and H. sapiens* display the best correspondence (lowest residuals, Figure S26) between their functional repertoire and phylogenetic position. As observed for the taxonomic analyses, terrestrial hosts again display a slightly better correlation than aquatic hosts (smaller residuals), in particular for the co-abundance of EggNOG categories (*Z*=2.2116, *P*=0.0267), CAZY (*Z*=2.0393, *P*=0.0414) and the co-occurrence of EggNOG categories (*Z*=2.7377, *P*=0.0061) and genes (*Z*=3.3062, *P*=0.0007; approximate Wilcoxon test) among hosts.

Finally, to reveal individual functions correlating to host phylogeny, we used the aforementioned Moran’s I eigenvector analyses with additional indicator analyses to narrow the potential clade associations. Interestingly, most functions that correlate to a specific host taxon/clade (1-3 taxa) are mainly restricted to vertebrate hosts or in combination with a vertebrate host (Table S18-S21). This pattern is repeated across all functional annotations used in this study. Examples include fucosyltransferases, fucosidases, polysaccharide binding proteins, as well as hyaluronate, xanthan, and chondroitin lyases that stem from CAZY (see Figure S31, Table S18). These functions are all related to glycan- and mucin degradation and interaction, which mediate many intimate host-bacterial interactions and are also observed in subsequent analyses based on general functional databases (EggNOG; Table S19, Table S20). Many other phylogenetically correlated functions appear to be driven by the vertebrate hosts as well, which likely reflects the high functional diversity within this group (see Figure 4 and Figure S23). Only *LPXC* and *LPXK* (EggNOG), genes involved in the biosynthesis of the outer membrane, are exclusively associated to the non-vertebrate hosts (LPXC: UDP-3-O-acyl-N-acetylglucosamine deacetylase, LPXK: Tetraacyldisaccharide 4’-kinase), as is an oxidative damage repair function (MSRA reductase) associated to *H. vulgaris* (Table S19, Figure S31). EggNOG category Q (secondary metabolites biosynthesis, transport and catabolism) is also characteristic of invertebrate hosts in addition to a small number of metabolic functions (*i.e*. dehydrogenases, mono oxygenase, fatty acid hydroxylase; MEGAN based; Table S20, Figure S31). More generally we observe a high number of genes of unknown function (S), carbohydrate transport and metabolism (G), replication, recombination and repair (L), cell wall/membrane/envelope biogenesis (M), and energy production and conversion (C) (Table S21 Figure S31). Finally, antibiotic resistance genes and virulence factors also show frequent phylogenetic and host specific signals (Table S19, S20; Figure S31).

## Discussion

Despite the great number of metagenomic studies published to date, which range in their focus on technical, analytical or biological aspects, our study represents a unique contribution given its breadth of different host samples analyzed with a panel of standardized methods. In particular, the tradeoffs between 16S rRNA gene amplicon- versus shotgun sequencing concerning amplification bias, functional information and both monetary and computational costs, warrant careful consideration when designing research projects. While 16S rRNA gene amplicon-based analyses are subject to considerable skepticism and criticism, we demonstrate that in many aspects similar, if not superior characterization of bacterial communities is achieved by these methods, although discrepancies associated with shotgun based data are largely dependent on the analytic pipeline. We also show, however, that important insight can be gained through the combination of taxonomic- and functional profiling, and that imputation-based functional profiles significantly differ from actual profiles. Our findings thus provide a guide for selecting an appropriate methodology for metagenomic analyses across a variety of metaorganisms. Finally, these data provide novel insight into the broad scale evolution of host-associated bacterial communities, which can be viewed as particularly reliable given the repeatability of observations (*e.g*. differences between aquatic and terrestrial hosts, indicator taxa) across methods.

Given the concerns regarding the accuracy of 16S rRNA gene amplicon sequencing, other studies such as that of Gohl *et al*. [8] performed systematic comparisons of different library preparation methods, and found superior results for a two step amplification procedure. This method offers the additional advantage that one panel of adapter/barcode sequences can be combined with any number of different primers. Our first analyses were based on a standard mock community including Gram positive and Gram negative bacteria from the Bacilli and Gamma Proteobacteria (eight species), as well as two fungi, which did not support an improvement of performance based on the two step protocol. However, a number of changes were made to the Gohl *et al*. [8] protocol to adapt it to our lab procedures (*e.g*. larger reaction volumes, polymerase, variable region, heterogeneity spacers) that may contribute to these discrepancies, in addition to our different and diverse set of samples and other factors with potential influence on the performance of amplicon sequencing [6–8, 30–32]. The complexity of the mock community, *i.e*. the number of taxa, distribution, and phylogenetic breadth, may also have an influence on the discovery of clear trends in amplification biases or detection limits for certain taxonomic groups [33]. Thus, the even and phylogenetically shallow mock community in our study may be less suited than the staggered and diverse mixtures used in other studies [8], but still provides valuable information on repeatability, primer biases, and accuracy [33]. Nonetheless, when applied to our range of complex host-associated communities, we also found that significant differences in most parameters were due to the variable region rather than amplification method, and in many cases biological signals were either improved- or limited to the one step protocol.

Additional sources of variation influencing the outcome of our 16S rRNA gene amplicon-based community profiling are the bioinformatic pipelines we employed, starting from trimming and merging to clustering and classification, which are stringent and incorporate more reliable *de novo* clustering algorithms [34] as well as different classification databases [35]. Heterogeneity among the different amplicon approaches is however far smaller than the observed heterogeneity between amplicon and shotgun methods, or within different shotgun analyses, as observed in other benchmarking studies [31]. Differences between shotgun approaches have been investigated in detail and also yield varying performances among classifiers, but in general find a comparatively high performance of MEGAN based approaches [9, 36, 37], which we also confirm in our study.

Given the limited number of studies that have compared imputed- and shotgun derived functional repertoires [5, 38], our study also provides important additional insights. As imputation by definition is data-dependent, the differential performance and prediction among hosts in our study may in large part be explained by the amount of bacteria isolated, sequenced, and deposited (16S rRNA or genome) from these hosts or their respective environments. This seems to be most critical for the aquatic hosts. Furthermore, we observe a clear effect of variable region on the prediction performance, which is most obvious based on the mock community. The PICRUSt algorithm was developed and tested using primers targeting V3V4 16S rRNA, thus optimization of the imputation algorithm might be biased towards this target over the V1V2 variable region. Although these performance differences, in particular the bias towards model organisms compared to less characterized communities (*e.g*. hypersaline microbial mats), were previously shown [5], our study provides additional, experimentally validated guidelines for a number of novel host taxa.

Interestingly, the strongest correspondence between bacterial community similarity and host genetic distance was detected at the bacterial order level for most of the employed methods. This may on the one hand reflect the deep phylogenetic relationships between our host taxa, such that turnover of bacterial taxa erodes phylosymbiosis over time [23, 24]. On the other hand, some of the more striking observations made among our host taxa are the differences between aquatic and terrestrial hosts, both at the level of alpha and beta diversity. Based on a molecular clock for the 16S rRNA gene of roughly 1% divergence per 50 million years [39], bacterial order level divergence corresponds well with the timing of animal terrestrialization (425-500 MYA) [40, 41]. Although evolutionary rates can widely vary among bacteria species [42], other studies of individual gut microbial lineages such as the *Enteroccoci* indicate that animal terrestrialization was indeed a likely driver of diversification [43]. Specifically the changing availability of carbohydrates in the host gut can be seen as a main driver of this diversification, which is consistent with the association of CAZY-based functional repertoires correlating to phylogenetic distance in our data set [23, 44].

In contrast to the patterns observed based on 16S rRNA gene amplicon-based profiles, the differentiation of bacterial communities according to host habitat was less pronounced based on functional genomic repertoires. This raises the possibility that the colonization of land by ancient animals required the acquisition of new, land-adapted bacterial lineages to perform some of the same ancestral functions. The overall observation of increased beta diversity among terrestrial-compared to aquatic hosts (Figure S19) could in part reflect differential acquisition among host lineages after colonizing land, although dispersal in the aquatic environment may on the other hand act as a greater homogenizing factor among aquatic hosts. The stronger correspondence between bacterial community- and host phylogenetic distance among terrestrial hosts is also generally consistent with this hypothesis. However, the higher alpha diversity and the slightly lower correspondence with the phylogenetic patterns in aquatic hosts may also indicate a higher influence of environmental bacteria or a lack of physiological control over bacterial communities.

Bacterial taxa and functions involved in carbohydrate utilization were among the most notable associations to individual hosts, groups of hosts, and/or host phylogenetic relationships. Taxa such as *Bacteroidales*, *Ruminococcaceae/ Ruminococcales*, and *Clostridia* associated to humans and/or mice include members known for a mucosal lifestyle, and these hosts also display the most diverse and abundant repertoire of carbohydrate active enzymes (particularly glycosylhydrolases) in their microbiome. Other examples include sialidases, esterases, and fucosyltransferases, as well as different extracellular structures that appear to be specific to aquatic hosts, indicating differences in mucus and glycan composition according to this host environment. Glycan structures provide a direct link between the microbial community and the host via attachment, nutrition, and communication [45, 46], and the composition of mucin and glycan structures themselves show strong evolutionary patterns and are distinct among taxonomic groups [44]. Thus, a high diversity of glycan structures within and between hosts may determine the specific sets carbohydrate facilitating enzymes of the respective microbial communities.

In addition to the bacterial carbohydrate hydrolases that digest surrounding host and dietary carbohydrates, we also identified a number of glycosyltransferases associated with capsular polysaccharide synthesis (Table S19, Table S20). This type of glycosylation is an important facilitator for host association and survival [47] and plays a crucial role in infections [48]. The capsule prevents opsonization and phagocytosis through the host immune system and gives the bacterium the ability to modulate its interaction with the host environment [47, 49]. This type of manipulation is performed by mutualists and pathogens alike [47, 50] via molecular mimicry and tolerogenic immune modulation [51, 52]. Bacterial glycan products like polysaccharide A (PSA) may also have direct benefits for the host, as it can interfere with the host immune system by increasing immunologic tolerance, or inhibit the binding of other microbes (*e.g. Helicobacter hepaticus* [53]). Thus, capsular and excreted glycan structures are important for the successful colonization and persistence in different environments [54, 55] and host organisms [47, 55].

## Conclusions

In summary, the systematic comparison of five different metagenomic sequencing methods applied to ten different holobiont yielded a number of novel technical and biological insights. Although important exceptions will exist, we demonstrate that broad scale biological patterns are largely consistent across these varying methods. While the richer information provided by shotgun sequencing is clearly desirable and is likely to surpass amplicon-based profiling techniques in the foreseeable future, technical variability among analytical pipelines currently surpasses that observed between different amplicon methods. As many aspects of differential performance in our study are host-specific (more detailed description of individual hosts can be found in the Supplementary Material), future development and benchmarking analyses would also benefit from a including a range of different host/environmental samples.

## Methods

### DNA extraction and 16S rRNA gene amplicon sequencing

Protocols for each host type are described in the Supplementary Material (see also Figure S18-S28). Each library (16S rRNA gene amplicon, shotgun) included at least one mock community sample based on the ZymoBIOMICS™ Microbial Community DNA Standard (Lot.: ZRC187324, ZRC187325) consisting of 8 bacterial species (*Pseudomonas aeruginosa* (10.4%), *Escherichia coli* (9.0%), *Salmonella enterica* (11.8%), *Lactobacillus fermentum* (10.3%), *Enterococcus faecalis* (14.1%), *Staphylococcus aureus* (14.6%), *Listeria monocytogenes* (13.2%), *Bacillus subtilis* (13.2%)) and two fungi (*Saccharomyces cerevisiae* (1.6%), *Cryptococcus neoformans* (1.8%)).

The 16S rRNA gene was amplified using uniquely barcoded primers flanking the V1 and V2 hypervariable regions (27F-338R) and V3V4 hypervariable regions (515F-806R) with fused MiSeq adapters and heterogeneity spacers in a 25 µl PCR [32]. For the traditional one step PCR protocol we used 4 µl of each forward and reverse primer (0.28 µM), 0.5 µl dNTPs (200 µM each), 0.25 µl Phusion Hot Start II High-Fidelity DNA Polymerase (0.5 Us), 5 µl of HF buffer (Thermo Fisher Scientific, Inc., Waltham, MA, USA) and 1 µl of undiluted DNA. PCRs were conducted with the following cycling conditions (98°C-30s, 30×[98°C-9s, 55°C-60s, 72°C-90s], 72°C-10 min) and checked on a 1.5 % agarose gel. Using a modified version of the recently published two step PCR protocol by Gohl *et al*. 2016, we employed for the first round of amplification fusion primers consisting of the 16S rRNA gene primers (V1V2, V3V4) and a part of the Illumina Nextera adapter with the following cycling conditions in a 25 µl PCR reaction (98°C-30s, 25×[98°C-10s, 55°C-30s, 72°C-60s], 72°C-10 min) [8]. Following the PCR was diluted 1:10 and 5µl of the solution were used in an additional reaction of 10 µl (98°C-30s, 10×[98°C-9s, 55°C-30s, 72°C-60s], 72°C-10 min) utilizing the Nextera adapter overhangs to ligate the Illumina adapter sequence and individual MIDs to the amplicons following the manufacturer’s instructions. The PCR protocol we used 1 µl of each forward and reverse primer (5 µM), 0.3 µl dNTPs (10 µM), 0.2 µl Phusion Hot Start II High-Fidelity DNA Polymerase (2 U/µl), 2 µl of 5×HF buffer (Thermo Fisher Scientific, Inc., Waltham, MA, USA) and 5 µl of the diluted PCR product. The concentration of the amplicons was estimated using a Gel Doc™ XR+ System coupled with Image Lab™ Software (BioRad, Hercules, CA USA) with 3 µl of O’GeneRulerTM 100 bp Plus DNA Ladder (Thermo Fisher Scientific, Inc., Waltham, MA, USA) as the internal standard for band intensity measurement. The samples of individual gels were pooled into approximately equimolar subpools as indicated by band intensity and measured with the Qubit dsDNA br Assay Kit (Life Technologies GmbH, Darmstadt, Germany). Sub pools were mixed in an equimolar fashion and stored at −20°C until sequencing.

Library preparation for shotgun sequencing was performed using the NexteraXT kit (Illumina) for fragmentation and multiplexing of input DNA following the manufacturer’s instructions. Amplicon sequencing was performed on the Illumina MiSeq platform with v3 chemistry (2×300 cycle kit), while shotgun sequencing was performed via 2×150bp Mid Output Kit at the IKMB Sequencing Center (CAU Kiel, Germany).

### Amplicon analysis

The respective V1V2 and V3V4 PCR primer sequences were removed from the sequencing data using *cutadapt* (v.1.8.3) [56]. Sequence data in FastQ format was quality trimmed using *sickle* (v.1.33) in paired-end mode with default settings and removing sequences dropping below 100bp after trimming [57]. Forward and reverse read were merged into a single amplicon read using VSEARCH allowing fragments with a length of 280-350 bp for V1V2 and 350-500 bp for V3V4 amplicons [58]. Sequence data was quality controlled using fastq_quality_filter (FastX Toolkit) retaining sequences with no more than 5% of per-base quality values below 30 and subsequently with VSEARCH discarding sequences with more than 1 expected errors [58, 59]. Reference guided chimera removal was performed using the gold.fa reference in VSEARCH (v2.4.3). The UTAX algorithm was used for a fast classification of the sequence data in order to remove sequences not assigned to the domains Bacteria or Archaea and exclude amplicon fragments from Chloroplasts [60]. Notably, only a total of 15 sequences were assigned to the domain Archaea, all found in two samples of human feces, accounting for less than 0.1% of the clean reads in theses samples. The entire cleaned sequence data was concatenated into a single file, dereplicated and processed with VSEARCH for OTU picking using the UCLUST algorithm [61] using a 97% similarity threshold. OTUs were again checked for chimeric sequences, now using the *de novo* implementation of the UCHIME algorithm in VSEARCH [58, 61, 62]. All clean sequence data of the samples were mapped back to the cleaned OTU sequences using VSEARCH. OTU sequences and clean sequences mapping to the OTUs were taxonomically annotated using the RDP classifier algorithm with the RDP training set 14 [63, 64]. Sequence data were normalized by selecting 10,000 random sequences per sample. Taxon-by-sample abundance tables were created for all taxonomic levels from Phylum to Genus, as well as for OTUs.

### PICRUSt functional imputations

Species level OTUs (97% similarity threshold) were further classified using the GreenGenes (August 2013) database [65] via RDP classifier as implemented in mothur (v1.39.5) and merged with the abundances into a biome file which was uploaded to the Galaxy PICRUSt v1.1.1 pipeline (http://galaxy.morganlangille.com/) to derive functional imputations (COG predicitions) [5]. To achieve accurate functional predictions samples with NSTI ≤ 0.15 (weighted Nearest Sequenced Taxon Index) were pruned from the data set, as recommended by the developers.

### Shotgun sequencing

Raw demultiplexed sequences were trimmed via Trimmomatic (v0.36) for low quality regions with a minimum length of 50 bp as well as for adaptor and remaining MID sequences [66]. After trimming reads were mapped to host specific genome databases and *ΦX* with additional retention databases containing all fully sequenced bacterial and metagenomic genomes (05-09-2015) via DeconSeq (v0.4.3) [67]. Single and paired sequences were repaired using the BBTools (v37.28) repair function [68]. Combined sequences were searched against the non-redundant NCBI database (28-07-2017) via DIAMOND [69] with (evalue cutoff 0.001, v0.8.28) and MEGAN [14] classifying hits by functions (EGGNOG-Oct2016) and taxa (May2017) (v6.6.1). MetaPhlan [15] (v1.7.7) and MetaPhlan2 [16] (v2.2.0) was used for taxonomic classification. Forward and reverse reads were mapped to the SIVLA non-redundant database (v123) via SortmeRNA [17, 70] (2.1b) and classified via RDP classifier and the RPD 16 database as implemented in mothur [71]. Kraken (v0.10.5-beta) database was constructed on complete and dusted genome sequences of all archaea (+scaffolds), bacteria, fungi (+scaffolds), protozoa (+scaffolds), viruses and full sequences of plasmids and plastids [13] (database 21-08-2017), which was used to classify raw reads as well as assembled contigs, which were used throughout the manuscript. For assemblies of single samples we used metaSPADES [72] (v3.9.1) using paired reads in addition to unpaired reads left from the previous steps. PROKKA (v1.12) was used for gene calling and initial genome annotation [73] using the metagenome option with additional identifying rRNAs and snRNA via barnap, ARAGORN [74], and Infernal [75]. ORFs were further annotated via EggNOG annotation via HMMER models implemented in the eggnog-mapper (v0.12.7) [20, 76], CAZY database via dbCAN (v5, 07/24/2016) and HMMER3 [21, 77]. Gene abundances were derived from mapping the all reads back to the predicted ORF via bowtie2 (v2.2.6) [78] and calculated TPM (transcripts per kilobase million) via SamTools (v1.5) [79].

18S rRNA genes were obtained from NCBI GeneBank and aligned via ClustalW (v1.4) [80] for host tree construction, which includes *A. aerophoba* (gi:51095211, AY5917991), *M. leidyi* (gi:14517703, AF2937001), *H. vulgaris* (gi:761889987, JN5940542), *A. aurita* (gi:14700050, AY0392081), *N. vectensis* (gi:13897746, AF2543821), *T. aestivum* (gi:15982656, AY0490401), *M. musculus* (gi:374088232, NR_0032783), *H. sapiens* (gi:36162, X032051), *D. melanogaster* (gi:939630477, NR_1335591), and *C.elegans* (gi:30525807, AY2681171). Phylogenetic distance was calculated via DNADIST (v3.5c) [81] and a maximum likelihood tree was constructed via FastTree v2.1 CAT+Γ model [82]. Accuracy was improved via increased minimum evolution rounds for initial tree search [-spr 4], more exhaustive tree search [-mlacc 2], and a slow initial tree search [-slownni].

### Statistical analysis

Statistical analyses were carried via R [83] (v3.4.3). Alpha diversity indices (richness, Shannon-Weaver index) and beta diversity metrics based on the shared presence (Jaccard distance)- or abundance (Bray-Curtis distance) of taxa were calculated in the *vegan* package [84] and ordinated via Principal Coordinate Analysis (PCoA, avoiding negative eigenvalues), or via non-metric multidimensional scaling (NMDS) using a maximum of 10000 random starts to obtain a minimally stressed configuration in three dimensions. Clusters were fit via an iterative process (10’000 permutations) tested for separation by direct gradient analysis via distance based Redundancy analyses and permutative ANOVA (10’000 permutations) [85, 86]. Univariate analyses were carried out with approximate Wilcoxon/Kruskal tests as implemented in *coin* [87] (10’000 permutations). Procrustes tests were used to relate pairwise community distances based on either different data sources such as functional repertoires or taxonomic composition, as well as phylogenetic distances [25, 88]. Moran’s I eigenvector technique was employed to correlate bacterial community members and their functions to phylogenetic divergence, as implemented in *ape* (10’000 permutations) [26, 89]. Indicator species analysis, employing the generalized indicator value (*IndVal.g*), was used to assess the predictive value of a taxon for each respective host phenotype/category as implemented in *indicspecies* [19]. Linear mixed models, as implemented in *nlme* were used to compare the influence of amplification method or variable region without the influence of the organism of origin [90]. We employed the Hommel- and Benjamini-Yekutieli adjustment of *P*-values when advised [91, 92].

## Declarations

### Ethics approval and consent to participate (Human samples)

Study participants were randomly recruited from inhabitants of Schleswig-Holstein (Germany) which were recruited for the PopGen cohort. Five individuals from the PopGen biobank (Schleswig-Holstein, Germany) were randomly selected among the healthy and unmedicated individuals and included in the study without corresponding meta-information. Study participants collected fecal samples at home in standard fecal tubes and shipped them immediately at room temperature or brought them to the collection center (within 24 h). Samples were stored at −80°C until processing. Human feces (N=4) were sampled and extracted following the procedures as described in Wang *et al*. 2016 [93]. A biopsy sample of the sigmoid colon was taken from a healthy control individual without macro- or microscopical inflammation (N=1) and DNA was extracted as described in Rausch *et al*. 2011 [94]. Investigators were blinded to sample identities and written, informed consent was obtained from all study participants before the study. All protocols were approved by the Ethics Committee of the Medical Faculty of Kiel and by the data protection officer of the University Hospital Schleswig-Holstein in adherence with the Declaration of Helsinki Principles.

### Ethics approval for animal and plant samples

Wild derived, hybrid mice were sacrificed according to the German animal welfare law and Federation of European Laboratory Animal Science Associations guidelines. Hybrid breeding stocks of wild derived *M. m. musculus* × *M. m. domesticus* hybrids captured in 2008 are kept at the Max Planck Institute Plön (11th lab generation). The approval for mouse husbandry and experiment was obtained from the local veterinary office “Veterinäramt Kreis Plön” (Permit: 1401-144/PLÖ-004697). All sampling, including invertebrate and plant samples, was performed in concordance with the German animal welfare law and Federation of European Laboratory Animal Science Associations guidelines. Further details for each host type are provided in the Supplementary Material.

### Consent for publication

Not applicable.

### Availability of data and material

Sequence- and meta-data are accessible under the study identifier PRJEB30924 (“https://www.ebi.ac.uk/ena”). Remaining DNA from non-human samples can be made available upon request. All human samples and information on their corresponding phenotypes have to be obtained from the PopGen Biobank Kiel (Schleswig-Holstein, Germany) through a Material Data Access Form. Information about the Material Data Access Form and how to apply can be found at: “https://www.uksh.de/p2n/Information+for+Researchers.html”.

### Author contributions

PRa, PRo, AF, TB, and JFB conceived and designed research. PRa and MR performed data analyses. PRa, MR, BH, SD, and JFB interpreted results and wrote the manuscript. PRa, MR, TD, KD, HD, SD, SF, JF, UHH, FAH, BH, MH, MJ, CJ, KABK, DL, AR, TBHR, TR, RAS, HS, RS, FS, ES, NWB, PRo, AF, TB, and JFB generated and interpreted host-specific data and gave intellectual input. All authors read and approved the final manuscript.

## Acknowledgements

We thank Katja Cloppenborg-Schmidt, Melanie Vollstedt, and Dr. Sven Künzel for excellent assistance and help during the development of the project and their constant drive to improve its quality.

