## Supplemental Material and Figures for "Comparative analysis of amplicon and metagenomic sequencing methods reveals key features in the evolution of animal metaorganisms"

**Supplementary Material:**

While a single classification pipeline was employed for all four 16S rRNA gene amplicon-based sequence profiles, community composition based on shotgun data was evaluated using five different available classification methods (Kraken [1], MEGAN [2], MetaPhlan [3], MetaPhlan2 [4] and SortmeRNA [5]). Briefly, SortmeRNA identifies 16S rRNA gene reads from metagenomic data, which can be classified using reference databases. MetaPhlan and MetaPhlan2 are abundance-based estimator programs. They work by using a database of marker genes (single-copy genes present in nearly all bacteria) and unique clade-specific genes, to allow fast classification and abundance estimation of organisms present in the sample. MetaPhlan2 is an updated version of MetaPhlan with an expanded database of more than 1 million genes, including additional marker genes from viral, archaeal, and eukaryotic species. A drawback for all three previous methods is a lot of reads are still unclassified and the metagenome sample is not utilized to its full potential.

In contrast to the specifically designed reference sets used in the aforementioned approaches, Kraken and MEGAN use the information from all reads generated in the sequencing. Kraken uses a user-specified library of genomes as a database, where the records consist of a *k*-mer and the lowest common ancestor (LCA) whose genomes contain the k-mer. By querying the database for each *k*-mer in a sequence, and then using the resulting set of LCA taxa an appropriate label for the sequence can be determined. This drastically speeds up the classification process, but requires absolute matches which sacrifices sensitivity. MEGAN first needs a preprocessing step where reads are compared against a database (usually the non-redundant NCBI GeneBank database) using a BLAST algorithm or another comparison tools to search the top hits for each short read (here DIAMOND [6]). MEGAN then uses the NCBI taxonomy of the matched database to classify the sequences using a LCA algorithm, where reads are assigned to taxa such that the taxonomical level of the assigned taxon reflects the level of conservation of the sequence. MEGAN has the advantage that it simultaneously performs a taxonomical and functional classification of the reads simultaneously due to the inherent information of the database hits.

**Supplementary Figures:**

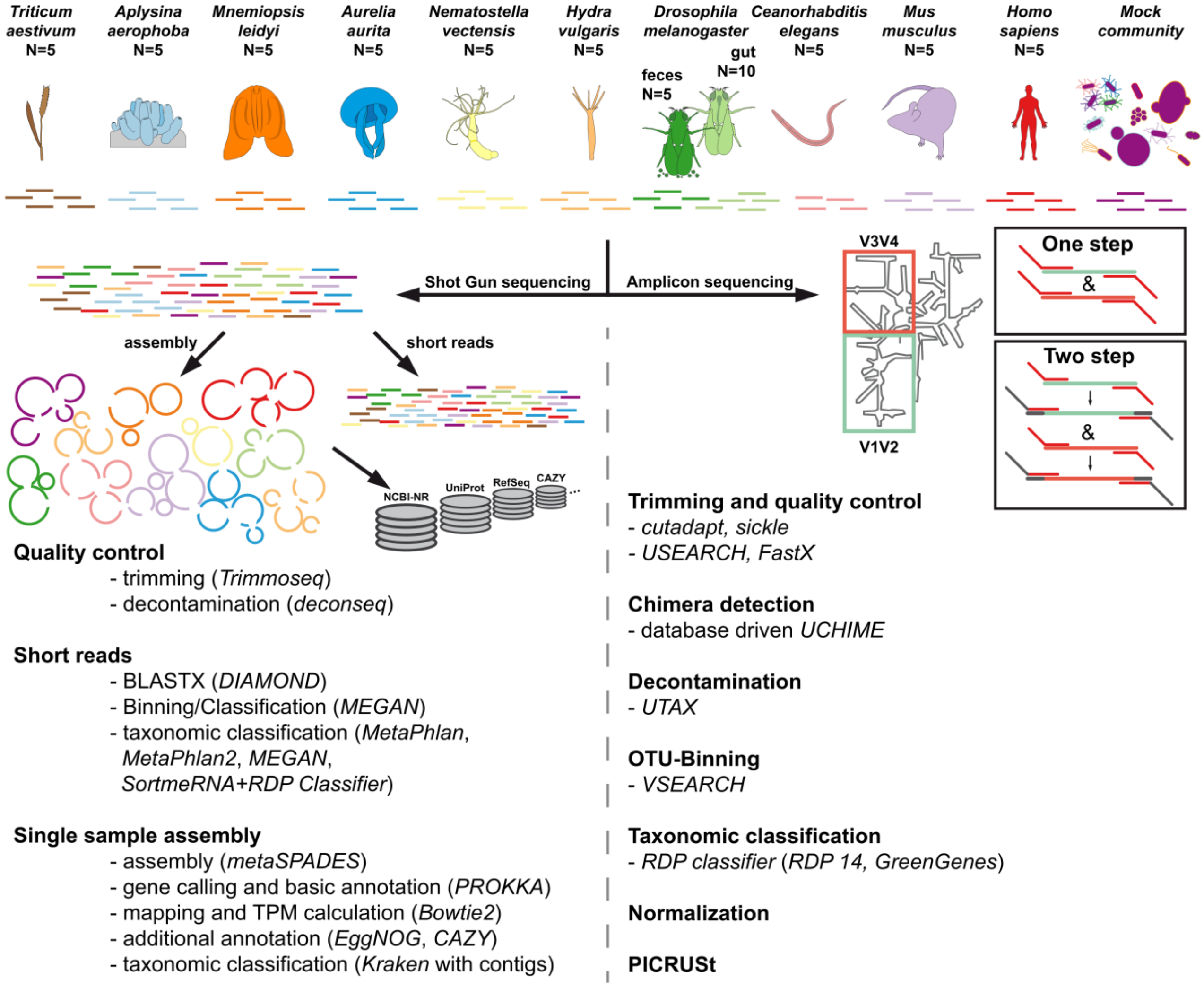


**Figure S1:** Visual abstract of the experiment, comparing different host organisms of the CRC 1182 investigated via different shotgun und 16S rRNA amplicon based methods (for final sample sizes see Table S1).

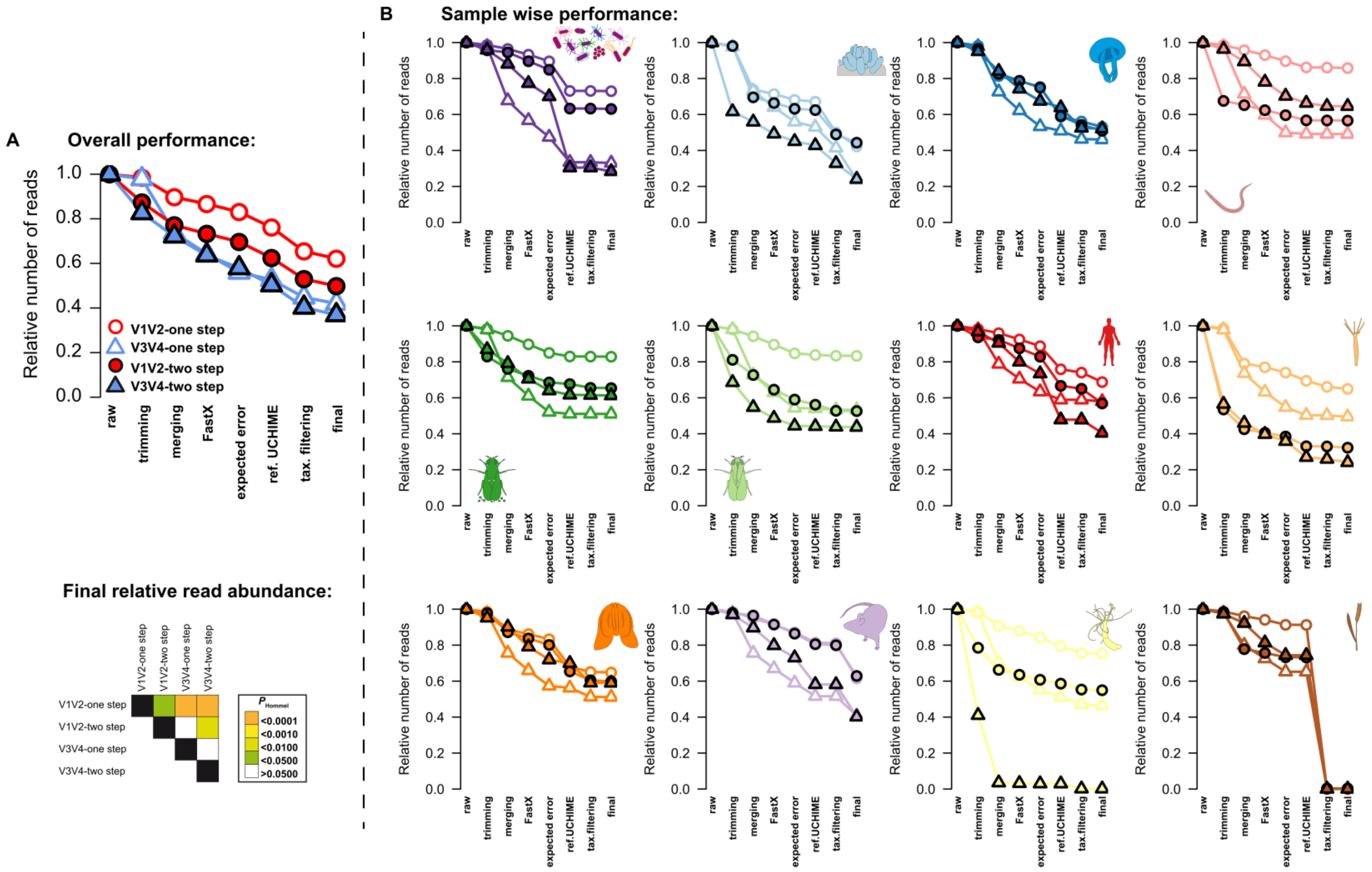


**Figure S2:** Average relative read count of the different 16S rRNA gene amplicon techniques (one step/two step, V1V2/V3V4) after each QC step employed in this study (for details please refer to the Methods section). (**A**) Average relative abundances of reads throughout the QC pipeline showing the significantly highest number of preserved reads in the V1V2-one step protocol (pairwise comparisons via pairwise *t*-Tests and Hommel *P*-value adjustment, significance levels are indicated by color). (**B**) Single plots display the average relative read counts across the different QC steps for each sample/host type (mock community, *A. aerophoba*, *A. aurita*, *C. elegans*, *D. melanogaster* feces, *D. melanogaster* gut tissue, *H. sapiens*, *H. vulgaris*, *M. leidyi*, *M. musculus*, *N. vectensis*, *T. aestivum*).

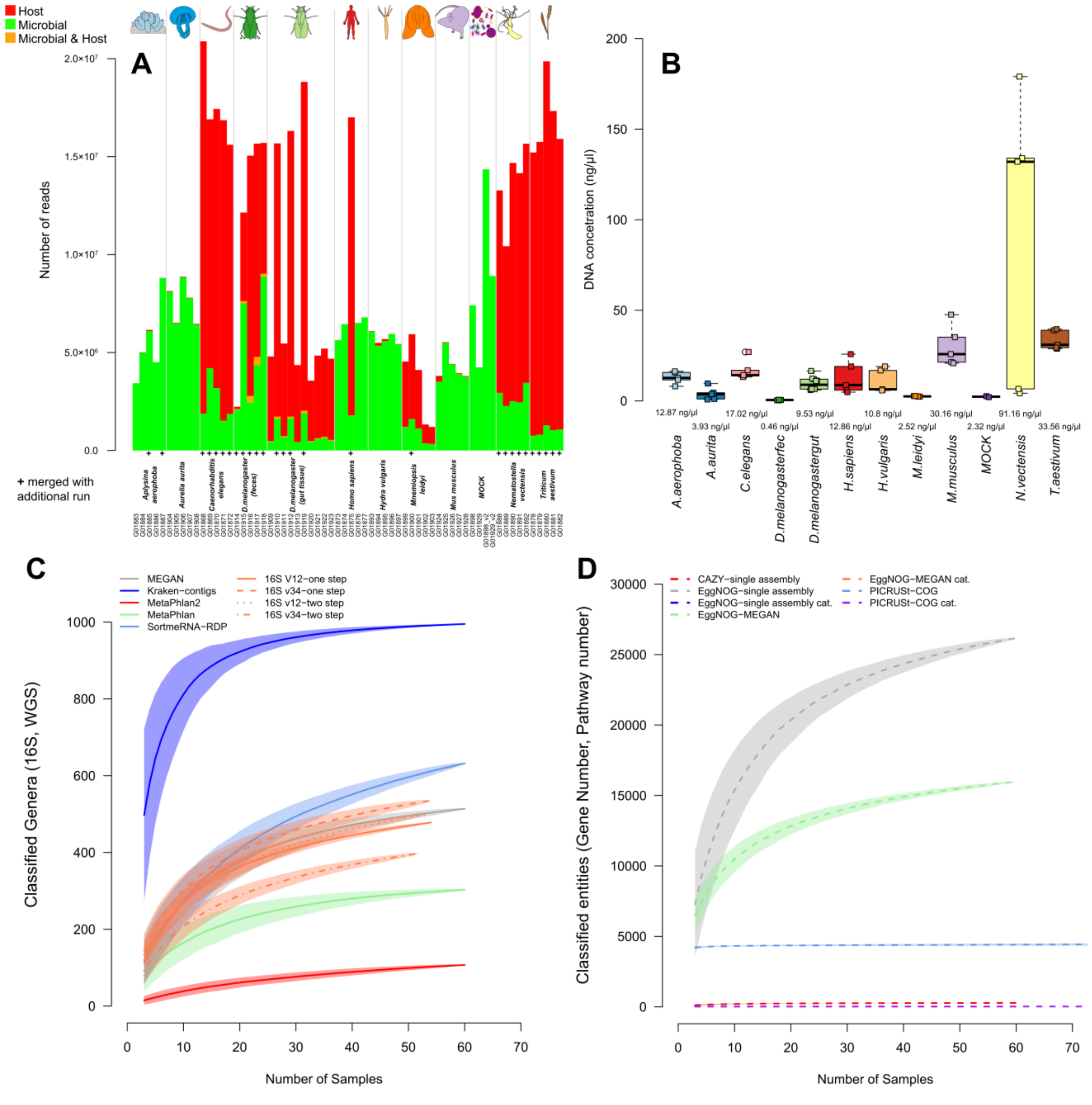


**Figure S3:** **(A)** Barplot displays the number of reads detected to be of host- or microbial origin for each sample. Resequenced samples are marked with a "+". (**B**) Average DNA concentration of samples (Qubit measurements). Collectors curves based on 1000 random re-samplings samples to see saturation of the number of **(C)** genera derived from shotgun and 16S rRNA gene amplicon techniques, as well as the number genes and functions **(D)** as derived from different shotgun based annotations and imputed functions (PICRUSt). Shading indicates standard deviations of the random samplings at each step (for sample sizes see Table S1).

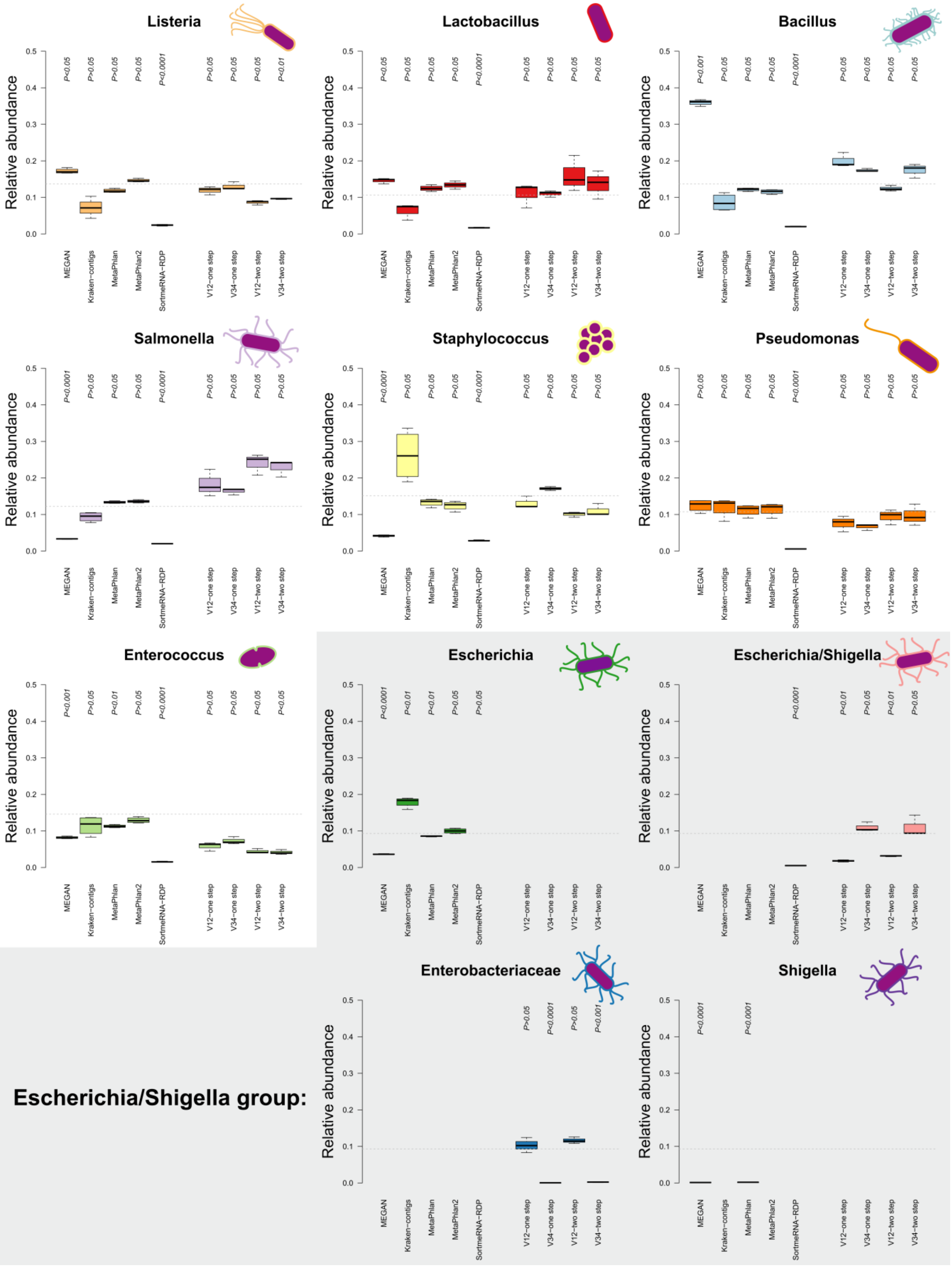
Listeria
Lactobacillus
Bacillus
Salmonella
Staphylococcus
Pseudomonas
Enterococcus
Escherichia
Escherichia/Shigella
Enterobacteriaceae
Shigella
Escherichia/Shigella group:
Relative abundance
0.0 0.1 0.2 0.3 0.4 0.5
MEGAN
Kraken−contigs
MetaPhlan
MetaPhlan2
SortmeRNA−RDP
V12−one step
V34−one step
V12−two step
V34−two step
Listeria: P<0.05 P>0.05 P>0.05 P>0.05 P<0.0001 P>0.05 P>0.05 P<0.05 P<0.01
Lactobacillus: P<0.05 P>0.05 P>0.05 P>0.05 P<0.0001 P>0.05 P>0.05 P>0.05 P>0.05
Bacillus: P<0.001 P>0.05 P<0.05 P>0.05 P<0.0001 P>0.05 P<0.05 P>0.05 P>0.05
Salmonella: P<0.0001 P>0.05 P<0.05 P>0.05 P<0.0001 P>0.05 P>0.05 P>0.05 P>0.05
Staphylococcus: P<0.0001 P>0.05 P>0.05 P>0.05 P<0.0001 P>0.05 P>0.05 P>0.05 P>0.05
Pseudomonas: P>0.05 P>0.05 P>0.05 P>0.05 P<0.0001 P>0.05 P>0.05 P>0.05 P>0.05
Enterococcus: P<0.001 P>0.05 P<0.01 P>0.05 P<0.0001 P>0.05 P>0.05 P<0.05 P<0.05
Escherichia: P<0.0001 P<0.01 P<0.01 P>0.05 P>0.05
Escherichia/Shigella: P<0.0001 P<0.01 P>0.05 P<0.01 P>0.05
Enterobacteriaceae: P>0.05 P<0.0001 P>0.05 P<0.0001
Shigella: P<0.0001 P<0.0001

**Figure S4:** Comparison of relative bacterial abundances of mock community samples to the expected relative abundances (dashed line) via one-sample Wilcoxon test (two-sided). Abundances are derived from 16S rRNA gene amplicon sequencing (V1V2, V3V4, one step, two step), MEGAN based classification (short reads), MetaPhlan (short reads), MetaPhlan2 (short reads), Kraken based classification (contigs), SortmeRNA (short reads). Sample sizes for the different approaches are $N_{shotgun}$=4, $N_{V1V2\text{-one step}}$=3, $N_{V1V2\text{-two step}}$=3, $N_{V3V4\text{-one step}}$=3, and $N_{V3V4\text{-two step}}$=3. *P*-values are corrected for multiple testing by Hommel *P*-value adjustment for each technique.

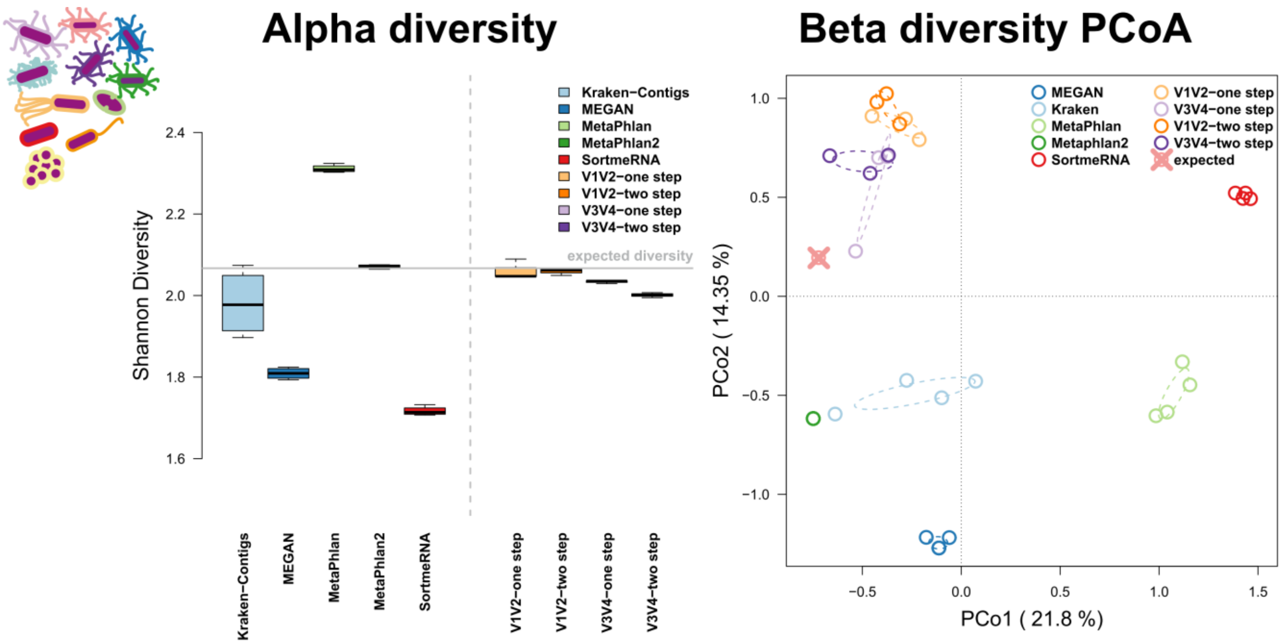


**Figure S5:** Bacterial alpha diversity Shannon H derived via different techniques in comparison to the expected community diversity. Principle Coordinate Analyses (PCoA) of community compositions based on the different shotgun and 16S rRNA gene amplicon techniques focusing on shared occurrences (Jaccard). Ellipses represent standard deviations of points within the respective groups. Sample sizes for the different approaches are $N_{shotgun}$=4, $N_{V1V2\text{-one step}}$=3, $N_{V1V2\text{-two step}}$=3, $N_{V3V4\text{-one step}}$=3, and $N_{V3V4\text{-two step}}$=3.

### Supplemental single host analyses:

#### Summary *Aplysina aerophoba* (Nardo, 1843):

*Aplysina aerophoba* is a Mediterranean-Atlantic member of the Verongida, which lives at light-exposed sites at depths from 5 to 15 m. *A. aerophoba*, as many demosponges, hosts a dense microbial community localized extracellularly in the mesohyl matrix which contributes to around one third of the sponge biomass [7, 8]. This sponge species serves as a model system to study basal host-microbial interactions and is further known for its biotechnological potential [9].

***Methods****:* Mediterranean *A. aerophoba* specimens were sampled offshore Girona, Spain, by scuba diving. Sponge specimens were rinsed with sterile sea water, fixed in RNAlater, and stored at -80°C. DNA was extracted from tissue via the Fast DNA Spin Kit for Soil (MP) with an additional ethanol precipitation step. No enrichment for bacterial DNA was used.

***Results****:* We see the highest alpha diversity in the shotgun profiles using Kraken, followed by MEGAN and SortmeRNA (Figure S6, left panel). The 16S rRNA gene amplicon profiles are more homogeneous and show a slightly lower diversity (Shannon index, Richness) for V1V2 compared to V3V4 and a slightly higher alpha diversity in samples amplified by the one step method. The principle coordinate analysis of the beta diversities show a clear separation of shotgun vs amplicon-sequenced samples based on abundance and co-occurrence of genera (Figure S6, middle panel) and displays clustering of MEGAN and Kraken based community profiles. Among the genera profiles derived from the 16S rRNA gene, there is no clear separation by variable region sequenced (*i.e.* V1V2 or V3V4) or amplification method. However, some samples based on the V3V4-two step method are clustering closely with MetaPhlan and MetaPhlan2 derived profiles (Figure S6, Middle panel). We can also identify a higher variation in the community profiles using derived from MetaPhlan and a very low variation in MetaPhlan2 which originates from its poor classification performance. In addition there is a noticeable higher variation detected in the V3V4 based profiles (Figure S6, right panel).

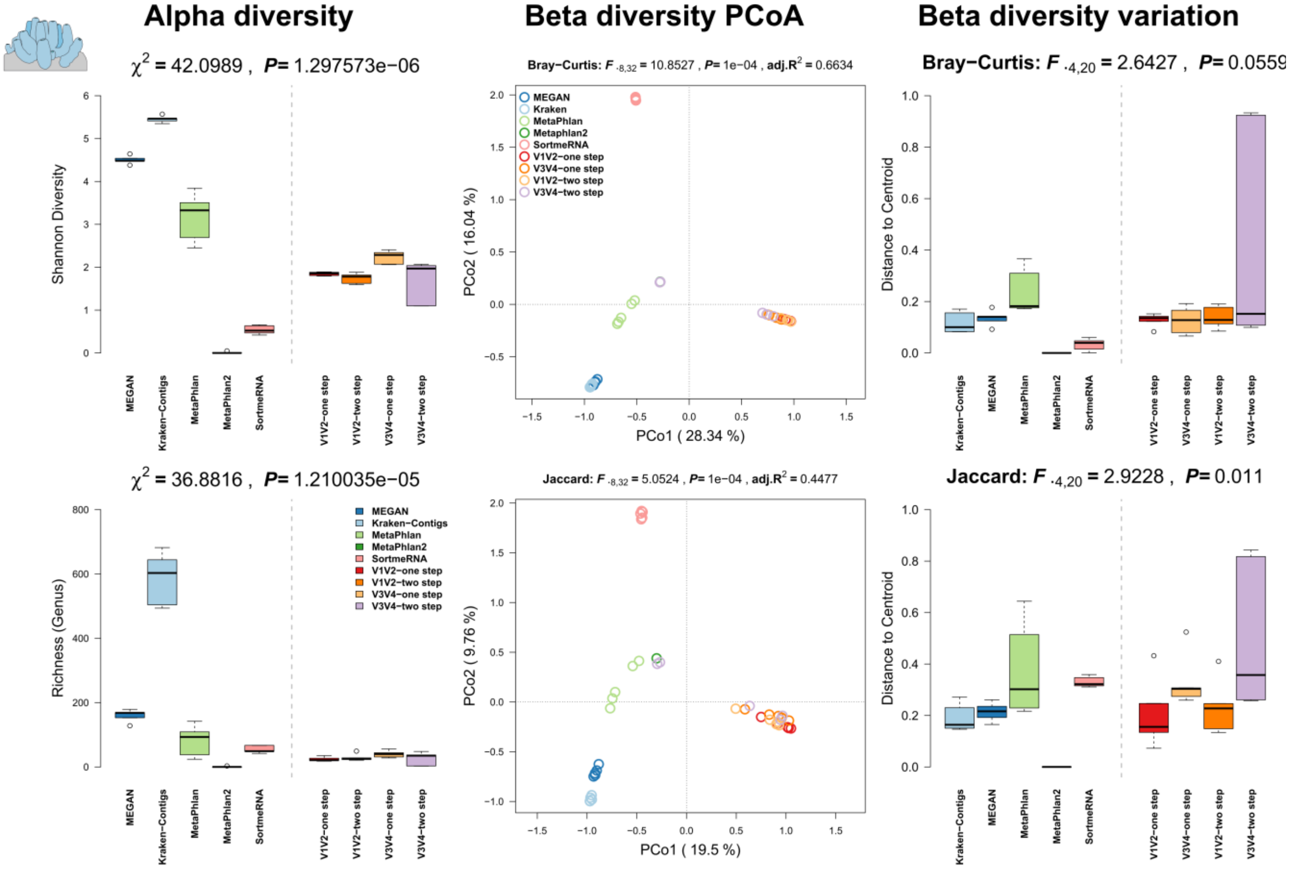


**Figure S6:** Comparison of community alpha diversity, composition, and variability of *Aplysina aerophoba* associated microbial communities among 16S rRNA gene amplicon and shotgun based pipelines. Comparisons are tested via approximate Kruskal-Wallis test [10] (alpha diversity), PERMANOVA [11, 12] (beta diversity), and permutative anova (variation of beta diversity). Community profiles were derived from 16S rRNA gene amplicon sequencing (V1V2, V3V4, one step, two step), MEGAN based classification (short reads), MetaPhlan (short reads), MetaPhlan2 (short reads), Kraken based classification (contigs), SortmeRNA (short reads). Sample sizes for the different approaches are $N_{shotgun}$=5, $N_{V1V2\text{-one step}}$=5, $N_{V1V2\text{-two step}}$=5, $N_{V3V4\text{-one step}}$=5, and $N_{V3V4\text{-two step}}$=5.

**Summary *Aurelia aurita* (Linnaeus, 1758):**

The moon jelly (*Aurelia aurita*) is a widely distributed pelagic scyphozoan found in almost all warm and temperate waters of coastal zones worldwide and is recognized as a key player in marine ecosystems [13]. Due to its ability to tolerate a wide range of environmental conditions, especially temperature and salinity, and its highly diverse food spectrum, it successfully colonizes different environments and often causes jellyfish blooms around the world [14].

***Methods**:* Individual *A. aurita* medusae (mean umbrella diameter 23 cm, N=5) were sampled from one location in the Eckernförder Bight, Baltic Sea (54.462654 N, 9.842743 E) in June 2016 by using a dip net. The animals were transported immediately to the laboratory, washed thoroughly with sterile filtered artificial seawater (ASW) to remove non-associated microbes. Pieces (2 × 2 cm) of the umbrella were cut out with a sterile scalpel. Dissociation of tissues was performed overnight at 4°C with 1 mg/mL collagenase (Sigma-Aldrich, St. Louis/USA). Homogenates were filtered through 10 µm Nylon gaze, followed by adding 0.1% IGEPAL CA-630 (Sigma-Aldrich, St. Louis/USA) and centrifugation of samples for 25 min at 300 × g at 4 °C. Supernatant including the prokaryotic fraction was centrifuged 5 min at 7,500 × g. DNA was extracted using the Wizard genomic purification kit (Promega, Madison, WI, USA). Pellets from eukaryotic/prokaryotic cell separation were homogenized in 480 µl 50mM EDTA and incubated at 37°C for 30 min after the addition of 10 mg/mL lysozyme (Carl Roth, Karlsruhe/Germany) and 60 Units Proteinase K (Life Technologies, Darmstadt/Germany). The remaining preparation steps were performed according to the manufacturer's protocol.

***Results****:* Alpha diversity estimates are mostly comparable between shotgun derived, (in particular Kraken, MEGAN, MetaPhlan) and the 16S rRNA gene amplicon -based community profiles (Figure S7, left panel). Kraken however, shows a far higher number of genera, which is evident in all other host species and is likely to be an artifact. The 16S rRNA gene amplicon profiles are homogeneous and only show a slightly higher diversity in the V3V4 based community profiles. The principle coordinate analyses show a relatively close clustering of MEGAN, Kraken and the V1V2 profiles based on shared abundances of genera, while there is a clear separation between shotgun- and amplicon-derived profiles when we consider the co-occurrence of bacterial genera among samples (Figure S7, middle panel). Within the 16S rRNA gene amplicon profiles, there is a separation by variable region sequenced based on abundance differences among samples (*i.e.* V1V2 or V3V4). When we focus on community variation we also see a higher variation in beta diversity noticeable in one-step PCR protocols compared to the two-step PCR protocol (Figure S7, right panel) and overall a relative comparable amount of variability detected by all methods, except SortmeRNA (Figure S7, right panel).

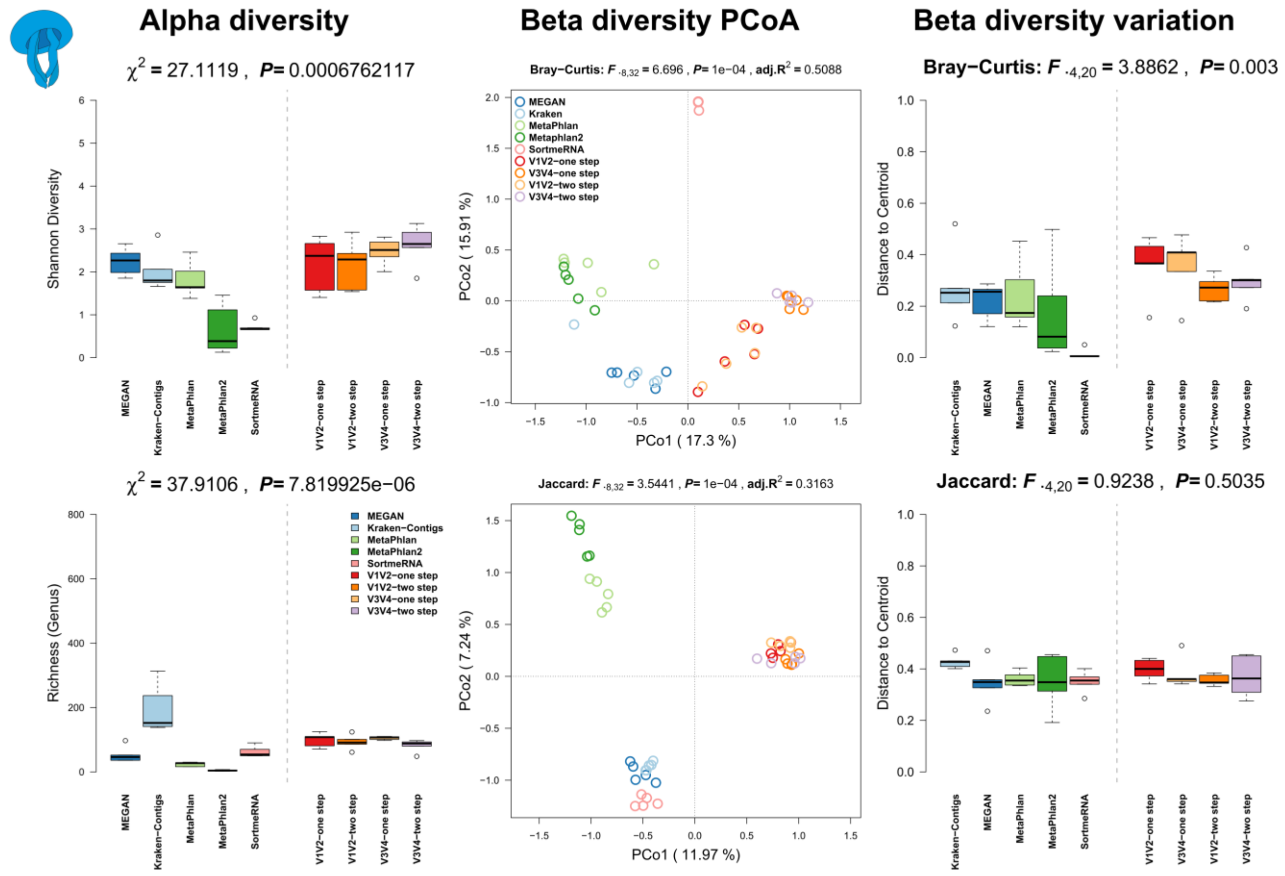


**Figure S7:** Comparison of community alpha diversity, composition, and variability of *Aurelia aurita* associated microbial communities among 16S rRNA gene amplicon and shotgun based pipelines. Comparisons are tested via approximate Kruskal-Wallis test (alpha diversity), PERMANOVA (beta diversity), and permutative anova (variation of beta diversity). Community profiles were derived from 16S rRNA gene amplicon sequencing (V1V2, V3V4, one step, two step), MEGAN based classification (short reads), MetaPhlan (short reads), MetaPhlan2 (short reads), Kraken based classification (contigs), SortmeRNA (short reads). Sample sizes for the different approaches are $N_{\text{shotgun}}$=5, $N_{\text{V1V2-one step}}$=5, $N_{\text{V1V2-two step}}$=5, $N_{\text{V3V4-one step}}$=5, and $N_{\text{V3V4-two step}}$=5.

**Summary *Caenorhabditis elegans* (Maupas, 1900):**

The nematode *Caenorhabditis elegans* provides a multitude of experimental advantages, such as small size, large-scale culturing, short generation time, transparency, genetic tractability, and thus, it has become one of the most widely used model organisms in biological research. In the laboratory *C. elegans* is almost exclusively cultivated mono-axenically on its food bacterium *E. coli* OP50. Therefore, almost all of the numerous studies with this nematode ignore a potential influence of the microbiome. In contrast, natural *C. elegans* are associated with a wide range of different microorganisms. Hence, wild caught and naturally colonized worms are of great interest to study the influence of a natural bacterial flora on nematode biology and fundamental developmental and genetic pathways.

***Methods**: C. elegans* were isolated directly from compost (n=2) and slugs found on the same compost (n=3) in the botanical garden in Kiel, Northern Germany (54°200N and 10°060E) in 2012, as previously described [15-17]. The samples were frozen in 30% glycerol-TSB and stored at -80°C. Worm samples were thawed and washed three times with M9 buffer with 0.05% Triton X-100 prior to DNA isolation [18]. DNA extractions from worm samples and a negative control (nuclease free water) were performed using the CTAB (Cetyl Trimethyl Ammonium Bromide) protocol as previously described [15, 17, 19]. No enrichment procedure for bacterial DNA was used.

***Results**:* We see the highest alpha diversity in the shotgun profiles using Kraken, followed by MetaPhlan, and the lowest diversity in profiles derived from MetaPhlan2 (Figure S8, left panel). The 16S rRNA gene amplicon profiles are quite homogeneous based on richness and complexity of their genus profiles. The principle coordinate analysis show no clear separation between shotgun- and amplicon-based profiles, which also strongly separate and vary within each method (see also Figure S8, right panel), in particular based on the abundance distribution of genera (Figure S8, middle panel). Co-occurrences of genera among samples show a degree of overlap between amplicon- and shotgun-derived profiles, except for MetaPhlan.

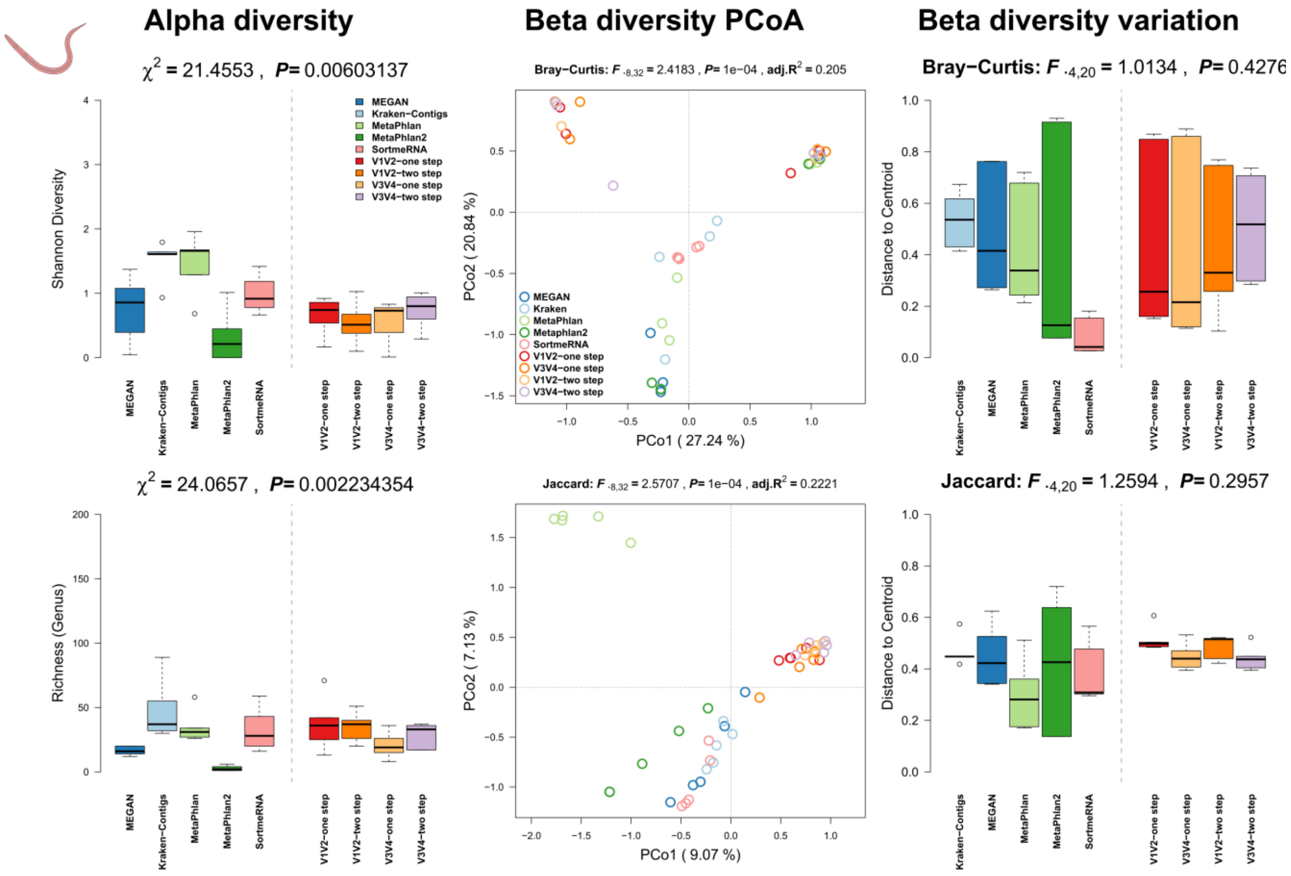


**Figure S8:** Comparison of community alpha diversity, composition, and variability of *Caenorhabditis elegans* associated microbial communities among 16S rRNA gene amplicon and shotgun based pipelines. Comparisons are tested via approximate Kruskal-Wallis test (alpha diversity), PERMANOVA (beta diversity), and permutative anova (variation of beta diversity). Community profiles were derived from 16S rRNA gene amplicon sequencing (V1V2, V3V4, one step, two step), MEGAN based classification (short reads), MetaPhlan (short reads), MetaPhlan2 (short reads), Kraken based classification (contigs), SortmeRNA (short reads). Sample sizes for the different approaches are $N_{shotgun}$=5, $N_{V1V2\text{-one step}}$=5, $N_{V1V2\text{-two step}}$=5, $N_{V3V4\text{-one step}}$=5, and $N_{V3V4\text{-two step}}$=5.

### Summary *Drosophila melanogaster* (Meigen, 1830):

*Drosophila melanogaster* is one of the best established model organisms for genetic and developmental investigations with a plethora of tools and protocols available for genetic, physiological and neurological manipulations and the first metazoan genome [20]. Due to its rather simple metagenome, its easy husbandry, and well established tools, *D. melanogaster* is becoming a model system for experimental microbial community analyses as well [21].

***Methods***: *D. melanogaster* (strain: $w^{1118}$) were sampled in different ways. Five samples of 5 and 10 female flies each were dissected as previously described and fecal spots of five independent culture vials were sampled and extracted with the MOBIO Soil Kit [22] to contrast the tissue associated and fecal microbial communities of *D. melanogaster*. No enrichment for bacterial DNA was used.

***Results****:* We see the highest alpha diversity in the shotgun profiles using Kraken, while all other methods clearly show lower diversity in the fecal, as well as gut samples of *D. melanogaster* (Figure S9, Figure S10, left panels). In the 16S rRNA gene amplicon profiles, the Shannon index is slightly higher for V1V2-two step compared to the other methods for amplicon generation. Principle coordinate analysis of the beta diversities shows overlap between 16S rRNA gene amplicon- and shotgun-derived profiles, while SortmeRNA and MetaPhlan are separated from those profiles in particular based on differences in the occurrences of genera (Figure S9, Figure S10, middle panels). There is slightly higher community variability in noticeable in amplicon- compared to shotgun-based profiles in the fecal spot samples, while in the gut samples we rather see an opposite trend (Figure S9, Figure S10, right panels).

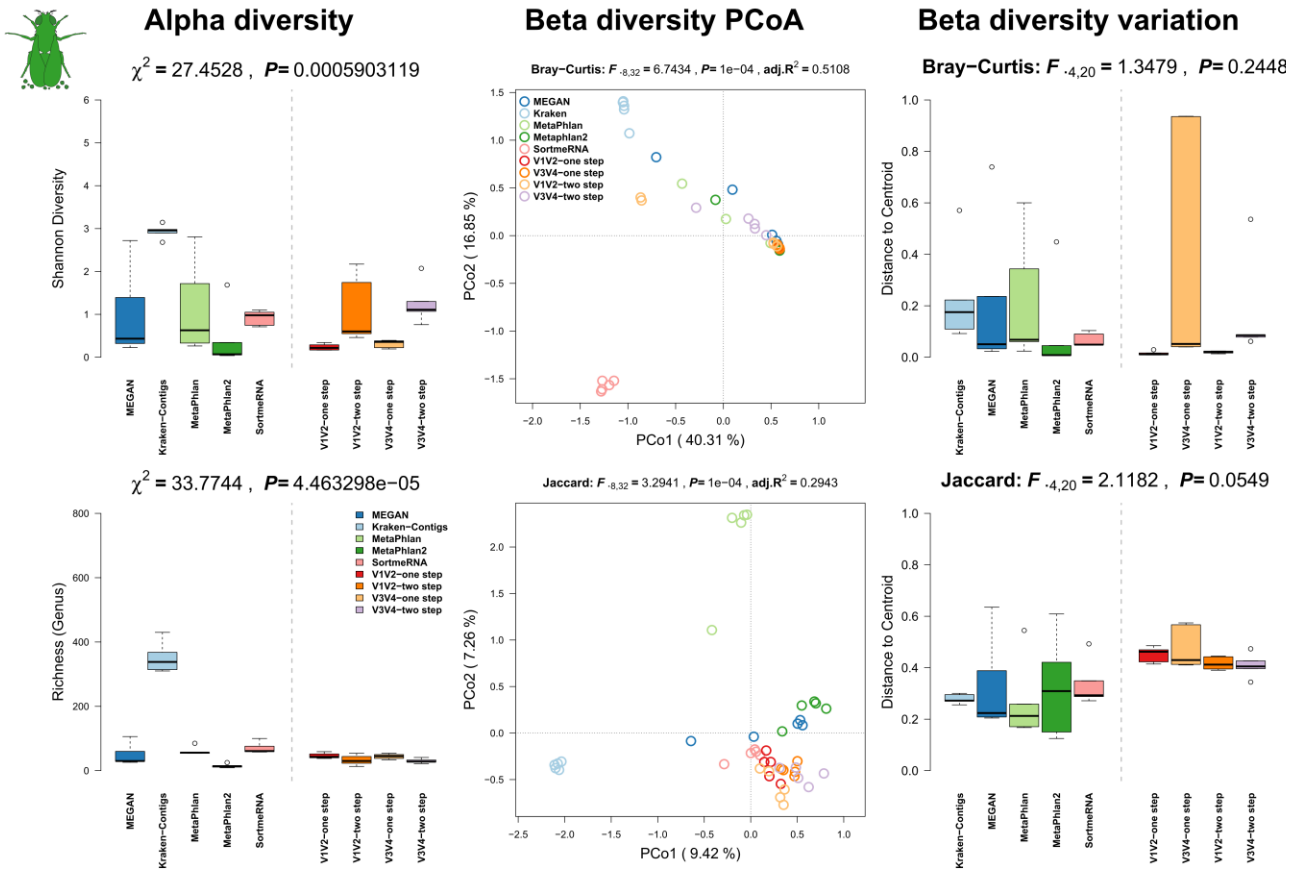


**Figure S9:** Comparison of community alpha diversity, composition, and variability of *Drosophila melanogaster* fecal microbial communities among 16S rRNA gene amplicon and shotgun based pipelines. Comparisons are tested via approximate Kruskal-Wallis test (alpha diversity), PERMANOVA (beta diversity), and permutative anova (variation of beta diversity). Community profiles were derived from 16S rRNA gene amplicon sequencing (V1V2, V3V4, one step, two step), MEGAN based classification (short reads), MetaPhlan (short reads), MetaPhlan2 (short reads), Kraken based classification (contigs), SortmeRNA (short reads). Sample sizes for the different approaches are $N_{shotgun}$=5, $N_{V1V2\text{-}one\ step}$=5, $N_{V1V2\text{-}two\ step}$=5, $N_{V3V4\text{-}one\ step}$=5, and $N_{V3V4\text{-}two\ step}$=5.

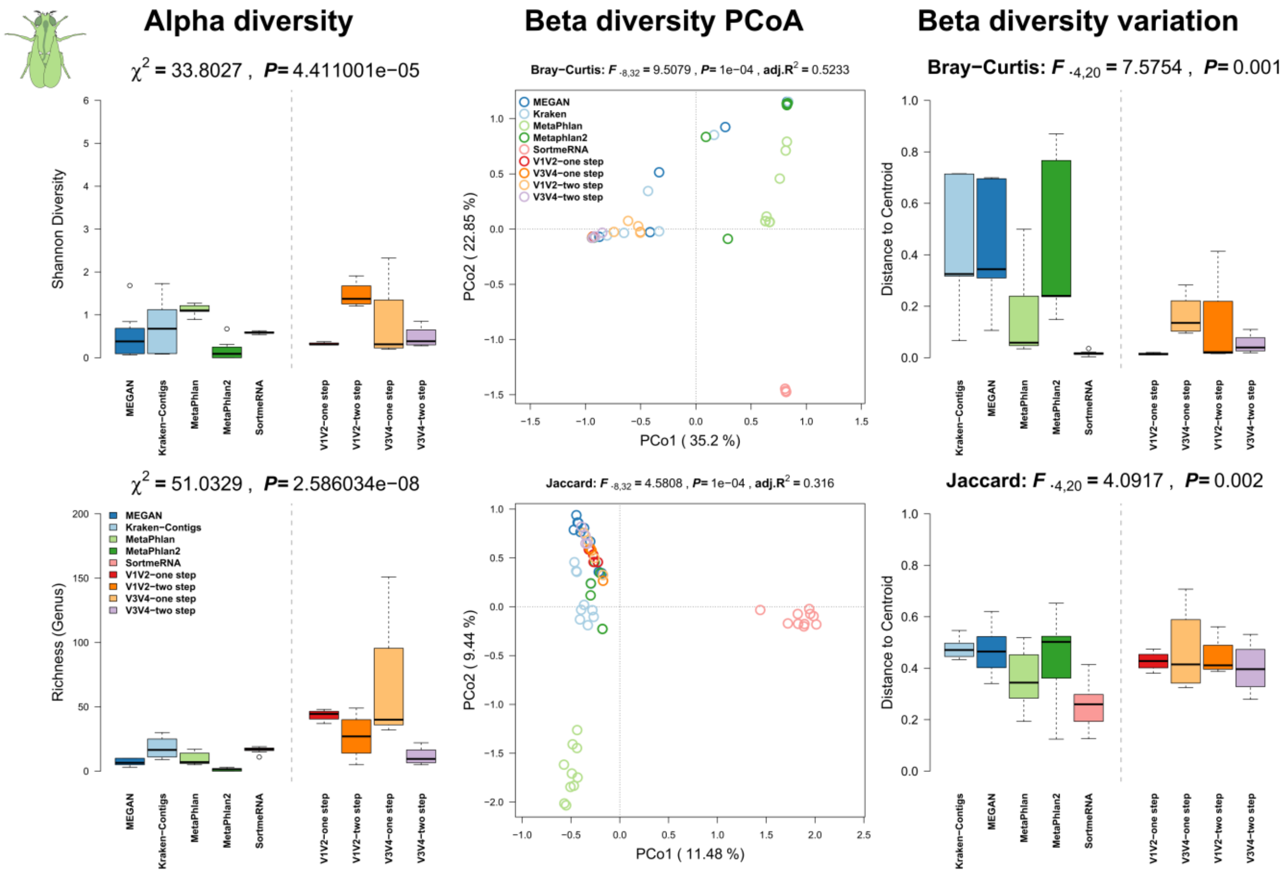


**Figure S10:** Comparison of community alpha diversity, composition, and variability of *Drosophila melanogaster* gut tissue associated microbial communities among 16S rRNA gene amplicon and shotgun based pipelines. Comparisons are tested via approximate Kruskal-Wallis test (alpha diversity), PERMANOVA (beta diversity), and permutative anova (variation of beta diversity). Community profiles were derived from 16S rRNA gene amplicon sequencing (V1V2, V3V4, one step, two step), MEGAN based classification (short reads), MetaPhlan (short reads), MetaPhlan2 (short reads), Kraken based classification (contigs), SortmeRNA (short reads). Sample sizes for the different approaches are $N_{shotgun}$=10, $N_{V1V2\text{-}one\ step}$=4, $N_{V1V2\text{-}two\ step}$=4, $N_{V3V4\text{-}one\ step}$=4, and $N_{V3V4\text{-}two\ step}$=4.

**Summary *Homo sapiens* (Linnaeus, 1758):**

The human (*Homo sapiens*) microbiome has been studied intensively and is presumed to be involved in wide array of diseases. Several microbiome studies have shown the influence of host genetics, environmental/life style variables, or diseases themselves [23, 24].

***Methods**:* Human feces (N=4) and biopsy samples (N=1) were sampled and extracted following the procedures as described in Wang *et al.* 2016 [23]. No enrichment for bacterial DNA was used.

***Results**:* We see again the highest alpha diversity in the shotgun profiles using Kraken, while all other methods in the shotgun and 16S rRNA gene amplicon profiles show very similar levels of complexity and richness. Interestingly, principle coordinate analyses show no clear separation between shotgun- and amplicon-derived profiles, except for SortmeRNA considering abundance and Kraken if we consider only occurrence of genera (Figure S11, middle panel). There is also no clear difference in community variation noticeable among methods (Figure S11, right panel).

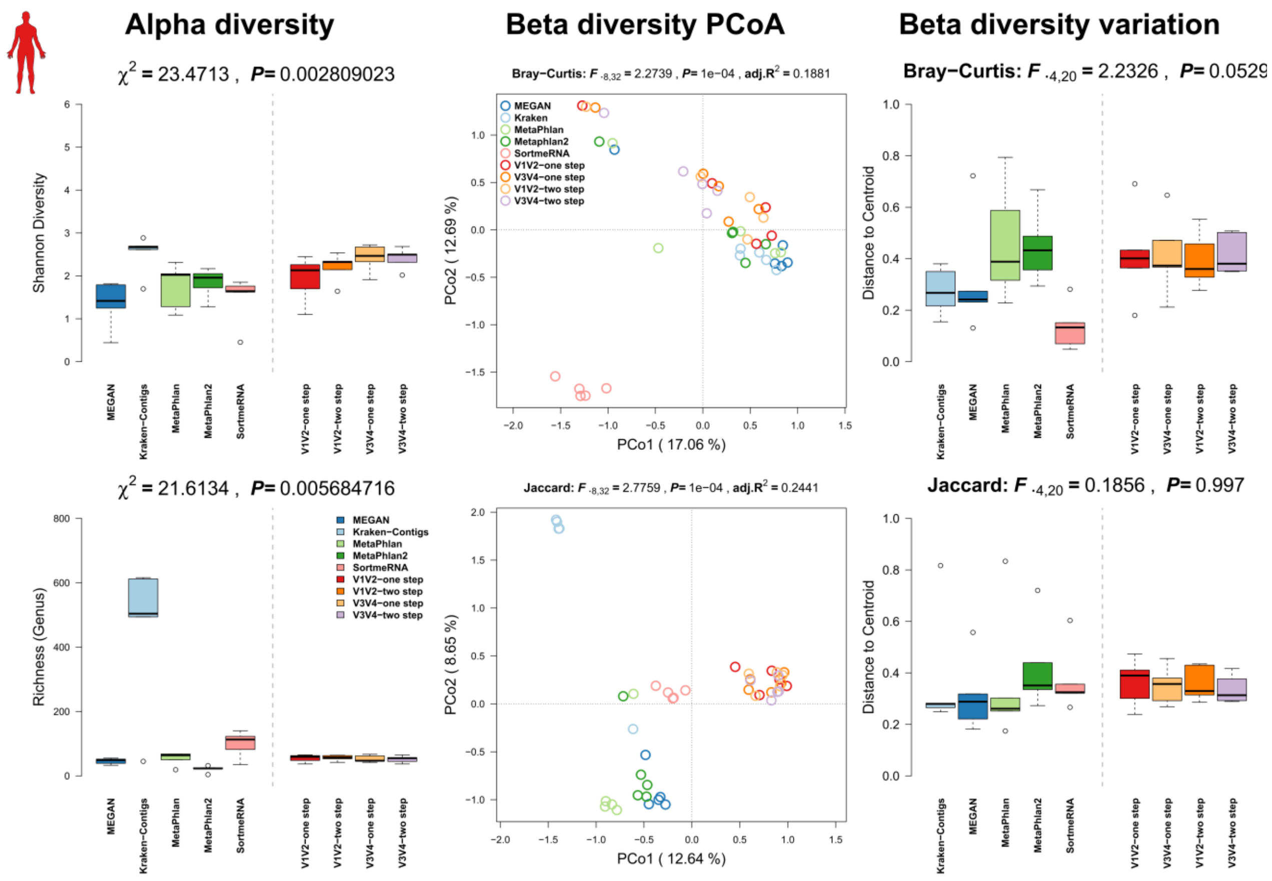


**Figure S11:** Comparison of community alpha diversity, composition, and variability of *Homo sapiens* associated microbial communities among 16S rRNA gene amplicon and shotgun based pipelines. Comparisons are tested via approximate Kruskal-Wallis test (alpha diversity), PERMANOVA (beta diversity), and permutative anova (variation of beta diversity). Community profiles were derived from 16S rRNA gene amplicon sequencing (V1V2, V3V4, one step, two step), MEGAN based classification (short reads), MetaPhlan (short reads), MetaPhlan2 (short reads), Kraken based classification (contigs), SortmeRNA (short reads). Sample sizes for the different approaches are $N_{shotgun}$=5, $N_{V1V2\text{-one step}}$=5, $N_{V1V2\text{-two step}}$=5, $N_{V3V4\text{-one step}}$=5, and $N_{V3V4\text{-two step}}$=5.

**Summary *Hydra vulgaris* (Pallas, 1766)*:***

The fresh-water cnidarian *Hydra vulgaris* is an established model organism in evolutionary developmental biology belonging to the Hydrozoa. Since *Hydra* is colonized by a stable and species specific bacterial community, it is also used for the study of host-microbe interactions [25]. Its body and tissue structure resembles in some ways vertebrate intestine with the endodermal epithelium and can be used to emulate dynamics as seen in vertebrates due to its ancestral relationship [26]. It also recently got a complete genome and has established methods for genetic manipulations [27, 28].

***Methods**:* Two hundred adult polyps of lab raised *H. vulgaris* were used for each sample and stored at -20°C until extraction via the QiaAMP stool kit following the manufacturer's instructions. All animals were cultured under constant, identical environmental conditions including culture medium, food (first-instar larvae of *Artemia nauplii*, fed three times per week) and temperature according to standard procedures. *H. vulgaris* polyps were cultured following standard operating procedure as described in Lenhoff & Brown (1970) [29]. No enrichment for bacterial DNA was used.

***Results**:* We see the highest alpha diversity in the shotgun profiles using Kraken, followed by MEGAN and MetaPhlan (Figure S12, left panel). In the 16S rRNA gene amplicon profiles, the Shannon index is slightly lower for V3V4 compared to V1V2. The principle coordinate analysis of the beta diversity shows no clear separation between shotgun- and amplicon-based profiles with the exception of SortmeRNA based profiles which are clearly separated based on abundances, while Kraken based profiles are strongly separated just based on differential occurrence of genera (Figure S12, middle panel). Co-occurrence also more clearly separates the amplicon- from the shotgun-derived profiles. Variation of communities is quite comparable among the different methods employed, however there is higher variation in V3V4 based samples (Figure S12, right panel).

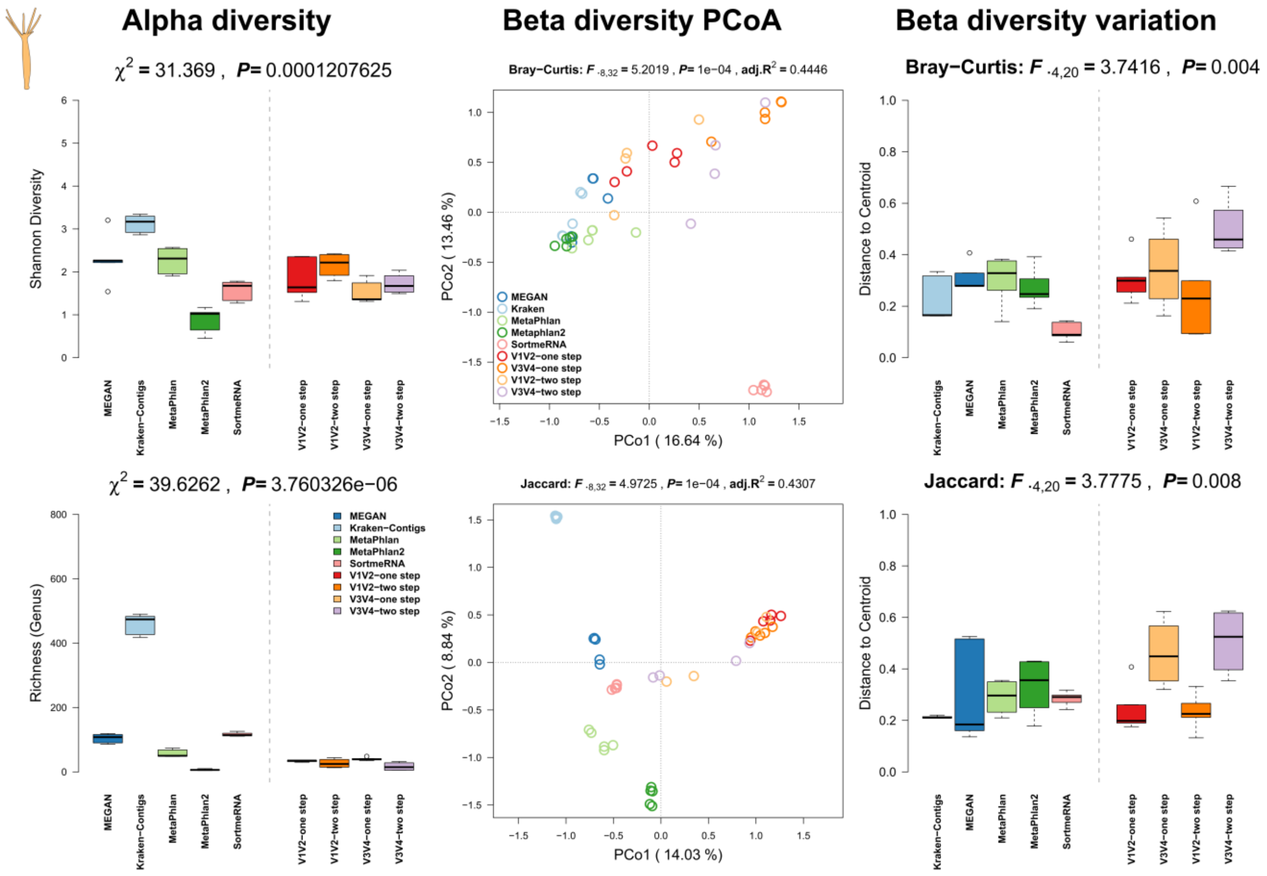


**Figure S12:** Comparison of community alpha diversity, composition, and variability of *Hydra vulgaris* associated microbial communities among 16S rRNA gene amplicon and shotgun based pipelines. Comparisons are tested via approximate Kruskal-Wallis test (alpha diversity), PERMANOVA (beta diversity), and permutative anova (variation of beta diversity). Community profiles were derived from 16S rRNA gene amplicon sequencing (V1V2, V3V4, one step, two step), MEGAN based classification (short reads), MetaPhlan (short reads), MetaPhlan2 (short reads), Kraken based classification (contigs), SortmeRNA (short reads). Sample sizes for the different approaches are $N_{shotgun}$=5, $N_{V1V2\text{-one step}}$=5, $N_{V1V2\text{-two step}}$=4, $N_{V3V4\text{-one step}}$=5, and $N_{V3V4\text{-two step}}$=4.

### Summary *Mnemiopsis leidyi* (Agassiz, 1865):

*Mnemiopsis leidyi* is a widely distributed lobate comb jelly (Ctenophora) native to the western Atlantic coasts. *M. leidyi* has become an ecologically and economically important invasive species as it recently expanded its range to the Black, Caspian, North- and Baltic Sea through human influence. *M. leidyi* was also established as model system to study regeneration, patterning, and luminescence, as well as the evolution of the metazoans. A recently finished genome is a recent addition to the set of resources available for *M. leidyi* [30].

***Methods**:* Individual *M. leidyi* (mean size 4 cm, N= 5) were sampled from one location in the Kiel Bight, Baltic Sea (54.330107 N, 10.149735 E) in September 2016 by using a dip net. The animals were transported immediately to the laboratory, washed thoroughly with sterile filtered artificial seawater (ASW) to remove non-associated microbes. Separation of eukaryotic and prokaryotic cells from whole animals as well as DNA extraction were performed as described in the *A. aurita* section.

***Results***: We can observe a drop out of MetaPhlan2, being unable to classify sequences obtained from *M. leidiy*. Furthermore we can observe the highest alpha diversity in the shotgun profiles using MEGAN and Kraken (Figure S13, left panel). In the 16S rRNA gene amplicon profiles we see a slightly higher diversity if V3V4 amplicons are analyzed as compared to V1V2. Principle coordinate analysis show a strong separation between methods, while particularly MEGAN, Kraken and samples based on V3V4 amplicons cluster close together. Profiles based on MetaPhlan, SortmeRNA, and V1V2 amplicons are strongly separated from each other (Figure S13, middle panel). MEGAN and Kraken show a higher variability in the abundance and co-occurrences in the community profiles, similar to the V1V2- and V3V4 two step amplicon methods (Figure S13, right panel).

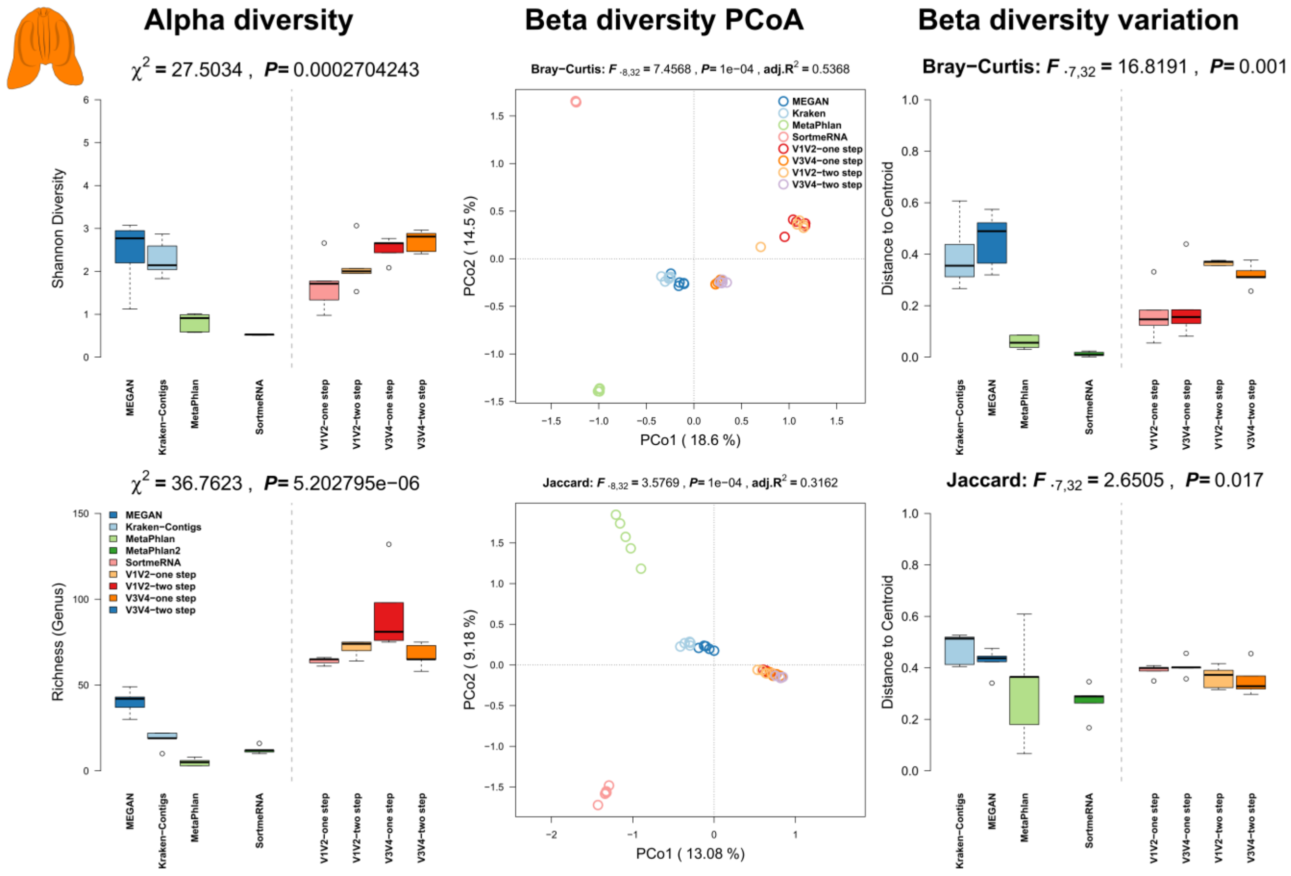


**Figure S13:** Comparison of community alpha diversity, composition, and variability of *Mnemiopsis leidyi* associated microbial communities among 16S rRNA gene amplicon and shotgun based pipelines. Comparisons are tested via approximate Kruskal-Wallis test (alpha diversity), PERMANOVA (beta diversity), and permutative anova (variation of beta diversity). Community profiles were derived from 16S rRNA gene amplicon sequencing (V1V2, V3V4, one step, two step), MEGAN based classification (short reads), MetaPhlan (short reads),

MetaPhlan2 (short reads), Kraken based classification (contigs), SortmeRNA (short reads). Sample sizes for the different approaches are $N_{shotgun}$=5, $N_{V1V2\text{-}one\ step}$=5, $N_{V1V2\text{-}two\ step}$=5, $N_{V3V4\text{-}one\ step}$=5, and $N_{V3V4\text{-}two\ step}$=5.

**Summary *Mus musculus* (Linnaeus, 1758):**

The house mouse (*Mus musculus*) is a widely used model organism in biology and medicine. As the mouse shares many similarities to humans in terms of anatomy, physiology and genetics, it has become the preferred mammalian model for genetic research, with a wide array of genetic and physiological tools available. It has also become a widely used model for microbiome studies.

***Methods:*** Five male hybrid mice were used for sampling. The mice originate from hybrid house mouse breeding stocks of wild derived *M. m. musculus* × *M. m. domesticus* hybrids captured in 2008, kept at the Max Planck Institute in Plön ($11^{th}$ lab generation at time of sampling). The stocks are derived from the Bavarian hybrid zone around Munich (Germany), in particular the locations FS (N=2), HA (N=1), and TU (N=2) [31]. Handling and killing of the mice was conducted according to the German animal welfare law and Federation of European Laboratory Animal Science Associations guidelines. All mice were sacrificed with $CO_2$ followed by cervical dislocation. Cecal content was preserved in 1ml of RNAlater and stored at -20°C after 12h at 5°C, following centrifugation and removal of supernatant (N=5). This was later extracted using the DNA/RNA AllPrep kit (Qiagen) according to the manufacture's protocol as described previously [32, 33], with an initial bead-beating step using Lysing Matrix E tubes (MP Biomedicals). No enrichment for bacterial DNA was used.

***Results:*** We see the highest alpha diversity in the shotgun profiles using Kraken, followed by MEGAN and MetaPhlan (Figure S14, left panel). In the 16S rRNA gene amplicon profiles, the Shannon index is slightly lower for V3V4 compared to V1V2. The principle coordinate analysis of the beta diversity shows a clear separation between shotgun- and amplicon-derived profiles (Figure S14, middle panel). Within the 16S rRNA gene amplicon profiles, there is a separation by variable region sequenced (*i.e.* V1V2 or V3V4). There is also a higher variation in beta diversity noticeable in one-step PCR compared to the two-step PCR protocol (Figure S14, right panel).

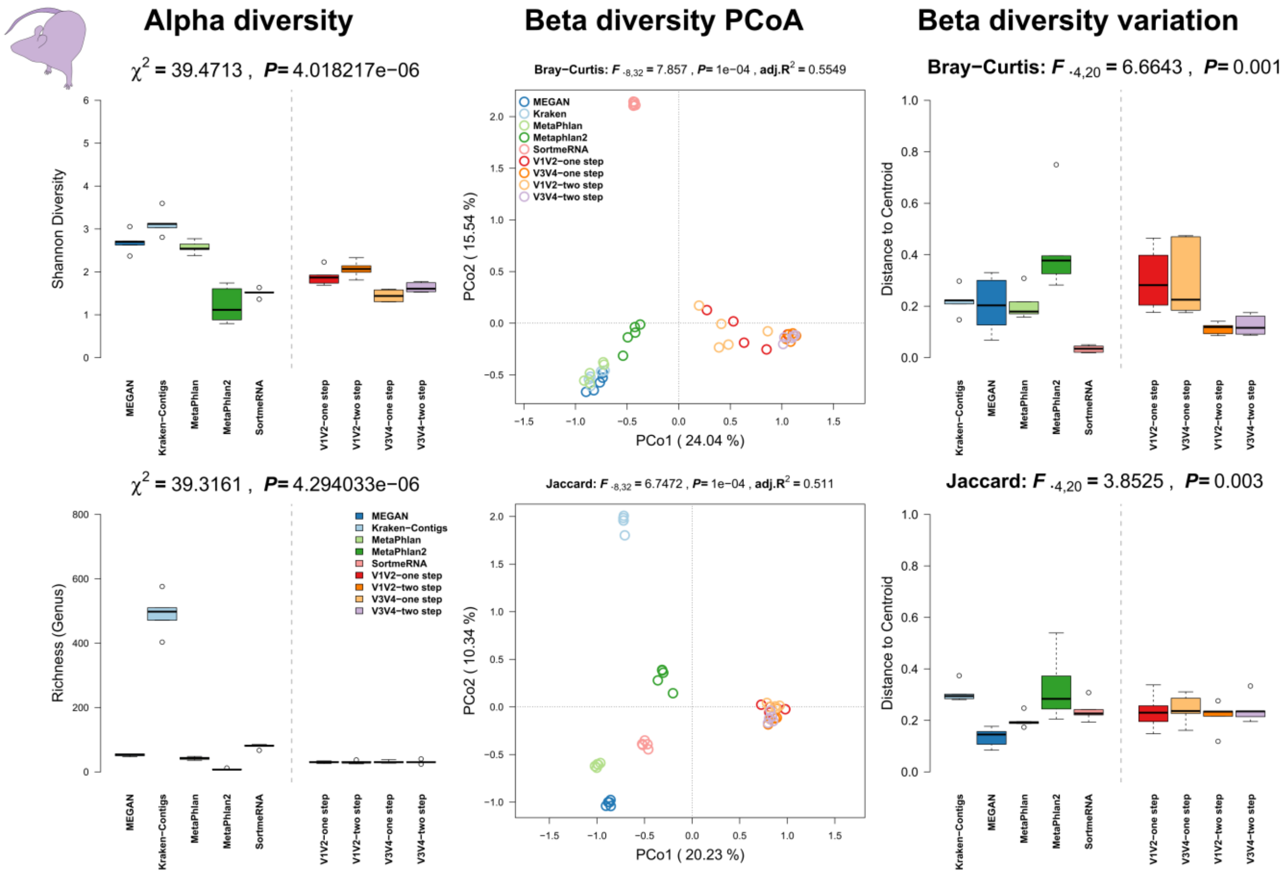


**Figure S14:** Comparison of community alpha diversity, composition, and variability of *Mus musculus* associated microbial communities among 16S rRNA gene amplicon and shotgun based pipelines. Comparisons are tested via approximate Kruskal-Wallis test (alpha diversity), PERMANOVA (beta diversity), and permutative anova (variation of beta diversity). Community profiles were derived from 16S rRNA gene amplicon sequencing (V1V2, V3V4, one step, two step), MEGAN based classification (short reads), MetaPhlan (short reads), MetaPhlan2 (short reads), Kraken based classification (contigs), SortmeRNA (short reads). Sample sizes for the different approaches are $N_{shotgun}$=5, $N_{V1V2\text{-one step}}$=5, $N_{V1V2\text{-two step}}$=5, $N_{V3V4\text{-one step}}$=5, and $N_{V3V4\text{-two step}}$=5.

**Summary *Nematostella vectensis* (Stephenson, 1935):**

The marine Starlet Sea Anemone *Nematostella vectensis* is a marine cnidarian (Anthozoa) living in the shallow coastal waters and marshes of Canada, the United States, and England. *N. vectensis* is a model organism for embryonic development and has an enormous adaptive potential to variable environmental factors. The animals can be cultured in the lab and reproduce sexually and asexually throughout the year.

***Methods**:* Five solitary adult lab raised polyps of *N. vectensis* were sampled and stored at -20°C until extraction via the QIAamp DNA Microbiome Kit following the manufacturer's instructions. All animals were cultured under constant, identical environmental conditions including culture medium (Artificial sea water, 16‰), food (first-instar larvae of *Artemia nauplii*, fed two times per week) and temperature according to standard procedures (18°C, in complete darkness). No enrichment for bacterial DNA was used.

***Results**:* In contrast to the other host organisms in this study we can see the highest alpha diversity in the one step amplicon methods compared to the rest of methods, followed by MEGAN and MetaPhlan (Figure S15, left panel). In the amplicon profiles V3V4-two step method produces the lowest alpha diversity comparable to estimates produced by MetaPhlan2. Principle coordinate analyses show only little differentiation between shotgun- and amplicon-sequenced samples with the exception of SortmeRNA and MetaPhlan (Figure S15, middle panel). Based on the co-occurrence of genera we can observe more differentiation between amplicon- and shotgun based profiles, however V3V4 communities still cluster close to the shotgun based communities. There is also a higher variation in beta diversity noticeable in the V3V4-two step PCR, as well as in Kraken and MEGAN based community profiles (Figure S15, right panel).

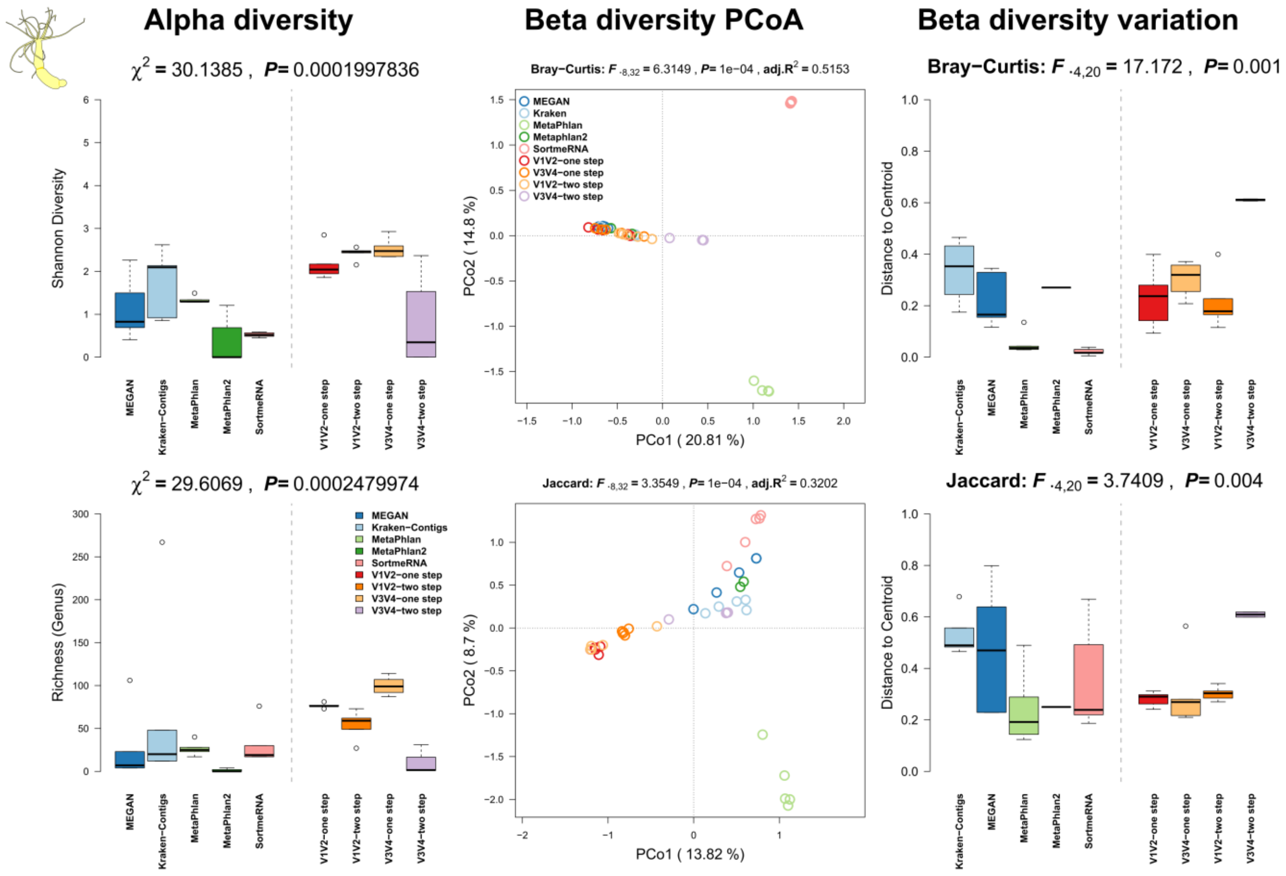


**Figure S15:** Comparison of community alpha diversity, composition, and variability of *Nematostella vectensis* associated microbial communities among 16S rRNA gene amplicon and shotgun based pipelines. Comparisons are tested via approximate Kruskal-Wallis test (alpha diversity), PERMANOVA (beta diversity), and permutative anova (variation of beta diversity). Community profiles were derived from 16S rRNA gene amplicon sequencing (V1V2, V3V4, one step, two step), MEGAN based classification (short reads), MetaPhlan (short reads), MetaPhlan2 (short reads), Kraken based classification (contigs), SortmeRNA (short reads).

Sample sizes for the different approaches are $N_{shotgun}$=5, $N_{V1V2\text{-}one\ step}$=5, $N_{V1V2\text{-}two\ step}$=5, $N_{V3V4\text{-}one\ step}$=5, and $N_{V3V4\text{-}two\ step}$=4.

**Summary *Triticum aestivum* (Linnaeus, 1758):**

*Triticum aestivum* a member of the *Poaceae* is the ancestral hybrid of *Triticum dicoccum* and *Aegilops tauschii* and, potentially first cultured 7000 yr. a.d. in the fertile crescent. It is one of the most widely distributed crops, used for bread, cereal and starch. Due to its industrial use it is of high genetic homogeneity. Grains and their associated microbial communities might be beneficial and interact during germination, as well as be a direct route of heritability from a fully grown plant to the next generation via the seed [34].

***Methods**:* DNA was extracted from grains (0.1 gram) of commercially purchased *Triticum aestivum* (Davert GmbH, brandname: Bioland Davert) (N=2) and farm raised *T. aestivum* (N=3, University farm CAU Kiel, Runal cultivar). 0.1 gram of seeds was washed in 70% ethanol and surface sterilized in 3% Sodium hypochlorite for 10 minutes. Afterwards grains were washed with sterile water. Surface sterilized grains were homogenized using Precellys (SK38; according to manufacturer's protocol). Homogenate was further processed using GeneJet Plant Genomic DNA Purification Kit (ThermoFisher Scientific). Quality of extracted DNA was tested on agarose gel. Presence of the bacterial 16S rRNA gene was confirmed by PCR using bacterila specific primers with less affinity to chloroplast signiatures (799F-1391R). No enrichment for bacterial DNA was used.

***Results**:* Although a high percentage of sequences has been lost due to host sequence contamination, we could still perform the majority of analyses. This also let in part to the drop out of MetaPhlan2 in the shotgun analyses. The highest alpha diversity has been recovered in the 16S rRNA gene amplicon based samples, particularly in the V1V2-two step (also highest richness) and V3V4-one step based amplicon samples. In the shotgun based community profiles we see the highest alpha diversity in Kraken, followed by MEGAN (Figure S16, left panel). The principle coordinate analysis of the beta diversity shows a clear separation between shotgun- and amplicon-derive profiles, with the strongest separation of MetaPhlan and SortmeRNA to the rest of samples (Figure S16, middle panel). Within the 16S rRNA gene amplicon profiles, there is a separation by variable region sequenced (*i.e.* V1V2 or V3V4). When only co-occurrences of taxa among samples is considered (Jaccard) we can see strong clustering of amplicon- and the shotgun based MEGAN and Kraken based profiles. On average the highest variability of communities can be observed among the 16S rRNA gene amplicon samples (Figure S16, right panel).

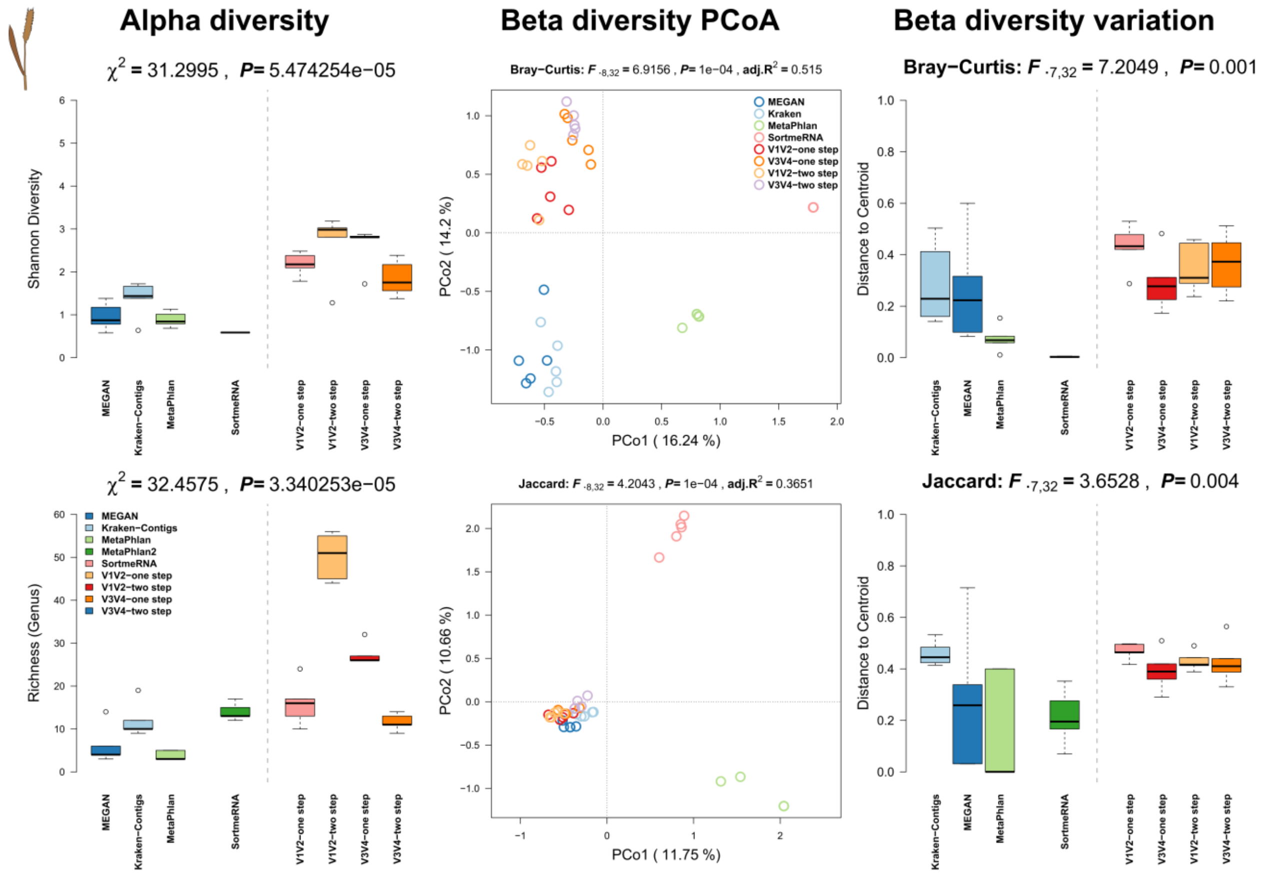


**Figure S16:** Comparison of community alpha diversity, composition, and variability of *Triticum aestivum* associated microbial communities among 16S rRNA gene amplicon and shotgun based pipelines. Comparisons are tested via approximate Kruskal-Wallis test (alpha diversity), PERMANOVA (beta diversity), and permutative anova (variation of beta diversity). Community profiles were derived from 16S rRNA gene amplicon sequencing (V1V2, V3V4, one step, two step), MEGAN based classification (short reads), MetaPhlan (short reads), MetaPhlan2 (short reads), Kraken based classification (contigs), SortmeRNA (short reads). Sample sizes for the different approaches are $N_{shotgun}$=5, $N_{V1V2\text{-one step}}$=5, $N_{V1V2\text{-two step}}$=5, $N_{V3V4\text{-one step}}$=5, and $N_{V3V4\text{-two step}}$=5.

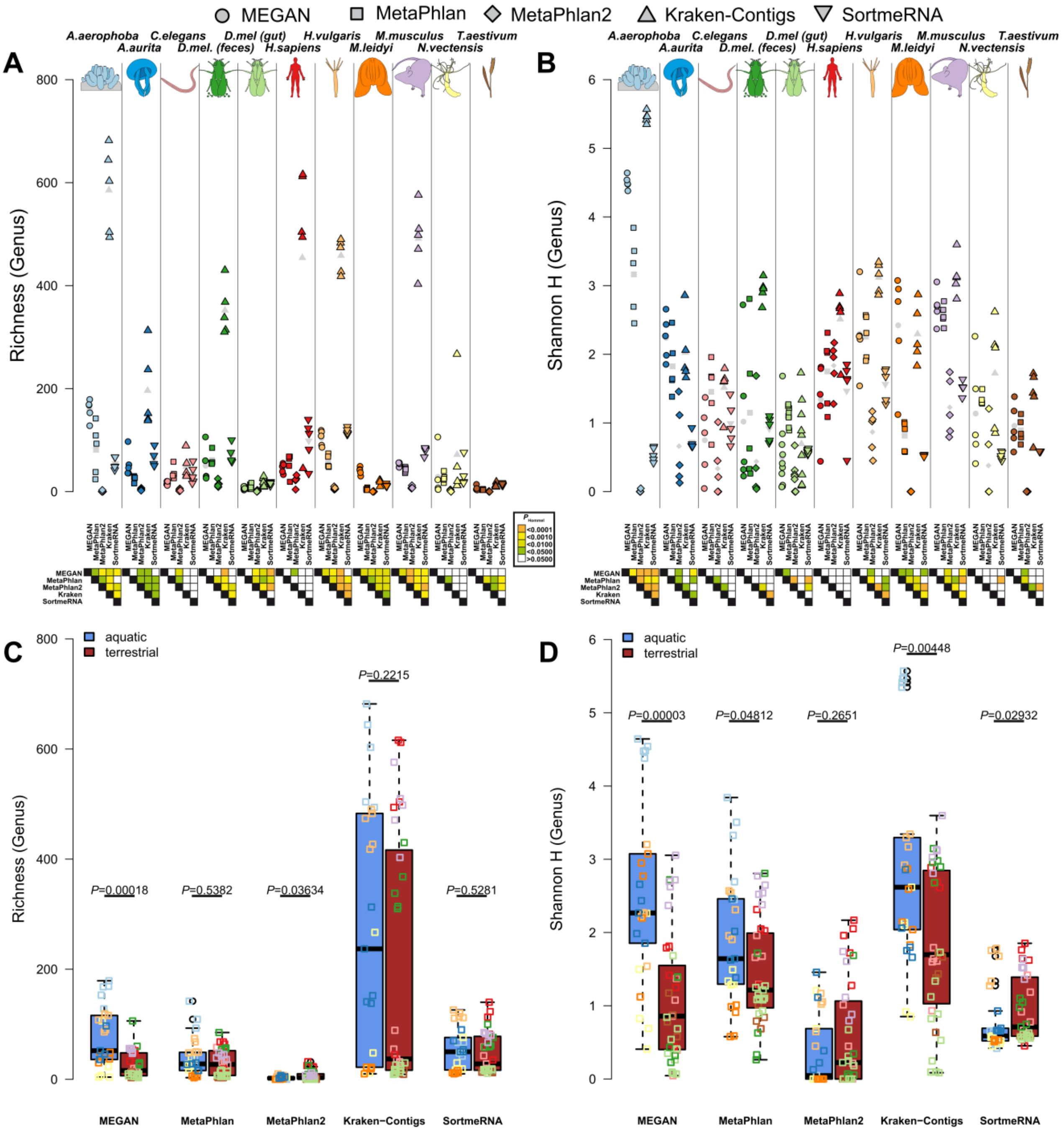


**Figure S17:** Comparison of bacterial community diversity estimates derived from different shotgun based classification pipelines. Pairwise comparisons are made among the different methods employed for each single host organism, via pairwise *t*-Tests without pooled standard variations, which show high heterogeneity of techniques within the host organisms. Mean values are shown in grey symbols in plots A and B. Significance levels after correction for multiple testing are indicated by color. Globally Shannon H differs between all methods except between MEGAN and MetaPhlan (LMM: $F_{4,285}$=58.2731, $P$<0.0001; random factor: organism) while in the case of richness Kraken shows a consistently higher diversity than all other techniques, as well as SortmeRNA compared to classifications based on MetaPhlan2 (LMM: $F_{4,285}$=58.2659,

$P<0.0001$; random factor: organism). Sample size for the host taxa is N=5, except for *D. melanogaster* gut tissue (N=10). The sample size of aquatic hosts is N=25, while terrestrial hosts have a sample size of N=35 (see Table S1).

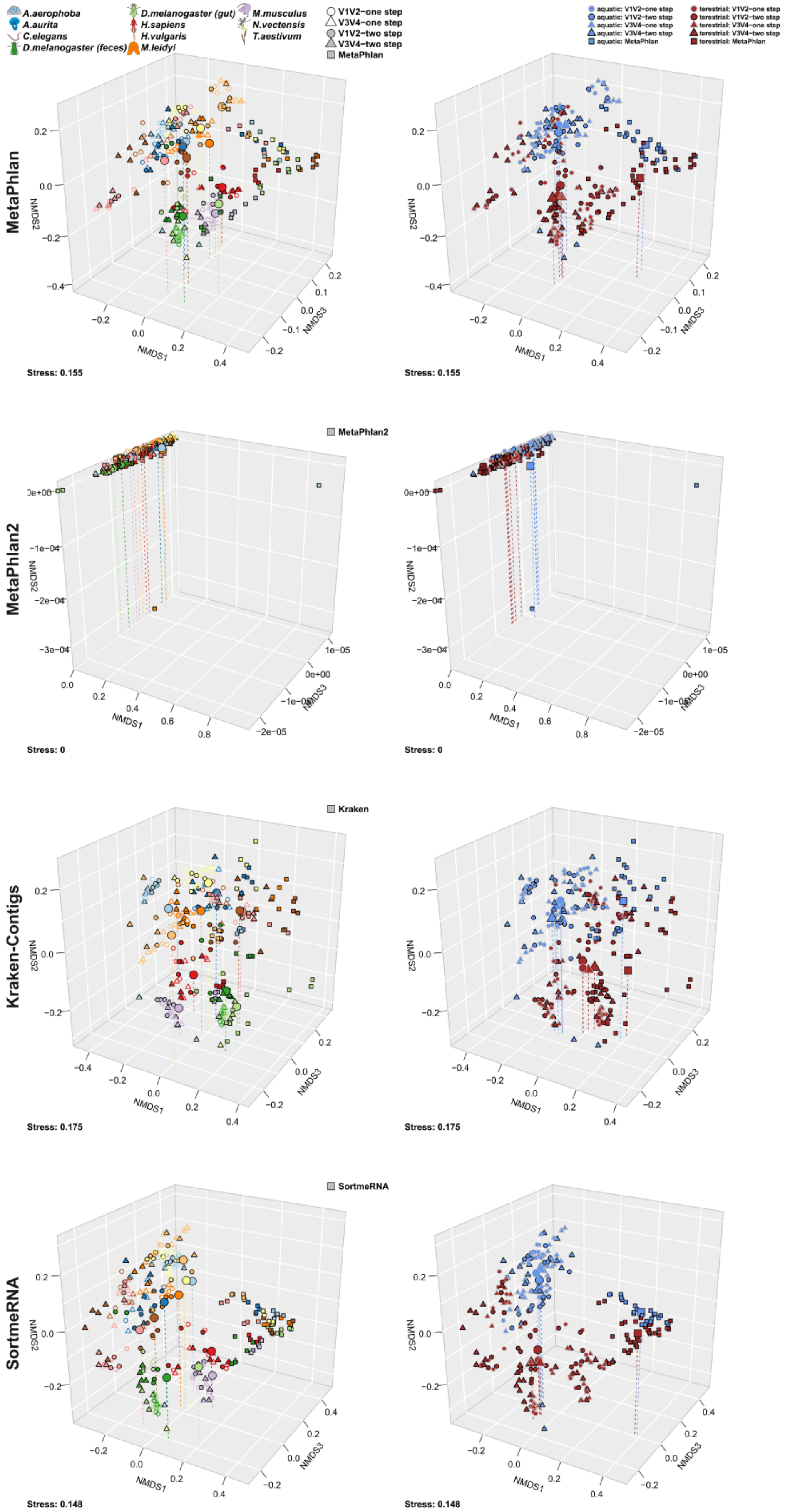
A.aerophoba
A.aurita
C.elegans
D.melanogaster (feces)
D.melanogaster (gut)
H.sapiens
H.vulgaris
M.leidyi
M.musculus
N.vectensis
T.aestivum
V1V2−one step
V3V4−one step
V1V2−two step
V3V4−two step
MetaPhlan
aquatic: V1V2−one step
aquatic: V1V2−two step
aquatic: V3V4−one step
aquatic: V3V4−two step
aquatic: MetaPhlan
terestrial: V1V2−one step
terestrial: V1V2−two step
terestrial: V3V4−one step
terestrial: V3V4−two step
terestrial: MetaPhlan
MetaPhlan
NMDS1
NMDS2
NMDS3
Stress: 0.155
Stress: 0.155
MetaPhlan2
MetaPhlan2
Stress: 0
Stress: 0
Kraken-Contigs
Kraken
Stress: 0.175
Stress: 0.175
SortmeRNA
SortmeRNA
Stress: 0.148
Stress: 0.148

**Figure S18:** Non-metric Multidimensional Scaling of Bray-Curtis distances derived from abundance differences of functional components highlighting functional differences between the host organisms (**A, C, E, G; see Table 3**) and host environments (**B, D, F, H; see Table 3**). Large symbols indicate the centroid of the respective host groups and vertical lines help to determine their position in space. Functional variation of communities based on pairwise Bray-Curtis distances within host organism groups and environmental groups. The sample size of aquatic hosts is N=25, while terrestrial hosts have a sample size of N=35 (see Table S1).

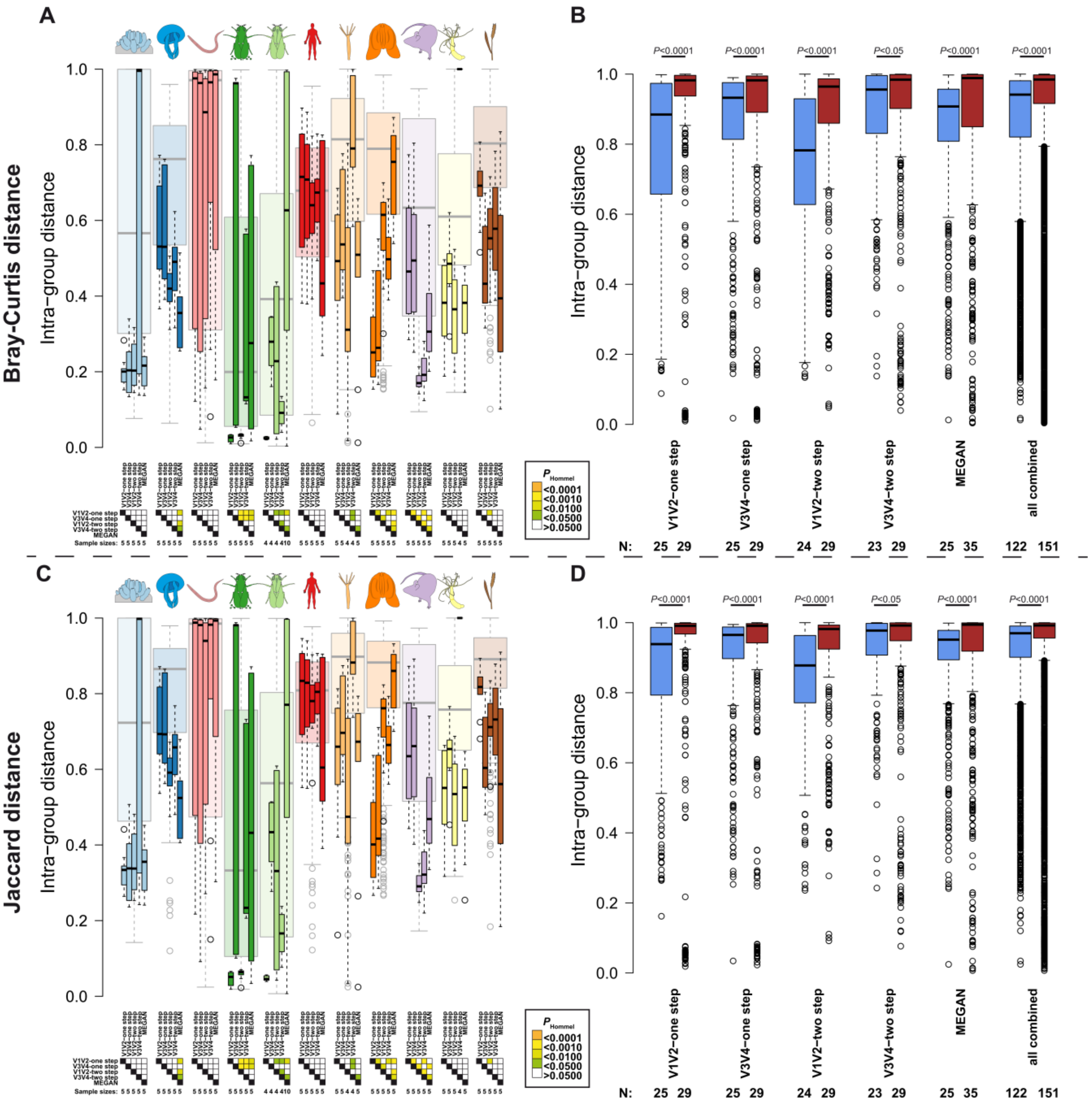

**Figure S19:** Boxplots visualize the pairwise distances based on shared abundance (**A, B**) and shared presence (**C, D**) between samples belonging to the respective host groups and methods (**A, C**), and host environments (**B, D**) among the different sequencing methods. Pairwise comparisons were computed via pairwise Wilcoxon tests (Hommel *P*-value adjustment) within host groups, and approximate Wilcoxon tests [10] between host environments. Significance levels after correction for multiple testing are indicated by color (for sample sizes see Table S1).

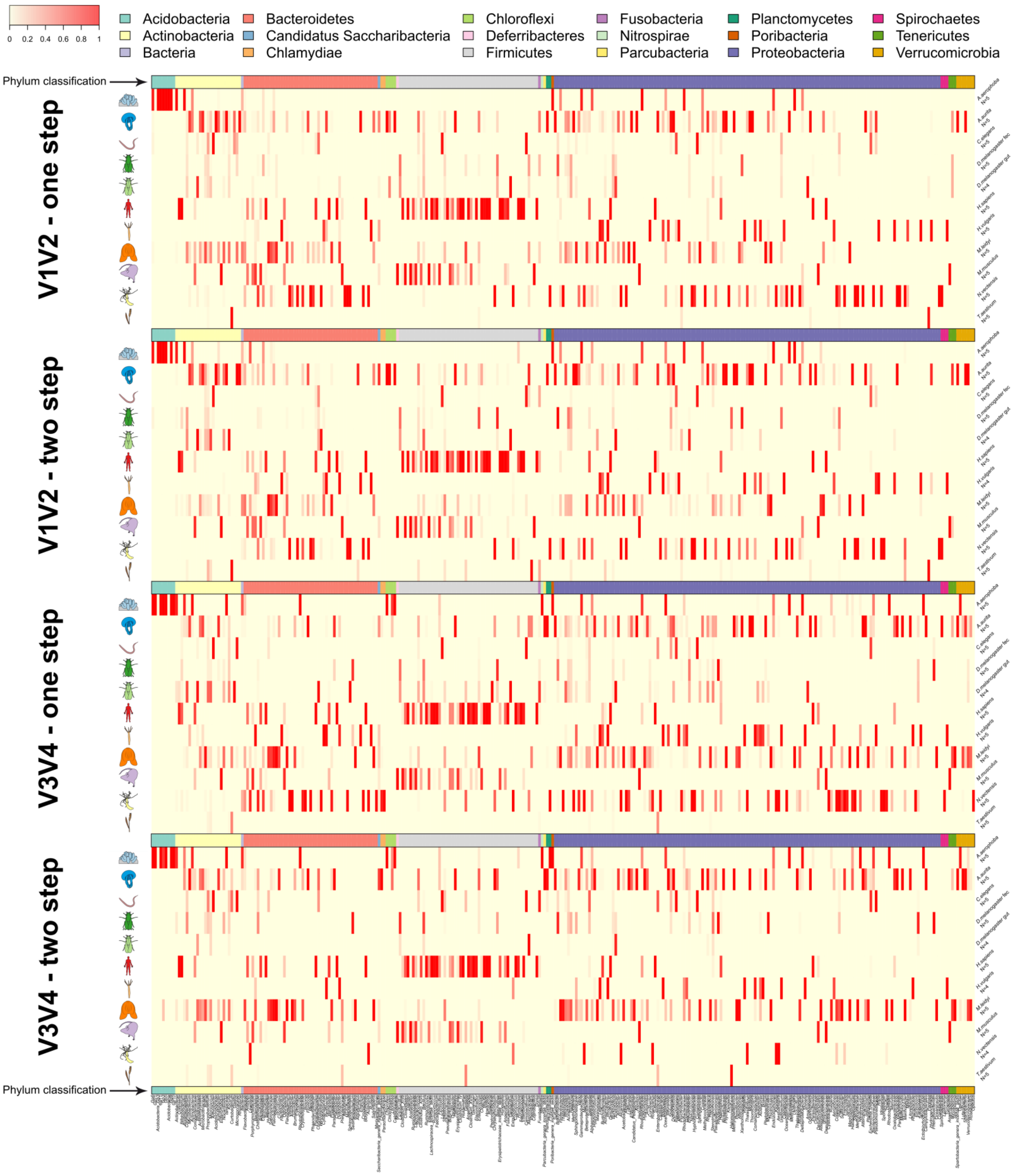

0 0.2 0.4 0.6 0.8 1
Acidobacteria
Actinobacteria
Bacteria
Bacteroidetes
Candidatus Saccharibacteria
Chlamydiae
Chloroflexi
Deferribacteres
Firmicutes
Fusobacteria
Nitrospirae
Parcubacteria
Planctomycetes
Poribacteria
Proteobacteria
Spirochaetes
Tenericutes
Verrucomicrobia
Phylum classification
V1V2 - one step
V1V2 - two step
V3V4 - one step
V3V4 - two step
Phylum classification

**Figure S20:** Heatmaps of indicator genera derived from the different 16S rRNA gene amplicon strategies. Heat color indicates relative abundance across sample groups and the color bar indicates phylum affiliation of the different indicators (**see Table S6**).

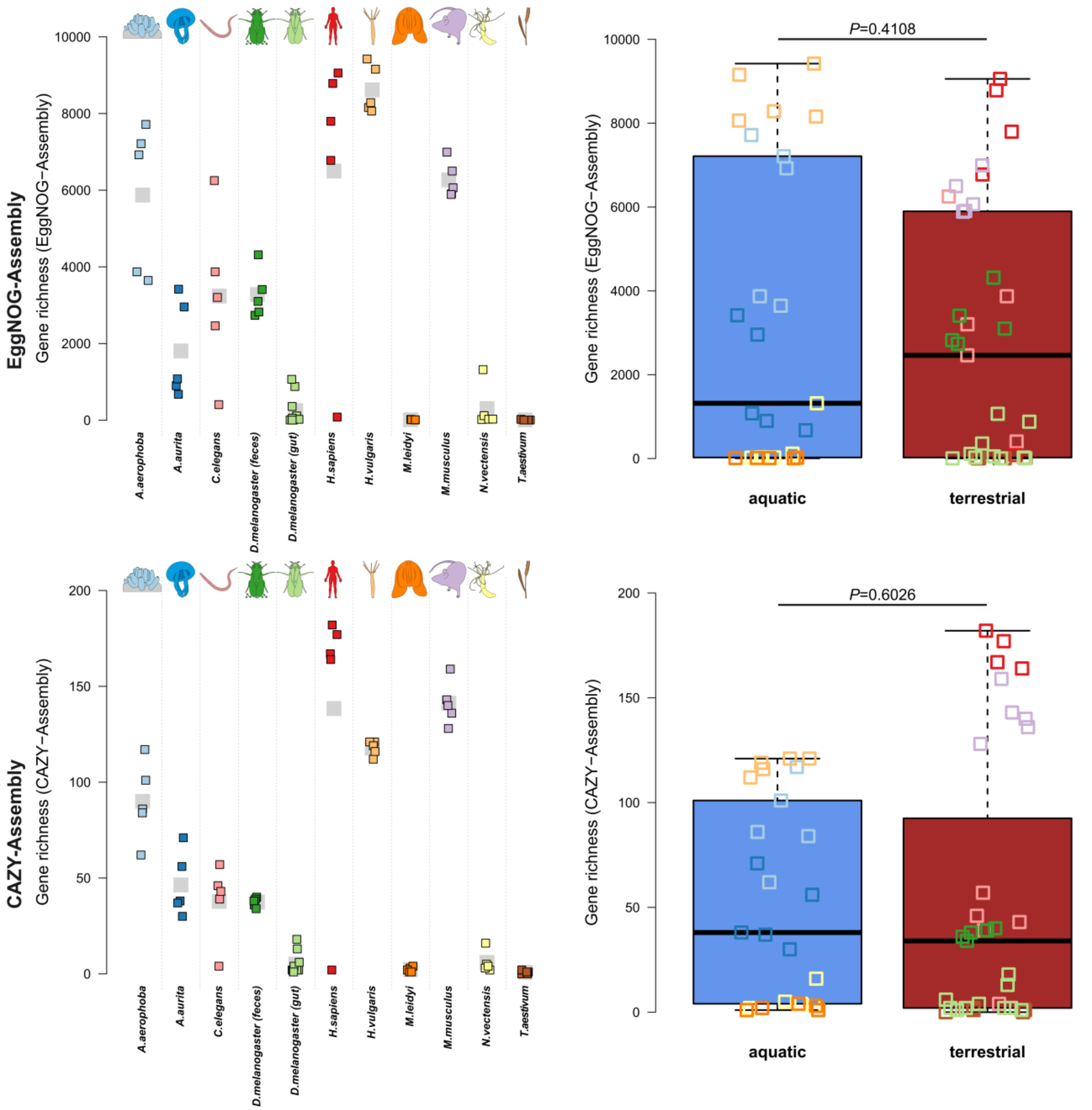


**Figure S21:** Functional diversity of EggNOG derived genes and CAZY predictions in the different organisms and host environments. Pairwise differences of functional diversity between hosts were tested via pairwise *t*-Tests with pooled standard variations. Approximate Wilcoxon tests were employed to compare the host environmental groups. Sample size for the host taxa is

N=5, except for D. melanogaster gut tissue (N=10). The sample size of aquatic hosts is N=25, while terrestrial hosts have a sample size of N=35 (see Table S1).

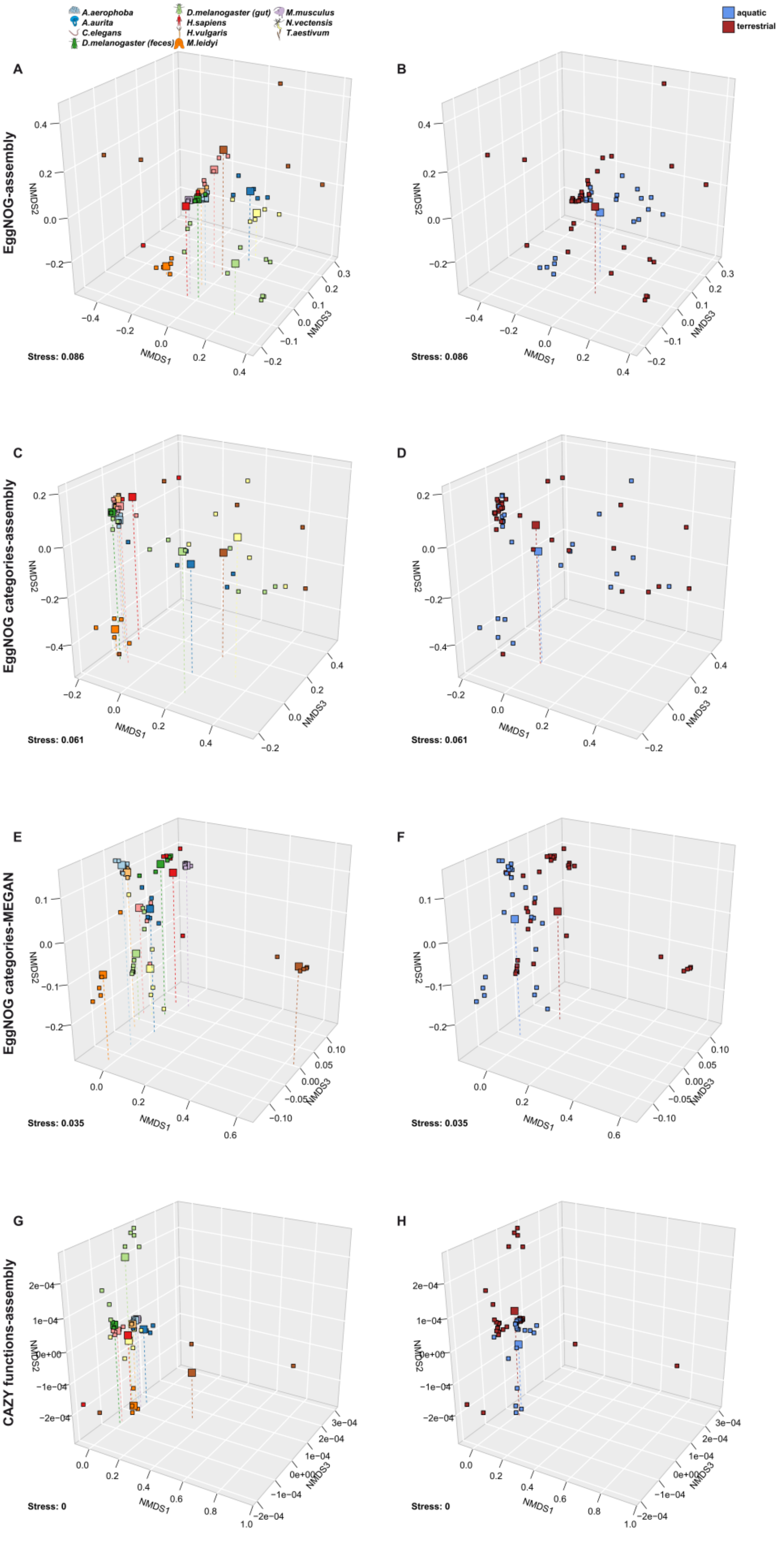
A.aerophoba
A.aurita
C.elegans
D.melanogaster (feces)
D.melanogaster (gut)
H.sapiens
H.vulgaris
M.leidyi
M.musculus
N.vectensis
T.aestivum
aquatic
terrestrial
A
B
EggNOG-assembly
Stress: 0.086
Stress: 0.086
C
D
EggNOG categories-assembly
Stress: 0.061
Stress: 0.061
E
F
EggNOG categories-MEGAN
Stress: 0.035
Stress: 0.035
G
H
CAZY functions-assembly
Stress: 0
Stress: 0
NMDS1
NMDS2
NMDS3

**Figure S22:** Non-metric Multidimensional Scaling of Bray-Curtis distances derived from abundance differences of functional components highlighting functional differences between the host organisms (**A, C, E, G; see Table 3**) and host environments (**B, D, F, H; see Table 3**). Large symbols indicate the centroid of the respective host groups and vertical lines help to determine their position in space. Functional variation of communities based on pairwise Bray-Curtis distances within host organism groups and environmental groups. Sample size for the host taxa is N=5, except for *D. melanogaster* gut tissue (N=10). The sample size of aquatic hosts is N=25, while terrestrial hosts have a sample size of N=35 **(**see Table S1**)**.

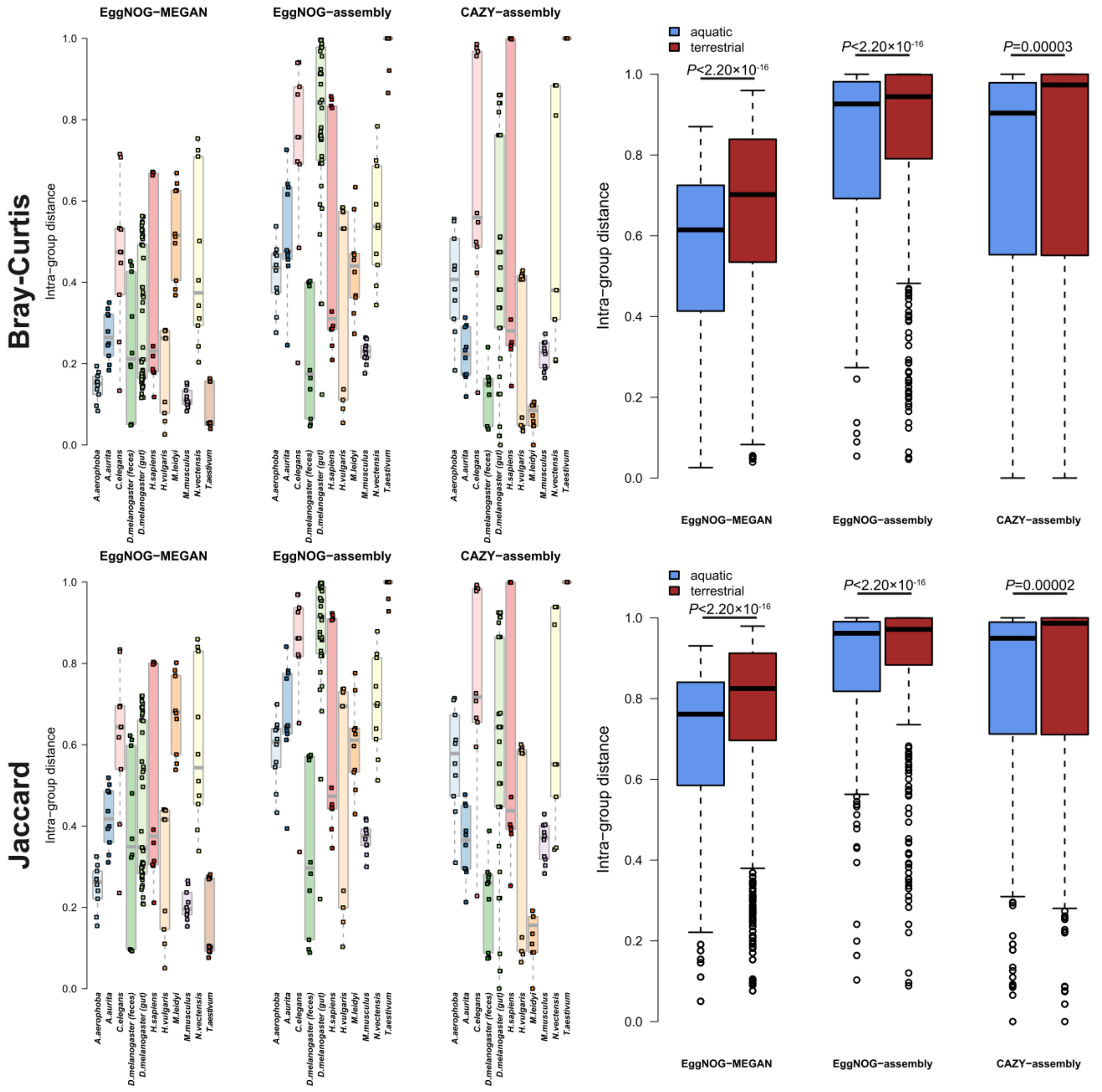

**Figure S23:** Functional variation of communities based on EggNOG derived genes and CAZY predictions based on pairwise Bray-Curtis distances within host organism groups and environmental groups. Terrestrial hosts show a significantly higher Functional variation than aquatic hosts. Sample size for the host taxa is N=5, except for *D. melanogaster* gut tissue (N=10). The sample size of aquatic hosts is N=25, while terrestrial hosts have a sample size of N=35 (see Table S1).

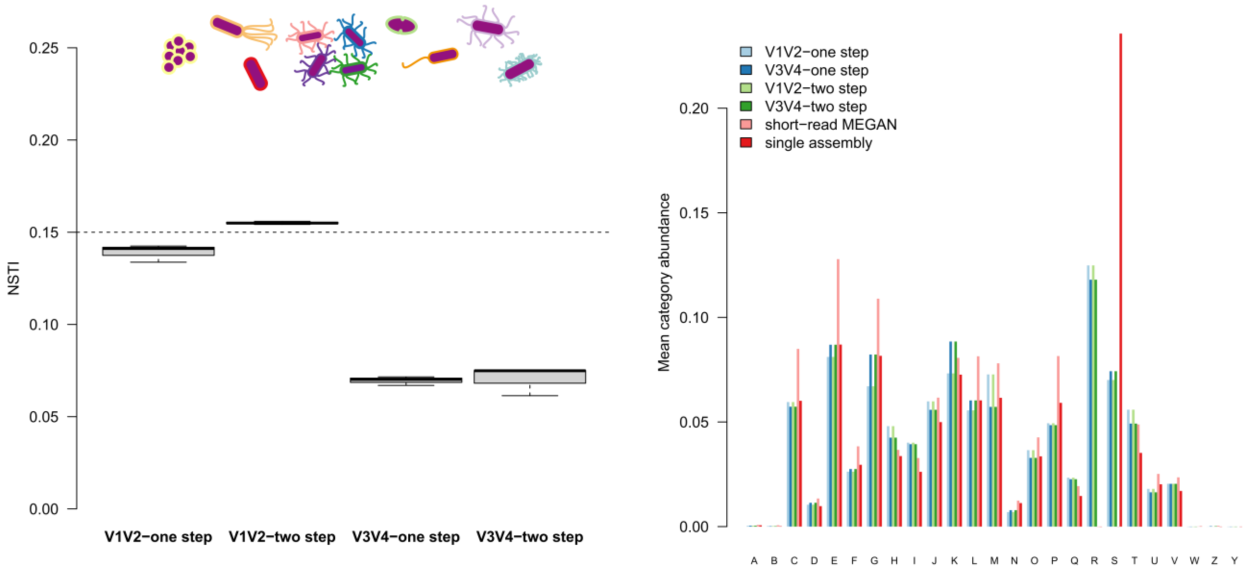


**Figure S24:** NSTI/imputation success differences between variable regions in the mock community samples. Barplot displays average abundances of the single functional categories derived by different 16S rRNA gene amplicon based and shotgun based techniques for mock community samples. Sample sizes for the different approaches are $N_{shotgun}$=4, $N_{V1V2\text{-one step}}$=3, $N_{V1V2\text{-two step}}$=3, $N_{V3V4\text{-one step}}$=3, and $N_{V3V4\text{-two step}}$=3.

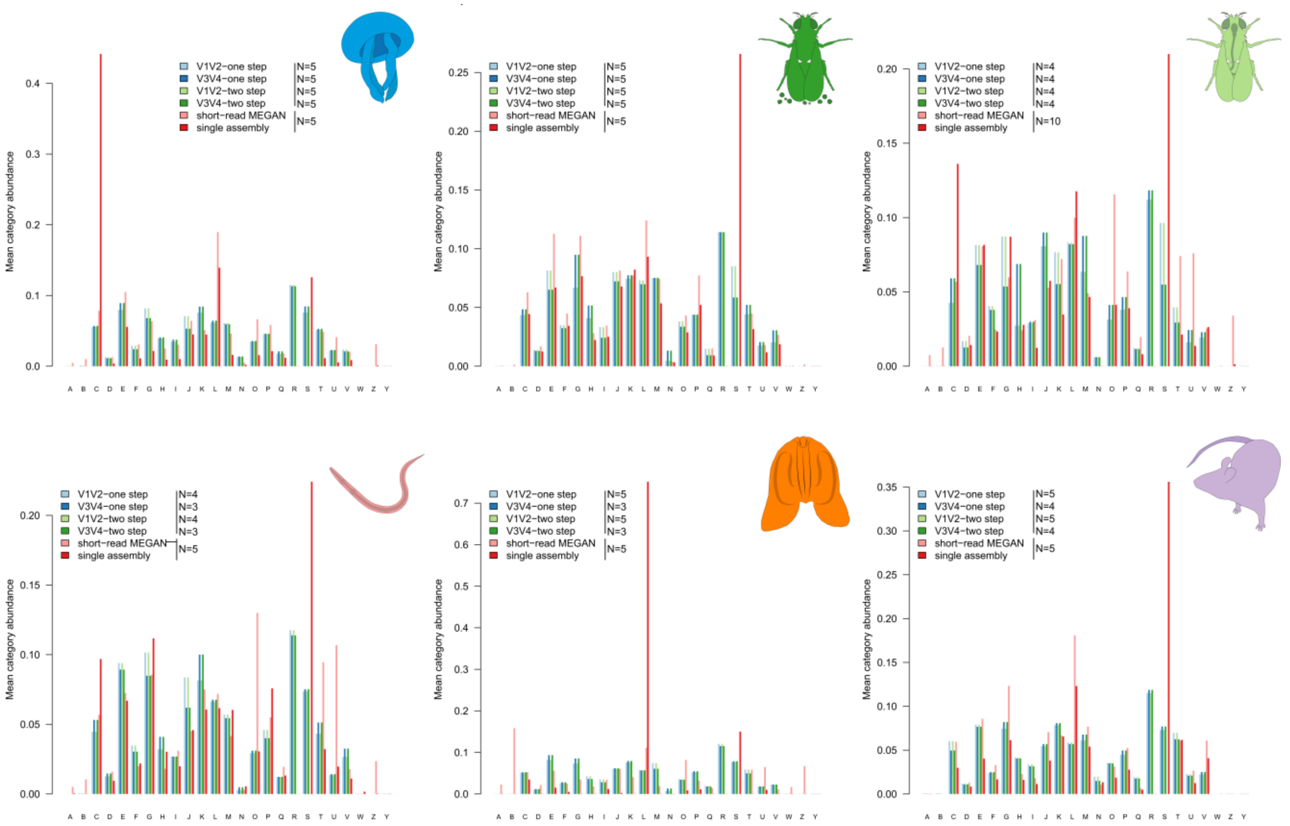


**Information Storage And Processing:**
**J**-Translation, ribosomal structure and biogenesis
**A**-RNA processing and modification
**K**-Transcription
**L**-Replication, recombination and repair
**B**-Chromatin structure and dynamics

**Cellular Processes And Signaling:**
**D**-Cell cycle control, cell division, chromosome partitioning
**Y**-Nuclear structure
**V**-Defense mechanisms
**T**-Signal transduction mechanisms
**M**-Cell wall/membrane/envelope biogenesis
N-Cell motility
**Z**-Cytoskeleton
**W**-Extracellular structures
**U**-Intracellular trafficking, secretion, and vesicular transport
**O**-Posttranslational modification, protein turnover, chaperones

**Metabolism:**
**C**-Energy production and conversion
**G**-Carbohydrate transport and metabolism
**E**-Amino acid transport and metabolism
**F**-Nucleotide transport and metabolism
**H**-Coenzyme transport and metabolism
**I**-Lipid transport and metabolism
**P**-Inorganic ion transport and metabolism
**Q**-Secondary metabolites biosynthesis, transport and catabolism

**Poorly Characterized:**
**R**-General function prediction only
**S**-Function unknown

**Figure S25:** Barplots display average abundances of the single functional categories derived by PICRUSt from the different 16S rRNA amplicon techniques and shotgun based annotations for specific host community samples with sufficient sample coverage.

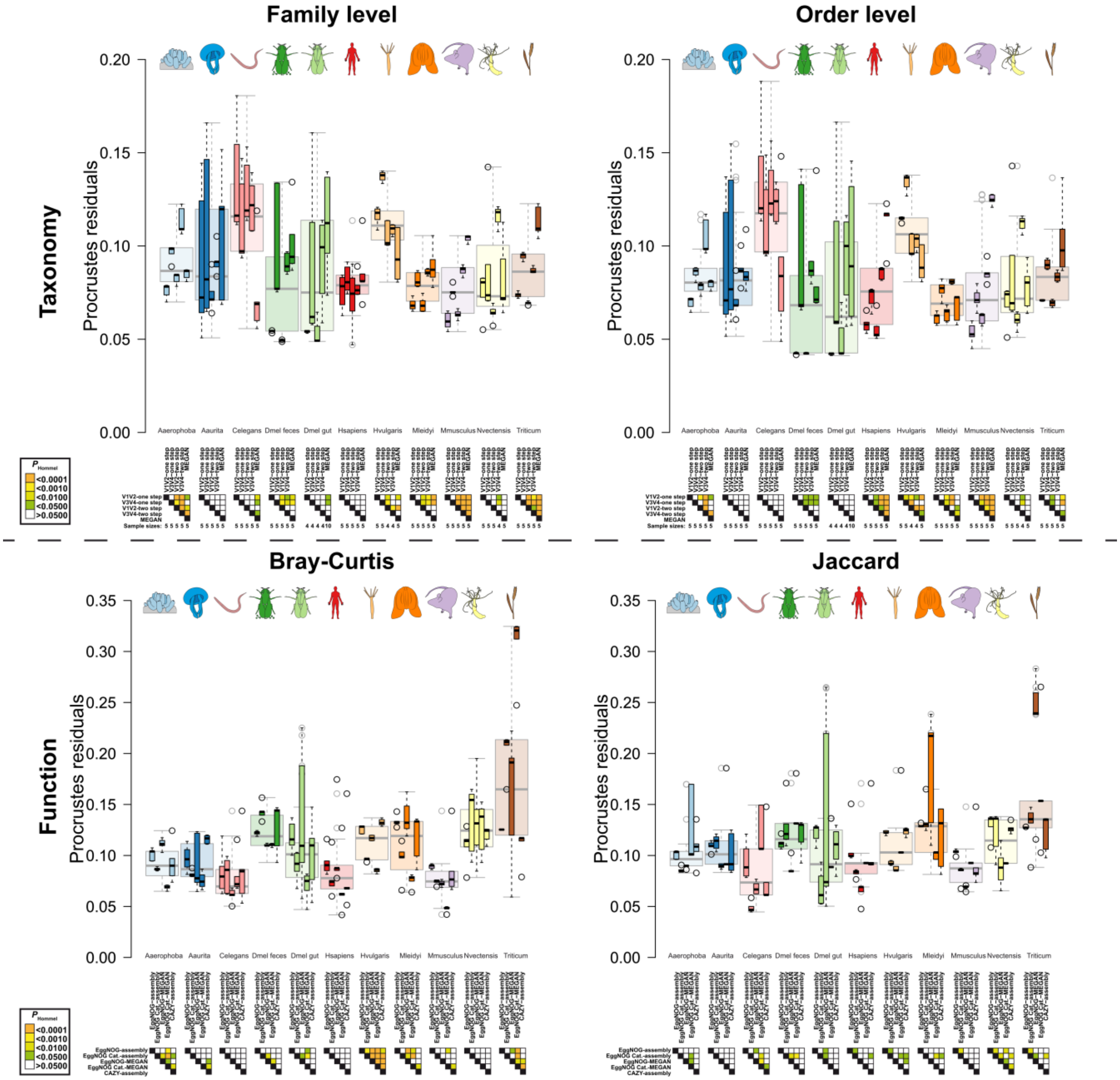


**Figure S26:** Upper panels visualize the residuals derived from Procrustes correlations between phylogenetic and taxonomic microbial community distances among host samples. Sample sizes are indicated below samples. Lower panels show residuals of the correlation between phylogenetic and community distances derived from functional profiles. Sample size for the host taxa is N=5, except for *D. melanogaster* gut tissue (N=10, see Table S1). Pairwise comparisons were computed via pairwise *t*-Tests (Hommel *P*-value adjustment) within host groups. Significance levels after correction for multiple testing are indicated by color.

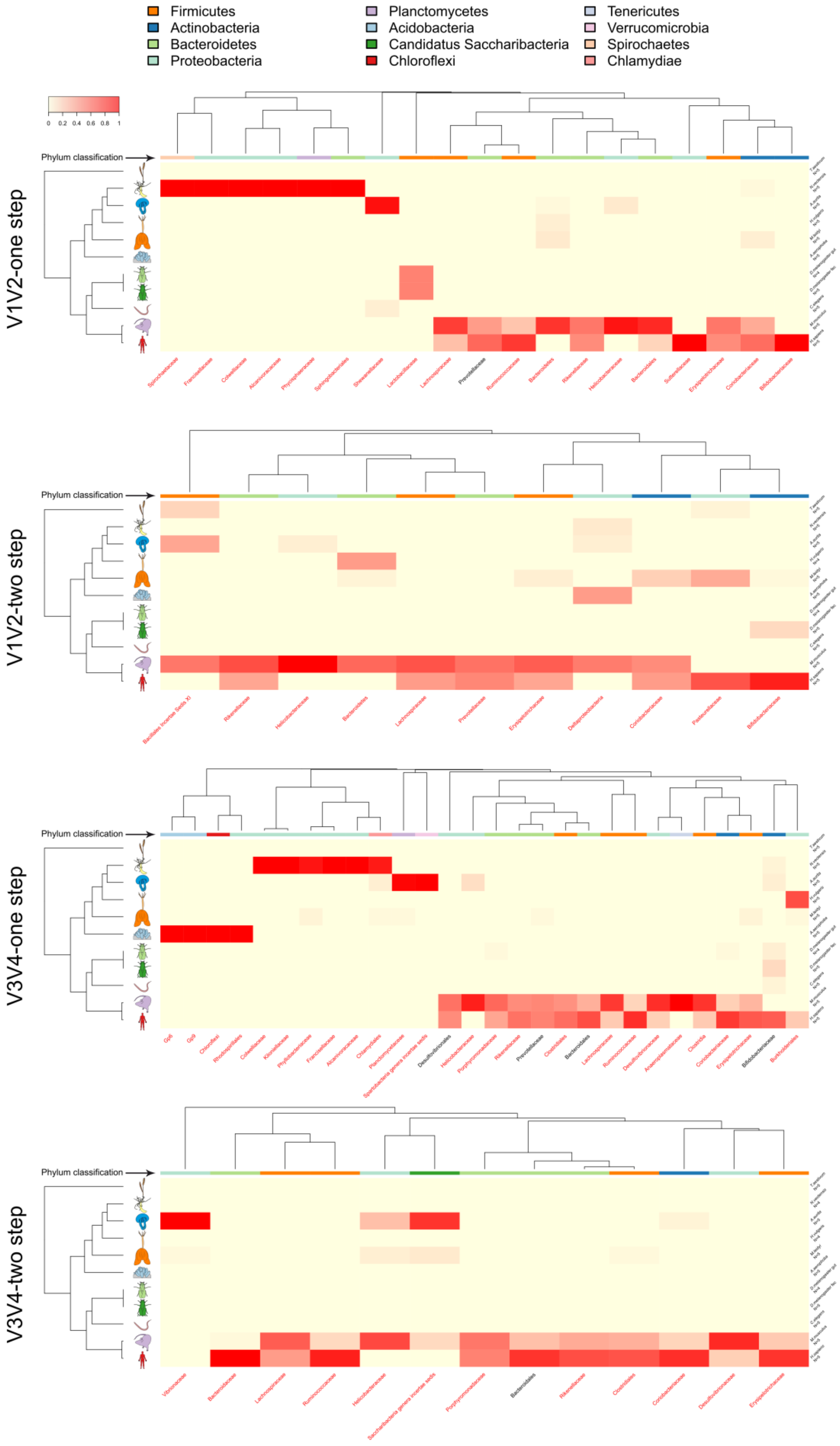

Firmicutes
Actinobacteria
Bacteroidetes
Proteobacteria
Planctomycetes
Acidobacteria
Candidatus Saccharibacteria
Chloroflexi
Tenericutes
Verrucomicrobia
Spirochaetes
Chlamydiae
0 0.2 0.4 0.6 0.8 1
Phylum classification
V1V2-one step
V1V2-two step
V3V4-one step
V3V4-two step

**Figure S27:** Heatmaps of family abundances derived from the different 16S rRNA gene amplicon strategies which associate significantly to phylogenetic distance, as determined by Moran's I test [35]. Heat colors indicate relative abundances across sample groups and the color bar indicates phylum affiliation of the different taxa (see Table S15). Row dendrogram depicts the 18S rRNA gene-based phylogeny of host species. The column dendrogram highlights the similarity of genera distributions as determined by Bray-Curtis distance and average neighbor clustering. Taxon names in red indicate a significant association as an indicator taxon (for sample sizes see Table S1).

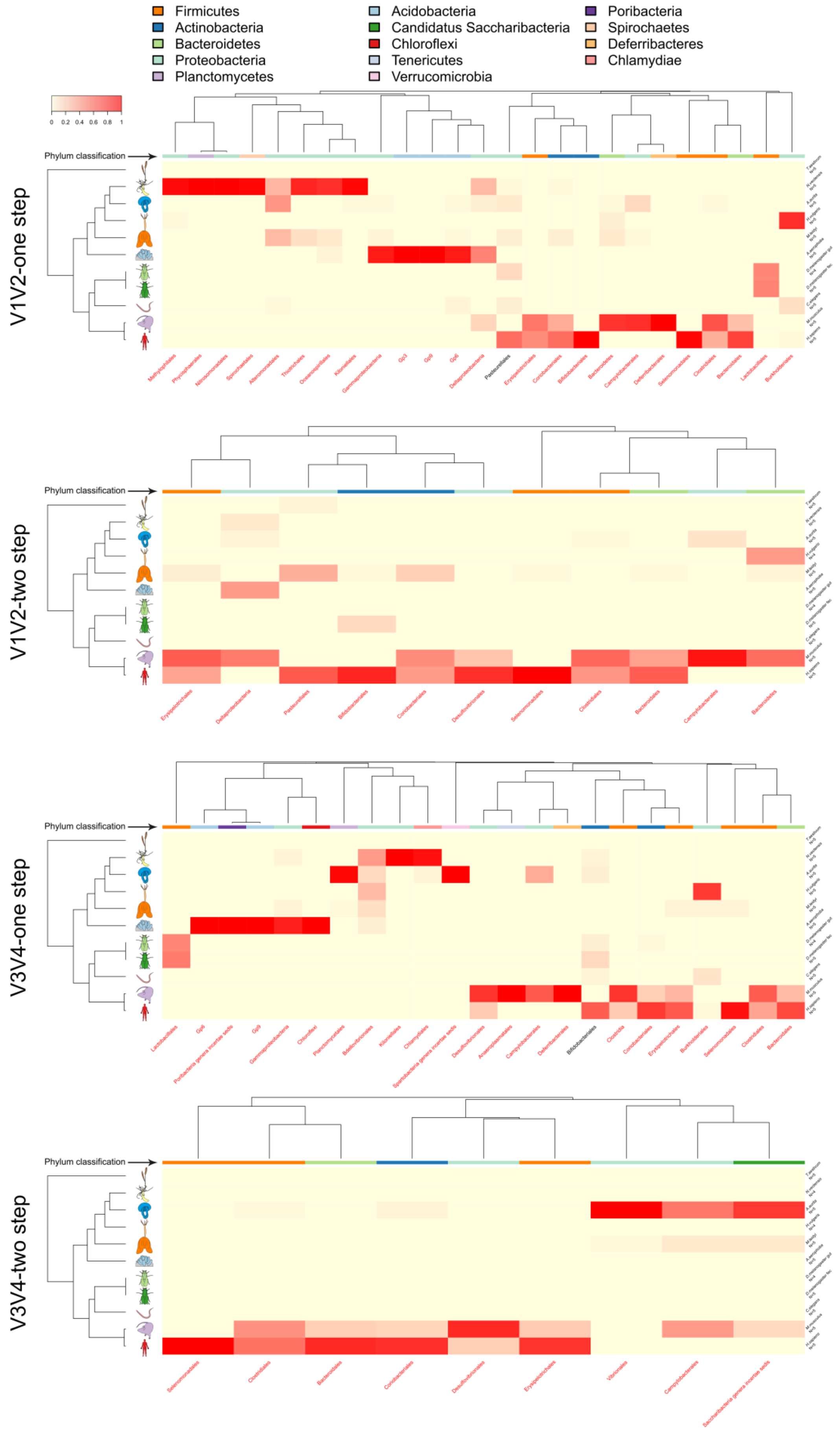
Firmicutes
Actinobacteria
Bacteroidetes
Proteobacteria
Planctomycetes
Acidobacteria
Candidatus Saccharibacteria
Chloroflexi
Tenericutes
Verrucomicrobia
Poribacteria
Spirochaetes
Deferribacteres
Chlamydiae
0 0.2 0.4 0.6 0.8 1
Phylum classification
V1V2-one step
Methylophilales
Phycisphaerales
Nitrosomonadales
Spirochaetales
Alteromonadales
Thiotrichales
Oceanospirillales
Kiloniellales
Gammaproteobacteria
Gp3
Gp9
Gp6
Deltaproteobacteria
Pasteurellales
Erysipelotrichales
Coriobacteriales
Bifidobacteriales
Bacteroidetes
Campylobacterales
Deferribacterales
Selenomonadales
Clostridiales
Bacteroidales
Lactobacillales
Burkholderiales
V1V2-two step
Erysipelotrichales
Deltaproteobacteria
Pasteurellales
Bifidobacteriales
Coriobacteriales
Desulfovibrionales
Selenomonadales
Clostridiales
Bacteroidales
Campylobacterales
Bacteroidales
V3V4-one step
Lactobacillales
Gp6
Poribacteria genera incertae sedis
Gp9
Gammaproteobacteria
Chloroflexi
Planctomycetales
Bdellovibrionales
Kiloniellales
Chlamydiales
Spartobacteria genera incertae sedis
Desulfovibrionales
Anaeroplasmatales
Campylobacterales
Deferribacterales
Bifidobacteriales
Clostridia
Coriobacteriales
Erysipelotrichales
Burkholderiales
Selenomonadales
Clostridiales
Bacteroidales
V3V4-two step
Selenomonadales
Clostridiales
Bacteroidales
Coriobacteriales
Desulfovibrionales
Erysipelotrichales
Vibrionales
Campylobacterales
Saccharibacteria genera incertae sedis

**Figure S28:** Heatmaps of order abundances derived from the different 16S rRNA gene amplicon strategies which associate significantly to phylogenetic distance, as determined by Moran's I test [35]. Heat colors indicate relative abundances across sample groups and the color bar indicates phylum affiliation of the different taxa (see Table S15). Row dendrogram depicts the 18S rRNA gene-based phylogeny of host species. The column dendrogram highlights the similarity of genera distributions as determined by Bray-Curtis distance and average neighbor clustering. Taxon names in red indicate a significant association as an indicator taxon (for sample sizes see Table S1).

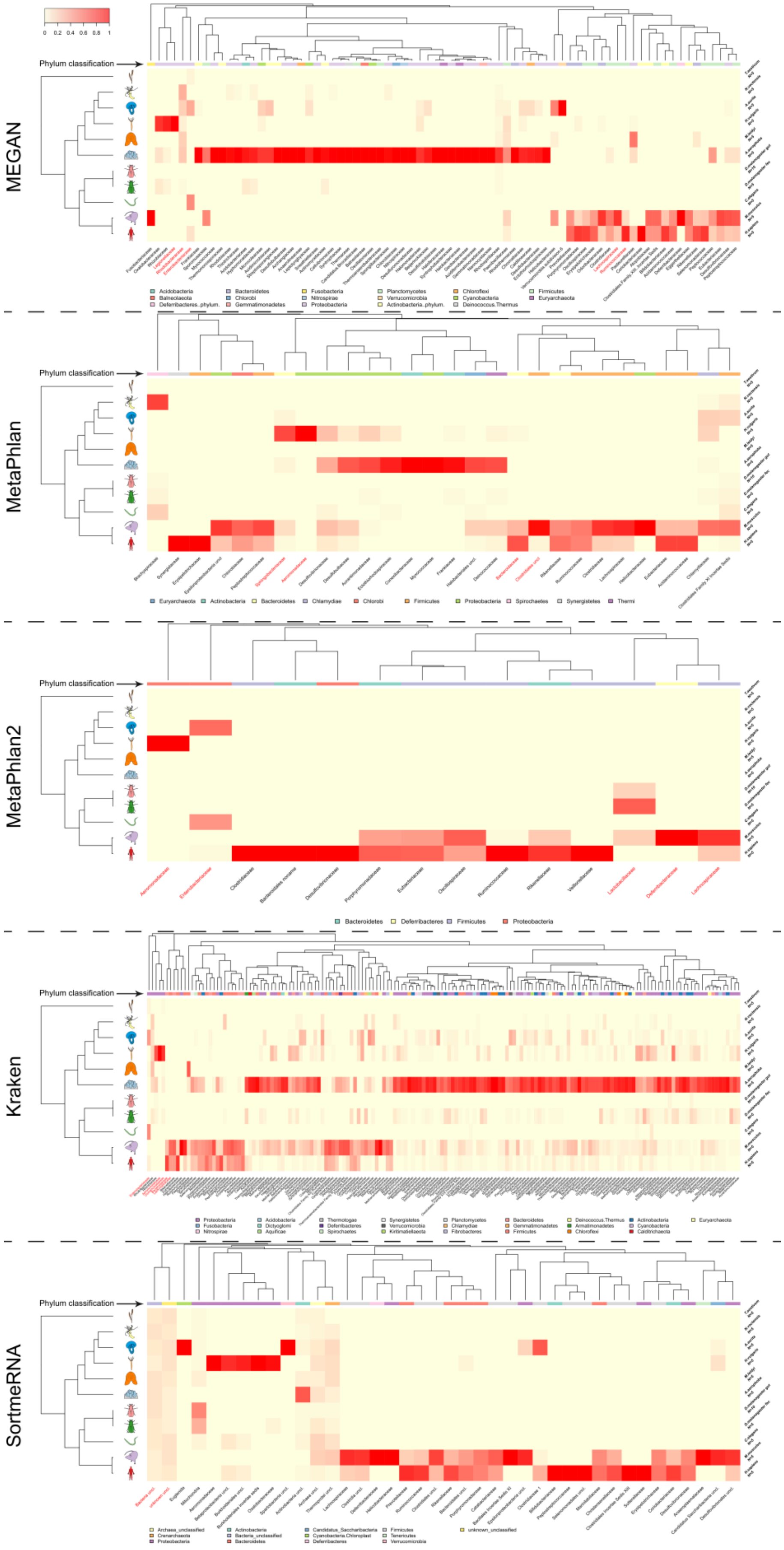
0 0.2 0.4 0.6 0.8 1
Phylum classification
MEGAN
Acidobacteria
Balneolaeota
Deferribacteres..phylum.
Bacteroidetes
Chlorobi
Gemmatimonadetes
Fusobacteria
Nitrospirae
Proteobacteria
Planctomycetes
Verrucomicrobia
Actinobacteria..phylum.
Chloroflexi
Cyanobacteria
Deinococcus.Thermus
Firmicutes
Euryarchaeota
MetaPhlan
Euryarchaeota
Actinobacteria
Bacteroidetes
Chlamydiae
Chlorobi
Firmicutes
Proteobacteria
Spirochaetes
Synergistetes
Thermi
MetaPhlan2
Bacteroidetes
Deferribacteres
Firmicutes
Proteobacteria
Kraken
SortmeRNA

**Figure S29:** Heatmaps of family abundances derived from the different shotgun strategies which associate significantly to phylogenetic distance, as determined by Moran's I test [35]. Heat color indicates relative abundance across sample groups and the color bar indicates phylum affiliation of the different indicators (see Table S16). Row dendrogram shows the 18S rRNA gene-based tree of host species (ML tree) and the column dendrogram highlights the similarity of genera distributions as determined by Bray-Curtis distance and average neighbor clustering. Taxon names in red indicate a significant association as an indicator taxon. Sample size for the host taxa is N=5, except for *D. melanogaster* gut tissue (N=10, see Table S1).

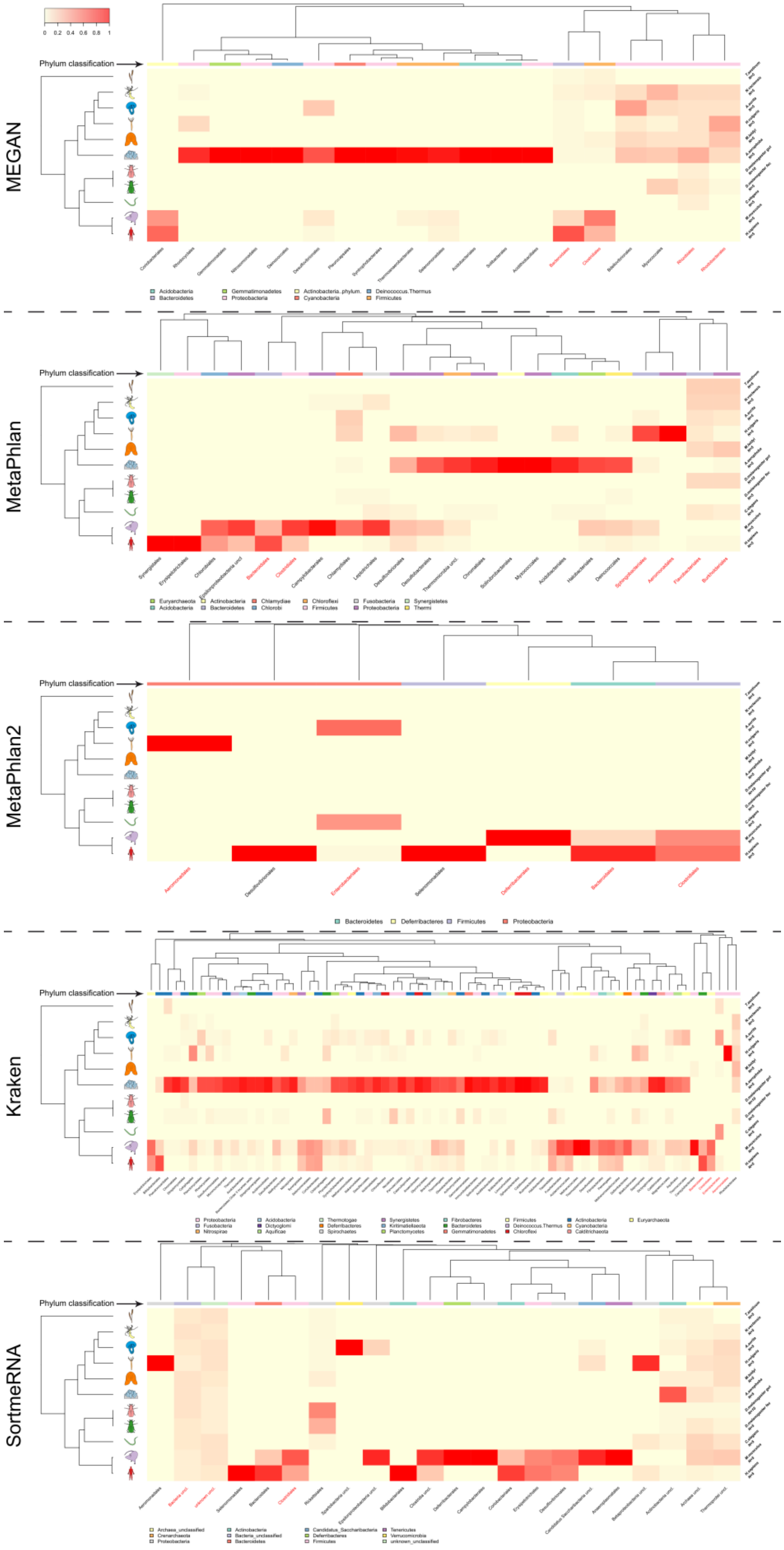

0 0.2 0.4 0.6 0.8 1
Phylum classification
MEGAN
Acidobacteria
Bacteroidetes
Gemmatimonadetes
Proteobacteria
Actinobacteria_phylum
Cyanobacteria
Deinococcus.Thermus
Firmicutes
MetaPhlan
Euryarchaeota
Acidobacteria
Actinobacteria
Bacteroidetes
Chlamydiae
Chlorobi
Chloroflexi
Firmicutes
Fusobacteria
Proteobacteria
Synergistetes
Thermi
MetaPhlan2
Bacteroidetes
Deferribacteres
Firmicutes
Proteobacteria
Kraken
SortmeRNA

**Figure S30:** Heatmaps of order abundances derived from the different shotgun strategies which associate significantly to phylogenetic distance, as determined by Moran's I test [35]. Heat color indicates relative abundance across sample groups and the color bar indicates phylum affiliation of the different indicators (see Table S16). Row dendrogram shows the 18S rRNA gene-based tree of host species (ML tree) and the column dendrogram highlights the similarity of genera distributions as determined by Bray-Curtis distance and average neighbor clustering. Taxon names in red indicate a significant association as an indicator taxon. Sample size for the host taxa is N=5, except for *D. melanogaster* gut tissue (N=10, see Table S1).

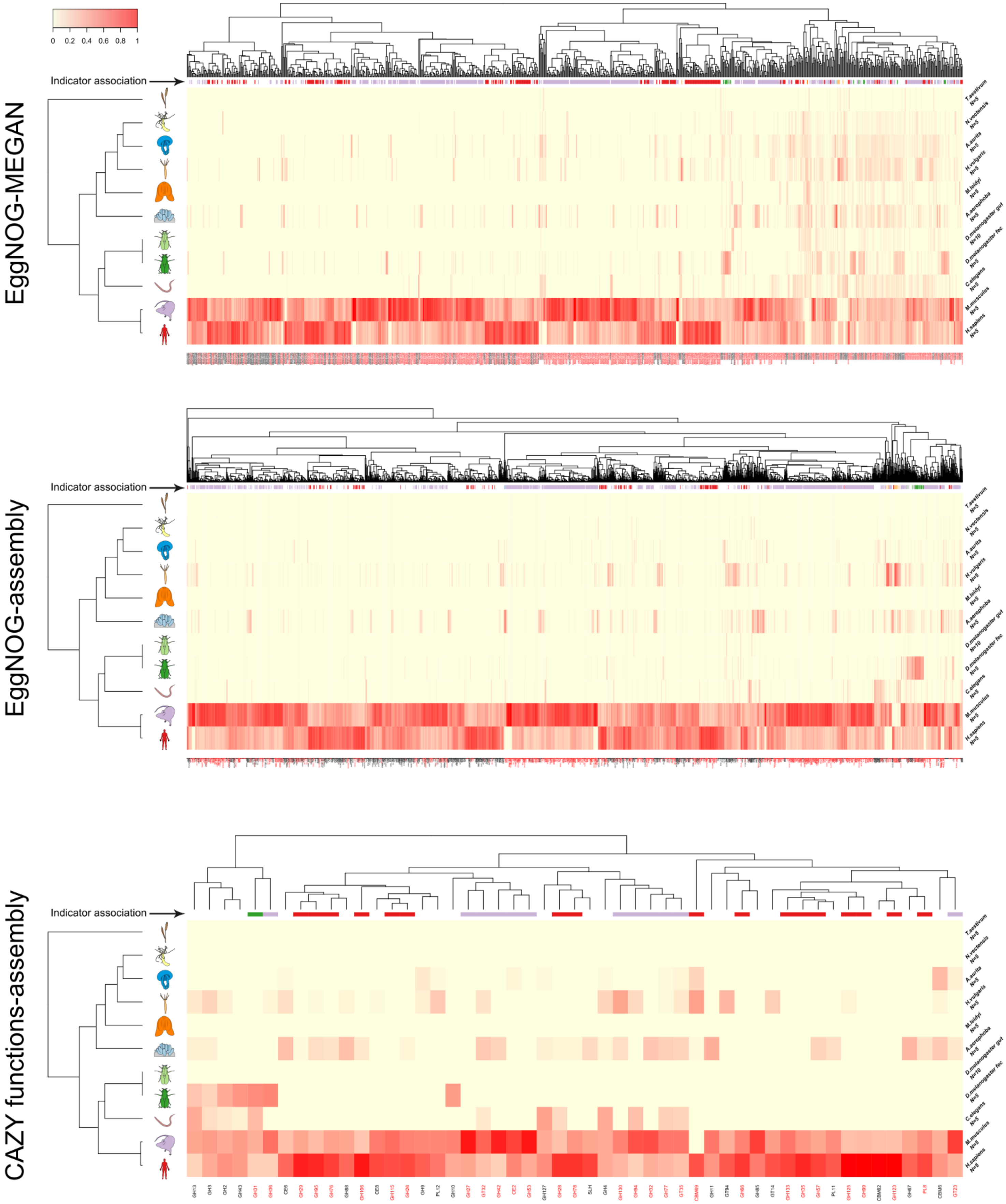
0
0.2
0.4
0.6
0.8
1
EggNOG-MEGAN
EggNOG-assembly
CAZY functions-assembly
Indicator association
T.aestivum N=5
N.vectensis N=5
A.aurita N=5
H.vulgaris N=5
M.leidyi N=5
A.aerophoba N=5
D.melanogaster gut N=10
D.melanogaster fec N=5
C.elegans N=5
M.musculus N=5
H.sapiens N=5
GH13 GH3 GH2 GH43 GH31 GH36 CE6 GH29 GH95 GH76 GH88 GH106 CE8 GH115 GH26 GH9 PL12 GH10 GH27 GT32 GH42 CE2 GH53 GH127 GH28 GH78 SLH GH4 GH130 GH94 GH32 GH77 GT35 CBM69 GH11 GT94 GH66 GH85 GT14 GH133 GH35 GH57 PL11 GH125 GH99 CBM62 GH123 GH67 PL8 CBM6 GT23

**Figure S31:** Heatmaps of function abundances derived from the different metagenomic functional annotation strategies associate significantly to phylogenetic distance, as determined by Moran's I test [35]. Heat colors indicate relative abundance across sample groups and the color bar indicates single host indicator associations (see Table S17-S20). Function names in red indicate a significant association as an indicator function. Row dendrogram shows the 18S rRNA gene-based tree of host species (ML tree) and the column dendrogram highlights the similarity of genera distributions as determined by Bray-Curtis distance and average neighbor clustering. Sample size for the host taxa is N=5, except for *D. melanogaster* gut tissue (N=10, see Table S1).

**Supplementary Tables:**

Table S1: Sample sizes of the 16S rRNA amplicon based- and shotgun based approaches for single host taxa and environments.

Table S2: Differences between expected and observed alpha diversity of the mock community, based on different shotgun- and 16S rRNA gene amplicon based community compositions (bacterial genus level), tested via a one-sample *t*-test (two-sided, *P*-values adjusted via Hommel procedure).

Table S3: Differences in community distance to the expected composition of the mock community between 16S rRNA gene amplicon and shotgun based techniques based on on shared abundance (Bray-Curtis) and shared presence (Jaccard) of bacterial genera. Tested via a one-sample *t*-test (two-sided, *P*-values adjusted via Hommel procedure).

Table S4: Pairwise mock community compositional differences between the 16S rRNA gene amplicon and shotgun based techniques focusing on shared abundance (Bray-Curtis) and shared presence (Jaccard) of bacterial genera (PERMANOVA *P*-values based on 10'000 permutations, *P*-values adjusted via Hommel procedure).

Table S5: Differences in alpha diversity estimates based on genus abundances between different hosts based on 16S rRNA amplicon and shotgun based techniques (pairwise *t*-test with Hommel adjusted *P*-values).

Table S6: Pairwise community compositional differences between the host species based on shared abundance of genera in the different amplicon techniques (Bray-Curtis). *P*-values are derived by PERMANOVA based on 10'000 permutations (*P*-values adjusted via Hommel procedure for each amplicon technique).

Table S7: Indicator genera for host taxa using different 16S rRNA gene amplicon techniques. Analyses are based on 10'000 permutations and *P*-value correction for each amplicon technique separately. Host taxa in grey highlight disagreement in their association and overlaps with shotgun techniques are highlighted.

Table S8: Indicator genera for host taxa using the different shotgun techniques. Analyses are based on 10’000 permutations and *P*-value correction for each technique separately. Host taxa in grey highlight disagreement in their association overlaps with the results from the 16S rRNA gene amplicon analyses are highlighted.

Table S9: Indicator genera for host environments using different 16S rRNA gene amplicon techniques. Analyses are based on 10’000 permutations and *P*-value correction for each 16S rRNA gene amplicon technique separately. Overlapping associations with shotgun techniques are highlighted.

Table S10: Indicator genera for host environments using the different shotgun techniques. Analyses are based on 10’000 permutations and *P*-value correction for each technique separately. Overlapping associations with amplicon techniques are highlighted.

Table S11: Indicator functions for host taxa using the EggNOG classification based on MEGAN. Analyses are based on 10’000 permutations with Benjamini-Yeketuli P-value correction.

Table S12: Indicator functions for host taxa using the EggNOG classification based on single sample assemblies and *emapper* annotation. Analyses are based on 10’000 permutations with Benjamini-Yeketuli *P*-value correction.

Table S13: Indicator functional categories for host taxa using the EggNOG classification based on single sample assemblies and emapper annotation and MEGAN. Analyses are based on 10’000 permutations with Benjamini-Yeketuli *P*-value correction.

Table S14: Indicator functions for host taxa using the CAZY classification based on single sample assemblies. Analyses are based on 10’000 permutations with Benjamini-Yeketuli *P*-value correction.

Table S15: Pairwise PERMANOVA comparisons between functional repertoires (EggNOG, COG) derived from single assemblies, MEGAN, and PICRUSt imputations (V1V2, V3V4, one step, two step).

Table S16: Moran’s eigenvector analyses for family and order abundances in combination with indicator analysis results for single and multiple hosts (maximum 3) using the 16S rRNA gene amplicon and metagenomic shotgun (MEGAN) based techniques (10’000 permutations). Repeated associations for single bacterial taxa, including their indicator associations are highlighted.

Table S17: Moran’s eigenvector analyses for family and order abundances in combination with indicator genera for single and multiple hosts (maximum 3) using the 16S and all shotgun based techniques (MEGAN, Kraken, MetaPhlan, MetaPhlan2, SortmeRNA;10’000 permutations). Repeated associations for single bacterial taxa, including their indicator associations are highlighted.

Table S18: Moran’s eigenvector analyses for CAZY functional abundances.

Table S19: Moran’s eigenvector analyses for EggNOG functional abundances derived from single sample assemblies.

Table S20: Moran's eigenvector analyses for EggNOG functional abundances derived from MEGAN.

Table S21: Moran's eigenvector analyses for EggNOG functional category abundances derived from MEGAN and single sample assemblies.
