## Supplemental Table S1 for "Comparative analysis of amplicon and metagenomic sequencing methods reveals key features in the evolution of animal metaorganisms"

|  |  |  | V1V2 |  | V3V4 |  |
| --- | --- | --- | --- | --- | --- | --- |
|  | Samples | shotgun | one step | two step | one step | two step |
| Sequencing | *A. aerophoba* | 5 | 5 | 5 | 5 | 5 |
|  | *A. aurita* | 5 | 5 | 5 | 5 | 5 |
|  | *C. elegans* | 5 | 5 | 5 | 5 | 5 |
|  | *D. melanogaster (feces)* | 5 | 5 | 5 | 5 | 5 |
|  | *D. melanogaster (gut)* | 10 | 4 | 4 | 4 | 4 |
|  | *H. sapiens* | 5 | 5 | 5 | 5 | 5 |
|  | *H. vulgaris* | 5 | 5 | 4 | 5 | 4 |
|  | *M. leidyi* | 5 | 5 | 5 | 5 | 5 |
|  | *M. musculus* | 5 | 5 | 5 | 5 | 5 |
|  | *N. vectensis* | 5 | 5 | 5 | 5 | 4 |
|  | *T. aestivum* | 5 | 5 | 5 | 5 | 5 |
|  | Mock | 4 | 3 | 3 | 3 | 3 |
|  | aquatic | 25 | 25 | 24 | 25 | 23 |
|  | terrestrial | 35 | 29 | 29 | 29 | 29 |
| PICRUSt | *A. aerophoba* | - | 1 | 1 | 1 | 1 |
|  | *A. aurita** | - | 5 | 5 | 5 | 5 |
|  | *C. elegans** | - | 4 | 4 | 3 | 3 |
|  | *D. melanogaster (feces)** | - | 5 | 5 | 5 | 5 |
|  | *D. melanogaster (gut)** | - | 4 | 4 | 4 | 4 |
|  | *H. sapiens* | - | 1 | 1 | 4 | 4 |
|  | *H. vulgaris* | - | 0 | 0 | 1 | 1 |
|  | *M. leidyi** | - | 5 | 5 | 3 | 3 |
|  | *M. musculus** | - | 5 | 5 | 4 | 4 |
|  | *N. vectensis* | - | 1 | 1 | 2 | 2 |
|  | *T. aestivum** | - | 2 | 2 | 4 | 4 |
|  | Mock*** | - | 3 | 3 | 3 | 3 |

* hosts with a sufficient number samples derived from PICRUSt
