## Supplemental Table S2 for "Comparative analysis of amplicon and metagenomic sequencing methods reveals key features in the evolution of animal metaorganisms"

| Sequencing |  | Shannon |  | Richness |  |
| --- | --- | --- | --- | --- | --- |
| technique | Data | *P* | *P*_Hommel_ | *P* | *P*_Hommel_ |
| shotgun | MEGAN | 0.00005 | 0.00032 | 0.00009 | 0.00062 |
|  | MetaPhlan | 0.00002 | 0.00013 | 0.01826 | 0.01826 |
|  | MetaPhlan2 | 0.19723 | 0.39446 | no difference | no difference |
|  | Kraken | 0.12836 | 0.36079 | 0.01069 | 0.01826 |
|  | SortmeRNA | 0.00001 | 0.00007 | 0.00036 | 0.00215 |
| amplicon | V1V2-one step | 0.71781 | 0.71781 | 0.00593 | 0.01791 |
|  | V3V4-one step | 0.00476 | 0.02856 | 0.01343 | 0.01826 |
|  | V1V2-two step | 0.06316 | 0.30065 | 0.00001 | 0.00006 |
|  | V3V4-two step | 0.24052 | 0.48105 | 0.00765 | 0.01826 |
