## Supplemental Table S3 for "Comparative analysis of amplicon and metagenomic sequencing methods reveals key features in the evolution of animal metaorganisms"

|  |  | Bray-Curtis |  | Jaccard |  |
| --- | --- | --- | --- | --- | --- |
| Group 1 | Group 2 | *P* | *P*_Hommel_ | *P* | *P*_Hommel_ |
| MEGAN | Kraken | 0.0083921 | 0.0503524 | 0.6885784 | 0.9078331 |
| MetaPhlan | Kraken | 3.73 × 10^-09^ | 0.0000001 | 0.0000013 | 0.0000301 |
| MetaPhlan2 | Kraken | 2.68 × 10^-10^ | 6.43 × 10^-09^ | 0.0000338 | 0.0007095 |
| SortmeRNA | Kraken | 3.16 × 10^-22^ | 9.81 × 10^-21^ | 0.0000007 | 0.0000174 |
| V1V2-one step | Kraken | 0.0001362 | 0.0016341 | 0.4354358 | 0.9078331 |
| V1V2-two step | Kraken | 0.1403239 | 0.1796733 | 0.3676752 | 0.9078331 |
| V3V4-one step | Kraken | 1.67 × 10^-11^ | 4.16 × 10^-10^ | 0.0092672 | 0.1390078 |
| V3V4-two step | Kraken | 0.0000005 | 0.0000073 | 0.0047803 | 0.0741375 |
| MetaPhlan | MEGAN | 1.98 × 10^-11^ | 4.96 × 10^-10^ | 0.0000033 | 0.0000752 |
| MetaPhlan2 | MEGAN | 2.20 × 10^-12^ | 5.94 × 10^-11^ | 0.0000125 | 0.0002742 |
| SortmeRNA | MEGAN | 1.90 × 10^-21^ | 5.51 × 10^-20^ | 0.0000018 | 0.0000413 |
| V1V2-one step | MEGAN | 0.0000002 | 0.0000039 | 0.2541565 | 0.9078331 |
| V1V2-two step | MEGAN | 0.0003449 | 0.0036676 | 0.2082728 | 0.9078331 |
| V3V4-one step | MEGAN | 2.86 × 10^-13^ | 8.02 × 10^-12^ | 0.003828 | 0.0612483 |
| V3V4-two step | MEGAN | 1.63 × 10^-09^ | 3.58 × 10^-08^ | 0.0019341 | 0.0328802 |
| MetaPhlan2 | MetaPhlan | 0.1796733 | 0.1796733 | 4.14 × 10^-11^ | 1.45 × 10^-09^ |
| SortmeRNA | MetaPhlan | 2.28 × 10^-24^ | 7.76 × 10^-23^ | 0.8026244 | 0.9078331 |
| V1V2-one step | MetaPhlan | 0.0006668 | 0.0063575 | 0.0000006 | 0.0000156 |
| V1V2-two step | MetaPhlan | 0.0000004 | 0.0000067 | 0.0000005 | 0.0000119 |
| V3V4-one step | MetaPhlan | 0.0012715 | 0.0114435 | 7.20 × 10^-09^ | 0.0000002 |
| V3V4-two step | MetaPhlan | 0.1309865 | 0.1796733 | 4.08 × 10^-09^ | 0.0000001 |
| SortmeRNA | MetaPhlan2 | 1.18 × 10^-24^ | 4.01 × 10^-23^ | 2.70 × 10^-11^ | 9.46 × 10^-10^ |
| V1V2-one step | MetaPhlan2 | 0.0000276 | 0.0003859 | 0.0006276 | 0.0119239 |
| V1V2-two step | MetaPhlan2 | 2.49 × 10^-08^ | 0.0000005 | 0.0008556 | 0.0162559 |
| V3V4-one step | MetaPhlan2 | 0.0254947 | 0.1019789 | 0.0687707 | 0.6189367 |
| V3V4-two step | MetaPhlan2 | 0.0091196 | 0.0547177 | 0.1174094 | 0.8708715 |
| V1V2-one step | SortmeRNA | 1.13 × 10^-22^ | 3.61 × 10^-21^ | 0.0000004 | 0.0000094 |
| V1V2-two step | SortmeRNA | 6.90 × 10^-22^ | 2.07 × 10^-20^ | 0.0000003 | 0.0000072 |
| V3V4-one step | SortmeRNA | 2.09 × 10^-24^ | 7.11 × 10^-23^ | 4.49 × 10^-09^ | 0.0000001 |
| V3V4-two step | SortmeRNA | 3.03 × 10^-23^ | 1.00 × 10^-21^ | 2.57 × 10^-09^ | 0.0000001 |
| V1V2-two step | V1V2-one step | 0.0091569 | 0.0549415 | 0.9078331 | 0.9078331 |
| V3V4-one step | V1V2-one step | 0.0000003 | 0.0000049 | 0.0680896 | 0.6128062 |
| V3V4-two step | V1V2-one step | 0.0371268 | 0.1485071 | 0.0398532 | 0.430501 |
| V3V4-one step | V1V2-two step | 8.31 × 10^-10^ | 1.87 × 10^-08^ | 0.0854235 | 0.7624695 |
| V3V4-two step | V1V2-two step | 0.0000404 | 0.000525 | 0.0507164 | 0.5071642 |
| V3V4-two step | V3V4-one step | 0.0000601 | 0.0007811 | 0.7938902 | 0.9078331 |

Shading highlights significant comparisons
