## Supplemental Table S4 for "Comparative analysis of amplicon and metagenomic sequencing methods reveals key features in the evolution of animal metaorganisms"

| Distance | Data 1 | Data 2 | *DF* | *F* | *P* | *P*_FDR_ | *R*^2^ | adj. *R*^2^ |
| --- | --- | --- | --- | --- | --- | --- | --- | --- |
| Bray-Curtis | Kraken | MEGAN | 1,6 | 19.9121 | 0.0275 | 0.0369 | 0.7684 | 0.7299 |
|  |  | MetaPhlan | 1,6 | 8.7955 | 0.0297 | 0.0369 | 0.5945 | 0.5269 |
|  |  | MetaPhlan2 | 1,6 | 7.8408 | 0.0288 | 0.0369 | 0.5665 | 0.4943 |
|  |  | SortmeRNA | 1,6 | 43.4914 | 0.0285 | 0.0369 | 0.8788 | 0.8586 |
|  |  | V1V2-one step | 1,5 | 9.0041 | 0.0286 | 0.0369 | 0.6430 | 0.5716 |
|  |  | V1V2-two step | 1,5 | 10.8181 | 0.0286 | 0.0369 | 0.6839 | 0.6207 |
|  |  | V3V4-one step | 1,5 | 10.5970 | 0.0286 | 0.0369 | 0.6794 | 0.6153 |
|  |  | V3V4-two step | 1,5 | 9.7830 | 0.0286 | 0.0369 | 0.6618 | 0.5941 |
|  | MEGAN | MetaPhlan | 1,6 | 37.6048 | 0.0260 | 0.0369 | 0.8624 | 0.8395 |
|  |  | MetaPhlan2 | 1,6 | 31.6016 | 0.0294 | 0.0369 | 0.8404 | 0.8138 |
|  |  | SortmeRNA | 1,6 | 165.8439 | 0.0287 | 0.0369 | 0.9651 | 0.9593 |
|  |  | V1V2-one step | 1,5 | 17.6692 | 0.0286 | 0.0369 | 0.7794 | 0.7353 |
|  |  | V1V2-two step | 1,5 | 26.1347 | 0.0286 | 0.0369 | 0.8394 | 0.8073 |
|  |  | V3V4-one step | 1,5 | 34.8718 | 0.0286 | 0.0369 | 0.8746 | 0.8495 |
|  |  | V3V4-two step | 1,5 | 20.1193 | 0.0286 | 0.0369 | 0.8009 | 0.7611 |
|  | MetaPhlan | MetaPhlan2 | 1,6 | 5.5451 | 0.0278 | 0.0369 | 0.4803 | 0.3937 |
|  |  | SortmeRNA | 1,6 | 134.1606 | 0.0311 | 0.0373 | 0.9572 | 0.9501 |
|  |  | V1V2-one step | 1,5 | 11.0571 | 0.0286 | 0.0369 | 0.6886 | 0.6263 |
|  |  | V1V2-two step | 1,5 | 15.1944 | 0.0286 | 0.0369 | 0.7524 | 0.7029 |
|  |  | V3V4-one step | 1,5 | 18.6144 | 0.0286 | 0.0369 | 0.7883 | 0.7459 |
|  |  | V3V4-two step | 1,5 | 12.8494 | 0.0286 | 0.0369 | 0.7199 | 0.6639 |
|  | MetaPhlan2 | SortmeRNA | 1,6 | 123.5607 | 0.0287 | 0.0369 | 0.9537 | 0.9460 |
|  |  | V1V2-one step | 1,5 | 10.6063 | 0.0286 | 0.0369 | 0.6796 | 0.6155 |
|  |  | V1V2-two step | 1,5 | 14.2274 | 0.0286 | 0.0369 | 0.7400 | 0.6879 |
|  |  | V3V4-one step | 1,5 | 17.3604 | 0.0286 | 0.0369 | 0.7764 | 0.7317 |
|  |  | V3V4-two step | 1,5 | 12.1932 | 0.0286 | 0.0369 | 0.7092 | 0.6510 |
|  | SortmeRNA | V1V2-one step | 1,5 | 53.9770 | 0.0286 | 0.0369 | 0.9152 | 0.8983 |
|  |  | V1V2-two step | 1,5 | 65.1984 | 0.0286 | 0.0369 | 0.9288 | 0.9145 |
|  |  | V3V4-one step | 1,5 | 114.4553 | 0.0286 | 0.0369 | 0.9581 | 0.9498 |
|  |  | V3V4-two step | 1,5 | 56.4162 | 0.0286 | 0.0369 | 0.9186 | 0.9023 |
|  | V1V2-one step | V1V2-two step | 1,4 | 2.9558 | 0.1000 | 0.1000 | 0.4249 | 0.2812 |
|  |  | V3V4-one step | 1,4 | 5.0552 | 0.1000 | 0.1000 | 0.5583 | 0.4478 |
|  |  | V3V4-two step | 1,4 | 3.6834 | 0.1000 | 0.1000 | 0.4794 | 0.3492 |
|  | V1V2-two step | V3V4-one step | 1,4 | 10.3244 | 0.1000 | 0.1000 | 0.7208 | 0.6509 |
|  |  | V3V4-two step | 1,4 | 3.7701 | 0.1000 | 0.1000 | 0.4852 | 0.3565 |
|  | V3V4-one step | V3V4-two step | 1,4 | 4.0413 | 0.1000 | 0.1000 | 0.5026 | 0.3782 |
| Jaccard | Kraken | MEGAN | 1,6 | 4.7135 | 0.0290 | 0.0375 | 0.4400 | 0.3466 |
|  |  | MetaPhlan | 1,6 | 4.3927 | 0.0258 | 0.0375 | 0.4227 | 0.3265 |
|  |  | MetaPhlan2 | 1,6 | 3.1463 | 0.0278 | 0.0375 | 0.3440 | 0.2347 |
|  |  | SortmeRNA | 1,6 | 6.0039 | 0.0317 | 0.0383 | 0.5002 | 0.4169 |
|  |  | V1V2-one step | 1,5 | 2.6229 | 0.0286 | 0.0375 | 0.3441 | 0.2129 |
|  |  | V1V2-two step | 1,5 | 2.8290 | 0.0286 | 0.0375 | 0.3614 | 0.2336 |
|  |  | V3V4-one step | 1,5 | 1.9825 | 0.0286 | 0.0375 | 0.2839 | 0.1407 |
|  |  | V3V4-two step | 1,5 | 2.6838 | 0.0286 | 0.0375 | 0.3493 | 0.2191 |
|  | MEGAN | MetaPhlan | 1,6 | 10.3012 | 0.0258 | 0.0375 | 0.6319 | 0.5706 |
|  |  | MetaPhlan2 | 1,6 | 47.0608 | 0.0292 | 0.0375 | 0.8869 | 0.8681 |
|  |  | SortmeRNA | 1,6 | 16.4193 | 0.0286 | 0.0375 | 0.7324 | 0.6878 |
|  |  | V1V2-one step | 1,5 | 9.2189 | 0.0286 | 0.0375 | 0.6484 | 0.5780 |
|  |  | V1V2-two step | 1,5 | 10.4510 | 0.0286 | 0.0375 | 0.6764 | 0.6117 |
|  |  | V3V4-one step | 1,5 | 7.5706 | 0.0286 | 0.0375 | 0.6022 | 0.5227 |
|  |  | V3V4-two step | 1,5 | 11.1724 | 0.0286 | 0.0375 | 0.6908 | 0.6290 |
|  | MetaPhlan | MetaPhlan2 | 1,6 | 13.8298 | 0.0257 | 0.0375 | 0.6974 | 0.6470 |
|  |  | SortmeRNA | 1,6 | 7.0631 | 0.0258 | 0.0375 | 0.5407 | 0.4641 |
|  |  | V1V2-one step | 1,5 | 5.2235 | 0.0286 | 0.0375 | 0.5109 | 0.4131 |
|  |  | V1V2-two step | 1,5 | 5.5890 | 0.0286 | 0.0375 | 0.5278 | 0.4334 |
|  |  | V3V4-one step | 1,5 | 4.8673 | 0.0286 | 0.0375 | 0.4933 | 0.3919 |
|  |  | V3V4-two step | 1,5 | 6.0857 | 0.0286 | 0.0375 | 0.5490 | 0.4588 |
|  | MetaPhlan2 | SortmeRNA | 1,6 | 22.0352 | 0.0319 | 0.0383 | 0.7860 | 0.7503 |
|  |  | V1V2-one step | 1,5 | 9.5392 | 0.0286 | 0.0375 | 0.6561 | 0.5873 |
|  |  | V1V2-two step | 1,5 | 11.1286 | 0.0286 | 0.0375 | 0.6900 | 0.6280 |
|  |  | V3V4-one step | 1,5 | 6.5090 | 0.0286 | 0.0375 | 0.5656 | 0.4787 |
|  |  | V3V4-two step | 1,5 | 10.6817 | 0.0286 | 0.0375 | 0.6812 | 0.6174 |
|  | SortmeRNA | V1V2-one step | 1,5 | 6.8379 | 0.0286 | 0.0375 | 0.5776 | 0.4932 |
|  |  | V1V2-two step | 1,5 | 7.5107 | 0.0286 | 0.0375 | 0.6003 | 0.5204 |
|  |  | V3V4-one step | 1,5 | 6.4127 | 0.0286 | 0.0375 | 0.5619 | 0.4743 |
|  |  | V3V4-two step | 1,5 | 8.1890 | 0.0286 | 0.0375 | 0.6209 | 0.5451 |
|  | V1V2-one step | V1V2-two step | 1,4 | 1.1205 | 0.3000 | 0.3000 | 0.2188 | 0.0235 |
|  |  | V3V4-one step | 1,4 | 1.8684 | 0.1000 | 0.1059 | 0.3184 | 0.1480 |
|  |  | V3V4-two step | 1,4 | 2.3890 | 0.1000 | 0.1059 | 0.3739 | 0.2174 |
|  | V1V2-two step | V3V4-one step | 1,4 | 1.7633 | 0.1000 | 0.1059 | 0.3060 | 0.1324 |
|  |  | V3V4-two step | 1,4 | 2.1644 | 0.1000 | 0.1059 | 0.3511 | 0.1889 |
|  | V3V4-one step | V3V4-two step | 1,4 | 1.3132 | 0.3000 | 0.3000 | 0.2472 | 0.0590 |
