## Supplemental Table S6 for "Comparative analysis of amplicon and metagenomic sequencing methods reveals key features in the evolution of animal metaorganisms"

| Data | Comparisons | *DF* | *F* | *P* | *P*_Hommel_ | *R*^2^ | adj. *R*^2^ |
| --- | --- | --- | --- | --- | --- | --- | --- |
| V1V2-one step | A.aerophoba/A.aurita | 1,8 | 6.7571 | 0.0063 | 0.0192 | 0.4579 | 0.3901 |
|  | A.aerophoba/C.elegans | 1,8 | 6.8457 | 0.0086 | 0.0258 | 0.4611 | 0.3938 |
|  | A.aerophoba/D.melanogaster(feces) | 1,8 | 40.1146 | 0.0082 | 0.0246 | 0.8337 | 0.8129 |
|  | A.aerophoba/D.melanogaster(gut) | 1,7 | 32.1926 | 0.0066 | 0.0198 | 0.8214 | 0.7959 |
|  | A.aerophoba/H.sapiens | 1,8 | 6.6935 | 0.0075 | 0.0225 | 0.4555 | 0.3875 |
|  | A.aerophoba/H.vulgaris | 1,8 | 10.8669 | 0.0067 | 0.0201 | 0.5760 | 0.5230 |
|  | A.aerophoba/M.leidyi | 1,8 | 7.1748 | 0.0080 | 0.0240 | 0.4728 | 0.4069 |
|  | A.aerophoba/M.musculus | 1,8 | 10.4923 | 0.0075 | 0.0225 | 0.5674 | 0.5133 |
|  | A.aerophoba/N.vectensis | 1,8 | 11.4223 | 0.0070 | 0.0210 | 0.5881 | 0.5366 |
|  | A.aerophoba/T.aestivum | 1,8 | 6.9159 | 0.0075 | 0.0225 | 0.4637 | 0.3966 |
|  | A.aurita/C.elegans | 1,8 | 2.5010 | 0.0400 | 0.0400 | 0.2382 | 0.1429 |
|  | A.aurita/D.melanogaster(feces) | 1,8 | 12.5533 | 0.0071 | 0.0213 | 0.6108 | 0.5621 |
|  | A.aurita/D.melanogaster(gut) | 1,7 | 9.8611 | 0.0088 | 0.0264 | 0.5848 | 0.5255 |
|  | A.aurita/H.sapiens | 1,8 | 3.7043 | 0.0085 | 0.0255 | 0.3165 | 0.2311 |
|  | A.aurita/H.vulgaris | 1,8 | 5.3480 | 0.0085 | 0.0255 | 0.4007 | 0.3257 |
|  | A.aurita/M.leidyi | 1,8 | 5.1386 | 0.0066 | 0.0198 | 0.3911 | 0.3150 |
|  | A.aurita/M.musculus | 1,8 | 5.3642 | 0.0079 | 0.0237 | 0.4014 | 0.3266 |
|  | A.aurita/N.vectensis | 1,8 | 4.0864 | 0.0081 | 0.0243 | 0.3381 | 0.2554 |
|  | A.aurita/T.aestivum | 1,8 | 3.7471 | 0.0079 | 0.0237 | 0.3190 | 0.2339 |
|  | C.elegans/D.melanogaster(feces) | 1,8 | 9.3734 | 0.0078 | 0.0234 | 0.5395 | 0.4820 |
|  | C.elegans/D.melanogaster(gut) | 1,7 | 7.3526 | 0.0065 | 0.0195 | 0.5123 | 0.4426 |
|  | C.elegans/H.sapiens | 1,8 | 3.1191 | 0.0090 | 0.0270 | 0.2805 | 0.1906 |
|  | C.elegans/H.vulgaris | 1,8 | 4.1899 | 0.0089 | 0.0267 | 0.3437 | 0.2617 |
|  | C.elegans/M.leidyi | 1,8 | 5.7953 | 0.0084 | 0.0252 | 0.4201 | 0.3476 |
|  | C.elegans/M.musculus | 1,8 | 4.3022 | 0.0074 | 0.0222 | 0.3497 | 0.2684 |
|  | C.elegans/N.vectensis | 1,8 | 5.0017 | 0.0069 | 0.0207 | 0.3847 | 0.3078 |
|  | C.elegans/T.aestivum | 1,8 | 2.9652 | 0.0088 | 0.0264 | 0.2704 | 0.1792 |
|  | D.melanogaster(feces)/D.melanogaster(gut) | 1,7 | 2.0198 | 0.0331 | 0.0400 | 0.2239 | 0.1131 |
|  | D.melanogaster(feces)/H.sapiens | 1,8 | 10.3448 | 0.0087 | 0.0261 | 0.5639 | 0.5094 |
|  | D.melanogaster(feces)/H.vulgaris | 1,8 | 16.2927 | 0.0091 | 0.0273 | 0.6707 | 0.6295 |
|  | D.melanogaster(feces)/M.leidyi | 1,8 | 28.3932 | 0.0072 | 0.0216 | 0.7802 | 0.7527 |
|  | D.melanogaster(feces)/M.musculus | 1,8 | 14.9828 | 0.0084 | 0.0252 | 0.6519 | 0.6084 |
|  | D.melanogaster(feces)/N.vectensis | 1,8 | 20.8783 | 0.0091 | 0.0273 | 0.7230 | 0.6883 |
|  | D.melanogaster(feces)/T.aestivum | 1,8 | 9.4941 | 0.0075 | 0.0225 | 0.5427 | 0.4855 |
|  | D.melanogaster(gut)/H.sapiens | 1,7 | 8.1258 | 0.0089 | 0.0267 | 0.5372 | 0.4711 |
|  | D.melanogaster(gut)/H.vulgaris | 1,7 | 12.8675 | 0.0078 | 0.0234 | 0.6477 | 0.5973 |
|  | D.melanogaster(gut)/M.leidyi | 1,7 | 22.5302 | 0.0074 | 0.0222 | 0.7630 | 0.7291 |
|  | D.melanogaster(gut)/M.musculus | 1,7 | 11.8103 | 0.0081 | 0.0243 | 0.6279 | 0.5747 |
|  | D.melanogaster(gut)/N.vectensis | 1,7 | 16.5405 | 0.0096 | 0.0288 | 0.7026 | 0.6602 |
|  | D.melanogaster(gut)/T.aestivum | 1,7 | 7.4592 | 0.0067 | 0.0201 | 0.5159 | 0.4467 |
|  | H.sapiens/H.vulgaris | 1,8 | 4.7768 | 0.0068 | 0.0204 | 0.3739 | 0.2956 |
|  | H.sapiens/M.leidyi | 1,8 | 5.8110 | 0.0089 | 0.0267 | 0.4208 | 0.3483 |
|  | H.sapiens/M.musculus | 1,8 | 3.4463 | 0.0082 | 0.0246 | 0.3011 | 0.2137 |
|  | H.sapiens/N.vectensis | 1,8 | 5.3730 | 0.0084 | 0.0252 | 0.4018 | 0.3270 |
|  | H.sapiens/T.aestivum | 1,8 | 3.2203 | 0.0092 | 0.0276 | 0.2870 | 0.1979 |
|  | H.vulgaris/M.leidyi | 1,8 | 9.1065 | 0.0076 | 0.0228 | 0.5323 | 0.4739 |
|  | H.vulgaris/M.musculus | 1,8 | 6.4230 | 0.0083 | 0.0249 | 0.4453 | 0.3760 |
|  | H.vulgaris/N.vectensis | 1,8 | 7.5359 | 0.0079 | 0.0237 | 0.4851 | 0.4207 |
|  | H.vulgaris/T.aestivum | 1,8 | 4.4645 | 0.0069 | 0.0207 | 0.3582 | 0.2780 |
|  | M.leidyi/M.musculus | 1,8 | 8.8666 | 0.0077 | 0.0231 | 0.5257 | 0.4664 |
|  | M.leidyi/N.vectensis | 1,8 | 9.4783 | 0.0085 | 0.0255 | 0.5423 | 0.4851 |
|  | M.leidyi/T.aestivum | 1,8 | 6.0292 | 0.0084 | 0.0252 | 0.4298 | 0.3585 |
|  | M.musculus/N.vectensis | 1,8 | 7.5725 | 0.0095 | 0.0285 | 0.4863 | 0.4221 |
|  | M.musculus/T.aestivum | 1,8 | 4.3456 | 0.0087 | 0.0261 | 0.3520 | 0.2710 |
|  | N.vectensis/T.aestivum | 1,8 | 5.1557 | 0.0083 | 0.0249 | 0.3919 | 0.3159 |
| V1V2-two step | A.aerophoba/A.aurita | 1,8 | 6.5506 | 0.0075 | 0.0450 | 0.4502 | 0.3815 |
|  | A.aerophoba/C.elegans | 1,8 | 7.3186 | 0.0060 | 0.0420 | 0.4778 | 0.4125 |
|  | A.aerophoba/D.melanogaster(feces) | 1,8 | 7.9092 | 0.0090 | 0.0472 | 0.4971 | 0.4343 |
|  | A.aerophoba/D.melanogaster(gut) | 1,7 | 8.2476 | 0.0074 | 0.0444 | 0.5409 | 0.4753 |
|  | A.aerophoba/H.sapiens | 1,8 | 6.8066 | 0.0078 | 0.0468 | 0.4597 | 0.3922 |
|  | A.aerophoba/H.vulgaris | 1,7 | 5.4813 | 0.0081 | 0.0472 | 0.4392 | 0.3590 |
|  | A.aerophoba/M.leidyi | 1,8 | 7.4016 | 0.0085 | 0.0472 | 0.4806 | 0.4156 |
|  | A.aerophoba/M.musculus | 1,8 | 10.1302 | 0.0079 | 0.0472 | 0.5587 | 0.5036 |
|  | A.aerophoba/N.vectensis | 1,8 | 5.1402 | 0.0098 | 0.0490 | 0.3912 | 0.3151 |
|  | A.aerophoba/T.aestivum | 1,8 | 9.5917 | 0.0090 | 0.0472 | 0.5452 | 0.4884 |
|  | A.aurita/C.elegans | 1,8 | 2.6866 | 0.0315 | 0.0642 | 0.2514 | 0.1578 |
|  | A.aurita/D.melanogaster(feces) | 1,8 | 3.8166 | 0.0088 | 0.0472 | 0.3230 | 0.2384 |
|  | A.aurita/D.melanogaster(gut) | 1,7 | 3.8328 | 0.0087 | 0.0472 | 0.3538 | 0.2615 |
|  | A.aurita/H.sapiens | 1,8 | 3.5440 | 0.0073 | 0.0441 | 0.3070 | 0.2204 |
|  | A.aurita/H.vulgaris | 1,7 | 2.8068 | 0.0058 | 0.0420 | 0.2862 | 0.1842 |
|  | A.aurita/M.leidyi | 1,8 | 4.0368 | 0.0089 | 0.0472 | 0.3354 | 0.2523 |
|  | A.aurita/M.musculus | 1,8 | 5.2448 | 0.0072 | 0.0441 | 0.3960 | 0.3205 |
|  | A.aurita/N.vectensis | 1,8 | 2.5633 | 0.0084 | 0.0472 | 0.2427 | 0.1480 |
|  | A.aurita/T.aestivum | 1,8 | 4.7303 | 0.0091 | 0.0472 | 0.3716 | 0.2930 |
|  | C.elegans/D.melanogaster(feces) | 1,8 | 3.4128 | 0.0067 | 0.0441 | 0.2990 | 0.2114 |
|  | C.elegans/D.melanogaster(gut) | 1,7 | 3.3385 | 0.0153 | 0.0571 | 0.3229 | 0.2262 |
|  | C.elegans/H.sapiens | 1,8 | 3.2184 | 0.0061 | 0.0427 | 0.2869 | 0.1977 |
|  | C.elegans/H.vulgaris | 1,7 | 2.2797 | 0.0428 | 0.0856 | 0.2457 | 0.1379 |
|  | C.elegans/M.leidyi | 1,8 | 5.5203 | 0.0098 | 0.0490 | 0.4083 | 0.3343 |
|  | C.elegans/M.musculus | 1,8 | 4.4817 | 0.0073 | 0.0441 | 0.3591 | 0.2789 |
|  | C.elegans/N.vectensis | 1,8 | 3.1165 | 0.0081 | 0.0472 | 0.2803 | 0.1904 |
|  | C.elegans/T.aestivum | 1,8 | 4.1037 | 0.0078 | 0.0468 | 0.3390 | 0.2564 |
|  | D.melanogaster(feces)/D.melanogaster(gut) | 1,7 | 1.0927 | 0.3038 | 0.3038 | 0.1350 | 0.0115 |
|  | D.melanogaster(feces)/H.sapiens | 1,8 | 3.6228 | 0.0054 | 0.0420 | 0.3117 | 0.2257 |
|  | D.melanogaster(feces)/H.vulgaris | 1,7 | 2.5160 | 0.0304 | 0.0642 | 0.2644 | 0.1593 |
|  | D.melanogaster(feces)/M.leidyi | 1,8 | 5.4092 | 0.0087 | 0.0472 | 0.4034 | 0.3288 |
|  | D.melanogaster(feces)/M.musculus | 1,8 | 4.5995 | 0.0078 | 0.0468 | 0.3651 | 0.2857 |
|  | D.melanogaster(feces)/N.vectensis | 1,8 | 3.3560 | 0.0067 | 0.0441 | 0.2955 | 0.2075 |
|  | D.melanogaster(feces)/T.aestivum | 1,8 | 4.2517 | 0.0079 | 0.0472 | 0.3470 | 0.2654 |
|  | D.melanogaster(gut)/H.sapiens | 1,7 | 3.5280 | 0.0076 | 0.0456 | 0.3351 | 0.2401 |
|  | D.melanogaster(gut)/H.vulgaris | 1,6 | 2.4110 | 0.0284 | 0.0642 | 0.2866 | 0.1678 |
|  | D.melanogaster(gut)/M.leidyi | 1,7 | 5.5165 | 0.0084 | 0.0472 | 0.4407 | 0.3608 |
|  | D.melanogaster(gut)/M.musculus | 1,7 | 4.5350 | 0.0076 | 0.0456 | 0.3932 | 0.3065 |
|  | D.melanogaster(gut)/N.vectensis | 1,7 | 3.2572 | 0.0160 | 0.0571 | 0.3176 | 0.2201 |
|  | D.melanogaster(gut)/T.aestivum | 1,7 | 4.1332 | 0.0070 | 0.0441 | 0.3712 | 0.2814 |
|  | H.sapiens/H.vulgaris | 1,7 | 2.4621 | 0.0077 | 0.0462 | 0.2602 | 0.1545 |
|  | H.sapiens/M.leidyi | 1,8 | 5.2489 | 0.0080 | 0.0472 | 0.3962 | 0.3207 |
|  | H.sapiens/M.musculus | 1,8 | 3.3252 | 0.0089 | 0.0472 | 0.2936 | 0.2053 |
|  | H.sapiens/N.vectensis | 1,8 | 3.1329 | 0.0076 | 0.0456 | 0.2814 | 0.1916 |
|  | H.sapiens/T.aestivum | 1,8 | 4.2834 | 0.0076 | 0.0456 | 0.3487 | 0.2673 |
|  | H.vulgaris/M.leidyi | 1,7 | 4.2092 | 0.0067 | 0.0441 | 0.3755 | 0.2863 |
|  | H.vulgaris/M.musculus | 1,7 | 3.0617 | 0.0075 | 0.0450 | 0.3043 | 0.2049 |
|  | H.vulgaris/N.vectensis | 1,7 | 2.1737 | 0.0159 | 0.0571 | 0.2370 | 0.1279 |
|  | H.vulgaris/T.aestivum | 1,7 | 2.9706 | 0.0079 | 0.0472 | 0.2979 | 0.1976 |
|  | M.leidyi/M.musculus | 1,8 | 7.6923 | 0.0081 | 0.0472 | 0.4902 | 0.4265 |
|  | M.leidyi/N.vectensis | 1,8 | 4.0730 | 0.0090 | 0.0472 | 0.3374 | 0.2545 |
|  | M.leidyi/T.aestivum | 1,8 | 6.9962 | 0.0075 | 0.0450 | 0.4665 | 0.3998 |
|  | M.musculus/N.vectensis | 1,8 | 4.3251 | 0.0073 | 0.0441 | 0.3509 | 0.2698 |
|  | M.musculus/T.aestivum | 1,8 | 5.6638 | 0.0080 | 0.0472 | 0.4145 | 0.3413 |
|  | N.vectensis/T.aestivum | 1,8 | 3.9181 | 0.0085 | 0.0472 | 0.3288 | 0.2448 |
| V3V4-one step | A.aerophoba/A.aurita | 1,8 | 10.8004 | 0.0079 | 0.0158 | 0.5745 | 0.5213 |
|  | A.aerophoba/C.elegans | 1,8 | 7.1365 | 0.0078 | 0.0156 | 0.4715 | 0.4054 |
|  | A.aerophoba/D.melanogaster(feces) | 1,8 | 35.8631 | 0.0083 | 0.0166 | 0.8176 | 0.7948 |
|  | A.aerophoba/D.melanogaster(gut) | 1,7 | 16.3971 | 0.0072 | 0.0150 | 0.7008 | 0.6581 |
|  | A.aerophoba/H.sapiens | 1,8 | 7.1021 | 0.0078 | 0.0156 | 0.4703 | 0.4041 |
|  | A.aerophoba/H.vulgaris | 1,8 | 11.7796 | 0.0063 | 0.0150 | 0.5955 | 0.5450 |
|  | A.aerophoba/M.leidyi | 1,8 | 8.2124 | 0.0081 | 0.0162 | 0.5066 | 0.4449 |
|  | A.aerophoba/M.musculus | 1,8 | 20.9174 | 0.0094 | 0.0188 | 0.7233 | 0.6888 |
|  | A.aerophoba/N.vectensis | 1,8 | 8.9339 | 0.0072 | 0.0150 | 0.5276 | 0.4685 |
|  | A.aerophoba/T.aestivum | 1,8 | 8.2251 | 0.0067 | 0.0150 | 0.5069 | 0.4453 |
|  | A.aurita/C.elegans | 1,8 | 4.2028 | 0.0077 | 0.0154 | 0.3444 | 0.2625 |
|  | A.aurita/D.melanogaster(feces) | 1,8 | 17.5530 | 0.0084 | 0.0168 | 0.6869 | 0.6478 |
|  | A.aurita/D.melanogaster(gut) | 1,7 | 9.3737 | 0.0073 | 0.0150 | 0.5725 | 0.5114 |
|  | A.aurita/H.sapiens | 1,8 | 5.0250 | 0.0079 | 0.0158 | 0.3858 | 0.3090 |
|  | A.aurita/H.vulgaris | 1,8 | 7.7982 | 0.0086 | 0.0172 | 0.4936 | 0.4303 |
|  | A.aurita/M.leidyi | 1,8 | 4.6541 | 0.0094 | 0.0188 | 0.3678 | 0.2888 |
|  | A.aurita/M.musculus | 1,8 | 12.4266 | 0.0074 | 0.0150 | 0.6084 | 0.5594 |
|  | A.aurita/N.vectensis | 1,8 | 7.3657 | 0.0090 | 0.0180 | 0.4794 | 0.4143 |
|  | A.aurita/T.aestivum | 1,8 | 5.7013 | 0.0087 | 0.0174 | 0.4161 | 0.3431 |
|  | C.elegans/D.melanogaster(feces) | 1,8 | 9.9465 | 0.0082 | 0.0164 | 0.5542 | 0.4985 |
|  | C.elegans/D.melanogaster(gut) | 1,7 | 5.7688 | 0.0081 | 0.0162 | 0.4518 | 0.3735 |
|  | C.elegans/H.sapiens | 1,8 | 3.3617 | 0.0093 | 0.0186 | 0.2959 | 0.2079 |
|  | C.elegans/H.vulgaris | 1,8 | 5.1391 | 0.0074 | 0.0150 | 0.3911 | 0.3150 |
|  | C.elegans/M.leidyi | 1,8 | 3.8341 | 0.0085 | 0.0170 | 0.3240 | 0.2395 |
|  | C.elegans/M.musculus | 1,8 | 7.6409 | 0.0091 | 0.0182 | 0.4885 | 0.4246 |
|  | C.elegans/N.vectensis | 1,8 | 5.5909 | 0.0075 | 0.0150 | 0.4114 | 0.3378 |
|  | C.elegans/T.aestivum | 1,8 | 3.7970 | 0.0063 | 0.0150 | 0.3219 | 0.2371 |
|  | D.melanogaster(feces)/D.melanogaster(gut) | 1,7 | 1.2056 | 0.1980 | 0.1980 | 0.1469 | 0.0251 |
|  | D.melanogaster(feces)/H.sapiens | 1,8 | 10.2590 | 0.0090 | 0.0180 | 0.5619 | 0.5071 |
|  | D.melanogaster(feces)/H.vulgaris | 1,8 | 19.0119 | 0.0096 | 0.0192 | 0.7038 | 0.6668 |
|  | D.melanogaster(feces)/M.leidyi | 1,8 | 12.3661 | 0.0086 | 0.0172 | 0.6072 | 0.5581 |
|  | D.melanogaster(feces)/M.musculus | 1,8 | 42.5522 | 0.0085 | 0.0170 | 0.8417 | 0.8220 |
|  | D.melanogaster(feces)/N.vectensis | 1,8 | 21.9116 | 0.0095 | 0.0190 | 0.7325 | 0.6991 |
|  | D.melanogaster(feces)/T.aestivum | 1,8 | 11.8267 | 0.0091 | 0.0182 | 0.5965 | 0.5461 |
|  | D.melanogaster(gut)/H.sapiens | 1,7 | 5.9145 | 0.0085 | 0.0170 | 0.4580 | 0.3805 |
|  | D.melanogaster(gut)/H.vulgaris | 1,7 | 10.1323 | 0.0099 | 0.0198 | 0.5914 | 0.5330 |
|  | D.melanogaster(gut)/M.leidyi | 1,7 | 6.8937 | 0.0100 | 0.0200 | 0.4962 | 0.4242 |
|  | D.melanogaster(gut)/M.musculus | 1,7 | 18.2554 | 0.0073 | 0.0150 | 0.7228 | 0.6832 |
|  | D.melanogaster(gut)/N.vectensis | 1,7 | 11.3091 | 0.0089 | 0.0178 | 0.6177 | 0.5631 |
|  | D.melanogaster(gut)/T.aestivum | 1,7 | 6.7668 | 0.0079 | 0.0158 | 0.4915 | 0.4189 |
|  | H.sapiens/H.vulgaris | 1,8 | 5.3437 | 0.0087 | 0.0174 | 0.4005 | 0.3255 |
|  | H.sapiens/M.leidyi | 1,8 | 3.9916 | 0.0064 | 0.0150 | 0.3329 | 0.2495 |
|  | H.sapiens/M.musculus | 1,8 | 5.7088 | 0.0082 | 0.0164 | 0.4164 | 0.3435 |
|  | H.sapiens/N.vectensis | 1,8 | 5.7373 | 0.0082 | 0.0164 | 0.4176 | 0.3448 |
|  | H.sapiens/T.aestivum | 1,8 | 3.8997 | 0.0075 | 0.0150 | 0.3277 | 0.2437 |
|  | H.vulgaris/M.leidyi | 1,8 | 5.8797 | 0.0075 | 0.0150 | 0.4236 | 0.3516 |
|  | H.vulgaris/M.musculus | 1,8 | 13.1163 | 0.0095 | 0.0190 | 0.6211 | 0.5738 |
|  | H.vulgaris/N.vectensis | 1,8 | 8.9099 | 0.0084 | 0.0168 | 0.5269 | 0.4678 |
|  | H.vulgaris/T.aestivum | 1,8 | 5.9706 | 0.0072 | 0.0150 | 0.4274 | 0.3558 |
|  | M.leidyi/M.musculus | 1,8 | 9.2979 | 0.0069 | 0.0150 | 0.5375 | 0.4797 |
|  | M.leidyi/N.vectensis | 1,8 | 5.9617 | 0.0091 | 0.0182 | 0.4270 | 0.3554 |
|  | M.leidyi/T.aestivum | 1,8 | 4.4791 | 0.0066 | 0.0150 | 0.3589 | 0.2788 |
|  | M.musculus/N.vectensis | 1,8 | 14.7449 | 0.0075 | 0.0150 | 0.6483 | 0.6043 |
|  | M.musculus/T.aestivum | 1,8 | 8.8512 | 0.0091 | 0.0182 | 0.5253 | 0.4659 |
|  | N.vectensis/T.aestivum | 1,8 | 6.4740 | 0.0082 | 0.0164 | 0.4473 | 0.3782 |
| V3V4-two step | A.aerophoba/A.aurita | 1,8 | 3.6151 | 0.0091 | 0.0637 | 0.3112 | 0.2251 |
|  | A.aerophoba/C.elegans | 1,8 | 2.6440 | 0.0077 | 0.0592 | 0.2484 | 0.1544 |
|  | A.aerophoba/D.melanogaster(feces) | 1,8 | 4.3367 | 0.0074 | 0.0592 | 0.3515 | 0.2705 |
|  | A.aerophoba/D.melanogaster(gut) | 1,7 | 3.9075 | 0.0080 | 0.0592 | 0.3582 | 0.2666 |
|  | A.aerophoba/H.sapiens | 1,8 | 3.0054 | 0.0070 | 0.0560 | 0.2731 | 0.1822 |
|  | A.aerophoba/H.vulgaris | 1,7 | 1.8196 | 0.0778 | 0.1398 | 0.2063 | 0.0929 |
|  | A.aerophoba/M.leidyi | 1,8 | 3.7680 | 0.0093 | 0.0651 | 0.3202 | 0.2352 |
|  | A.aerophoba/M.musculus | 1,8 | 5.5203 | 0.0098 | 0.0672 | 0.4083 | 0.3343 |
|  | A.aerophoba/N.vectensis | 1,7 | 1.5572 | 0.1398 | 0.1398 | 0.1820 | 0.0651 |
|  | A.aerophoba/T.aestivum | 1,8 | 3.2375 | 0.0078 | 0.0592 | 0.2881 | 0.1991 |
|  | A.aurita/C.elegans | 1,8 | 3.6508 | 0.0073 | 0.0584 | 0.3134 | 0.2275 |
|  | A.aurita/D.melanogaster(feces) | 1,8 | 6.5580 | 0.0085 | 0.0595 | 0.4505 | 0.3818 |
|  | A.aurita/D.melanogaster(gut) | 1,7 | 6.2319 | 0.0097 | 0.0672 | 0.4710 | 0.3954 |
|  | A.aurita/H.sapiens | 1,8 | 4.5552 | 0.0086 | 0.0602 | 0.3628 | 0.2832 |
|  | A.aurita/H.vulgaris | 1,7 | 2.9862 | 0.0091 | 0.0637 | 0.2990 | 0.1989 |
|  | A.aurita/M.leidyi | 1,8 | 4.6526 | 0.0078 | 0.0592 | 0.3677 | 0.2887 |
|  | A.aurita/M.musculus | 1,8 | 8.6980 | 0.0083 | 0.0592 | 0.5209 | 0.4610 |
|  | A.aurita/N.vectensis | 1,7 | 2.6768 | 0.0080 | 0.0592 | 0.2766 | 0.1733 |
|  | A.aurita/T.aestivum | 1,8 | 4.9675 | 0.0076 | 0.0592 | 0.3831 | 0.3060 |
|  | C.elegans/D.melanogaster(feces) | 1,8 | 4.1904 | 0.0078 | 0.0592 | 0.3437 | 0.2617 |
|  | C.elegans/D.melanogaster(gut) | 1,7 | 3.8621 | 0.0073 | 0.0584 | 0.3556 | 0.2635 |
|  | C.elegans/H.sapiens | 1,8 | 2.9430 | 0.0076 | 0.0592 | 0.2689 | 0.1776 |
|  | C.elegans/H.vulgaris | 1,7 | 1.8643 | 0.0384 | 0.1152 | 0.2103 | 0.0975 |
|  | C.elegans/M.leidyi | 1,8 | 3.8723 | 0.0072 | 0.0576 | 0.3262 | 0.2419 |
|  | C.elegans/M.musculus | 1,8 | 5.4850 | 0.0077 | 0.0592 | 0.4067 | 0.3326 |
|  | C.elegans/N.vectensis | 1,7 | 1.6707 | 0.0847 | 0.1398 | 0.1927 | 0.0773 |
|  | C.elegans/T.aestivum | 1,8 | 3.1819 | 0.0064 | 0.0533 | 0.2846 | 0.1951 |
|  | D.melanogaster(feces)/D.melanogaster(gut) | 1,7 | 3.8647 | 0.0222 | 0.0960 | 0.3557 | 0.2637 |
|  | D.melanogaster(feces)/H.sapiens | 1,8 | 4.9766 | 0.0102 | 0.0672 | 0.3835 | 0.3064 |
|  | D.melanogaster(feces)/H.vulgaris | 1,7 | 3.1889 | 0.0159 | 0.0795 | 0.3130 | 0.2148 |
|  | D.melanogaster(feces)/M.leidyi | 1,8 | 6.1797 | 0.0085 | 0.0595 | 0.4358 | 0.3653 |
|  | D.melanogaster(feces)/M.musculus | 1,8 | 9.0784 | 0.0066 | 0.0533 | 0.5316 | 0.4730 |
|  | D.melanogaster(feces)/N.vectensis | 1,7 | 2.8872 | 0.0146 | 0.0768 | 0.2920 | 0.1909 |
|  | D.melanogaster(feces)/T.aestivum | 1,8 | 5.1212 | 0.0076 | 0.0592 | 0.3903 | 0.3141 |
|  | D.melanogaster(gut)/H.sapiens | 1,7 | 4.5236 | 0.0083 | 0.0592 | 0.3926 | 0.3058 |
|  | D.melanogaster(gut)/H.vulgaris | 1,6 | 2.8949 | 0.0270 | 0.1080 | 0.3255 | 0.2130 |
|  | D.melanogaster(gut)/M.leidyi | 1,7 | 6.0195 | 0.0087 | 0.0609 | 0.4623 | 0.3855 |
|  | D.melanogaster(gut)/M.musculus | 1,7 | 8.6571 | 0.0072 | 0.0576 | 0.5529 | 0.4890 |
|  | D.melanogaster(gut)/N.vectensis | 1,6 | 2.5880 | 0.0292 | 0.1129 | 0.3014 | 0.1849 |
|  | D.melanogaster(gut)/T.aestivum | 1,7 | 4.5792 | 0.0067 | 0.0536 | 0.3955 | 0.3091 |
|  | H.sapiens/H.vulgaris | 1,7 | 2.2074 | 0.0076 | 0.0592 | 0.2397 | 0.1311 |
|  | H.sapiens/M.leidyi | 1,8 | 4.5382 | 0.0091 | 0.0637 | 0.3620 | 0.2822 |
|  | H.sapiens/M.musculus | 1,8 | 3.9634 | 0.0076 | 0.0592 | 0.3313 | 0.2477 |
|  | H.sapiens/N.vectensis | 1,7 | 1.9716 | 0.0086 | 0.0602 | 0.2198 | 0.1083 |
|  | H.sapiens/T.aestivum | 1,8 | 3.7181 | 0.0072 | 0.0576 | 0.3173 | 0.2320 |
|  | H.vulgaris/M.leidyi | 1,7 | 2.8369 | 0.0075 | 0.0592 | 0.2884 | 0.1867 |
|  | H.vulgaris/M.musculus | 1,7 | 4.1566 | 0.0061 | 0.0533 | 0.3726 | 0.2829 |
|  | H.vulgaris/N.vectensis | 1,6 | 1.1379 | 0.0296 | 0.1129 | 0.1594 | 0.0193 |
|  | H.vulgaris/T.aestivum | 1,7 | 2.2941 | 0.0079 | 0.0592 | 0.2468 | 0.1392 |
|  | M.leidyi/M.musculus | 1,8 | 8.4731 | 0.0087 | 0.0609 | 0.5144 | 0.4537 |
|  | M.leidyi/N.vectensis | 1,7 | 2.5934 | 0.0077 | 0.0592 | 0.2703 | 0.1661 |
|  | M.leidyi/T.aestivum | 1,8 | 4.8134 | 0.0057 | 0.0513 | 0.3757 | 0.2976 |
|  | M.musculus/N.vectensis | 1,7 | 3.7426 | 0.0098 | 0.0672 | 0.3484 | 0.2553 |
|  | M.musculus/T.aestivum | 1,8 | 6.7572 | 0.0070 | 0.0560 | 0.4579 | 0.3901 |
|  | N.vectensis/T.aestivum | 1,7 | 2.0760 | 0.0075 | 0.0592 | 0.2287 | 0.1186 |

Shading highlights insignificant comparisons after correction for multiple testing.
