## Supplemental Table S7 for "Comparative analysis of amplicon and metagenomic sequencing methods reveals key features in the evolution of animal metaorganisms"

| Genus | Data | Association | IndVal.g | *P* | *P*_FDR_ | Overlap with shotgun results |
| --- | --- | --- | --- | --- | --- | --- |
| *Acetatifactor* | V1V2-one step | *M.musculus* | 0.7746 | 0.0040 | 0.0088 |  |
|  | V1V2-two step |  | 0.8165 | 0.0007 | 0.0024 |  |
|  | V3V4-one step |  | 0.9678 | 0.0001 | 0.0004 |  |
|  | V3V4-two step |  | 0.9129 | 0.0001 | 0.0005 |  |
| *Acetobacter* | V1V2-one step | *D.mel.(gut)* | 0.9025 | 0.0001 | 0.0006 | MEGAN |
|  | V1V2-two step |  | 0.8628 | 0.0006 | 0.0021 | MetaPhlan |
|  | V3V4-one step |  | 1.0000 | 0.0001 | 0.0004 | MetaPhlan2 |
|  | V3V4-two step |  | 0.6672 | 0.0263 | 0.0456 | Kraken |
| *Acetobacteraceae* | V1V2-two step | *M.leidyi* | 0.7400 | 0.0019 | 0.0052 |  |
|  | V3V4-two step |  | 0.7118 | 0.0074 | 0.0155 |  |
| *Achromobacter* | V1V2-one step | *M.leidyi* | 0.7097 | 0.0244 | 0.0435 |  |
| *Acidimicrobiaceae* | V1V2-one step | *A.aurita* | 0.7746 | 0.0040 | 0.0088 |  |
|  | V1V2-two step |  | 0.7071 | 0.0043 | 0.0104 |  |
| *Acidimicrobiales* | V1V2-one step | *A.aerophoba* | 0.9929 | 0.0001 | 0.0006 |  |
|  | V1V2-two step |  | 0.9941 | 0.0001 | 0.0005 |  |
|  | V3V4-one step |  | 0.9270 | 0.0001 | 0.0004 |  |
|  | V3V4-two step |  | 0.5745 | 0.0237 | 0.0417 |  |
| *Acidobacteria* | V1V2-one step | *A.aerophoba* | 0.7746 | 0.0030 | 0.0069 |  |
| *Acidobacteria Gp3* | V1V2-one step | *A.aerophoba* | 1.0000 | 0.0001 | 0.0006 |  |
| *Acidovorax* | V1V2-one step | *H.vulgaris* | 0.9649 | 0.0001 | 0.0006 |  |
|  | V3V4-one step |  | 0.9185 | 0.0001 | 0.0004 |  |
| *Acinetobacter* | V1V2-two step | *D.mel.(feces)* | 0.7184 | 0.0246 | 0.0456 |  |
| *Actinobacteria* | V1V2-one step | *A.aerophoba* | 0.9752 | 0.0001 | 0.0006 |  |
|  | V1V2-two step |  | 0.9322 | 0.0002 | 0.0008 |  |
|  | V3V4-one step |  | 0.9218 | 0.0003 | 0.0008 |  |
|  | V3V4-two step |  | 0.6708 | 0.0081 | 0.0167 |  |
| *Actinobacteria,1* | V1V2-one step | *A.aurita* | 0.7130 | 0.0023 | 0.0054 |  |
|  | V1V2-two step | *A.aurita* | 0.8106 | 0.0005 | 0.0019 |  |
|  | V3V4-one step | *A.aerophoba* | 0.9742 | 0.0001 | 0.0004 |  |
|  | V3V4-two step | *A.aerophoba* | 0.7126 | 0.0096 | 0.0189 |  |
| *Actinomycetales* | V1V2-one step | *A.aurita* | 0.7204 | 0.0054 | 0.0108 |  |
|  | V1V2-two step |  | 0.6881 | 0.0023 | 0.0060 |  |
|  | V3V4-one step |  | 0.7541 | 0.0003 | 0.0008 |  |
|  | V3V4-two step |  | 0.7378 | 0.0006 | 0.0023 |  |
| *Aequorivita* | V1V2-one step | *N.vectensis* | 0.7746 | 0.0030 | 0.0069 |  |
| *Aerococcaceae* | V3V4-two step | *D.mel.(gut)* | 0.7071 | 0.0116 | 0.0218 |  |
| *Aeromonas* | V1V2-one step | *H.vulgaris* | 0.9375 | 0.0002 | 0.0007 | MEGAN |
|  | V1V2-two step |  | 0.8848 | 0.0007 | 0.0024 | MetaPhlan |
|  | V3V4-one step |  | 0.9307 | 0.0001 | 0.0004 | MetaPhlan2,Kraken |
| *Aestuariispira* | V3V4-one step | *N.vectensis* | 0.9857 | 0.0001 | 0.0004 |  |
| *Afipia* | V1V2-one step | *H.vulgaris* | 1.0000 | 0.0001 | 0.0006 |  |
|  | V1V2-two step |  | 0.6642 | 0.0087 | 0.0175 |  |
| *Alcaligenaceae* | V1V2-one step | *N.vectensis* | 1.0000 | 0.0002 | 0.0007 |  |
| *Alcanivorax* | V1V2-one step | *N.vectensis* | 1.0000 | 0.0002 | 0.0007 |  |
|  | V1V2-two step |  | 1.0000 | 0.0001 | 0.0005 |  |
|  | V3V4-one step |  | 1.0000 | 0.0001 | 0.0004 |  |
| *Algiphilus* | V1V2-one step | *N.vectensis* | 0.8944 | 0.0002 | 0.0007 |  |
|  | V3V4-one step |  | 0.8944 | 0.0003 | 0.0008 |  |
| *Algoriphagus* | V1V2-two step | *A.aurita* | 0.6642 | 0.0092 | 0.0182 |  |
| *Aliivibrio* | V1V2-one step | *A.aurita* | 0.8944 | 0.0004 | 0.0012 |  |
|  | V1V2-two step |  | 0.7237 | 0.0028 | 0.0071 |  |
|  | V3V4-one step |  | 0.8944 | 0.0003 | 0.0008 |  |
| *Alistipes* | V1V2-one step | *M.musculus* | 0.7295 | 0.0044 | 0.0093 |  |
|  | V1V2-two step | *M.musculus* | 0.7992 | 0.0035 | 0.0087 |  |
|  | V3V4-one step | *H.sapiens* | 0.7392 | 0.0023 | 0.0049 |  |
|  | V3V4-two step | *H.sapiens* | 0.8193 | 0.0010 | 0.0034 |  |
| *Alphaproteobacteria* | V1V2-one step | *A.aerophoba* | 0.8239 | 0.0001 | 0.0006 |  |
|  | V1V2-two step | *N.vectensis* | 0.7713 | 0.0037 | 0.0091 |  |
|  | V3V4-one step | *A.aerophoba* | 0.6639 | 0.0014 | 0.0031 |  |
|  | V3V4-two step | *A.aurita* | 0.6112 | 0.0223 | 0.0397 |  |
| *Alteromonadaceae* | V3V4-one step | *M.leidyi* | 0.7385 | 0.0036 | 0.0073 |  |
| *Alteromonadales* | V1V2-two step | *M.leidyi* | 0.7659 | 0.0029 | 0.0073 |  |
|  | V3V4-one step | *A.aurita* | 0.7746 | 0.0054 | 0.0099 |  |
|  | V3V4-two step | *A.aurita* | 0.7071 | 0.0047 | 0.0112 |  |
| *Alteromonas* | V1V2-one step | *M.leidyi* | 0.8406 | 0.0001 | 0.0006 |  |
|  | V1V2-two step |  | 0.6838 | 0.0086 | 0.0174 |  |
|  | V3V4-one step |  | 0.9019 | 0.0001 | 0.0004 |  |
|  | V3V4-two step |  | 0.9997 | 0.0001 | 0.0005 |  |
| *Amaricoccus* | V3V4-one step | *D.mel.(gut)* | 0.6776 | 0.0136 | 0.0233 |  |
| *Amphritea* | V3V4-one step | *A.aurita* | 0.8536 | 0.0001 | 0.0004 |  |
|  | V3V4-two step |  | 0.6804 | 0.0047 | 0.0112 |  |
| *Anaeroplasma* | V3V4-one step | *M.musculus* | 0.7746 | 0.0038 | 0.0075 |  |
|  | V3V4-two step |  | 0.7071 | 0.0055 | 0.0123 |  |
| *Anaerostipes* | V1V2-one step | *H.sapiens* | 0.8875 | 0.0005 | 0.0014 |  |
|  | V1V2-two step |  | 0.9129 | 0.0001 | 0.0005 |  |
|  | V3V4-one step |  | 0.9982 | 0.0001 | 0.0004 |  |
|  | V3V4-two step |  | 0.9129 | 0.0001 | 0.0005 |  |
| *Anaerotruncus* | V1V2-two step | *H.sapiens* | 0.8165 | 0.0009 | 0.0030 |  |
|  | V3V4-one step |  | 0.6000 | 0.0207 | 0.0345 |  |
|  | V3V4-two step |  | 0.6325 | 0.0167 | 0.0309 |  |
| *Arcobacter* | V1V2-one step | *A.aurita* | 0.7859 | 0.0002 | 0.0007 |  |
|  | V1V2-two step |  | 0.8336 | 0.0005 | 0.0019 |  |
|  | V3V4-one step |  | 0.8183 | 0.0005 | 0.0012 |  |
|  | V3V4-two step |  | 0.8676 | 0.0001 | 0.0005 |  |
| *Arenibacter* | V1V2-one step | *N.vectensis* | 0.5721 | 0.0237 | 0.0425 |  |
|  | V3V4-two step | *M.leidyi* | 0.6325 | 0.0287 | 0.0489 |  |
| *Arthrobacter* | V1V2-one step | *C.elegans* | 0.8926 | 0.0022 | 0.0052 |  |
|  | V3V4-one step |  | 0.7716 | 0.0130 | 0.0226 |  |
|  | V3V4-two step |  | 0.9116 | 0.0008 | 0.0030 |  |
| *Aureispira* | V1V2-one step | *N.vectensis* | 1.0000 | 0.0002 | 0.0007 |  |
|  | V1V2-two step |  | 0.8933 | 0.0001 | 0.0005 |  |
|  | V3V4-one step |  | 1.0000 | 0.0001 | 0.0004 |  |
| *Bacillus* | V1V2-one step | *D.mel.(gut)* | 0.5716 | 0.0144 | 0.0270 |  |
|  | V1V2-two step | *D.mel.(gut)* | 0.7315 | 0.0030 | 0.0075 |  |
|  | V3V4-one step | *D.mel.(gut)* | 0.7149 | 0.0010 | 0.0023 |  |
|  | V3V4-two step | *D.mel.(feces)* | 0.8588 | 0.0001 | 0.0005 |  |
| *Bacteria* | V1V2-one step | *M.leidyi* | 0.7288 | 0.0001 | 0.0006 |  |
|  | V1V2-two step | *M.leidyi* | 0.6850 | 0.0001 | 0.0005 |  |
|  | V3V4-one step | *A.aerophoba* | 0.8099 | 0.0001 | 0.0004 |  |
|  | V3V4-two step | *A.aerophoba* | 0.6717 | 0.0283 | 0.0485 |  |
| *Bacteriovoracaceae* | V1V2-one step | *N.vectensis* | 0.8944 | 0.0002 | 0.0007 |  |
|  | V1V2-two step |  | 0.5979 | 0.0273 | 0.0494 |  |
| *Bacteroidales* | V1V2-one step | *M.musculus* | 0.9077 | 0.0001 | 0.0006 |  |
|  | V1V2-two step |  | 0.9029 | 0.0001 | 0.0005 |  |
| *Bacteroides* | V1V2-one step | *H.sapiens* | 0.9717 | 0.0001 | 0.0006 | MEGAN |
|  | V1V2-two step |  | 0.9475 | 0.0001 | 0.0005 | MetaPhlan |
|  | V3V4-one step |  | 0.9747 | 0.0001 | 0.0004 | MetaPhlan2 |
|  | V3V4-two step |  | 0.9856 | 0.0001 | 0.0005 | Kraken |
| *Bacteroidetes* | V1V2-one step | *M.musculus* | 0.8849 | 0.0001 | 0.0006 |  |
|  | V1V2-two step | *M.musculus* | 0.7372 | 0.0057 | 0.0125 |  |
|  | V3V4-one step | *H.vulgaris* | 0.9697 | 0.0003 | 0.0008 |  |
| *Barnesiella* | V1V2-one step | *H.sapiens* | 0.8874 | 0.0004 | 0.0012 |  |
|  | V1V2-two step |  | 0.9741 | 0.0001 | 0.0005 |  |
|  | V3V4-one step |  | 0.7503 | 0.0016 | 0.0035 |  |
|  | V3V4-two step |  | 0.8984 | 0.0002 | 0.0009 |  |
| *Bdellovibrio* | V1V2-one step | *H.vulgaris* | 0.7888 | 0.0006 | 0.0016 |  |
|  | V1V2-two step |  | 0.7071 | 0.0068 | 0.0139 |  |
|  | V3V4-one step |  | 1.0000 | 0.0001 | 0.0004 |  |
| *Betaproteobacteria* | V1V2-one step | *M.leidyi* | 0.8798 | 0.0001 | 0.0006 |  |
|  | V1V2-two step | *M.leidyi* | 0.9356 | 0.0001 | 0.0005 |  |
|  | V3V4-one step | *A.aurita* | 0.6701 | 0.0021 | 0.0045 |  |
| *Bifidobacterium* | V1V2-one step | *H.sapiens* | 0.8944 | 0.0003 | 0.0010 |  |
|  | V1V2-two step |  | 0.7262 | 0.0018 | 0.0052 |  |
|  | V3V4-two step |  | 0.8165 | 0.0011 | 0.0037 |  |
| *Bizionia* | V3V4-one step | *N.vectensis* | 0.8944 | 0.0002 | 0.0006 |  |
| *Blautia* | V1V2-one step | *H.sapiens* | 0.9796 | 0.0001 | 0.0006 |  |
|  | V1V2-two step |  | 0.8923 | 0.0007 | 0.0024 |  |
|  | V3V4-one step |  | 0.8878 | 0.0001 | 0.0004 |  |
|  | V3V4-two step |  | 1.0000 | 0.0001 | 0.0005 |  |
| *Bosea* | V3V4-one step | *H.vulgaris* | 0.8627 | 0.0005 | 0.0012 |  |
| *Bradyrhizobium* | V1V2-one step | *D.mel.(feces)* | 0.7362 | 0.0012 | 0.0031 | Kraken |
|  | V1V2-two step |  | 0.7701 | 0.0019 | 0.0052 |  |
|  | V3V4-one step |  | 0.7385 | 0.0025 | 0.0053 |  |
|  | V3V4-two step |  | 0.9337 | 0.0001 | 0.0005 |  |
| *Brochothrix* | V3V4-one step | *D.mel.(feces)* | 0.6852 | 0.0087 | 0.0154 |  |
| *Brumimicrobium* | V3V4-one step | *M.leidyi* | 0.6606 | 0.0137 | 0.0233 |  |
| *Burkholderia* | V1V2-one step | *D.mel.(gut)* | 0.5446 | 0.0169 | 0.0310 |  |
|  | V1V2-two step | *M.leidyi* | 0.7456 | 0.0026 | 0.0067 |  |
|  | V3V4-one step | *M.leidyi* | 0.7375 | 0.0002 | 0.0006 |  |
|  | V3V4-two step | *M.leidyi* | 0.8533 | 0.0001 | 0.0005 |  |
| *Burkholderiales* | V1V2-one step | *H.vulgaris* | 0.9282 | 0.0001 | 0.0006 |  |
|  | V3V4-one step |  | 0.8326 | 0.0002 | 0.0006 |  |
| *Butyricicoccus* | V1V2-one step | *M.musculus* | 0.8724 | 0.0001 | 0.0006 |  |
|  | V1V2-two step |  | 0.7206 | 0.0018 | 0.0052 |  |
|  | V3V4-one step |  | 0.7227 | 0.0011 | 0.0025 |  |
|  | V3V4-two step |  | 0.8346 | 0.0001 | 0.0005 |  |
| *Butyricimonas* | V1V2-one step | *H.sapiens* | 0.7746 | 0.0043 | 0.0091 |  |
|  | V1V2-two step |  | 0.7071 | 0.0065 | 0.0138 |  |
| *Caenimonas* | V1V2-one step | *H.vulgaris* | 0.8341 | 0.0006 | 0.0016 |  |
|  | V1V2-two step |  | 0.7071 | 0.0068 | 0.0139 |  |
| *Caldilineaceae* | V1V2-one step | *A.aurita* | 0.8944 | 0.0001 | 0.0006 |  |
|  | V1V2-two step | *A.aurita* | 0.9129 | 0.0001 | 0.0005 |  |
|  | V3V4-one step | *A.aerophoba* | 1.0000 | 0.0002 | 0.0006 |  |
|  | V3V4-two step | *A.aerophoba* | 0.8765 | 0.0003 | 0.0013 |  |
| *Campylobacterales* | V3V4-one step | *A.aurita* | 0.8944 | 0.0003 | 0.0008 |  |
|  | V3V4-two step |  | 0.9129 | 0.0002 | 0.0009 |  |
| *Candidatus Pelagibacter* | V1V2-two step | *M.leidyi* | 0.7416 | 0.0002 | 0.0008 |  |
|  | V3V4-one step |  | 0.9557 | 0.0001 | 0.0004 |  |
|  | V3V4-two step |  | 0.9567 | 0.0001 | 0.0005 |  |
| *Carnobacterium* | V3V4-two step | *C.elegans* | 0.7071 | 0.0049 | 0.0114 |  |
| *Castellaniella* | V3V4-one step | *N.vectensis* | 1.0000 | 0.0001 | 0.0004 |  |
| *Caulobacter* | V3V4-one step | *H.vulgaris* | 0.5958 | 0.0213 | 0.0352 |  |
| *Chitinophagaceae* | V1V2-one step | *N.vectensis* | 0.8920 | 0.0002 | 0.0007 |  |
|  | V1V2-two step |  | 0.6261 | 0.0234 | 0.0436 |  |
|  | V3V4-one step |  | 0.9447 | 0.0001 | 0.0004 |  |
| *Chlamydiales* | V3V4-one step | *N.vectensis* | 0.9450 | 0.0001 | 0.0004 |  |
| *Chloroflexales* | V1V2-two step | *A.aurita* | 0.7071 | 0.0053 | 0.0118 |  |
| *Chloroflexi* | V1V2-one step | *A.aurita* | 0.7746 | 0.0040 | 0.0088 |  |
|  | V1V2-two step | *A.aurita* | 0.8165 | 0.0007 | 0.0024 |  |
|  | V3V4-one step | *A.aerophoba* | 1.0000 | 0.0002 | 0.0006 |  |
|  | V3V4-two step | *A.aerophoba* | 0.7696 | 0.0030 | 0.0075 |  |
| *Chromatiaceae* | V1V2-two step | *A.aurita* | 0.7071 | 0.0047 | 0.0109 |  |
| *Chromatiales* | V3V4-one step | *A.aerophoba* | 1.0000 | 0.0002 | 0.0006 |  |
|  | V3V4-two step |  | 0.7746 | 0.0024 | 0.0064 |  |
| *Chryseobacterium* | V1V2-one step | *C.elegans* | 0.6838 | 0.0127 | 0.0241 |  |
|  | V3V4-one step | *D.mel.(gut)* | 0.6919 | 0.0081 | 0.0145 |  |
|  | V3V4-two step | *C.elegans* | 0.7143 | 0.0060 | 0.0131 |  |
| *Chryseomicrobium* | V1V2-one step | *N.vectensis* | 1.0000 | 0.0002 | 0.0007 |  |
|  | V3V4-one step |  | 0.7833 | 0.0009 | 0.0021 |  |
| *Clostridia* | V3V4-one step | *M.musculus* | 0.8927 | 0.0001 | 0.0004 |  |
|  | V3V4-two step |  | 0.9401 | 0.0001 | 0.0005 |  |
| *Clostridiales* | V1V2-one step | *M.musculus* | 0.9793 | 0.0001 | 0.0006 |  |
|  | V1V2-two step | *M.musculus* | 0.9805 | 0.0001 | 0.0005 |  |
|  | V3V4-one step | *H.sapiens* | 0.6693 | 0.0186 | 0.0314 |  |
|  | V3V4-two step | *M.musculus* | 0.8680 | 0.0019 | 0.0055 |  |
| *Clostridium IV* | V1V2-one step | *H.sapiens* | 1.0000 | 0.0001 | 0.0006 |  |
|  | V1V2-two step |  | 1.0000 | 0.0001 | 0.0005 |  |
|  | V3V4-one step |  | 0.9074 | 0.0001 | 0.0004 |  |
|  | V3V4-two step |  | 0.8385 | 0.0005 | 0.0020 |  |
| *Clostridium XlVa* | V1V2-one step | *H.sapiens* | 0.9513 | 0.0002 | 0.0007 |  |
|  | V1V2-two step | *H.sapiens* | 0.9648 | 0.0002 | 0.0008 |  |
|  | V3V4-one step | *M.musculus* | 0.9570 | 0.0001 | 0.0004 |  |
|  | V3V4-two step | *M.musculus* | 0.9097 | 0.0001 | 0.0005 |  |
| *Clostridium XlVb* | V1V2-one step | *M.musculus* | 0.6273 | 0.0182 | 0.0332 |  |
|  | V1V2-two step | *H.sapiens* | 0.7037 | 0.0028 | 0.0071 |  |
|  | V3V4-one step | *H.sapiens* | 0.8341 | 0.0005 | 0.0012 |  |
|  | V3V4-two step | *H.sapiens* | 0.7683 | 0.0012 | 0.0038 |  |
| *Clostridium XVIII* | V1V2-one step | *M.musculus* | 0.7906 | 0.0036 | 0.0083 |  |
|  | V1V2-two step | *M.musculus* | 0.8201 | 0.0020 | 0.0054 |  |
|  | V3V4-one step | *H.sapiens* | 0.8132 | 0.0010 | 0.0023 |  |
|  | V3V4-two step | *H.sapiens* | 0.7471 | 0.0025 | 0.0066 |  |
| *Collinsella* | V1V2-one step | *H.sapiens* | 0.7746 | 0.0043 | 0.0091 |  |
|  | V3V4-one step |  | 0.7108 | 0.0090 | 0.0158 |  |
|  | V3V4-two step |  | 0.8165 | 0.0012 | 0.0038 |  |
| *Colwelliaceae* | V3V4-one step | *N.vectensis* | 0.9970 | 0.0001 | 0.0004 |  |
| *Comamonadaceae* | V1V2-one step | *H.vulgaris* | 0.9294 | 0.0002 | 0.0007 |  |
|  | V1V2-two step |  | 0.7257 | 0.0053 | 0.0118 |  |
|  | V3V4-one step |  | 0.9904 | 0.0001 | 0.0004 |  |
| *Comamonas* | V1V2-one step | *C.elegans* | 0.8935 | 0.0018 | 0.0044 |  |
|  | V1V2-two step |  | 0.9724 | 0.0016 | 0.0047 |  |
| *Coprobacillus* | V1V2-two step | *M.musculus* | 0.7071 | 0.0047 | 0.0109 |  |
| *Coprobacter* | V1V2-one step | *H.sapiens* | 0.7746 | 0.0049 | 0.0099 |  |
| *Coprococcus* | V3V4-one step | *H.sapiens* | 0.8944 | 0.0002 | 0.0006 |  |
| *Coriobacteriaceae* | V1V2-one step | *M.musculus* | 0.7625 | 0.0004 | 0.0012 |  |
|  | V1V2-two step | *M.musculus* | 0.8797 | 0.0001 | 0.0005 |  |
|  | V3V4-one step | *H.sapiens* | 0.6581 | 0.0308 | 0.0496 |  |
|  | V3V4-two step | *M.musculus* | 0.6729 | 0.0054 | 0.0123 |  |
| *Corynebacterium* | V1V2-one step | *M.leidyi* | 0.6015 | 0.0167 | 0.0308 |  |
|  | V3V4-two step | *D.mel.(feces)* | 0.6167 | 0.0235 | 0.0416 |  |
| *Coxiella* | V1V2-one step | *A.aerophoba* | 0.7071 | 0.0022 | 0.0052 |  |
|  | V1V2-two step | *A.aerophoba* | 0.8612 | 0.0001 | 0.0005 |  |
|  | V3V4-one step | *N.vectensis* | 0.6882 | 0.0017 | 0.0037 |  |
|  | V3V4-two step | *A.aurita* | 0.8165 | 0.0012 | 0.0038 |  |
| *Coxiellaceae* | V1V2-one step | *N.vectensis* | 0.8944 | 0.0003 | 0.0010 |  |
|  | V1V2-two step |  | 0.7746 | 0.0015 | 0.0045 |  |
| *Croceibacter* | V1V2-one step | *N.vectensis* | 0.9891 | 0.0002 | 0.0007 |  |
|  | V1V2-two step |  | 0.8944 | 0.0001 | 0.0005 |  |
|  | V3V4-one step |  | 0.8944 | 0.0003 | 0.0008 |  |
| *Crocinitomix* | V3V4-one step | *M.leidyi* | 0.8412 | 0.0002 | 0.0006 |  |
|  | V3V4-two step | *A.aurita* | 0.7427 | 0.0030 | 0.0075 |  |
| *Cryomorphaceae* | V1V2-one step | *N.vectensis* | 0.9272 | 0.0002 | 0.0007 |  |
|  | V1V2-two step | *N.vectensis* | 0.7166 | 0.0014 | 0.0043 |  |
|  | V3V4-one step | *N.vectensis* | 0.9403 | 0.0001 | 0.0004 |  |
|  | V3V4-two step | *M.leidyi* | 0.7272 | 0.0014 | 0.0042 |  |
| *Curtobacterium* | V1V2-two step | *T.aestivum* | 1.0000 | 0.0001 | 0.0005 |  |
| *Curvibacter* | V1V2-one step | *H.vulgaris* | 0.9947 | 0.0001 | 0.0006 |  |
|  | V1V2-two step | *H.vulgaris* | 0.9883 | 0.0013 | 0.0041 |  |
|  | V3V4-one step | *M.leidyi* | 0.6019 | 0.0030 | 0.0062 |  |
| *Cystobacteraceae* | V3V4-one step | *N.vectensis* | 0.8944 | 0.0003 | 0.0008 |  |
| *Cytophaga* | V1V2-two step | *N.vectensis* | 0.7746 | 0.0015 | 0.0045 |  |
| *Cytophagaceae* | V1V2-one step | *H.vulgaris* | 0.8746 | 0.0006 | 0.0016 |  |
|  | V3V4-one step |  | 0.9989 | 0.0001 | 0.0004 |  |
| *Cytophagales* | V3V4-one step | *N.vectensis* | 0.8660 | 0.0001 | 0.0004 |  |
| *Deltaproteobacteria* | V1V2-one step | *A.aerophoba* | 0.6979 | 0.0013 | 0.0033 |  |
|  | V3V4-one step |  | 0.9731 | 0.0001 | 0.0004 |  |
|  | V3V4-two step |  | 0.7083 | 0.0052 | 0.0120 |  |
| *Desulfovibrio* | V1V2-one step | *H.sapiens* | 0.7746 | 0.0043 | 0.0091 |  |
|  | V3V4-one step |  | 0.7746 | 0.0045 | 0.0086 |  |
|  | V3V4-two step |  | 0.7071 | 0.0040 | 0.0098 |  |
| *Desulfovibrionaceae* | V1V2-two step | *H.sapiens* | 0.7071 | 0.0065 | 0.0138 |  |
|  | V3V4-one step | *M.musculus* | 0.7681 | 0.0039 | 0.0076 |  |
|  | V3V4-two step | *M.musculus* | 0.7058 | 0.0041 | 0.0099 |  |
| *Desulfovibrionales* | V1V2-two step | *M.musculus* | 0.7071 | 0.0045 | 0.0107 |  |
|  | V3V4-two step |  | 0.6782 | 0.0065 | 0.0140 |  |
| *Dialister* | V1V2-one step | *H.sapiens* | 0.7739 | 0.0055 | 0.0109 |  |
|  | V3V4-one step |  | 0.8944 | 0.0003 | 0.0008 |  |
|  | V3V4-two step |  | 0.8165 | 0.0012 | 0.0038 |  |
| *Diplorickettsia* | V1V2-two step | *A.aurita* | 0.7071 | 0.0060 | 0.0130 |  |
|  | V3V4-one step |  | 0.6211 | 0.0188 | 0.0316 |  |
|  | V3V4-two step |  | 0.8165 | 0.0009 | 0.0032 |  |
| *Dorea* | V1V2-one step | *H.sapiens* | 0.8944 | 0.0004 | 0.0012 |  |
|  | V1V2-two step |  | 0.9083 | 0.0003 | 0.0012 |  |
|  | V3V4-one step |  | 1.0000 | 0.0001 | 0.0004 |  |
|  | V3V4-two step |  | 0.9969 | 0.0001 | 0.0005 |  |
| *Dyadobacter* | V1V2-two step | *H.vulgaris* | 0.7071 | 0.0068 | 0.0139 |  |
|  | V3V4-one step |  | 0.7078 | 0.0211 | 0.0350 |  |
| *Ectothiorhodospiraceae* | V3V4-one step | *A.aerophoba* | 0.7746 | 0.0037 | 0.0074 |  |
| *Eisenbergiella* | V3V4-one step | *H.sapiens* | 0.7327 | 0.0098 | 0.0171 |  |
| *Emticicia* | V3V4-one step | *H.vulgaris* | 0.8944 | 0.0004 | 0.0010 |  |
| *Endozoicomonas* | V3V4-one step | *A.aerophoba* | 0.7268 | 0.0020 | 0.0043 |  |
| *Enhydrobacter* | V1V2-one step | *M.leidyi* | 0.7098 | 0.0003 | 0.0010 |  |
|  | V1V2-two step | *D.mel.(gut)* | 0.8108 | 0.0022 | 0.0059 |  |
|  | V3V4-one step | *D.mel.(feces)* | 0.6203 | 0.0202 | 0.0338 |  |
| *Enterobacteriaceae* | V1V2-one step | *A.aurita* | 0.9646 | 0.0011 | 0.0028 |  |
| *Enterococcus* | V3V4-one step | *M.leidyi* | 0.6565 | 0.0255 | 0.0418 |  |
| *Enterorhabdus* | V1V2-one step | *M.musculus* | 1.0000 | 0.0001 | 0.0006 |  |
|  | V1V2-two step |  | 1.0000 | 0.0001 | 0.0005 |  |
|  | V3V4-one step |  | 1.0000 | 0.0001 | 0.0004 |  |
|  | V3V4-two step |  | 1.0000 | 0.0001 | 0.0005 |  |
| *Enterovibrio* | V1V2-one step | *A.aurita* | 0.7746 | 0.0048 | 0.0099 |  |
|  | V3V4-one step |  | 0.7454 | 0.0008 | 0.0019 |  |
|  | V3V4-two step |  | 0.6926 | 0.0019 | 0.0055 |  |
| *Erysipelotrichaceae* | V1V2-two step | *H.sapiens* | 0.6054 | 0.0217 | 0.0407 |  |
| *Erysipelotrichaceae incertae sedis* | V3V4-one step | *M.musculus* | 0.6583 | 0.0134 | 0.0232 |  |
| *Escherichia/Shigella* | V1V2-one step | *C.elegans* | 0.8131 | 0.0010 | 0.0026 | MEGAN,Kraken |
|  | V1V2-two step |  | 0.7781 | 0.0016 | 0.0047 | MEGAN,Kraken |
|  | V3V4-one step |  | 0.8731 | 0.0007 | 0.0017 | MEGAN,Kraken |
|  | V3V4-two step |  | 0.8645 | 0.0007 | 0.0026 | MEGAN,Kraken |
| *Exiguobacterium* | V1V2-one step | *N.vectensis* | 1.0000 | 0.0002 | 0.0007 |  |
|  | V1V2-two step |  | 0.9427 | 0.0001 | 0.0005 |  |
|  | V3V4-one step |  | 0.9972 | 0.0001 | 0.0004 |  |
| *Faecalibacterium* | V1V2-one step | *H.sapiens* | 0.9990 | 0.0001 | 0.0006 |  |
|  | V1V2-two step |  | 0.9841 | 0.0001 | 0.0005 |  |
|  | V3V4-one step |  | 0.9969 | 0.0001 | 0.0004 |  |
|  | V3V4-two step |  | 0.9989 | 0.0001 | 0.0005 |  |
| *Faecalicoccus* | V1V2-one step | *H.sapiens* | 0.7746 | 0.0049 | 0.0099 |  |
| *Firmicutes* | V1V2-one step | *M.musculus* | 0.9267 | 0.0001 | 0.0006 |  |
|  | V1V2-two step |  | 0.9022 | 0.0001 | 0.0005 |  |
| *Flavobacteriaceae* | V1V2-one step | *M.leidyi* | 0.6621 | 0.0008 | 0.0022 |  |
|  | V1V2-two step | *M.leidyi* | 0.7396 | 0.0003 | 0.0012 |  |
|  | V3V4-one step | *N.vectensis* | 0.6600 | 0.0019 | 0.0041 |  |
|  | V3V4-two step | *M.leidyi* | 0.8518 | 0.0001 | 0.0005 |  |
| *Flavobacteriales* | V1V2-one step | *M.leidyi* | 0.7375 | 0.0021 | 0.0051 |  |
|  | V1V2-two step |  | 0.7626 | 0.0003 | 0.0012 |  |
|  | V3V4-one step |  | 0.8496 | 0.0003 | 0.0008 |  |
|  | V3V4-two step |  | 0.8271 | 0.0001 | 0.0005 |  |
| *Flavobacterium* | V1V2-one step | *M.leidyi* | 0.8652 | 0.0001 | 0.0006 |  |
|  | V1V2-two step |  | 0.7595 | 0.0002 | 0.0008 |  |
|  | V3V4-one step |  | 0.9691 | 0.0001 | 0.0004 |  |
|  | V3V4-two step |  | 0.9886 | 0.0001 | 0.0005 |  |
| *Flavonifractor* | V1V2-one step | *H.sapiens* | 1.0000 | 0.0001 | 0.0006 |  |
|  | V1V2-two step |  | 1.0000 | 0.0001 | 0.0005 |  |
|  | V3V4-one step |  | 0.8449 | 0.0005 | 0.0012 |  |
|  | V3V4-two step |  | 0.9634 | 0.0001 | 0.0005 |  |
| *Flectobacillus* | V1V2-one step | *H.vulgaris* | 0.8931 | 0.0004 | 0.0012 |  |
|  | V3V4-one step |  | 0.8944 | 0.0004 | 0.0010 |  |
| *Fluviicola* | V1V2-one step | *A.aurita* | 0.7494 | 0.0049 | 0.0099 |  |
|  | V1V2-two step |  | 0.6934 | 0.0023 | 0.0060 |  |
|  | V3V4-one step |  | 0.8944 | 0.0004 | 0.0010 |  |
|  | V3V4-two step |  | 0.7071 | 0.0048 | 0.0112 |  |
| *Francisella* | V1V2-one step | *N.vectensis* | 1.0000 | 0.0002 | 0.0007 |  |
|  | V1V2-two step |  | 0.9997 | 0.0001 | 0.0005 |  |
|  | V3V4-one step |  | 0.9991 | 0.0001 | 0.0004 |  |
| *Fusicatenibacter* | V1V2-one step | *H.sapiens* | 0.8478 | 0.0009 | 0.0024 |  |
|  | V1V2-two step |  | 0.8738 | 0.0002 | 0.0008 |  |
|  | V3V4-one step |  | 0.8944 | 0.0001 | 0.0004 |  |
|  | V3V4-two step |  | 1.0000 | 0.0001 | 0.0005 |  |
| *Fusobacterium* | V1V2-one step | *A.aurita* | 0.6450 | 0.0072 | 0.0141 |  |
| *Gaiella* | V3V4-one step | *D.mel.(gut)* | 0.7071 | 0.0047 | 0.0089 |  |
| *Gammaproteobacteria* | V1V2-one step | *A.aerophoba* | 0.9435 | 0.0001 | 0.0006 |  |
|  | V1V2-two step |  | 0.9620 | 0.0001 | 0.0005 |  |
|  | V3V4-one step |  | 0.9310 | 0.0001 | 0.0004 |  |
|  | V3V4-two step |  | 0.6477 | 0.0198 | 0.0359 |  |
| *Gemella* | V1V2-one step | *M.leidyi* | 0.6617 | 0.0079 | 0.0152 |  |
|  | V1V2-two step | *M.musculus* | 0.5923 | 0.0151 | 0.0294 |  |
| *Gemmiger* | V3V4-one step | *H.sapiens* | 1.0000 | 0.0001 | 0.0004 |  |
|  | V3V4-two step |  | 1.0000 | 0.0001 | 0.0005 |  |
| *Gemmobacter* | V3V4-one step | *H.vulgaris* | 0.9309 | 0.0001 | 0.0004 |  |
| *Geobacillus* | V1V2-two step | *D.mel.(gut)* | 0.7071 | 0.0071 | 0.0144 |  |
| *Gp10* | V3V4-one step | *A.aerophoba* | 0.8944 | 0.0003 | 0.0008 |  |
| *Gp11* | V1V2-one step | *A.aerophoba* | 0.8944 | 0.0002 | 0.0007 |  |
|  | V1V2-two step |  | 0.8944 | 0.0002 | 0.0008 |  |
|  | V3V4-one step |  | 0.8944 | 0.0004 | 0.0010 |  |
| *Gp21* | V3V4-one step | *A.aerophoba* | 1.0000 | 0.0002 | 0.0006 |  |
|  | V3V4-two step |  | 0.7746 | 0.0024 | 0.0064 |  |
| *Gp3* | V1V2-one step | *A.aerophoba* | 1.0000 | 0.0001 | 0.0006 |  |
|  | V1V2-two step |  | 1.0000 | 0.0001 | 0.0005 |  |
|  | V3V4-one step |  | 0.8944 | 0.0003 | 0.0008 |  |
|  | V3V4-two step |  | 0.6808 | 0.0090 | 0.0180 |  |
| *Gp5* | V1V2-one step | *A.aerophoba* | 1.0000 | 0.0001 | 0.0006 |  |
|  | V1V2-two step |  | 0.8944 | 0.0002 | 0.0008 |  |
|  | V3V4-one step |  | 1.0000 | 0.0002 | 0.0006 |  |
| *Gp6* | V1V2-one step | *A.aerophoba* | 0.9425 | 0.0001 | 0.0006 |  |
|  | V1V2-two step |  | 0.9653 | 0.0001 | 0.0005 |  |
|  | V3V4-one step |  | 0.9914 | 0.0002 | 0.0006 |  |
|  | V3V4-two step |  | 0.7708 | 0.0034 | 0.0084 |  |
| *Gp9* | V1V2-one step | *A.aerophoba* | 0.9994 | 0.0001 | 0.0006 |  |
|  | V1V2-two step |  | 1.0000 | 0.0001 | 0.0005 |  |
|  | V3V4-one step |  | 1.0000 | 0.0002 | 0.0006 |  |
|  | V3V4-two step |  | 0.7746 | 0.0024 | 0.0064 |  |
| *Haemophilus* | V1V2-two step | *H.sapiens* | 0.6969 | 0.0057 | 0.0125 |  |
|  | V3V4-two step |  | 0.6252 | 0.0203 | 0.0366 |  |
| *Halioglobus* | V3V4-one step | *M.leidyi* | 0.9346 | 0.0001 | 0.0004 |  |
|  | V3V4-two step |  | 0.7422 | 0.0021 | 0.0060 |  |
| *Halobacteriovorax* | V3V4-one step | *N.vectensis* | 0.9476 | 0.0001 | 0.0004 |  |
| *Halomonadaceae* | V1V2-two step | *D.mel.(feces)* | 0.7766 | 0.0143 | 0.0280 |  |
| *Halomonas* | V1V2-two step | *D.mel.(feces)* | 0.7171 | 0.0173 | 0.0330 |  |
|  | V3V4-one step | *N.vectensis* | 0.7746 | 0.0038 | 0.0075 |  |
|  | V3V4-two step | *M.leidyi* | 0.7977 | 0.0001 | 0.0005 |  |
| *Hathewaya* | V1V2-one step | *A.aurita* | 1.0000 | 0.0001 | 0.0006 |  |
|  | V1V2-two step |  | 0.9995 | 0.0001 | 0.0005 |  |
|  | V3V4-one step |  | 1.0000 | 0.0001 | 0.0004 |  |
|  | V3V4-two step |  | 1.0000 | 0.0001 | 0.0005 |  |
| *Helicobacter* | V1V2-one step | *M.musculus* | 0.9996 | 0.0001 | 0.0006 |  |
|  | V1V2-two step |  | 0.9995 | 0.0001 | 0.0005 |  |
|  | V3V4-one step |  | 0.9995 | 0.0001 | 0.0004 |  |
|  | V3V4-two step |  | 1.0000 | 0.0001 | 0.0005 |  |
| *Hoeflea* | V3V4-one step | *N.vectensis* | 0.9652 | 0.0001 | 0.0004 |  |
| *Holdemania* | V1V2-one step | *H.sapiens* | 1.0000 | 0.0001 | 0.0006 |  |
|  | V1V2-two step |  | 1.0000 | 0.0001 | 0.0005 |  |
|  | V3V4-one step |  | 0.7746 | 0.0040 | 0.0077 |  |
|  | V3V4-two step |  | 0.9129 | 0.0001 | 0.0005 |  |
| *Howardella* | V1V2-two step | *H.sapiens* | 0.7071 | 0.0048 | 0.0110 |  |
|  | V3V4-two step |  | 0.7071 | 0.0060 | 0.0131 |  |
| *Hungatella* | V1V2-two step | *H.sapiens* | 0.8165 | 0.0009 | 0.0030 |  |
| *Hydrogenophaga* | V1V2-one step | *H.vulgaris* | 0.9877 | 0.0001 | 0.0006 |  |
| *Hymenobacter* | V1V2-one step | *M.leidyi* | 0.9451 | 0.0001 | 0.0006 |  |
| *Hyphomicrobium* | V3V4-one step | *A.aurita* | 0.7322 | 0.0073 | 0.0131 |  |
|  | V3V4-two step |  | 0.6810 | 0.0079 | 0.0164 |  |
| *Hyphomonadaceae* | V1V2-one step | *N.vectensis* | 0.9113 | 0.0002 | 0.0007 |  |
|  | V1V2-two step |  | 0.9946 | 0.0001 | 0.0005 |  |
|  | V3V4-one step |  | 1.0000 | 0.0001 | 0.0004 |  |
| *Hyphomonas* | V3V4-one step | *M.leidyi* | 0.7229 | 0.0061 | 0.0111 |  |
| *Iamiaceae* | V1V2-one step | *A.aurita* | 1.0000 | 0.0001 | 0.0006 |  |
|  | V1V2-two step | *A.aurita* | 0.9097 | 0.0001 | 0.0005 |  |
|  | V3V4-one step | *A.aerophoba* | 0.9992 | 0.0002 | 0.0006 |  |
|  | V3V4-two step | *A.aerophoba* | 0.7633 | 0.0027 | 0.0070 |  |
| *Idiomarina* | V1V2-one step | *N.vectensis* | 0.7746 | 0.0040 | 0.0088 |  |
| *Ilumatobacter* | V1V2-one step | *A.aurita* | 0.7055 | 0.0018 | 0.0044 |  |
|  | V1V2-two step |  | 0.5840 | 0.0173 | 0.0330 |  |
|  | V3V4-one step |  | 0.9094 | 0.0003 | 0.0008 |  |
|  | V3V4-two step |  | 0.7980 | 0.0011 | 0.0037 |  |
| *Intestinimonas* | V1V2-one step | *M.musculus* | 0.9951 | 0.0001 | 0.0006 |  |
|  | V1V2-two step |  | 0.9886 | 0.0001 | 0.0005 |  |
|  | V3V4-one step |  | 0.7303 | 0.0008 | 0.0019 |  |
|  | V3V4-two step |  | 0.7504 | 0.0006 | 0.0023 |  |
| *Jannaschia* | V3V4-one step | *N.vectensis* | 0.8460 | 0.0006 | 0.0015 |  |
| *Kangiella* | V1V2-one step | *N.vectensis* | 0.8944 | 0.0002 | 0.0007 |  |
| *Kiloniella* | V1V2-one step | *N.vectensis* | 0.9881 | 0.0002 | 0.0007 |  |
|  | V1V2-two step |  | 0.9965 | 0.0001 | 0.0005 |  |
|  | V3V4-one step |  | 0.9998 | 0.0001 | 0.0004 |  |
| *Kocuria* | V3V4-one step | *D.mel.(gut)* | 0.6268 | 0.0068 | 0.0123 |  |
| *Lachnospiracea incertae sedis* | V1V2-one step | *H.sapiens* | 0.9947 | 0.0001 | 0.0006 |  |
|  | V1V2-two step |  | 0.9783 | 0.0001 | 0.0005 |  |
|  | V3V4-one step |  | 0.9746 | 0.0001 | 0.0004 |  |
|  | V3V4-two step |  | 1.0000 | 0.0001 | 0.0005 |  |
| *Lachnospiraceae* | V1V2-one step | *M.musculus* | 0.9281 | 0.0001 | 0.0006 |  |
|  | V1V2-two step |  | 0.8917 | 0.0001 | 0.0005 |  |
|  | V3V4-one step |  | 0.9379 | 0.0001 | 0.0004 |  |
|  | V3V4-two step |  | 0.8755 | 0.0001 | 0.0005 |  |
| *Lactobacillus* | V1V2-one step | *D.mel.(feces)* | 0.7030 | 0.0006 | 0.0016 | MEGAN |
|  | V1V2-two step | *D.mel.(gut)* | 0.6764 | 0.0135 | 0.0266 | Kraken |
|  | V3V4-one step | *D.mel.(feces)* | 0.7233 | 0.0009 | 0.0021 | MetaPhlan |
|  | V3V4-two step | *D.mel.(feces)* | 0.8926 | 0.0001 | 0.0005 | MetaPhlan2 |
| *Lactococcus* | V3V4-two step | *C.elegans* | 0.7066 | 0.0082 | 0.0168 |  |
| *Legionella* | V1V2-one step | *H.vulgaris* | 0.9717 | 0.0001 | 0.0006 |  |
|  | V1V2-two step |  | 0.8401 | 0.0065 | 0.0138 |  |
|  | V3V4-one step |  | 0.9886 | 0.0001 | 0.0004 |  |
|  | V3V4-two step |  | 0.8309 | 0.0067 | 0.0142 |  |
| *Leisingera* | V3V4-one step | *M.leidyi* | 0.8944 | 0.0001 | 0.0004 |  |
| *Leptonema* | V3V4-one step | *N.vectensis* | 0.7746 | 0.0046 | 0.0087 |  |
| *Lewinella* | V3V4-one step | *N.vectensis* | 0.9359 | 0.0001 | 0.0004 |  |
| *Litoreibacter* | V3V4-one step | *A.aurita* | 0.9732 | 0.0001 | 0.0004 |  |
|  | V3V4-two step |  | 0.8796 | 0.0006 | 0.0023 |  |
| *Litorilinea* | V3V4-one step | *A.aerophoba* | 0.7845 | 0.0011 | 0.0025 |  |
|  | V3V4-two step | *A.aurita* | 0.6594 | 0.0067 | 0.0142 |  |
| *Loktanella* | V3V4-one step | *N.vectensis* | 0.8716 | 0.0001 | 0.0004 |  |
| *Luteolibacter* | V3V4-one step | *M.leidyi* | 0.6595 | 0.0276 | 0.0450 |  |
| *Malikia* | V1V2-one step | *H.vulgaris* | 0.6708 | 0.0154 | 0.0287 |  |
| *Mariniflexile* | V1V2-one step | *N.vectensis* | 1.0000 | 0.0002 | 0.0007 |  |
|  | V1V2-two step |  | 0.9833 | 0.0001 | 0.0005 |  |
|  | V3V4-one step |  | 1.0000 | 0.0001 | 0.0004 |  |
| *Marinobacter* | V1V2-one step | *N.vectensis* | 0.9946 | 0.0002 | 0.0007 |  |
|  | V1V2-two step | *N.vectensis* | 0.6382 | 0.0090 | 0.0179 |  |
|  | V3V4-one step | *M.leidyi* | 0.8773 | 0.0001 | 0.0004 |  |
|  | V3V4-two step | *M.leidyi* | 0.9967 | 0.0001 | 0.0005 |  |
| *Marinobacterium* | V3V4-one step | *M.leidyi* | 0.8944 | 0.0001 | 0.0004 |  |
|  | V3V4-two step |  | 1.0000 | 0.0001 | 0.0005 |  |
| *Marinomonas* | V1V2-one step | *M.leidyi* | 0.8241 | 0.0006 | 0.0016 |  |
|  | V1V2-two step |  | 0.5891 | 0.0189 | 0.0356 |  |
|  | V3V4-two step |  | 0.7740 | 0.0019 | 0.0055 |  |
| *Maritalea* | V1V2-one step | *N.vectensis* | 0.9453 | 0.0002 | 0.0007 |  |
|  | V1V2-two step |  | 1.0000 | 0.0001 | 0.0005 |  |
|  | V3V4-one step |  | 1.0000 | 0.0001 | 0.0004 |  |
| *Marivita* | V3V4-one step | *M.leidyi* | 0.8463 | 0.0001 | 0.0004 |  |
|  | V3V4-two step |  | 0.9976 | 0.0001 | 0.0005 |  |
| *Marixanthomonas* | V3V4-one step | *N.vectensis* | 0.7746 | 0.0038 | 0.0075 |  |
| *Mesonia* | V1V2-one step | *N.vectensis* | 1.0000 | 0.0002 | 0.0007 |  |
|  | V1V2-two step |  | 0.7746 | 0.0015 | 0.0045 |  |
|  | V3V4-one step |  | 1.0000 | 0.0001 | 0.0004 |  |
| *Methylococcaceae* | V1V2-two step | *N.vectensis* | 0.7746 | 0.0019 | 0.0052 |  |
| *Methylocystis* | V3V4-one step | *A.aurita* | 1.0000 | 0.0001 | 0.0004 |  |
|  | V3V4-two step |  | 0.8165 | 0.0005 | 0.0020 |  |
| *Methyloparacoccus* | V1V2-one step | *A.aurita* | 0.8944 | 0.0004 | 0.0012 |  |
|  | V1V2-two step |  | 0.9129 | 0.0001 | 0.0005 |  |
|  | V3V4-one step |  | 0.8944 | 0.0002 | 0.0006 |  |
|  | V3V4-two step |  | 0.9129 | 0.0002 | 0.0009 |  |
| *Methylophaga* | V1V2-one step | *N.vectensis* | 1.0000 | 0.0002 | 0.0007 |  |
|  | V1V2-two step |  | 0.8778 | 0.0001 | 0.0005 |  |
| *Methylophilaceae* | V1V2-one step | *N.vectensis* | 0.9909 | 0.0002 | 0.0007 |  |
|  | V1V2-two step | *N.vectensis* | 0.8924 | 0.0001 | 0.0005 |  |
|  | V3V4-one step | *M.leidyi* | 0.9930 | 0.0001 | 0.0004 |  |
|  | V3V4-two step | *M.leidyi* | 0.9937 | 0.0001 | 0.0005 |  |
| *Methylophilus* | V1V2-one step | *H.vulgaris* | 0.7746 | 0.0047 | 0.0098 |  |
|  | V1V2-two step |  | 0.7071 | 0.0068 | 0.0139 |  |
|  | V3V4-one step |  | 0.9877 | 0.0001 | 0.0004 |  |
| *Methylotenera* | V3V4-one step | *N.vectensis* | 1.0000 | 0.0001 | 0.0004 |  |
| *Microbacteriaceae* | V1V2-one step | *A.aurita* | 0.8276 | 0.0016 | 0.0040 |  |
|  | V1V2-two step |  | 0.9276 | 0.0001 | 0.0005 |  |
|  | V3V4-one step |  | 0.8520 | 0.0001 | 0.0004 |  |
|  | V3V4-two step |  | 0.7095 | 0.0014 | 0.0042 |  |
| *Micrococcaceae* | V3V4-two step | *C.elegans* | 0.7071 | 0.0048 | 0.0112 |  |
| *Micrococcus* | V1V2-one step | *M.leidyi* | 0.5861 | 0.0187 | 0.0339 |  |
|  | V3V4-one step | *D.mel.(gut)* | 0.7097 | 0.0032 | 0.0066 |  |
| *Moritella* | V1V2-two step | *A.aurita* | 0.8165 | 0.0012 | 0.0039 |  |
|  | V3V4-one step |  | 0.7721 | 0.0083 | 0.0147 |  |
|  | V3V4-two step |  | 0.7071 | 0.0055 | 0.0123 |  |
| *Mucispirillum* | V1V2-one step | *M.musculus* | 0.8941 | 0.0003 | 0.0010 | MetaPhlan2 |
|  | V1V2-two step |  | 0.9129 | 0.0001 | 0.0005 |  |
|  | V3V4-one step |  | 0.8940 | 0.0002 | 0.0006 |  |
|  | V3V4-two step |  | 0.9129 | 0.0001 | 0.0005 |  |
| *Mycobacterium* | V1V2-one step | *A.aurita* | 0.8797 | 0.0002 | 0.0007 |  |
|  | V1V2-two step |  | 0.6354 | 0.0261 | 0.0478 |  |
|  | V3V4-one step |  | 0.7646 | 0.0012 | 0.0027 |  |
|  | V3V4-two step |  | 0.8023 | 0.0011 | 0.0037 |  |
| *Mycoplasma* | V3V4-one step | *M.leidyi* | 0.8338 | 0.0008 | 0.0019 |  |
|  | V3V4-two step |  | 0.8940 | 0.0002 | 0.0009 |  |
| *Myxococcales* | V3V4-one step | *N.vectensis* | 0.6401 | 0.0136 | 0.0233 |  |
|  | V3V4-two step | *D.mel.(feces)* | 0.7559 | 0.0025 | 0.0066 |  |
| *Neptunomonas* | V1V2-two step | *M.leidyi* | 0.7448 | 0.0014 | 0.0043 |  |
| *Nitratireductor* | V3V4-one step | *N.vectensis* | 0.6831 | 0.0039 | 0.0076 |  |
| *Nitrosomonas* | V1V2-one step | *N.vectensis* | 0.8944 | 0.0003 | 0.0010 |  |
|  | V1V2-two step |  | 1.0000 | 0.0001 | 0.0005 |  |
|  | V3V4-one step |  | 1.0000 | 0.0001 | 0.0004 |  |
| *Nitrospira* | V3V4-one step | *A.aerophoba* | 1.0000 | 0.0002 | 0.0006 |  |
| *Nocardioides* | V1V2-one step | *M.leidyi* | 0.7895 | 0.0001 | 0.0006 |  |
| *Oceanicaulis* | V3V4-one step | *M.leidyi* | 0.8944 | 0.0002 | 0.0006 |  |
|  | V3V4-two step |  | 0.7746 | 0.0014 | 0.0042 |  |
| *Oceanisphaera* | V1V2-one step | *A.aurita* | 0.9677 | 0.0001 | 0.0006 |  |
|  | V1V2-two step |  | 0.9614 | 0.0001 | 0.0005 |  |
|  | V3V4-one step |  | 0.7746 | 0.0061 | 0.0111 |  |
|  | V3V4-two step |  | 0.9129 | 0.0002 | 0.0009 |  |
| *Oceanospirillaceae* | V1V2-one step | *N.vectensis* | 1.0000 | 0.0002 | 0.0007 |  |
|  | V1V2-two step |  | 0.8869 | 0.0001 | 0.0005 |  |
| *Oceanospirillales* | V3V4-one step | *A.aerophoba* | 0.7746 | 0.0037 | 0.0074 |  |
|  | V3V4-two step |  | 0.6841 | 0.0061 | 0.0133 |  |
| *Odoribacter* | V1V2-one step | *H.sapiens* | 0.7298 | 0.0015 | 0.0038 |  |
|  | V1V2-two step | *H.sapiens* | 0.7087 | 0.0026 | 0.0067 |  |
|  | V3V4-one step | *M.musculus* | 0.9124 | 0.0001 | 0.0004 |  |
|  | V3V4-two step | *M.musculus* | 0.8591 | 0.0001 | 0.0005 |  |
| *Opitutae* | V3V4-one step | *A.aerophoba* | 0.9630 | 0.0002 | 0.0006 |  |
|  | V3V4-two step |  | 0.7368 | 0.0029 | 0.0075 |  |
| *Opitutus* | V3V4-one step | *N.vectensis* | 0.7746 | 0.0036 | 0.0073 |  |
| *Oscillibacter* | V1V2-one step | *H.sapiens* | 0.7453 | 0.0010 | 0.0026 |  |
|  | V1V2-two step | *H.sapiens* | 0.8226 | 0.0001 | 0.0005 |  |
|  | V3V4-one step | *M.musculus* | 0.7211 | 0.0008 | 0.0019 |  |
|  | V3V4-two step | *H.sapiens* | 0.7094 | 0.0009 | 0.0032 |  |
| *Oxalobacteraceae* | V1V2-one step | *H.vulgaris* | 0.9958 | 0.0001 | 0.0006 |  |
|  | V1V2-two step |  | 0.6716 | 0.0188 | 0.0356 |  |
| *Paenibacillus* | V1V2-two step | *T.aestivum* | 0.7071 | 0.0045 | 0.0107 |  |
| *Panacagrimonas* | V1V2-one step | *N.vectensis* | 0.7746 | 0.0042 | 0.0091 |  |
| *Pantoea* | V1V2-one step | *T.aestivum* | 0.9599 | 0.0002 | 0.0007 | MEGAN |
|  | V1V2-two step |  | 0.9978 | 0.0001 | 0.0005 |  |
| *Parabacteroides* | V1V2-one step | *H.sapiens* | 0.9255 | 0.0004 | 0.0012 |  |
|  | V1V2-two step |  | 0.8943 | 0.0002 | 0.0008 |  |
|  | V3V4-one step |  | 1.0000 | 0.0001 | 0.0004 |  |
|  | V3V4-two step |  | 1.0000 | 0.0001 | 0.0005 |  |
| *Parachlamydiaceae* | V3V4-one step | *N.vectensis* | 0.9899 | 0.0001 | 0.0004 |  |
| *Paracoccus* | V3V4-two step | *D.mel.(feces)* | 0.9345 | 0.0001 | 0.0005 |  |
| *Parasegetibacter* | V3V4-one step | *A.aurita* | 0.8022 | 0.0006 | 0.0015 |  |
|  | V3V4-two step |  | 0.8165 | 0.0009 | 0.0032 |  |
| *Parasutterella* | V1V2-one step | *H.sapiens* | 0.8767 | 0.0006 | 0.0016 |  |
|  | V1V2-two step |  | 0.9104 | 0.0002 | 0.0008 |  |
|  | V3V4-one step |  | 0.7668 | 0.0038 | 0.0075 |  |
|  | V3V4-two step |  | 0.8123 | 0.0059 | 0.0131 |  |
| *Parcubacteria genera incertae sedis* | V3V4-one step | *A.aurita* | 0.8701 | 0.0003 | 0.0008 |  |
|  | V3V4-two step |  | 0.8165 | 0.0009 | 0.0032 |  |
| *Pedobacter* | V1V2-two step | *H.vulgaris* | 0.7071 | 0.0068 | 0.0139 |  |
|  | V3V4-one step |  | 0.9754 | 0.0001 | 0.0004 |  |
|  | V3V4-two step |  | 0.7071 | 0.0100 | 0.0189 |  |
| *Peptoniphilaceae* | V3V4-one step | *D.mel.(gut)* | 0.7071 | 0.0042 | 0.0081 |  |
| *Peredibacter* | V1V2-two step | *A.aurita* | 0.7071 | 0.0046 | 0.0109 |  |
|  | V3V4-one step | *M.leidyi* | 0.7435 | 0.0015 | 0.0033 |  |
|  | V3V4-two step | *M.leidyi* | 0.6325 | 0.0292 | 0.0494 |  |
| *Phaeodactylibacter* | V1V2-one step | *N.vectensis* | 0.8992 | 0.0002 | 0.0007 |  |
|  | V1V2-two step |  | 0.8062 | 0.0006 | 0.0021 |  |
| *Phenylobacterium* | V3V4-one step | *D.mel.(feces)* | 0.9620 | 0.0001 | 0.0004 |  |
|  | V3V4-two step |  | 0.9991 | 0.0001 | 0.0005 |  |
| *Photobacterium* | V3V4-one step | *A.aurita* | 0.8687 | 0.0005 | 0.0012 |  |
|  | V3V4-two step |  | 0.9129 | 0.0003 | 0.0013 |  |
| *Phycisphaera* | V1V2-one step | *N.vectensis* | 1.0000 | 0.0002 | 0.0007 |  |
|  | V1V2-two step |  | 0.8815 | 0.0001 | 0.0005 |  |
|  | V3V4-one step |  | 0.8944 | 0.0003 | 0.0008 |  |
| *Planctomycetaceae* | V1V2-one step | *A.aurita* | 0.9986 | 0.0001 | 0.0006 |  |
|  | V1V2-two step |  | 1.0000 | 0.0001 | 0.0005 |  |
|  | V3V4-one step |  | 0.9877 | 0.0001 | 0.0004 |  |
|  | V3V4-two step |  | 0.8920 | 0.0002 | 0.0009 |  |
| *Planktomarina* | V3V4-one step | *M.leidyi* | 0.9764 | 0.0001 | 0.0004 |  |
|  | V3V4-two step |  | 0.9639 | 0.0001 | 0.0005 |  |
| *Polaribacter* | V1V2-one step | *M.leidyi* | 0.9081 | 0.0003 | 0.0010 |  |
|  | V1V2-two step | *M.leidyi* | 0.8580 | 0.0006 | 0.0021 |  |
|  | V3V4-one step | *N.vectensis* | 0.9517 | 0.0001 | 0.0004 |  |
| *Poribacteria genera incertae sedis* | V1V2-one step | *A.aerophoba* | 0.9999 | 0.0001 | 0.0006 |  |
|  | V1V2-two step |  | 0.9776 | 0.0001 | 0.0005 |  |
|  | V3V4-one step |  | 1.0000 | 0.0002 | 0.0006 |  |
|  | V3V4-two step |  | 0.7746 | 0.0024 | 0.0064 |  |
| *Porphyromonadaceae* | V1V2-two step | *M.musculus* | 0.5847 | 0.0162 | 0.0313 |  |
|  | V3V4-one step |  | 0.9183 | 0.0001 | 0.0004 |  |
|  | V3V4-two step |  | 0.8559 | 0.0001 | 0.0005 |  |
| *Porticoccus* | V3V4-one step | *N.vectensis* | 0.7746 | 0.0048 | 0.0090 |  |
| *Prevotellaceae* | V1V2-one step | *M.musculus* | 0.8466 | 0.0022 | 0.0052 |  |
|  | V1V2-two step |  | 0.8850 | 0.0013 | 0.0041 |  |
| *Propionibacterium* | V1V2-two step | *T.aestivum* | 0.5358 | 0.0266 | 0.0484 |  |
|  | V3V4-one step | *D.mel.(gut)* | 0.8131 | 0.0095 | 0.0166 |  |
|  | V3V4-two step | *C.elegans* | 0.6711 | 0.0173 | 0.0318 |  |
| *Proteobacteria* | V1V2-one step | *A.aerophoba* | 0.6793 | 0.0005 | 0.0014 |  |
|  | V1V2-two step | *A.aerophoba* | 0.6874 | 0.0002 | 0.0008 |  |
|  | V3V4-one step | *M.leidyi* | 0.9580 | 0.0001 | 0.0004 |  |
|  | V3V4-two step | *M.leidyi* | 0.9725 | 0.0001 | 0.0005 |  |
| *Pseudoalteromonas* | V1V2-one step | *N.vectensis* | 0.7427 | 0.0002 | 0.0007 |  |
|  | V1V2-two step | *N.vectensis* | 0.7719 | 0.0002 | 0.0008 |  |
|  | V3V4-one step | *N.vectensis* | 0.6899 | 0.0005 | 0.0012 |  |
|  | V3V4-two step | *M.leidyi* | 0.7213 | 0.0002 | 0.0009 |  |
| *Pseudoflavonifractor* | V1V2-one step | *H.sapiens* | 1.0000 | 0.0001 | 0.0006 |  |
|  | V1V2-two step |  | 1.0000 | 0.0001 | 0.0005 |  |
| *Pseudomonadaceae* | V1V2-two step | *H.vulgaris* | 0.5925 | 0.0260 | 0.0478 |  |
| *Pseudomonas* | V1V2-one step | *C.elegans* | 0.8599 | 0.0129 | 0.0243 | MEGAN,MetaPhlan |
|  | V1V2-two step |  | 0.8280 | 0.0018 | 0.0052 | MEGAN,MetaPhlan |
|  | V3V4-one step |  | 0.9174 | 0.0049 | 0.0091 | MEGAN,MetaPhlan |
|  | V3V4-two step |  | 0.9865 | 0.0010 | 0.0034 | MEGAN,MetaPhlan |
| *Psychrobacter* | V1V2-one step | *N.vectensis* | 0.8481 | 0.0004 | 0.0012 |  |
|  | V1V2-two step |  | 0.7267 | 0.0047 | 0.0109 |  |
|  | V3V4-one step |  | 0.8494 | 0.0006 | 0.0015 |  |
| *Psychromonas* | V1V2-one step | *A.aurita* | 0.7746 | 0.0050 | 0.0101 |  |
|  | V1V2-two step |  | 0.7071 | 0.0050 | 0.0113 |  |
| *Ralstonia* | V1V2-one step | *D.mel.(gut)* | 0.6091 | 0.0066 | 0.0130 |  |
|  | V3V4-one step | *M.leidyi* | 0.7138 | 0.0001 | 0.0004 |  |
|  | V3V4-two step | *M.leidyi* | 0.8134 | 0.0002 | 0.0009 |  |
| *Reyranella* | V1V2-one step | *H.vulgaris* | 0.8193 | 0.0015 | 0.0038 |  |
|  | V3V4-one step |  | 0.8790 | 0.0007 | 0.0017 |  |
| *Rheinheimera* | V1V2-one step | *A.aurita* | 0.7006 | 0.0101 | 0.0194 |  |
|  | V1V2-two step |  | 0.8676 | 0.0002 | 0.0008 |  |
|  | V3V4-two step |  | 0.7827 | 0.0024 | 0.0064 |  |
| *Rhizobiaceae* | V3V4-one step | *H.vulgaris* | 1.0000 | 0.0001 | 0.0004 |  |
|  | V3V4-two step |  | 0.7071 | 0.0100 | 0.0189 |  |
| *Rhizobiales* | V1V2-two step | *A.aurita* | 0.7188 | 0.0019 | 0.0052 |  |
| *Rhodanobacter* | V3V4-one step | *D.mel.(feces)* | 1.0000 | 0.0001 | 0.0004 |  |
|  | V3V4-two step |  | 0.8944 | 0.0001 | 0.0005 |  |
| *Rhodobacteraceae* | V1V2-one step | *A.aerophoba* | 0.9659 | 0.0008 | 0.0022 |  |
|  | V1V2-two step | *A.aerophoba* | 0.9379 | 0.0007 | 0.0024 |  |
|  | V3V4-one step | *H.vulgaris* | 0.7423 | 0.0001 | 0.0004 |  |
| *Rhodococcus* | V1V2-one step | *D.mel.(gut)* | 0.9201 | 0.0001 | 0.0006 |  |
|  | V1V2-two step |  | 0.8595 | 0.0006 | 0.0021 |  |
|  | V3V4-one step |  | 0.9634 | 0.0001 | 0.0004 |  |
|  | V3V4-two step |  | 0.6751 | 0.0188 | 0.0344 |  |
| *Rhodoferax* | V1V2-one step | *H.vulgaris* | 0.9934 | 0.0001 | 0.0006 |  |
|  | V1V2-two step |  | 0.8544 | 0.0008 | 0.0027 |  |
|  | V3V4-one step |  | 0.9946 | 0.0001 | 0.0004 |  |
| *Rhodoluna* | V1V2-one step | *A.aurita* | 0.8944 | 0.0004 | 0.0012 |  |
|  | V1V2-two step |  | 0.8067 | 0.0004 | 0.0015 |  |
| *Rhodospirillaceae* | V3V4-one step | *A.aerophoba* | 0.8787 | 0.0001 | 0.0004 |  |
| *Rhodospirillales* | V3V4-one step | *A.aerophoba* | 0.9987 | 0.0002 | 0.0006 |  |
|  | V3V4-two step |  | 0.7571 | 0.0030 | 0.0075 |  |
| *Rhodothermaceae* | V3V4-one step | *A.aerophoba* | 1.0000 | 0.0002 | 0.0006 |  |
|  | V3V4-two step |  | 0.7746 | 0.0024 | 0.0064 |  |
| *Rickettsiaceae* | V1V2-one step | *A.aurita* | 0.9656 | 0.0001 | 0.0006 |  |
|  | V1V2-two step |  | 0.9825 | 0.0001 | 0.0005 |  |
| *Robiginitalea* | V3V4-two step | *M.leidyi* | 0.7746 | 0.0014 | 0.0042 |  |
| *Romboutsia* | V1V2-one step | *H.sapiens* | 0.7746 | 0.0043 | 0.0091 |  |
|  | V3V4-two step |  | 0.6455 | 0.0073 | 0.0154 |  |
| *Roseburia* | V1V2-one step | *H.sapiens* | 0.8944 | 0.0003 | 0.0010 |  |
|  | V1V2-two step |  | 0.9708 | 0.0001 | 0.0005 |  |
|  | V3V4-one step |  | 1.0000 | 0.0001 | 0.0004 |  |
|  | V3V4-two step |  | 1.0000 | 0.0001 | 0.0005 |  |
| *Roseibacillus* | V3V4-one step | *A.aurita* | 0.6268 | 0.0144 | 0.0244 |  |
|  | V3V4-two step | *M.leidyi* | 0.9625 | 0.0001 | 0.0005 |  |
| *Roseovarius* | V3V4-one step | *N.vectensis* | 0.9332 | 0.0001 | 0.0004 |  |
| *Rothia* | V1V2-one step | *M.leidyi* | 0.6393 | 0.0230 | 0.0414 |  |
|  | V3V4-one step |  | 0.7441 | 0.0292 | 0.0472 |  |
|  | V3V4-two step |  | 0.6721 | 0.0100 | 0.0189 |  |
| *Ruegeria* | V3V4-two step | *M.leidyi* | 0.7348 | 0.0084 | 0.0170 |  |
| *Ruminococcaceae* | V1V2-one step | *H.sapiens* | 0.8435 | 0.0001 | 0.0006 |  |
|  | V1V2-two step |  | 0.9019 | 0.0001 | 0.0005 |  |
|  | V3V4-one step |  | 0.8658 | 0.0003 | 0.0008 |  |
|  | V3V4-two step |  | 0.8831 | 0.0001 | 0.0005 |  |
| *Ruminococcus* | V1V2-one step | *H.sapiens* | 0.6325 | 0.0158 | 0.0293 |  |
|  | V3V4-two step |  | 0.6799 | 0.0240 | 0.0419 |  |
| *Ruminococcus2* | V1V2-one step | *H.sapiens* | 0.8944 | 0.0003 | 0.0010 |  |
|  | V1V2-two step |  | 0.9129 | 0.0001 | 0.0005 |  |
|  | V3V4-one step |  | 0.9783 | 0.0001 | 0.0004 |  |
|  | V3V4-two step |  | 0.9010 | 0.0001 | 0.0005 |  |
| *Runella* | V1V2-one step | *H.vulgaris* | 0.8944 | 0.0003 | 0.0010 |  |
| *Saccharibacteria genera incertae sedis* | V1V2-one step | *A.aurita* | 0.7035 | 0.0048 | 0.0099 |  |
|  | V1V2-two step |  | 0.8326 | 0.0012 | 0.0039 |  |
|  | V3V4-one step |  | 0.6546 | 0.0073 | 0.0131 |  |
|  | V3V4-two step |  | 0.8751 | 0.0001 | 0.0005 |  |
| *Salinirepens* | V1V2-one step | *A.aurita* | 0.7746 | 0.0028 | 0.0066 |  |
|  | V1V2-two step | *M.leidyi* | 0.7362 | 0.0003 | 0.0012 |  |
|  | V3V4-one step | *M.leidyi* | 0.9858 | 0.0001 | 0.0004 |  |
|  | V3V4-two step | *M.leidyi* | 0.9931 | 0.0001 | 0.0005 |  |
| *Salinisphaera* | V3V4-one step | *N.vectensis* | 0.7746 | 0.0036 | 0.0073 |  |
| *Salmonella* | V3V4-one step | *M.leidyi* | 0.5674 | 0.0278 | 0.0451 |  |
| *Saprospiraceae* | V1V2-one step | *M.leidyi* | 0.6411 | 0.0122 | 0.0233 |  |
|  | V3V4-one step |  | 0.7796 | 0.0007 | 0.0017 |  |
|  | V3V4-two step |  | 0.8368 | 0.0004 | 0.0017 |  |
| *Sediminibacterium* | V3V4-one step | *D.mel.(feces)* | 0.6882 | 0.0244 | 0.0401 |  |
| *Shewanella* | V1V2-one step | *A.aurita* | 0.9630 | 0.0001 | 0.0006 |  |
|  | V3V4-one step |  | 0.7627 | 0.0008 | 0.0019 |  |
|  | V3V4-two step |  | 0.6590 | 0.0123 | 0.0229 |  |
| *Shinella* | V3V4-one step | *H.vulgaris* | 0.9970 | 0.0001 | 0.0004 |  |
|  | V3V4-two step |  | 0.7071 | 0.0100 | 0.0189 |  |
| *Spartobacteria* | V1V2-two step | *A.aurita* | 0.7071 | 0.0043 | 0.0104 |  |
|  | V3V4-one step |  | 0.9042 | 0.0001 | 0.0004 |  |
|  | V3V4-two step |  | 0.8862 | 0.0003 | 0.0013 |  |
| *Spartobacteria genera incertae sedis* | V1V2-one step | *A.aurita* | 0.8944 | 0.0004 | 0.0012 |  |
|  | V1V2-two step |  | 1.0000 | 0.0001 | 0.0005 |  |
|  | V3V4-one step |  | 0.9996 | 0.0001 | 0.0004 |  |
|  | V3V4-two step |  | 0.9982 | 0.0001 | 0.0005 |  |
| *Sphaerotilus* | V1V2-one step | *H.vulgaris* | 0.9987 | 0.0001 | 0.0006 |  |
|  | V1V2-two step |  | 0.9695 | 0.0001 | 0.0005 |  |
|  | V3V4-one step |  | 1.0000 | 0.0001 | 0.0004 |  |
| *Sphingobacteriales* | V1V2-one step | *N.vectensis* | 1.0000 | 0.0002 | 0.0007 |  |
|  | V1V2-two step | *N.vectensis* | 0.8944 | 0.0001 | 0.0005 |  |
|  | V3V4-one step | *D.mel.(gut)* | 0.7071 | 0.0046 | 0.0087 |  |
| *Sphingomonadaceae* | V3V4-one step | *H.vulgaris* | 0.7734 | 0.0003 | 0.0008 |  |
| *Sphingomonas* | V3V4-one step | *M.leidyi* | 0.7955 | 0.0027 | 0.0057 |  |
| *Spirochaetaceae* | V1V2-one step | *N.vectensis* | 0.9929 | 0.0001 | 0.0006 |  |
|  | V1V2-two step |  | 0.9892 | 0.0001 | 0.0005 |  |
| *Spiroplasma* | V3V4-one step | *M.leidyi* | 0.9996 | 0.0001 | 0.0004 |  |
|  | V3V4-two step |  | 0.9999 | 0.0001 | 0.0005 |  |
| *Spirosoma* | V3V4-one step | *M.leidyi* | 0.7689 | 0.0030 | 0.0062 |  |
|  | V3V4-two step |  | 0.7746 | 0.0019 | 0.0055 |  |
| *Spongiibacter* | V1V2-two step | *N.vectensis* | 0.7746 | 0.0019 | 0.0052 |  |
| *Staphylococcus* | V1V2-one step | *A.aurita* | 0.7432 | 0.0040 | 0.0088 |  |
|  | V3V4-two step |  | 0.6958 | 0.0086 | 0.0173 |  |
| *Stenotrophomonas* | V3V4-two step | *D.mel.(feces)* | 0.5855 | 0.0268 | 0.0462 |  |
| *Streptococcus* | V1V2-two step | *M.leidyi* | 0.6867 | 0.0021 | 0.0057 |  |
|  | V3V4-two step |  | 0.6535 | 0.0211 | 0.0378 |  |
| *Sulfitobacter* | V3V4-one step | *N.vectensis* | 0.8296 | 0.0002 | 0.0006 |  |
|  | V3V4-two step | *M.leidyi* | 0.7709 | 0.0032 | 0.0080 |  |
| *Sulfurimonas* | V1V2-one step | *A.aurita* | 0.8944 | 0.0002 | 0.0007 |  |
|  | V1V2-two step |  | 1.0000 | 0.0001 | 0.0005 |  |
|  | V3V4-one step |  | 1.0000 | 0.0001 | 0.0004 |  |
|  | V3V4-two step |  | 0.9650 | 0.0002 | 0.0009 |  |
| *Sulfurovum* | V1V2-one step | *A.aurita* | 1.0000 | 0.0001 | 0.0006 |  |
|  | V1V2-two step |  | 1.0000 | 0.0001 | 0.0005 |  |
|  | V3V4-one step |  | 0.8702 | 0.0003 | 0.0008 |  |
|  | V3V4-two step |  | 0.7441 | 0.0015 | 0.0045 |  |
| *Sutterella* | V1V2-one step | *H.sapiens* | 0.7736 | 0.0071 | 0.0139 |  |
| *Thalassobaculum* | V3V4-one step | *A.aurita* | 0.8216 | 0.0003 | 0.0008 |  |
|  | V3V4-two step |  | 0.8165 | 0.0005 | 0.0020 |  |
| *Thalassolituus* | V3V4-one step | *N.vectensis* | 0.9855 | 0.0001 | 0.0004 |  |
| *Thalassotalea* | V1V2-one step | *N.vectensis* | 0.9984 | 0.0002 | 0.0007 |  |
|  | V1V2-two step |  | 0.8944 | 0.0001 | 0.0005 |  |
|  | V3V4-one step |  | 0.9956 | 0.0001 | 0.0004 |  |
| *Thiomicrospira* | V1V2-one step | *A.aurita* | 0.8944 | 0.0002 | 0.0007 |  |
| *Thiotrichales* | V1V2-one step | *M.leidyi* | 0.8944 | 0.0002 | 0.0007 |  |
|  | V1V2-two step |  | 1.0000 | 0.0001 | 0.0005 |  |
|  | V3V4-one step |  | 0.9973 | 0.0001 | 0.0004 |  |
|  | V3V4-two step |  | 1.0000 | 0.0001 | 0.0005 |  |
| *Turneriella* | V1V2-one step | *H.vulgaris* | 0.7701 | 0.0075 | 0.0146 |  |
|  | V1V2-two step |  | 0.7071 | 0.0068 | 0.0139 |  |
|  | V3V4-one step |  | 0.7746 | 0.0033 | 0.0068 |  |
|  | V3V4-two step |  | 0.7071 | 0.0100 | 0.0189 |  |
| *Undibacterium* | V3V4-one step | *H.vulgaris* | 0.9641 | 0.0001 | 0.0004 |  |
|  | V3V4-two step |  | 0.7071 | 0.0100 | 0.0189 |  |
| *Verrucomicrobiaceae* | V3V4-one step | *M.leidyi* | 0.7384 | 0.0052 | 0.0096 |  |
| *Vibrio* | V1V2-one step | *N.vectensis* | 0.7579 | 0.0010 | 0.0026 | MEGAN,Kraken |
|  | V1V2-two step | *A.aurita* | 0.7868 | 0.0001 | 0.0005 | MetaPhlan2 |
|  | V3V4-one step | *N.vectensis* | 0.9222 | 0.0001 | 0.0004 | MEGAN,Kraken |
|  | V3V4-two step | *A.aurita* | 0.9594 | 0.0001 | 0.0005 | MetaPhlan2 |
| *Vibrionaceae* | V1V2-one step | *N.vectensis* | 0.7746 | 0.0040 | 0.0088 |  |
|  | V1V2-two step | *A.aurita* | 0.7071 | 0.0049 | 0.0112 |  |
|  | V3V4-one step | *A.aurita* | 0.9999 | 0.0001 | 0.0004 |  |
|  | V3V4-two step | *A.aurita* | 0.9092 | 0.0002 | 0.0009 |  |
| *Vitellibacter* | V1V2-one step | *N.vectensis* | 1.0000 | 0.0002 | 0.0007 |  |
| *Vogesella* | V1V2-one step | *H.vulgaris* | 0.9802 | 0.0001 | 0.0006 |  |
|  | V1V2-two step |  | 0.9970 | 0.0001 | 0.0005 |  |
|  | V3V4-one step |  | 0.9923 | 0.0001 | 0.0004 |  |
|  | V3V4-two step |  | 0.7071 | 0.0100 | 0.0189 |  |
| *Winogradskyella* | V1V2-one step | *N.vectensis* | 0.7746 | 0.0042 | 0.0091 |  |
|  | V3V4-one step |  | 0.7746 | 0.0048 | 0.0090 |  |
| *Xanthomonadaceae* | V1V2-one step | *D.mel.(feces)* | 0.8283 | 0.0001 | 0.0006 |  |
|  | V1V2-two step |  | 0.7400 | 0.0056 | 0.0124 |  |
