## Supplemental Table S8 for "Comparative analysis of amplicon and metagenomic sequencing methods reveals key features in the evolution of animal metaorganisms"

| Genus | Classifier | Association | IndVal.g | *P* | *P*_FDR_ | Overlap amplicon results |
| --- | --- | --- | --- | --- | --- | --- |
| *Acetobacter* | MEGAN | *D.mel (gut)* | 0.7060 | 0.0041 | 0.0050 | V1V2-one step |
|  | MetaPhlan |  | 0.6324 | 0.0117 | 0.0143 | V1V2-two step |
|  | MetaPhlan2 |  | 0.6325 | 0.0122 | 0.0174 | V3V4-one step |
|  | Kraken |  | 0.7023 | 0.0314 | 0.0314 | V3V4-two step |
| *Aeromonas* | MEGAN | *H.vulgaris* | 0.9850 | 0.0001 | 0.0002 | V1V2-one step |
|  | MetaPhlan |  | 1.0000 | 0.0001 | 0.0003 | V1V2-two step |
|  | MetaPhlan2 |  | 1.0000 | 0.0001 | 0.0002 | V3V4-one step |
|  | Kraken |  | 0.9859 | 0.0001 | 0.0002 |  |
| *Bacteria uncl.* | SortmeRNA | *D.mel (gut)* | 0.3237 | 0.0077 | 0.0154 |  |
| *Bacteroides* | MEGAN | *H.sapiens* | 0.8673 | 0.0001 | 0.0002 | V1V2-one step |
|  | MetaPhlan |  | 0.8168 | 0.0003 | 0.0005 | V1V2-two step |
|  | MetaPhlan2 |  | 0.9530 | 0.0001 | 0.0002 | V3V4-one step |
|  | Kraken |  | 0.8441 | 0.0001 | 0.0002 | V3V4-two step |
| *Blautia* | MetaPhlan | *M.musculus* | 0.9727 | 0.0001 | 0.0003 |  |
| *Bradyrhizobium* | Kraken | *D.mel (feces)* | 0.9274 | 0.0001 | 0.0002 | V1V2-one step  V1V2-two step  V3V4-one step  V3V4-two step |
| *Candidatus Carsonella* | MetaPhlan | *T.aestivum* | 0.4570 | 0.0002 | 0.0004 |  |
| *Candidatus Sulcia* | MetaPhlan | *T.aestivum* | 0.4501 | 0.0169 | 0.0186 |  |
| *Candidatus Zinderia* | MetaPhlan | *M.leidyi* | 0.4644 | 0.0001 | 0.0003 |  |
| *Cutibacterium* | Kraken | *M.leidyi* | 0.6996 | 0.0002 | 0.0003 |  |
| *Escherichia* | MetaPhlan | *A.aurita* | 0.7064 | 0.0022 | 0.0030 |  |
|  | MetaPhlan2 | *A.aurita* | 0.7488 | 0.0005 | 0.0010 |  |
|  | MEGAN | *C.elegans* | 0.8717 | 0.0003 | 0.0004 | V1V2-one step,V1V2-two step |
|  | Kraken | *C.elegans* | 0.7583 | 0.0123 | 0.0148 | V3V4-one step,V3V4-two step |
| *Exiguobacterium* | Kraken | *T.aestivum* | 0.9959 | 0.0001 | 0.0002 |  |
| *Lachnoclostridium* | Kraken | *M.musculus* | 0.9105 | 0.0001 | 0.0002 |  |
| *Lactobacillus* | MEGAN | *D.mel (feces)* | 0.7410 | 0.0001 | 0.0002 | V1V2-one step |
|  | MetaPhlan | *D.mel (feces)* | 0.9514 | 0.0001 | 0.0003 | V3V4-one step |
|  | MetaPhlan2 | *D.mel (feces)* | 0.7933 | 0.0001 | 0.0002 | V3V4-two step |
|  | Kraken | *D.mel (gut)* | 0.7551 | 0.0006 | 0.0009 | V1V2-two step |
| *Mucispirillum* | MetaPhlan2 | *M.musculus* | 0.8944 | 0.0001 | 0.0002 | V1V2-one step  V1V2-two step  V3V4-one step  V3V4-two step |
| *Pantoea* | MEGAN | *T.aestivum* | 0.9978 | 0.0001 | 0.0002 | V1V2-one step  V1V2-two step |
| *Paracoccus* | MEGAN | *M.leidyi* | 0.9282 | 0.0001 | 0.0002 |  |
| *Pseudomonas* | MEGAN | *C.elegans* | 0.8087 | 0.0109 | 0.0120 | V1V2-one step,V1V2-two step |
|  | MetaPhlan | *C.elegans* | 0.6767 | 0.0235 | 0.0235 | V3V4-one step,V3V4-two step |
|  | MetaPhlan2 | *H.vulgaris* | 0.6510 | 0.0049 | 0.0082 |  |
|  | Kraken | *H.vulgaris* | 0.5713 | 0.0211 | 0.0230 |  |
| *Ralstonia* | MEGAN | *T.aestivum* | 0.8855 | 0.0001 | 0.0002 |  |
|  | Kraken |  | 0.8247 | 0.0001 | 0.0002 |  |
| *Sphingobacteriaceae uncl.* | MetaPhlan | *H.vulgaris* | 0.8397 | 0.0003 | 0.0005 |  |
| *unknown/unclassified* | SortmeRNA | *T.aestivum* | 0.3358 | 0.0226 | 0.0226 |  |
| *Vibrio* | MetaPhlan2 | *A.aurita* | 0.5675 | 0.0380 | 0.0475 | V1V2-two step,V3V4-two step |
|  | MEGAN | *N.vectensis* | 0.7760 | 0.0001 | 0.0002 | V1V2-one step |
|  | Kraken | *N.vectensis* | 0.7169 | 0.0016 | 0.0021 | V3V4-one step |
