## Supplemental Table S9 for "Comparative analysis of amplicon and metagenomic sequencing methods reveals key features in the evolution of animal metaorganisms"

| Genus | Method | Association | *IndVal.g* | *P* | *P*_FDR_ | Overlap with shotgun results |
| --- | --- | --- | --- | --- | --- | --- |
| *Acetobacteraceae* | V1V2-two step | aquatic | 0.4472 | 0.0121 | 0.0474 |  |
|  | V3V4-two step |  | 0.4993 | 0.0057 | 0.0297 |  |
| *Acidimicrobiales* | V1V2-one step | aquatic | 0.5648 | 0.0079 | 0.0471 |  |
|  | V1V2-two step |  | 0.5643 | 0.0044 | 0.0261 |  |
|  | V3V4-one step |  | 0.7211 | 0.0001 | 0.0022 |  |
|  | V3V4-two step |  | 0.6770 | 0.0001 | 0.0015 |  |
| *Acidovorax* | V3V4-one step | aquatic | 0.7119 | 0.0003 | 0.0043 |  |
| *Actinobacteria* | V1V2-one step | aquatic | 0.5640 | 0.0026 | 0.0219 |  |
|  | V1V2-two step |  | 0.5919 | 0.0011 | 0.0112 |  |
| *Actinobacteria.1* | V1V2-one step | aquatic | 0.7724 | 0.0002 | 0.0033 |  |
|  | V1V2-two step |  | 0.7483 | 0.0001 | 0.0020 |  |
|  | V3V4-one step |  | 0.7211 | 0.0001 | 0.0022 |  |
|  | V3V4-two step |  | 0.7360 | 0.0001 | 0.0015 |  |
| *Actinomycetales* | V1V2-two step | aquatic | 0.6473 | 0.0056 | 0.0289 |  |
|  | V3V4-one step |  | 0.6618 | 0.0008 | 0.0086 |  |
|  | V3V4-two step |  | 0.7055 | 0.0003 | 0.0036 |  |
| *Aeromonas* | V1V2-one step | aquatic | 0.7463 | 0.0032 | 0.0233 | Kraken |
|  | V1V2-two step |  | 0.6873 | 0.0005 | 0.0066 | MEGAN |
|  | V3V4-one step |  | 0.6590 | 0.0003 | 0.0043 | MetaPhlan2 |
|  | V3V4-two step |  | 0.5774 | 0.0002 | 0.0025 |  |
| *Aestuariispira* | V3V4-one step | aquatic | 0.6000 | 0.0003 | 0.0043 |  |
| *Alcaligenaceae* | V3V4-one step | aquatic | 0.5657 | 0.0011 | 0.0096 |  |
| *Alcanivorax* | V1V2-two step | aquatic | 0.4472 | 0.0107 | 0.0455 |  |
|  | V3V4-one step |  | 0.4472 | 0.0176 | 0.0486 |  |
| *Aliivibrio* | V1V2-two step | aquatic | 0.4472 | 0.0122 | 0.0474 |  |
| *Alistipes* | V1V2-two step | terrestrial | 0.7323 | 0.0035 | 0.0252 |  |
|  | V3V4-one step |  | 0.5871 | 0.0033 | 0.0179 |  |
|  | V3V4-two step |  | 0.6030 | 0.0023 | 0.0155 |  |
| *Alphaproteobacteria* | V1V2-one step | aquatic | 0.9375 | 0.0001 | 0.0022 |  |
|  | V1V2-two step |  | 0.9376 | 0.0001 | 0.0020 |  |
|  | V3V4-one step |  | 0.9990 | 0.0001 | 0.0022 |  |
|  | V3V4-two step |  | 0.8411 | 0.0001 | 0.0015 |  |
| *Alteromonadaceae* | V1V2-two step | aquatic | 0.4899 | 0.0043 | 0.0261 |  |
| *Alteromonas* | V1V2-one step | aquatic | 0.7201 | 0.0001 | 0.0022 |  |
|  | V1V2-two step |  | 0.7480 | 0.0001 | 0.0020 |  |
|  | V3V4-one step |  | 0.6317 | 0.0013 | 0.0104 |  |
|  | V3V4-two step |  | 0.5000 | 0.0038 | 0.0214 |  |
| *Amphritea* | V3V4-one step | aquatic | 0.5292 | 0.0037 | 0.0195 |  |
|  | V3V4-two step |  | 0.5401 | 0.0013 | 0.0116 |  |
| *Arcobacter* | V1V2-one step | aquatic | 0.5657 | 0.0015 | 0.0150 |  |
|  | V1V2-two step |  | 0.6000 | 0.0006 | 0.0073 |  |
|  | V3V4-one step |  | 0.4899 | 0.0075 | 0.0311 |  |
|  | V3V4-two step |  | 0.6455 | 0.0001 | 0.0015 |  |
| *Aureispira* | V1V2-two step | aquatic | 0.4472 | 0.0107 | 0.0455 |  |
|  | V3V4-one step |  | 0.4472 | 0.0176 | 0.0486 |  |
| *Bacillus* | V3V4-one step | terrestrial | 0.5821 | 0.0040 | 0.0205 |  |
| *Bacteria* | V1V2-one step | aquatic | 0.9442 | 0.0001 | 0.0022 |  |
|  | V1V2-two step |  | 0.9463 | 0.0001 | 0.0020 |  |
|  | V3V4-one step |  | 0.9332 | 0.0001 | 0.0022 |  |
|  | V3V4-two step |  | 0.8092 | 0.0002 | 0.0025 |  |
| *Bacteriovoracaceae* | V3V4-one step | aquatic | 0.5292 | 0.0032 | 0.0179 |  |
| *Bacteroidales* | V1V2-one step | terrestrial | 0.6427 | 0.0018 | 0.0164 |  |
|  | V1V2-two step |  | 0.6004 | 0.0105 | 0.0455 |  |
| *Bacteroides* | V1V2-two step | terrestrial | 0.8530 | 0.0026 | 0.0210 |  |
|  | V3V4-one step |  | 0.7176 | 0.0067 | 0.0304 |  |
|  | V3V4-two step |  | 0.6273 | 0.0026 | 0.0171 |  |
| *Bacteroidetes* | V3V4-one step | aquatic | 0.9158 | 0.0001 | 0.0022 |  |
| *Barnesiella* | V3V4-one step | terrestrial | 0.5872 | 0.0023 | 0.0154 |  |
|  | V3V4-two step |  | 0.5774 | 0.0032 | 0.0196 |  |
| *Bdellovibrio* | V3V4-one step | aquatic | 0.4472 | 0.0179 | 0.0486 |  |
| *Betaproteobacteria* | V1V2-one step | aquatic | 0.7542 | 0.0002 | 0.0033 |  |
|  | V1V2-two step |  | 0.7182 | 0.0001 | 0.0020 |  |
| *Brevundimonas* | V3V4-one step | terrestrial | 0.4913 | 0.0108 | 0.0405 |  |
| *Brumimicrobium* | V3V4-one step | aquatic | 0.4899 | 0.0065 | 0.0301 |  |
| *Burkholderiales* | V1V2-one step | aquatic | 0.6325 | 0.0002 | 0.0033 |  |
|  | V1V2-two step |  | 0.5962 | 0.0005 | 0.0066 |  |
| *Butyricicoccus* | V1V2-one step | terrestrial | 0.5571 | 0.0043 | 0.0282 |  |
|  | V1V2-two step |  | 0.5774 | 0.0032 | 0.0240 |  |
|  | V3V4-one step |  | 0.6159 | 0.0010 | 0.0091 |  |
|  | V3V4-two step |  | 0.6030 | 0.0022 | 0.0152 |  |
| *Caldilineaceae* | V1V2-two step | aquatic | 0.4472 | 0.0120 | 0.0474 |  |
|  | V3V4-one step |  | 0.4472 | 0.0165 | 0.0486 |  |
|  | V3V4-two step |  | 0.4564 | 0.0105 | 0.0426 |  |
| *Campylobacterales* | V3V4-two step | aquatic | 0.4564 | 0.0098 | 0.0408 |  |
| *Candidatus Pelagibacter* | V1V2-two step | aquatic | 0.5657 | 0.0008 | 0.0093 |  |
|  | V3V4-one step |  | 0.6000 | 0.0009 | 0.0089 |  |
|  | V3V4-two step |  | 0.6455 | 0.0001 | 0.0015 |  |
| *Castellaniella* | V3V4-one step | aquatic | 0.4472 | 0.0176 | 0.0486 |  |
| *Chitinophagaceae* | V1V2-one step | aquatic | 0.7113 | 0.0004 | 0.0062 |  |
|  | V1V2-two step |  | 0.6721 | 0.0013 | 0.0124 |  |
|  | V3V4-one step |  | 0.5801 | 0.0164 | 0.0486 |  |
| *Chlamydiales* | V3V4-one step | aquatic | 0.6614 | 0.0001 | 0.0022 |  |
| *Chloroflexi* | V3V4-one step | aquatic | 0.4472 | 0.0165 | 0.0486 |  |
|  | V3V4-two step |  | 0.5774 | 0.0005 | 0.0055 |  |
| *Chromatiales* | V3V4-one step |  | 0.4472 | 0.0165 | 0.0486 |  |
| *Chryseomicrobium* | V3V4-one step | aquatic | 0.4899 | 0.0068 | 0.0304 |  |
| *Clostridia* | V3V4-one step | terrestrial | 0.4913 | 0.0112 | 0.0412 |  |
| *Clostridiales* | V1V2-one step | terrestrial | 0.6941 | 0.0016 | 0.0150 |  |
|  | V1V2-two step |  | 0.7584 | 0.0010 | 0.0105 |  |
| *Clostridium IV* | V3V4-one step | terrestrial | 0.5872 | 0.0019 | 0.0134 |  |
| *Clostridium XlVa* | V1V2-one step | terrestrial | 0.5872 | 0.0019 | 0.0168 |  |
|  | V1V2-two step |  | 0.6940 | 0.0021 | 0.0184 |  |
|  | V3V4-one step |  | 0.6693 | 0.0008 | 0.0086 |  |
|  | V3V4-two step |  | 0.6030 | 0.0022 | 0.0152 |  |
| *Clostridium XlVb* | V1V2-two step | terrestrial | 0.5505 | 0.0054 | 0.0283 |  |
|  | V3V4-one step |  | 0.5571 | 0.0061 | 0.0295 |  |
|  | V3V4-two step |  | 0.5222 | 0.0115 | 0.0453 |  |
| *Colwelliaceae* | V3V4-one step | aquatic | 0.5292 | 0.0033 | 0.0179 |  |
| *Comamonadaceae* | V1V2-one step | aquatic | 0.6626 | 0.0001 | 0.0022 |  |
|  | V1V2-two step |  | 0.5641 | 0.0010 | 0.0105 |  |
|  | V3V4-one step |  | 0.7994 | 0.0002 | 0.0034 |  |
| *Coriobacteriaceae* | V1V2-two step | terrestrial | 0.5774 | 0.0022 | 0.0187 |  |
|  | V3V4-one step |  | 0.5532 | 0.0060 | 0.0295 |  |
| *Coxiella* | V1V2-one step | aquatic | 0.5978 | 0.0005 | 0.0075 |  |
|  | V1V2-two step |  | 0.5987 | 0.0002 | 0.0035 |  |
|  | V3V4-one step |  | 0.6000 | 0.0001 | 0.0022 |  |
| *Croceibacter* | V1V2-one step | aquatic | 0.4899 | 0.0083 | 0.0478 |  |
| *Crocinitomix* | V3V4-one step | aquatic | 0.5657 | 0.0014 | 0.0107 |  |
|  | V3V4-two step |  | 0.4564 | 0.0098 | 0.0408 |  |
| *Cryomorphaceae* | V1V2-one step | aquatic | 0.5917 | 0.0011 | 0.0124 |  |
|  | V1V2-two step |  | 0.6564 | 0.0001 | 0.0020 |  |
|  | V3V4-one step |  | 0.6928 | 0.0001 | 0.0022 |  |
|  | V3V4-two step |  | 0.6770 | 0.0001 | 0.0015 |  |
| *Curvibacter* | V1V2-one step | aquatic | 0.7207 | 0.0027 | 0.0221 |  |
|  | V1V2-two step |  | 0.7454 | 0.0058 | 0.0295 |  |
|  | V3V4-one step |  | 0.6253 | 0.0104 | 0.0394 |  |
| *Cytophagaceae* | V1V2-one step | aquatic | 0.5961 | 0.0012 | 0.0131 |  |
|  | V3V4-one step |  | 0.4899 | 0.0076 | 0.0311 |  |
| *Cytophagales* | V3V4-one step | aquatic | 0.4899 | 0.0077 | 0.0311 |  |
| *Deltaproteobacteria* | V1V2-one step | aquatic | 0.6880 | 0.0011 | 0.0124 |  |
|  | V3V4-one step |  | 0.6303 | 0.0009 | 0.0089 |  |
|  | V3V4-two step |  | 0.4919 | 0.0085 | 0.0404 |  |
| *Dyadobacter* | V3V4-one step | aquatic | 0.4472 | 0.0169 | 0.0486 |  |
| *Endozoicomonas* | V3V4-one step | aquatic | 0.6325 | 0.0002 | 0.0034 |  |
|  | V3V4-two step |  | 0.5000 | 0.0030 | 0.0188 |  |
| *Enhydrobacter* | V1V2-two step | terrestrial | 0.6288 | 0.0082 | 0.0404 |  |
|  | V3V4-one step |  | 0.5760 | 0.0156 | 0.0486 |  |
| *Enterobacteriaceae* | V3V4-one step | terrestrial | 0.7237 | 0.0088 | 0.0340 |  |
| *Enterovibrio* | V3V4-one step | aquatic | 0.5981 | 0.0008 | 0.0086 |  |
|  | V3V4-two step |  | 0.5401 | 0.0013 | 0.0116 |  |
| *Exiguobacterium* | V1V2-two step | aquatic | 0.5265 | 0.0031 | 0.0238 |  |
|  | V3V4-one step |  | 0.4895 | 0.0160 | 0.0486 |  |
| *Filomicrobium* | V3V4-one step | aquatic | 0.4472 | 0.0167 | 0.0486 |  |
| *Flaviramulus* | V3V4-one step | aquatic | 0.4899 | 0.0078 | 0.0311 |  |
| *Flavobacteriaceae* | V1V2-one step | aquatic | 0.9117 | 0.0001 | 0.0022 |  |
|  | V1V2-two step |  | 0.8649 | 0.0001 | 0.0020 |  |
|  | V3V4-one step |  | 0.8687 | 0.0001 | 0.0022 |  |
|  | V3V4-two step |  | 0.7621 | 0.0001 | 0.0015 |  |
| *Flavobacteriales* | V1V2-one step | aquatic | 0.8718 | 0.0001 | 0.0022 |  |
|  | V1V2-two step |  | 0.8242 | 0.0001 | 0.0020 |  |
|  | V3V4-one step |  | 0.7201 | 0.0001 | 0.0022 |  |
|  | V3V4-two step |  | 0.5774 | 0.0008 | 0.0082 |  |
| *Flavobacterium* | V1V2-one step | aquatic | 0.7685 | 0.0002 | 0.0033 |  |
|  | V1V2-two step |  | 0.7419 | 0.0001 | 0.0020 |  |
|  | V3V4-one step |  | 0.8636 | 0.0008 | 0.0086 |  |
|  | V3V4-two step |  | 0.7344 | 0.0001 | 0.0015 |  |
| *Flavonifractor* | V3V4-one step | terrestrial | 0.4913 | 0.0119 | 0.0433 |  |
| *Fluviicola* | V1V2-two step | aquatic | 0.4899 | 0.0039 | 0.0256 |  |
| *Francisella* | V1V2-two step | aquatic | 0.5291 | 0.0036 | 0.0252 |  |
| *Gammaproteobacteria* | V1V2-one step | aquatic | 0.9991 | 0.0001 | 0.0022 |  |
|  | V1V2-two step |  | 0.9570 | 0.0001 | 0.0020 |  |
|  | V3V4-one step |  | 0.9988 | 0.0001 | 0.0022 |  |
|  | V3V4-two step |  | 0.8372 | 0.0001 | 0.0015 |  |
| *Gemmobacter* | V3V4-one step | aquatic | 0.5657 | 0.0010 | 0.0091 |  |
| *Gp21* | V3V4-one step | aquatic | 0.4472 | 0.0165 | 0.0486 |  |
| *Gp3* | V1V2-two step | aquatic | 0.4472 | 0.0098 | 0.0455 |  |
| *Gp5* | V3V4-one step | aquatic | 0.4472 | 0.0165 | 0.0486 |  |
| *Gp6* | V1V2-one step | aquatic | 0.5873 | 0.0034 | 0.0237 |  |
|  | V1V2-two step |  | 0.5205 | 0.0030 | 0.0236 |  |
| *Gp9* | V1V2-two step | aquatic | 0.4472 | 0.0098 | 0.0455 |  |
|  | V3V4-one step |  | 0.4472 | 0.0165 | 0.0486 |  |
| *Halioglobus* | V3V4-one step | aquatic | 0.4845 | 0.0151 | 0.0486 |  |
|  | V3V4-two step |  | 0.4564 | 0.0117 | 0.0455 |  |
| *Halobacteriovorax* | V3V4-one step | aquatic | 0.6633 | 0.0001 | 0.0022 |  |
| *Hathewaya* | V1V2-two step | aquatic | 0.5292 | 0.0020 | 0.0180 |  |
|  | V3V4-one step |  | 0.4472 | 0.0175 | 0.0486 |  |
|  | V3V4-two step |  | 0.5000 | 0.0029 | 0.0186 |  |
| *Hoeflea* | V3V4-one step | aquatic | 0.5641 | 0.0036 | 0.0193 |  |
| *Hydrogenophaga* | V1V2-one step | aquatic | 0.4899 | 0.0060 | 0.0386 |  |
| *Hyphomicrobium* | V3V4-one step | aquatic | 0.5631 | 0.0032 | 0.0179 |  |
| *Hyphomonadaceae* | V1V2-one step | aquatic | 0.4899 | 0.0072 | 0.0443 |  |
|  | V1V2-two step |  | 0.4899 | 0.0041 | 0.0261 |  |
|  | V3V4-one step |  | 0.4472 | 0.0176 | 0.0486 |  |
| *Hyphomonas* | V3V4-one step | aquatic | 0.5292 | 0.0032 | 0.0179 |  |
| *Iamiaceae* | V1V2-two step | aquatic | 0.4899 | 0.0047 | 0.0261 |  |
| *Ilumatobacter* | V1V2-one step | aquatic | 0.5292 | 0.0028 | 0.0224 |  |
|  | V1V2-two step |  | 0.6000 | 0.0003 | 0.0047 |  |
|  | V3V4-one step |  | 0.6172 | 0.0054 | 0.0274 |  |
|  | V3V4-two step |  | 0.6768 | 0.0001 | 0.0015 |  |
| *Intestinimonas* | V3V4-one step | terrestrial | 0.4913 | 0.0111 | 0.0412 |  |
|  | V3V4-two step |  | 0.5222 | 0.0085 | 0.0404 |  |
| *Jannaschia* | V3V4-one step | aquatic | 0.4472 | 0.0157 | 0.0486 |  |
| *Kiloniella* | V1V2-one step | aquatic | 0.5292 | 0.0035 | 0.0239 |  |
|  | V1V2-two step |  | 0.5288 | 0.0065 | 0.0325 |  |
| *Lachnospiracea incertae sedis* | V1V2-two step | terrestrial | 0.5222 | 0.0092 | 0.0446 |  |
| *Lachnospiraceae* | V1V2-one step | terrestrial | 0.7862 | 0.0011 | 0.0124 |  |
|  | V1V2-two step |  | 0.6950 | 0.0036 | 0.0252 |  |
|  | V3V4-one step |  | 0.8293 | 0.0015 | 0.0112 |  |
|  | V3V4-two step |  | 0.6023 | 0.0083 | 0.0404 |  |
| *Lactobacillus* | V1V2-one step | terrestrial | 0.8297 | 0.0030 | 0.0229 | Kraken |
|  | V1V2-two step |  | 0.7969 | 0.0046 | 0.0261 | MEGAN |
|  | V3V4-one step |  | 0.7425 | 0.0032 | 0.0179 | MetaPhlan |
|  | V3V4-two step |  | 0.7780 | 0.0004 | 0.0046 | MetaPhlan2 |
| *Legionella* | V1V2-one step | aquatic | 0.8235 | 0.0001 | 0.0022 |  |
|  | V1V2-two step |  | 0.6925 | 0.0001 | 0.0020 |  |
|  | V3V4-one step |  | 0.8000 | 0.0001 | 0.0022 |  |
|  | V3V4-two step |  | 0.6122 | 0.0001 | 0.0015 |  |
| *Lewinella* | V3V4-one step | aquatic | 0.4899 | 0.0074 | 0.0311 |  |
| *Litoreibacter* | V3V4-one step | aquatic | 0.7209 | 0.0006 | 0.0078 |  |
|  | V3V4-two step |  | 0.6770 | 0.0001 | 0.0015 |  |
| *Litorilinea* | V3V4-one step | aquatic | 0.5657 | 0.0007 | 0.0085 |  |
|  | V3V4-two step |  | 0.5401 | 0.0016 | 0.0130 |  |
| *Loktanella* | V3V4-one step | aquatic | 0.6928 | 0.0001 | 0.0022 |  |
| *Luteolibacter* | V3V4-one step | aquatic | 0.4899 | 0.0070 | 0.0309 |  |
| *Mariniflexile* | V1V2-two step | aquatic | 0.4899 | 0.0039 | 0.0256 |  |
|  | V3V4-one step |  | 0.4472 | 0.0176 | 0.0486 |  |
| *Marinobacter* | V1V2-one step | aquatic | 0.4899 | 0.0073 | 0.0443 |  |
|  | V3V4-one step |  | 0.6325 | 0.0002 | 0.0034 |  |
|  | V3V4-two step |  | 0.4551 | 0.0097 | 0.0408 |  |
| *Marinobacterium* | V3V4-two step | aquatic | 0.4564 | 0.0097 | 0.0408 |  |
| *Marinomonas* | V1V2-one step | aquatic | 0.5657 | 0.0016 | 0.0150 |  |
|  | V3V4-one step |  | 0.6304 | 0.0012 | 0.0098 |  |
| *Maritalea* | V1V2-one step | aquatic | 0.5292 | 0.0033 | 0.0235 |  |
|  | V1V2-two step |  | 0.4472 | 0.0107 | 0.0455 |  |
|  | V3V4-one step |  | 0.4472 | 0.0176 | 0.0486 |  |
| *Marivita* | V3V4-one step | aquatic | 0.6928 | 0.0001 | 0.0022 |  |
|  | V3V4-two step |  | 0.5401 | 0.0019 | 0.0142 |  |
| *Mesonia* | V3V4-one step | aquatic | 0.4472 | 0.0176 | 0.0486 |  |
| *Methylocystis* | V3V4-one step | aquatic | 0.4472 | 0.0175 | 0.0486 |  |
| *Methyloparacoccus* | V1V2-two step | aquatic | 0.4472 | 0.0120 | 0.0474 |  |
|  | V3V4-two step |  | 0.4564 | 0.0080 | 0.0401 |  |
| *Methylophaga* | V1V2-two step | aquatic | 0.4899 | 0.0043 | 0.0261 |  |
| *Methylophilaceae* | V1V2-two step | aquatic | 0.4472 | 0.0111 | 0.0464 |  |
|  | V3V4-two step |  | 0.5401 | 0.0019 | 0.0142 |  |
| *Methylophilus* | V3V4-one step | aquatic | 0.4899 | 0.0082 | 0.0324 |  |
| *Methylotenera* | V3V4-one step | aquatic | 0.4472 | 0.0176 | 0.0486 |  |
| *Microbacteriaceae* | V1V2-one step | aquatic | 0.6534 | 0.0020 | 0.0173 |  |
|  | V1V2-two step |  | 0.5971 | 0.0018 | 0.0167 |  |
|  | V3V4-one step |  | 0.7672 | 0.0002 | 0.0034 |  |
|  | V3V4-two step |  | 0.6728 | 0.0001 | 0.0015 |  |
| *Mycobacterium* | V1V2-one step | aquatic | 0.6525 | 0.0016 | 0.0150 |  |
|  | V1V2-two step |  | 0.6633 | 0.0001 | 0.0020 |  |
|  | V3V4-one step |  | 0.4899 | 0.0063 | 0.0295 |  |
| *Mycoplasma* | V3V4-one step | aquatic | 0.6920 | 0.0024 | 0.0155 |  |
|  | V3V4-two step |  | 0.6455 | 0.0001 | 0.0015 |  |
| *Nitratireductor* | V3V4-one step | aquatic | 0.5657 | 0.0014 | 0.0107 |  |
| *Nitrosomonas* | V1V2-two step | aquatic | 0.4472 | 0.0107 | 0.0455 |  |
|  | V3V4-one step |  | 0.4472 | 0.0176 | 0.0486 |  |
| *Nitrospira* | V3V4-one step | aquatic | 0.4472 | 0.0165 | 0.0486 |  |
| *Oceanisphaera* | V3V4-two step | aquatic | 0.4564 | 0.0099 | 0.0408 |  |
| *Oceanospirillaceae* | V1V2-two step | aquatic | 0.4472 | 0.0102 | 0.0455 |  |
| *Oceanospirillales* | V1V2-one step | aquatic | 0.6633 | 0.0001 | 0.0022 |  |
|  | V1V2-two step |  | 0.5657 | 0.0009 | 0.0101 |  |
| *Odoribacter* | V1V2-one step | terrestrial | 0.5571 | 0.0036 | 0.0241 |  |
|  | V3V4-one step |  | 0.5872 | 0.0018 | 0.0132 |  |
|  | V3V4-two step |  | 0.6030 | 0.0013 | 0.0116 |  |
| *Opitutae* | V3V4-one step | aquatic | 0.5657 | 0.0012 | 0.0098 |  |
| *Oscillibacter* | V1V2-one step | terrestrial | 0.6159 | 0.0009 | 0.0114 |  |
|  | V1V2-two step |  | 0.6470 | 0.0012 | 0.0118 |  |
|  | V3V4-one step |  | 0.5847 | 0.0024 | 0.0155 |  |
|  | V3V4-two step |  | 0.6028 | 0.0021 | 0.0152 |  |
| *Parachlamydiaceae* | V3V4-one step | aquatic | 0.5657 | 0.0007 | 0.0085 |  |
| *Parasutterella* | V1V2-two step | terrestrial | 0.5771 | 0.0103 | 0.0455 |  |
| *Pedobacter* | V3V4-one step | aquatic | 0.5253 | 0.0063 | 0.0295 |  |
| *Peredibacter* | V3V4-one step | aquatic | 0.4899 | 0.0072 | 0.0311 |  |
| *Phaeodactylibacter* | V1V2-one step | aquatic | 0.4899 | 0.0083 | 0.0478 |  |
|  | V1V2-two step |  | 0.4472 | 0.0112 | 0.0464 |  |
| *Phenylobacterium* | V3V4-one step | terrestrial | 0.4913 | 0.0139 | 0.0486 |  |
| *Photobacterium* | V3V4-two step | aquatic | 0.4564 | 0.0098 | 0.0408 |  |
| *Phycisphaera* | V1V2-two step | aquatic | 0.4899 | 0.0038 | 0.0256 |  |
| *Planctomycetaceae* | V1V2-one step | aquatic | 0.4899 | 0.0073 | 0.0443 |  |
|  | V1V2-two step |  | 0.4899 | 0.0048 | 0.0261 |  |
|  | V3V4-one step |  | 0.4899 | 0.0073 | 0.0311 |  |
| *Planktomarina* | V3V4-one step | aquatic | 0.5624 | 0.0092 | 0.0352 |  |
|  | V3V4-two step |  | 0.6455 | 0.0001 | 0.0015 |  |
| *Polaribacter* | V1V2-one step | aquatic | 0.6633 | 0.0001 | 0.0022 |  |
|  | V1V2-two step |  | 0.5657 | 0.0005 | 0.0066 |  |
|  | V3V4-one step |  | 0.5657 | 0.0009 | 0.0089 |  |
| *Poribacteria genera incertae sedis* | V3V4-one step | aquatic | 0.4472 | 0.0165 | 0.0486 |  |
| *Porphyromonadaceae* | V3V4-one step | terrestrial | 0.6645 | 0.0063 | 0.0295 |  |
|  | V3V4-two step |  | 0.5222 | 0.0076 | 0.0388 |  |
| *Prevotellaceae* | V1V2-two step | terrestrial | 0.6264 | 0.0121 | 0.0474 |  |
| *Proteobacteria* | V1V2-one step | aquatic | 0.9387 | 0.0001 | 0.0022 |  |
|  | V1V2-two step |  | 0.9267 | 0.0001 | 0.0020 |  |
|  | V3V4-one step |  | 0.8659 | 0.0005 | 0.0067 |  |
|  | V3V4-two step |  | 0.7356 | 0.0001 | 0.0015 |  |
| *Pseudoalteromonas* | V1V2-one step | aquatic | 0.7746 | 0.0001 | 0.0022 |  |
|  | V1V2-two step |  | 0.7951 | 0.0001 | 0.0020 |  |
|  | V3V4-one step |  | 0.7739 | 0.0001 | 0.0022 |  |
|  | V3V4-two step |  | 0.7066 | 0.0001 | 0.0015 |  |
| *Reyranella* | V1V2-one step | aquatic | 0.5278 | 0.0031 | 0.0231 |  |
|  | V3V4-one step |  | 0.5975 | 0.0019 | 0.0134 |  |
| *Rheinheimera* | V1V2-two step | aquatic | 0.4899 | 0.0053 | 0.0283 |  |
| *Rhizobiaceae* | V3V4-one step | aquatic | 0.4472 | 0.0179 | 0.0486 |  |
| *Rhizobiales* | V1V2-two step | aquatic | 0.4472 | 0.0128 | 0.0490 |  |
| *Rhodobacteraceae* | V1V2-one step | aquatic | 0.7456 | 0.0002 | 0.0033 |  |
|  | V1V2-two step |  | 0.7746 | 0.0001 | 0.0020 |  |
|  | V3V4-one step |  | 0.9710 | 0.0001 | 0.0022 |  |
|  | V3V4-two step |  | 0.7566 | 0.0001 | 0.0015 |  |
| *Rhodoferax* | V1V2-one step | aquatic | 0.5990 | 0.0015 | 0.0150 |  |
|  | V3V4-one step |  | 0.4899 | 0.0078 | 0.0311 |  |
| *Rhodoluna* | V1V2-two step | aquatic | 0.4472 | 0.0131 | 0.0491 |  |
| *Rhodospirillaceae* | V3V4-one step | aquatic | 0.7337 | 0.0004 | 0.0056 |  |
| *Rhodospirillales* | V3V4-one step | aquatic | 0.5292 | 0.0029 | 0.0175 |  |
|  | V3V4-two step |  | 0.5000 | 0.0043 | 0.0228 |  |
| *Rhodothermaceae* | V3V4-one step | aquatic | 0.4472 | 0.0165 | 0.0486 |  |
| *Rickettsiaceae* | V1V2-one step | aquatic | 0.6000 | 0.0001 | 0.0022 |  |
|  | V1V2-two step |  | 0.6000 | 0.0006 | 0.0073 |  |
| *Roseibacillus* | V3V4-one step | aquatic | 0.5657 | 0.0022 | 0.0150 |  |
|  | V3V4-two step |  | 0.5000 | 0.0038 | 0.0214 |  |
| *Roseovarius* | V3V4-one step | aquatic | 0.5657 | 0.0012 | 0.0098 |  |
| *Ruegeria* | V3V4-one step | aquatic | 0.4899 | 0.0061 | 0.0295 |  |
|  | V3V4-two step |  | 0.4564 | 0.0108 | 0.0432 |  |
| *Ruminococcaceae* | V1V2-one step | terrestrial | 0.7638 | 0.0001 | 0.0022 |  |
|  | V1V2-two step |  | 0.7521 | 0.0023 | 0.0191 |  |
|  | V3V4-one step |  | 0.6422 | 0.0039 | 0.0203 |  |
|  | V3V4-two step |  | 0.6244 | 0.0092 | 0.0408 |  |
| *Saccharibacteria genera incertae sedis* | V3V4-one step | aquatic | 0.7375 | 0.0068 | 0.0304 |  |
|  | V3V4-two step |  | 0.6124 | 0.0087 | 0.0407 |  |
| *Salinirepens* | V1V2-one step | aquatic | 0.5657 | 0.0007 | 0.0096 |  |
|  | V1V2-two step |  | 0.6000 | 0.0004 | 0.0060 |  |
|  | V3V4-one step |  | 0.5982 | 0.0029 | 0.0175 |  |
|  | V3V4-two step |  | 0.5401 | 0.0017 | 0.0134 |  |
| *Saprospiraceae* | V1V2-one step | aquatic | 0.6000 | 0.0006 | 0.0086 |  |
|  | V1V2-two step |  | 0.6000 | 0.0003 | 0.0047 |  |
|  | V3V4-one step |  | 0.5966 | 0.0011 | 0.0096 |  |
|  | V3V4-two step |  | 0.5401 | 0.0016 | 0.0130 |  |
| *Shinella* | V3V4-one step | aquatic | 0.4897 | 0.0169 | 0.0486 |  |
| *Spartobacteria* | V3V4-one step | aquatic | 0.4899 | 0.0085 | 0.0332 |  |
|  | V3V4-two step |  | 0.5000 | 0.0034 | 0.0200 |  |
| *Spartobacteria genera incertae sedis* | V1V2-two step | aquatic | 0.4899 | 0.0048 | 0.0261 |  |
|  | V3V4-one step |  | 0.4898 | 0.0185 | 0.0499 |  |
|  | V3V4-two step |  | 0.5401 | 0.0012 | 0.0116 |  |
| *Sphaerotilus* | V3V4-one step | aquatic | 0.4472 | 0.0179 | 0.0486 |  |
| *Sphingomonadaceae* | V3V4-one step | aquatic | 0.6012 | 0.0010 | 0.0091 |  |
| *Spirochaetaceae* | V1V2-one step | aquatic | 0.6316 | 0.0029 | 0.0226 |  |
|  | V1V2-two step |  | 0.6631 | 0.0002 | 0.0035 |  |
| *Spiroplasma* | V3V4-two step | aquatic | 0.5000 | 0.0034 | 0.0200 |  |
| *Sulfitobacter* | V3V4-one step | aquatic | 0.7196 | 0.0002 | 0.0034 |  |
|  | V3V4-two step |  | 0.5000 | 0.0043 | 0.0228 |  |
| *Sulfurimonas* | V1V2-two step | aquatic | 0.4899 | 0.0048 | 0.0261 |  |
|  | V3V4-one step |  | 0.4472 | 0.0175 | 0.0486 |  |
|  | V3V4-two step |  | 0.5401 | 0.0016 | 0.0130 |  |
| *Sulfurovum* | V1V2-two step | aquatic | 0.4899 | 0.0048 | 0.0261 |  |
|  | V3V4-one step |  | 0.4472 | 0.0176 | 0.0486 |  |
|  | V3V4-two step |  | 0.5401 | 0.0008 | 0.0082 |  |
| *Thalassobaculum* | V3V4-one step | aquatic | 0.5292 | 0.0033 | 0.0179 |  |
| *Thalassolituus* | V3V4-one step | aquatic | 0.4896 | 0.0157 | 0.0486 |  |
| *Thalassotalea* | V3V4-one step | aquatic | 0.5292 | 0.0026 | 0.0162 |  |
| *Thiotrichales* | V1V2-two step | aquatic | 0.4472 | 0.0129 | 0.0490 |  |
|  | V3V4-two step |  | 0.4564 | 0.0097 | 0.0408 |  |
| *Undibacterium* | V3V4-one step | aquatic | 0.5978 | 0.0021 | 0.0146 |  |
| *Verrucomicrobiaceae* | V3V4-two step | aquatic | 0.5000 | 0.0042 | 0.0228 |  |
| *Vibrio* | V1V2-one step | aquatic | 0.7998 | 0.0001 | 0.0022 | Kraken |
|  | V1V2-two step |  | 0.8481 | 0.0001 | 0.0020 | MEGAN |
|  | V3V4-one step |  | 0.7745 | 0.0003 | 0.0043 | MetaPhlan2 |
|  | V3V4-two step |  | 0.7071 | 0.0001 | 0.0015 |  |
| *Vibrionaceae* | V3V4-one step | aquatic | 0.5292 | 0.0025 | 0.0159 |  |
|  | V3V4-two step |  | 0.6124 | 0.0002 | 0.0025 |  |
| *Vogesella* | V1V2-one step | aquatic | 0.5657 | 0.0009 | 0.0114 |  |
|  | V3V4-one step |  | 0.4899 | 0.0073 | 0.0311 |  |
