## Supplemental Table S10 for "Comparative analysis of amplicon and metagenomic sequencing methods reveals key features in the evolution of animal metaorganisms"

| Genus | Method | Association | *IndVal.g* | *P* | *P*_FDR_ | Overlap with amplicon results |
| --- | --- | --- | --- | --- | --- | --- |
| *Aeromonas* | Kraken | aquatic | 0.7984 | 0.0056 | 0.0224 | V1V2-one step,V1V2-two step |
|  | MEGAN |  | 0.7746 | 0.0001 | 0.0004 | V3V4-one step |
|  | MetaPhlan2 |  | 0.4472 | 0.0101 | 0.0337 | V3V4-two step |
| *Lactobacillus* | Kraken | terrestrial | 0.8276 | 0.0014 | 0.0084 | V1V2-one step |
|  | MEGAN |  | 0.8757 | 0.0001 | 0.0004 | V1V2-two step |
|  | MetaPhlan |  | 0.6325 | 0.0011 | 0.0121 | V3V4-one step |
|  | MetaPhlan2 |  | 0.6761 | 0.0006 | 0.0045 | V3V4-two step |
| *Paracoccus* | MEGAN | aquatic | 0.7869 | 0.0005 | 0.0014 |  |
| *Vibrio* | Kraken | aquatic | 0.8940 | 0.0001 | 0.0012 | V1V2-one step,V1V2-two step |
|  | MEGAN |  | 0.9164 | 0.0001 | 0.0004 | V3V4-one step |
|  | MetaPhlan2 |  | 0.5292 | 0.0009 | 0.0045 | V3V4-two step |
